# Exact linear theory of perturbation response in a space- and feature-dependent cortical circuit model

**DOI:** 10.1101/2024.12.27.630558

**Authors:** Ho Yin Chau, Kenneth D. Miller, Agostina Palmigiano

**Affiliations:** Center for Theoretical Neuroscience, College of Physicians and Surgeons and Mortimer B. Zuckerman Mind Brain Behavior Institute, Columbia University, New York, NY; Dept. of Neuroscience, Swartz Program in Theoretical Neuroscience, Kavli Institute for Brain Science, College of Physicians and Surgeons and Mortimer B. Zuckerman Mind Brain Behavior Institute, Columbia University, New York; Gatsby Computational Neuroscience Unit, University College London

**Keywords:** Recurrent neural networks, Optogenetic perturbation, Mouse Primary Visual Cortex

## Abstract

What are the principles that govern the responses of cortical networks to their inputs and the emergence of these responses from recurrent connectivity? Recent experiments have probed these questions by measuring cortical responses to two-photon optogenetic perturbations of single cells in the mouse primary visual cortex. A robust theoretical framework is needed to determine the implications of these responses for cortical recurrence. Here we propose a novel analytical approach: a formulation of the dependence of cell-type-specific connectivity on spatial distance that yields an exact solution for the linear perturbation response of a model with multiple cell types and space- and feature-dependent connectivity. Importantly and unlike previous approaches, the solution is valid in regimes of strong as well as weak intra-cortical coupling. Analysis reveals the structure of connectivity implied by various features of single-cell perturbation responses, such as the surprisingly narrow spatial radius of nearby excitation beyond which inhibition dominates, the number of transitions between mean excitation and inhibition thereafter, and the dependence of these responses on feature preferences. Comparison of these results to existing optogenetic perturbation data yields constraints on cell-type-specific connection strengths and their tuning dependence. Finally, we provide experimental predictions regarding the response of inhibitory neurons to single-cell perturbations and the modulation of perturbation response by neuronal gain.

**Significance Statement:** The cerebral cortex is strongly recurrently connected with complex wiring rules. This circuitry can now be probed by studying responses to optogenetic perturbations of one or small numbers of cells. However, we currently lack a general theory connecting these responses to underlying circuitry. Here we develop a novel, exactly solvable theory to determine responses to small perturbations from the underlying connectivity. Analysis of these equations reveals simple rules that govern perturbation response patterns. Comparison with experimental data yields new constraints on the connectivity parameters. The theory yields predictions for the responses of unmeasured cell types and in new experimental conditions.

---

In recent years, methods have been developed to optogenetically perturb small sets of neurons to probe recurrent neuronal circuitry. Such experiments in layers 2/3 (L2/3) of the mouse primary visual cortex (V1) have found that perturbation responses depend in complex ways on the spatial locations and orientation tunings of both the perturbed and the unperturbed neurons (1–7).

A common approach to making sense of this rich structure is to model mouse V1 L2/3 with a linear, recurrently-connected firing rate model in which connectivity strength depends on the spatial location, orientation tuning, and cell type of the pre- and post-synaptic neurons (2). While such models provide much simpler descriptions than biophysical spiking models and are analytically tractable for weak connectivity, there is still a lack of a more general understanding of how the perturbation response is related to the underlying connectivity structure.

Here we introduce a novel analytical approach to the problem. First, we show that a certain exponential-like spatial connectivity kernel is a good descriptor of the product of connection probability and synaptic strength. This choice of kernel allows us to derive an exact solution for the linear perturbation response of recurrently connected networks with multiple cell types that is valid regardless of the strength of recurrent coupling. For a linear network to be stable, the largest eigenvalue of its weight matrix must have real part *<* 1; but the spectral radius – the largest absolute value of an eigenvalue – may be arbitrarily large, and typically increases with the strength of recurrent coupling. Previous analytical methods have relied on a spectral radius *<* 1, *i*.*e*. on sufficiently weak coupling. Our method, which works for arbitrary spectral radius, allows the investigation of perturbation responses of inhibition stabilized networks (ISNs) (8, 9) – networks with recurrent excitation strong enough to be unstable without feedback inhibition. ISNs appear to describe cortical circuits (10) and commonly have spectral radius *>* 1.

The general solution for the circuit involving an arbitrary number of cell-types and connectivity length scales is complex, and does not easily provide intuitive insight. However, for the case of an excitatory/inhibitory (E-I) network in which connectivity width depends only on presynaptic cell types, we discover simple mathematical rules relating connectivity structure to single-cell perturbation response. This allows us to infer constraints that the cortical connectivity must satisfy to explain existing optogenetic perturbation data.

In particular, perturbation of a single excitatory neuron causes excitation of nearby neurons and suppression of neurons further away (1). Intuitively this might suggest that inhibitory connections are more spatially widespread than excitatory connections, but anatomical findings indicate the opposite (11, 12). Our theory shows that the pattern of nearby excitation and broad suppression arises when the strength of inhibition is sufficiently strong, regardless of whether inhibitory connections have a broader or narrower spatial spread than excitatory connections. More generally, we can quantitatively predict the spatial layout of excitatory and suppressive responses from network parameters.

Perturbation of a single excitatory neuron also causes, on average, stronger suppression of neurons with similar orientation tuning preference than those with opposite tuning preference (1); we call this perturbation response “opposite-favoring”. We find that feature-specific disynaptic inhibition (E →I → E) must be like-to-like (*i*.*e*. neurons with similar preferred features are more strongly connected) in order to generate opposite-favoring mean responses and, more generally, to generate perturbation responses that are opposite-favoring at any distance.

Our theory yields several experimentally testable predictions. First, existing studies of excitatory neuron perturbations primarily studied the responses of excitatory neurons. We predict that inhibitory neuron responses should have less suppression and spatially broader nearby excitation than excitatory responses. Second, perturbations may be performed with or without simultaneous presentation of visual stimuli. We predict that the absence of visual stimuli, which reduces firing rate and hence neuronal gain, can yield feature-tuning dependence of perturbation response opposite to that when visual stimuli are present. Finally, we predict that the absence of visual stimuli should generally result in responses with less suppression and broader nearby excitation, possibly eliminating the presence of zero-crossings altogether.

## Results

We study responses to moderate single-cell perturbations. Because these perturbations are small, we expect a linear theory to be adequate. To this end, we consider a linear recurrent neuronal network with *N*_*c*_ cell types in *d* spatial dimensions, where each neuron has a feature tuning preference and selectivity (Fig. 1A). We assume that cells are homogeneously distributed in space and feature tuning preference. Each neuron is uniquely indexed by four variables, a four-tuple (*α, µ*, ***x***, *θ*). *α* is an integer ranging from 0 to *N*_*c*_ − 1 that indicates the cell-type; *µ* is a number between 0 and 1 that indicates the cell’s feature selectivity (*i*.*e*. how well tuned the neuron is); ***x*** is a *d*-dimensional vector that represents the spatial location of the cell, and finally *θ* is a circular variable between −*π* and *π* that represents feature tuning preference. The firing rate of neuron (*α, µ*, ***x***, *θ*) at time *t* is written *r*_*α*_(*µ*, ***x***, *θ, t*), while the connectivity weight between postsynaptic neuron (*α, µ*, ***x***, *θ*) and presynaptic neuron (*β, ν*, ***y***, *ϕ*) is denoted *W*_*αβ*_(*µ, ν*, ***x*** − ***y***, *θ* − *ϕ*). Feature selectivity is assigned independently to each neuron and may be arbitrarily distributed with density *P*_*α*_(*µ*). The external input to each neuron is denoted *h*_*α*_(*µ*, ***x***, *θ*). For the single-cell perturbations we are considering, *h* is a delta function given by Eq. 11 (Materials and Methods). Taking the continuum limit for our analytical work, the dynamical equation of the network that determines *r*_*α*_(*µ*, ***x***, *θ, t*) is given by Eq. 12 (Materials and Methods). We are primarily interested in the steady-state of Eq. 12, *r*_*α*_(*µ*, ***x***, *θ*) = lim_*t*→*∞*_ *r*_*α*_(*µ*, ***x***, *θ, t*), which exists if and only if the network is stable, and is given by the equilibrium equation

**Fig. 1.**
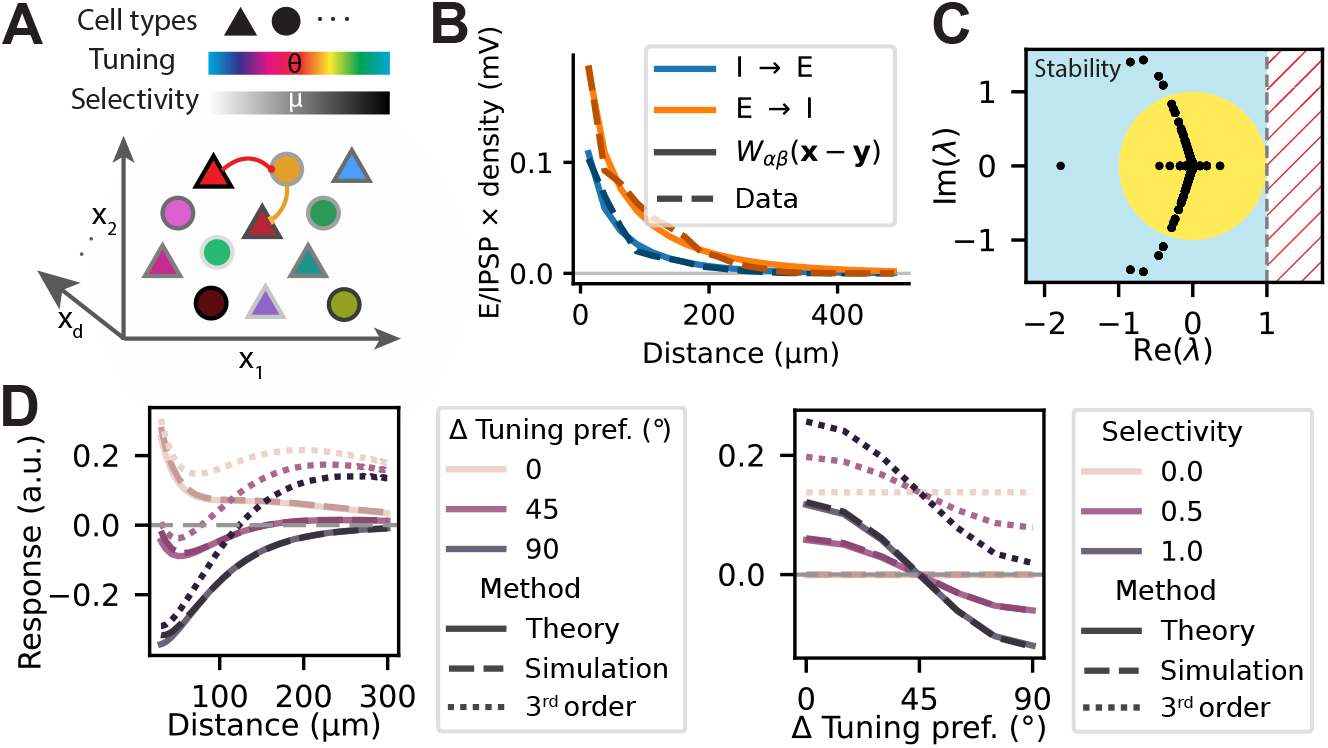
Model setup and theory. **A)** Schematic of model. Neurons are located in a *d*-dimensional space with *N*_*c*_ cell types, and have feature tuning preferences *θ* and selectivities *µ*. **B)** Connectivity function in a simplified 2D model, *W*_*αβ*_ (***x*** − ***y***) (Eq. 3), fitted to the product of connection density (11) and connection strength (13) between excitatory and inhibitory neurons. **C)** Eigenvalues of the connectivity matrix of the model shown in D). All eigenvalues lie in the region of stability (blue, Re(*λ*) *<* 1), for which our theory applies, but some lie outside the region of convergence (yellow, |*λ*| *<* 1) for the matrix inverse expansion 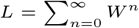 used in existing theoretical analyses of perturbation response. **D)** Comparison between theory, simulation, and 3^rd^ order matrix inverse expansion of single-cell perturbation response in an E-I model with spectral radius of 1.8. Left: Response of excitatory neurons as a function of distance to the perturbed excitatory neuron, for different feature tuning preferences. Right: Same, but as a function of difference in tuning preference for different feature selectivities, averaged over neurons 30-300 µm away from the perturbed neuron.

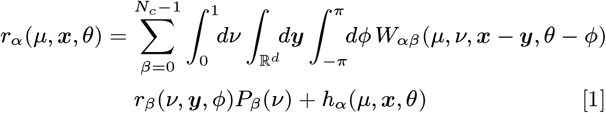

Here, the first term on the right represents the total recurrent input, which is a sum over input from neurons of all cell types *β*, feature selectivities *ν*, spatial locations ***y***, and feature tuning preferences *ϕ*. The term involves the convolution of connectivity with firing rates, both in space and feature, which is the continuum analog of a matrix multiplication. We will use the word “kernel” to refer to a function that is used in a convolution, as in the connectivity here. In general, for an arbitrary choice of *W*, there is no closed-form analytical solution for *r*_*α*_(*µ*, ***x***, *θ*). Our key insight is that *W* can be chosen such that it captures the spatial dependence of the product of the connection probability and the synaptic strength between cells (Fig. 1B), and admits a closed-form analytical solution for *r*_*α*_(*µ*, ***x***, *θ*), as we now explain.

We will make the common assumption that the dependence of *W* on space and feature can be factorized. The spatial dependence is commonly modeled as a Gaussian kernel (14– 20), in accordance with the approximately Gaussian spatial profile of connection probability measured in mouse V1 L2/3 (11, 12). However, this choice of spatial kernel neglects the spatial decay of synaptic strength (13) and does not admit a closed-form solution for Eq. 1. Instead, we propose a new form of spatial kernel that does allow a closed-form solution, as follows. We let *σ* be the connectivity length scale, and define *λ* = *σ*^−2^. We then define the spatial kernel to be the Green’s function (the functional inverse) of the operator *λ* −∇^2^ in *d*-dimensions, which we call *G*_*d*_(*s*; *λ*), or equivalently *G*_*d*_(*s*; *σ*^−2^), where *s* = ∥***x*** − ***y***∥ is the spatial distance in *d*-dimensions. Specifically, *G*_*d*_ is a monotonic, exponentiallydecaying kernel given by

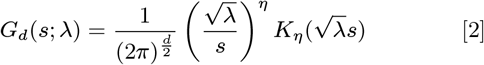

where 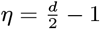 and *K*_*η*_ (*z*) is the modified Bessel function of the second kind with order *η* (SI section 2). In 1 and 3 dimensions, *G*_d_ (*s*; *λ*) is proportional to 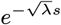 and 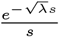 respectively; the shape of *G*_*d*_(*s*; *λ*) in 2 dimensions is shown by the solid lines in Fig. 1B. We combine data from (11) and (13) to compute the product of connection density and connection strength as a function of distance between excitatory and inhibitory neurons in mouse V1 L2/3. We find that our kernel can well capture this dependence (Fig. 1B; Materials and Methods), with best-fit E →I and I →E connectivity widths given by *σ*_E_ = (150 ± 11) µm and *σ*_I_ = (108 ± 8) µm respectively.

### Derivation for a simplified model

To illustrate the key ideas behind our derivation of the linear response for the full model, we first consider a simplified model whose connectivity depends only on the cell type and spatial location of the pre- and post-synaptic neurons, and whose connectivity width depends only on the pre-synaptic cell type. For this simplified model, the connectivity function is given by

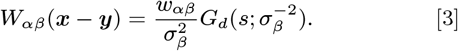

where *s* = ∥***x*** − ***y***∥, and the division by 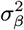 ensures that the integral of *W*_*αβ*_ over space is *w*_*αβ*_.

To solve for the system’s linear response to a perturbation, we use the standard bra-ket notation (Materials and Methods) to rewrite Eq. 1 in a more abstract form

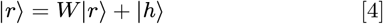

where |*r*⟩, |*h*⟩ are the firing rate function and the perturbing input function respectively, and *W* is the linear integral operator that acts on *r* according to Eq. 1. More precisely, |*r*⟩ is an *N*_*c*_ × 1-vector of firing rate functions *r*_*α*_(***x***), one for each cell type *α* (and similarly for |*h*⟩); *W* is an *N*_*c*_ × *N*_*c*_ matrix of spatial kernels, with individual elements given by Eq. 3; and *W r* indicates the combination of matrix-vector multiplication and spatial convolution. The response vector can be written as *r* = (*I* − *W*)^−1^ |*h*⟩ where *I* is the identity operator. Our goal is to compute the operator *L* := (*I* − *W*)^−1^ (where the inverse represents the combination of matrix inverse and functional inverse).

The most common approach is to compute the perturbative expansion of the linear response operator in the form of a Neumann series 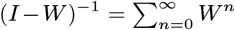 (2, 21–25). However, this approach suffers from two key issues: 1) the series does not converge for operators *W* whose spectral radius is greater than 1 (Fig. 1C), and thus is not typically applicable to biological networks with strong recurrent connectivity, such as ISNs; and 2) even when the series converges, the number of terms required for a good approximation may be large, thus failing to provide a simple description of the relationship between connectivity and perturbation response.

The choice of spatial kernel *G*_*d*_ allows for exact computation of the inverse *L* = (*I* − *W*)^−1^. Because *G*_*d*_(*s*; *σ*^−2^) is the Green’s function of *σ*^−2^ − ∇^2^, convolution of the firing rates by the spatial kernel *G*_*d*_(*s*; *σ*^−2^) is equivalent to applying the inverse operator (*σ*^−2^ − ∇^2^)^−1^ to the firing rates. Thus, if we define ***W*** as the *N*_*c*_ × *N*_*c*_ matrix with elements *w*_*αβ*_ and **Σ** as the *N*_*c*_ × *N*_*c*_ diagonal matrix with elements 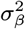, then we may write the connectivity operator *W* in Eq. 3 as:

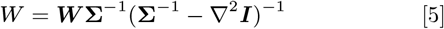

where **Σ**^−1^ − ∇^2^***I*** represents a *N*_*c*_ × *N*_*c*_ diagonal matrix of spatial operators with elements 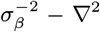, such that the inverse operator (**Σ**^−1^ − ∇^2^***I***)^−1^ acts on |*r*⟩ by applying 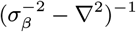 to *r*_*β*_(***x***) for each cell type *β* (see SI section 3 for a more thorough explanation, and Eq. S28 for a more formal approach to defining *W* using tensor products).

By the Woodbury matrix (operator) identity (26), (*I* − *UC*^−1^)^−1^ = *I* + *U* (*C* − *U*)^−1^ for any operators *U, C*. Thus, if we take *U* = ***W* Σ**^−1^ and *C* = **Σ**^−1^ − ∇^2^***I***, and assume that (***I*** − ***W***)**Σ**^−1^ is diagonalizable as ***Q*Λ*Q***^−1^, then we obtain an explicit expression for the linear response operator L:

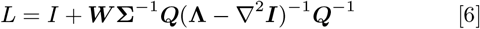

The identity operator *I* only affects a neuron’s response to its own perturbation, so 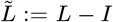 determines a neuron’s response to perturbation of a different neuron.

As *G*_*d*_(*s*; *λ*) is the Green’s function of *λ*− ∇ ^2^ for a scalar lambda, multiplying any function by (*λ* − ∇^2^)^−1^ is equivalent to convolving it with *G*_*d*_(*s*; *λ*). We can thus express 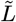 in terms of cell-type indices and spatial variables as

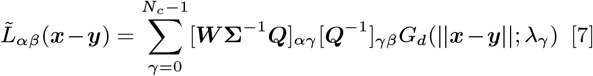

where *λ*_*γ*_ are the diagonal entries of **Λ**, *i*.*e*. the eigenvalues of (***I*** − ***W***)**Σ**^−1^. This is the response of neuron (*α*, ***x***) to a unit amplitude perturbation of a different neuron (*β*, ***y***).

### The full model

We define the connectivity function of the full space- and feature-dependent model by

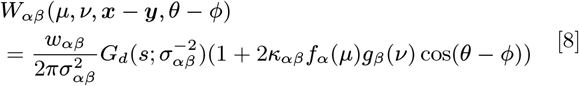

where *κ*_*αβ*_ lies between -0.5 and 0.5, and *f*_*α*_, *g*_*α*_ are monotonically increasing functions over the interval 0 to 1, such that *f*_*α*_(0) = *g*_*α*_(0) = 0, *f*_*α*_(1) = *g*_*α*_(1) = 1. The sign of *κ*_*αβ*_ determines whether connectivity is correlated or anticorrelated with difference in feature preference, *i*.*e*. whether the connection is like-to-like (*κ*_*αβ*_ *>* 0) or anti-like-to-like (*κ*_*αβ*_ *<* 0), whereas the functions *f*_*α*_ and *g*_*α*_ determine the strength of this correlation as a function of feature selectivity. Under this choice of *W*, the steady-state response of *r*_*α*_(*µ*, ***x***, *θ*) to a unit amplitude single-cell perturbation of a different neuron (*β, ν*, ***y***, *ϕ*) is (SI section 4)

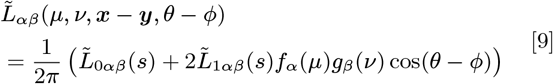

where 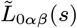 and 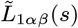 are defined by

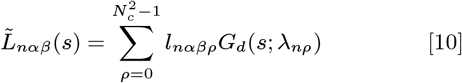

with *l*_*nαβρ*_ and *λ*_*nρ*_ given by Eq. S49 and S42 respectively. This generalizes Eq. 7 to allow connectivity widths to depend on both pre- and post-synaptic cell types and to include feature-preference dependence.

Since Eq. 9 is exact, we expect a close agreement between our theory and numerical simulations of the model regardless of the spectral radius of the connectivity matrix. Indeed, we obtain near perfect agreement between our theory and numerical simulations for the single cell perturbation response in an E-I model with two spatial dimensions and a spectral radius of 1.8 (Fig. 1C, D; Materials and Methods). For comparison, we also computed the perturbation response using the Neumann series expansion of the matrix inverse up to 3^rd^ order (*i*.*e*. 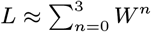). This is the minimum order at which the responses of excitatory neurons to the perturbation of a single excitatory neuron depend on all connectivity weights (including I →I weights). As expected, the series expansion severely diverges from simulations due to the spectral radius being greater than 1 (Fig. 1D).

### Mean response of unperturbed neurons

In experiments, perturbing a single pyramidal neuron results in mean suppression of the unperturbed neurons (1), suggesting that inhibitory connections are sufficiently strong in order to overcome recurrent excitation. To derive the precise conditions under which mean suppression occurs, we integrate Eq. 9 over all its continuous variables (Eq. S55, SI section 5). In the case of an E-I model studied here, it can be shown that for single-cell excitatory neuron perturbations, unperturbed excitatory neurons are suppressed on average if and only if det(***W***) *> w*_EE_, or equivalently, |*w*_EI_| *w*_IE_ *> w*_EE_(|*w*_II_| + 1), while inhibitory neurons are always excited on average (SI section 8). Thus the observation of mean suppression of unperturbed neurons implies that the disynaptic E → I → E inhibition must be stronger than the product of E → E excitation and I → I inhibition.

### Spatial profile of perturbation response

Single-cell perturbations of pyramidal neurons produce a Mexican-hat-shaped response as a function of distance, where neurons near the perturbed site are excited and neurons farther away are suppressed (1). Intuitively, this would suggest a connectivity motif of narrow excitation and broad inhibition. However, recent mouse V1 L2/3 connectivity data shows that the opposite is true: E → I and I → E connections are narrower than E →E connections (11, 12). Furthermore, the length scale of E →E connectivity (standard deviation ≈125 µm for a Gaussian spatial profile, 12) is significantly broader than the spatial radius of nearby excitation (≈70 µm, 1), and an even shorter radius of excitation (≈35 µm) is seen for multi-cell perturbations, which could not be explained by a model with a Gaussian spatial profile for each connection (2). Thus we set out to investigate the conditions under which such small radii of nearby excitation can arise in our model with realistic connectivity length scales.

### Number of spatial zero crossings

The Mexican-hat-shaped spatial profile of perturbation response implies that the response crosses zero from nearby activation to suppression at least once, or in other words, that there is at least one zero crossing in the response as a function of distance from the perturbation. It is conceivable that the response changes sign more than once, but that these zero crossings cannot be detected due to measurement noise. Thus, the question of whether or not the model can exhibit a Mexican-hat-shaped perturbation response can be broken into two mathematical sub-problems: whether or not nearby neurons are activated, and whether or not there exists at least one zero crossing in the response as a function of distance. We find that for all networks with 2 or more spatial dimensions, single-cell excitatory neuron perturbations always activates nearby neurons, in the mathematical sense that neurons arbitrarily close to the perturbed cell are activated (SI section 9).

To proceed further, we assume that connectivity width depends only on pre-synaptic cell type. In this case, we find that E-I models may exhibit either 0, 1, or infinitely many zero crossings (SI section 10A). The exact behavior is determined by both connectivity width and connectivity strength via the two eigenvalues *λ*_*γ*_ of the 2 × 2 matrix (***I*** − ***W***)**Σ**^−1^. These are the same eigenvalues which characterize the perturbation response in the simplified model (Eq. 7). If *λ*_0_, *λ*_1_ are complex conjugates, then the responses of both excitatory and inhibitory neurons must exhibit infinitely many zero crossings. If *λ*_*γ*_ are real and the network is an ISN with two or more spatial dimensions, the condition for the excitatory neuron response to have exactly one zero crossing is that the smaller of the two eigenvalues, *λ*_0_, satisfies 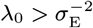. A similar condition for inhibitory neurons is 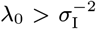 (SI Corollary 10.4). Thus, not only can the model exhibit a Mexican-hat-shaped perturbation response, but we are also able to determine the precise conditions under which it occurs.

The mathematical conditions on the number of zero crossings can be formulated in terms of the connectivity strengths *w*_*αβ*_ and the ratio of inhibitory to excitatory connectivity width 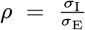. These conditions, and the following results presented in Figures 2 and 3, assume that the E-I network is an ISN with two or more spatial dimensions whose connectivity widths depend only on presynaptic cell type. We first consider the special case in which the inhibitory and excitatory spatial kernels have equal width (*ρ* = 1). In this case, the number of zero crossings of excitatory neuron response can be represented as a phase diagram in terms of the trace and determinant of ***W*** (Fig. 2A). This diagram reveals simple principles governing the number of zero crossings: First, the existence of at least one zero-crossing implies det(***W***) = |*w*_EI_|*w*_IE_ − *w*_EE_|*w*_II_| must be positive, and hence disynaptic E → I → E inhibition must be stronger than the product of E → E and I → I connections. Second, I → I connections must be stronger than E →E (so that the x-axis value is negative) for the response to exhibit exactly one zero crossing. This suggests I →I connections may have the regularizing role of suppressing spatial oscillations.

**Fig. 2.**
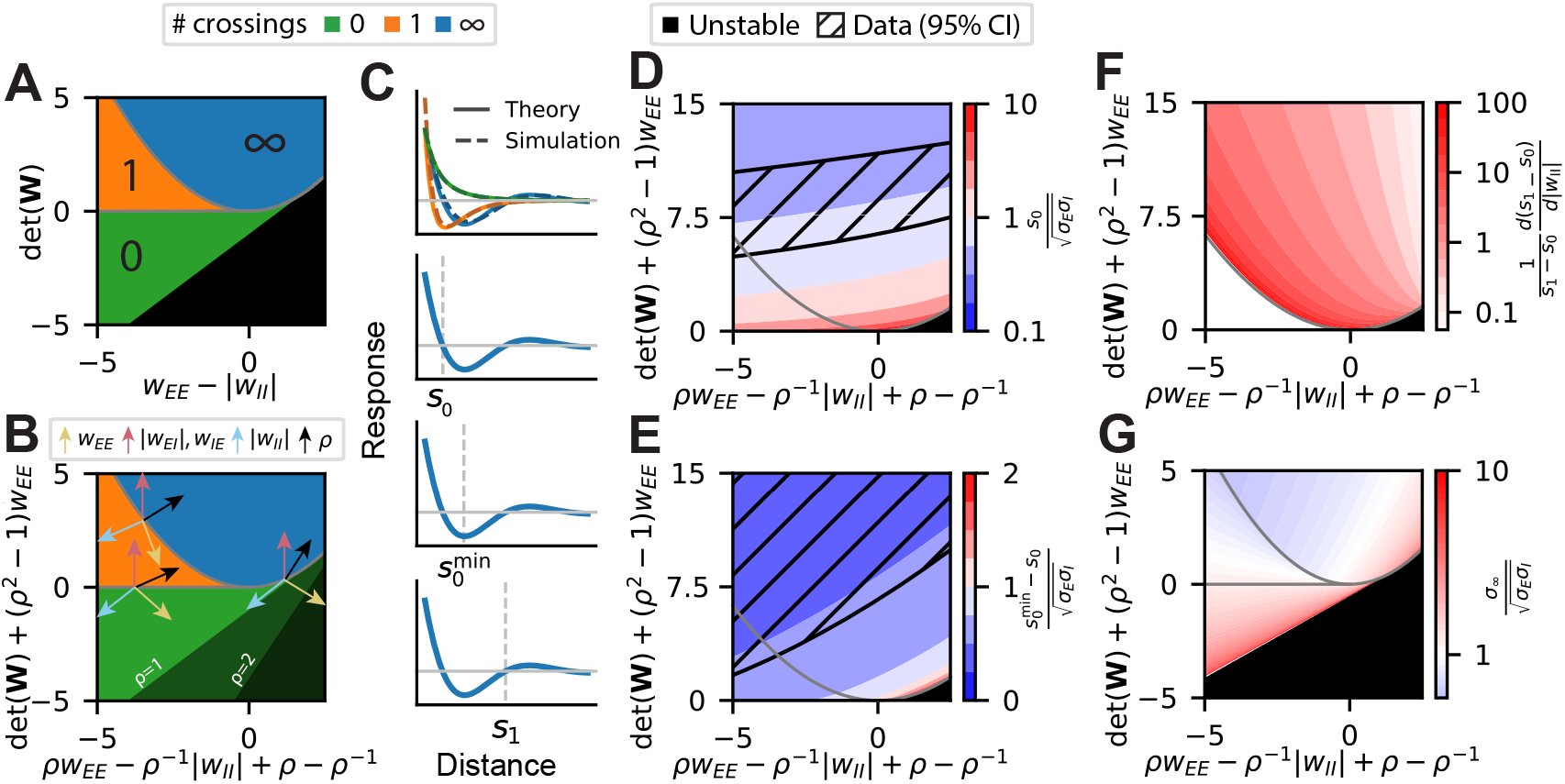
Spatial profile of excitatory neuron response to single-cell perturbation in E-I ISNs. **A)** Phase diagram of the number of zero crossings in the perturbation response as a function of distance from the perturbation for networks with *ρ*=1 (*i*.*e. σ*_*E*_ = *σ*_*I*_). Shaded region in black indicates dynamically unstablility. The phase boundaries between 0, 1, and *∞* are given by *y* = 0 and 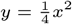. **B)** Phase diagram of the number of zero crossings for networks with arbitrary *ρ*. The instability region is dependent on *ρ*, with boundary *y ρ x* −2*ρ* for *x* ≤ *ρ* and 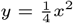 for *x >* 2ρ. Areas shaded in dark green and even darker green illustrate the instability regions for *ρ* and *ρ* respectively. Arrows indicate changes in number of zero crossings induced by perturbations of each parameter at the phase boundaries. **C)** Top panel: Example perturbation responses as a function of distance within each of the three phase regions, computed either with the theory or with numerical simulations. Theory curves are suitably normalized for comparison, and the same normalization factors are applied to the corresponding simulation curves.. Remaining panels: Illustration of the quantities 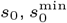, and *s*_1_ as plotted in D-F. **D)** Location to the first zero-crossing, *s*_0_, as a fraction of the connectivity length scale 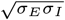 for 2-dimensional models with *w*_EE_ = 5, *ρ* = 0.72. Hatched region indicates 95% confidence interval of this fraction as estimated from experimental data (1, 11, 13; Materials and Methods). Grey line indicates the boundary between 1 and *∞* zero crossings as seen in A and B. **E)** Similar to D, but the distance from the first zero crossing *s*_0_ to the first minimum 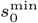 is plotted. **F)** Similar to D, but the derivative of the distance from the first zero-crossing *s*_0_ to the second zero-crossing *s*_1_ with respect to recurrent inhibition strength, normalized by this distance, is plotted. **G)** Asymptotic decay length scale *σ*_*∞*_ for models with *ρ* = 0.72. Note that unlike D-F, this variable is independent of the specific choice of *w*_EE_ and the number of spatial dimensions *d*. Panels D-F are computed for 2-dimensional models; panels A and B are valid for 2 or more dimensions.

**Fig. 3.**
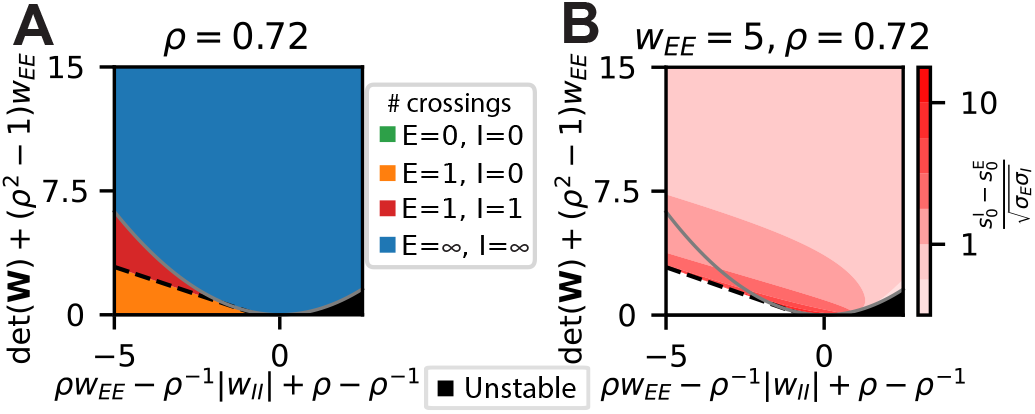
Spatial profile of excitatory vs. inhibitory neuron response to single-cell perturbation in E-I ISNs with 2 or more spatial dimensions. **A)** Phase diagram of the number of E and I zero crossings for networks with *ρ* = 0.72. Green: Neither E nor I exhibit zero crossings. Orange: E exhibits 1 zero crossing, I exhibits 0. Red: Both E and I exhibit 1 zero crossing. Blue: Both E and I exhibit infinitely many zero crossings. Dashed line is given by *y* = (*ρ* − *ρ*^−1^)(*x* − (*ρ* − *ρ*^−1^)). **B)** Distance between the first zero crossing of excitatory neuron response and the first zero crossing of inhibitory neuron response in networks with two spatial dimensions and *w*_EE_ = 5, *ρ* = 0.72.

Our phase diagram can be generalized to the case of arbitrary *ρ* by modifying the axes (Fig. 2B; SI section 10B). To gain intuition about this phase diagram, we analyze the change in the number of zero crossings induced by increasing each of the connectivity parameters at the phase boundaries (SI section 10C), as indicated by the arrows in Fig. 2B. We find that increasing *ρ* (*i*.*e*. broadening inhibitory connections) or the strength of E → I or I →E connections encourages the formation of zero crossings, while increasing the strength of E →E or I → I connections has the opposite effect. Furthermore, the phase diagram reveals that the principles we obtained for the case of *ρ* = 1 can be generalized: First, the existence of at least one zero crossing implies det(***W***) must be greater than (1 − *ρ*^2^)*w*_EE_, and thus must be positive for networks with *ρ <* 1. Second, for the network response to exhibit exactly one zero crossing, I →I connections must be stronger than *ρ*^2^*w*_EE_ + (*ρ*^2^ − 1), and thus must be stronger than E →E connections if *ρ >* 1. Finally, since this phase diagram is valid for all *ρ*, this implies that as long as the disynaptic E →I →E connections are sufficiently strong (*i*.*e*. the y-axis value is sufficiently large), the model can exhibit at least one zero crossing regardless of the relative widths of excitatory and inhibitory connections.

### Spatial radius of nearby excitation

The location of the first zero crossing (*i*.*e*. the spatial radius of nearby excitation, Fig. 2C), *s*_0_, has been measured at approximately 70 µm (1), which is significantly narrower than the connection probability length scale at around 100 – 125 µm (11, 12). Can this be explained by our model? To address this we compute *s*_0_ at different points of the phase space as a fraction of the connectivity length scale 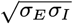. Since this quantity is not fully determined by the x- and y-axes of Fig. 2B, we compute it for different combinations of *w*_EE_ and *ρ* (Fig. S1). The specific case of *w*_EE_ = 5, *ρ* = 0.72 is illustrated in Fig. 2D, where the value of 0.72 is our best estimate of *ρ* obtained from the fitted connectivity kernels in Fig. 1B. The value of *w*_EE_ is chosen so that it is within the distribution of values of *w*_EE_ obtained by fitting our model to data (see Validation of theoretical insights in fitted models; Fig. S6A) and the network is an ISN. These numerical results show that *s*_0_ is negatively correlated with det(***W***), and that the determinant must be considerably greater than 0 (*i*.*e*., disynaptic E →I → E inhibition must be significantly stronger than the product of E →E and I →I connections) in order to explain the narrow Mexican-hat-shaped response profile observed by (1). Note that this condition is more stringent than the necessary condition det(***W***) *>* 0 for the existence of at least one zero crossing when *ρ <* 1.

### Spatial location of maximum suppression

Further constraints on the connectivity parameters can be inferred by considering the distance to the first local minimum 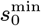 of the perturbation response (Fig. 2C), which we expect to be the spatial location of maximum suppression. Unlike the location of the first zero crossing *s*_0_, the additional distance to the first minimum, 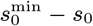, is moderately invariant to the specific choice of *w*_EE_ and *ρ* (Fig. S2; Fig. 2E shows the specific case of *w*_EE_ = 5, *ρ* = 0.72). Combined with the observation that the contour lines of 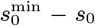 are diagonal, this implies a correlation between the values of det(***W***) and tr(***W***) that can explain the data.

Experimental data places 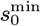 at around 110 µm (1), so that 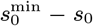 is around 40 µm, which is less than half of the connectivity width length scale of 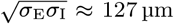 as measured from Fig. 1B. This would place the network in the darker blue region – roughly, the upper left triangle – of Fig. 2E, which overlaps considerably with the appropriate region of Fig. 2D as determined above.

### Frequency of spatial oscillations

As we have shown, the region of phase space with only one zero-crossing requires sufficiently strong I →I connectivity (Fig. 2B). This suggests that recurrent inhibition is important for suppressing spatial oscillations. This intuition can be made precise by considering the the derivative of the distance from the first zero-crossing *s*_0_ to the second zero-crossing *s*_1_ (Fig. 2C) with respect to |*w*_II_|. A positive derivative indicates that increasing the strength of recurrent inhibition reduces the frequency of spatial oscillations. As expected from the intuition, we numerically observe a positive derivative regardless of the model parameters (Fig. 2F, S3). In fact, it can be proven that in one or three spatial dimensions, this derivative must be positive (SI section 11B), and we conjecture that this results also holds in two spatial dimensions.

### Stability and spatial decay length scale

Finally we consider the rate at which the perturbation response decays with distance. Since the response is a non-monotonic function of distance, we measure its asymptotic decay length scale *σ*_*∞*_, defined such that the perturbation response decays asymptotically as 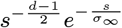 as *s* → *∞*. Under the assumption of fast inhibition, we find an interesting relationship between *σ*_*∞*_ and the overall stability of the network: the closer the network is to the edge of instability, the longer the decay length scale (Fig. 2G, SI section 13). This relationship is fully general, applying to networks with arbitrary number of cell types and arbitrary connectivity widths and spatial dimensions. Thus, assuming sufficiently fast inhibition, observation of a decay length scale of the same order of magnitude as, or smaller than, the connectivity length scale would suggest that the network is reasonably far from the edge of instability.

### Inhibitory neuron response

Thus far we have focused on the perturbation responses of excitatory neurons. This is because, to the best of our knowledge, existing simultaneous two-photon optogenetics and calcium imaging experiments in mouse V1 either do not discriminate between the responses of excitatory and inhibitory neurons, or only measure the responses of excitatory neurons (1–3, 5). However, the responses of inhibitory neurons encode important information about the recurrent connectivity: for example, whether the cortical circuit is an ISN can be determined by a paradoxical effect whereby inhibitory neurons are suppressed by optogenetic stimulation of inhibitory neurons (8–10, 27, 28).

We find that, in response to perturbation of a single excitatory cell, the responses of inhibitory neurons are tightly related to those of excitatory neurons. Consider, again, an E-I ISN in two or higher dimensions with connectivity widths depending only on pre-synaptic cell type. We can show that 1) the excitatory neuron response is oscillatory (having an infinite number of zero crossings as a function of distance) if and only if inhibitory neuron response is also oscillatory, and 2) if excitatory neuron response exhibits a single zero-crossing as a function of distance, then inhibitory neuron response must also exhibit a single zero-crossing unless 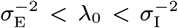 (SI section 10E). These relations are summarized by Fig. 3A. Thus we expect the responses of inhibitory neurons to likely exhibit the same number of zero-crossings as excitatory neurons. Violations of this expectation, however, would suggest that the connection strengths between E and I satisfy tight inequalities.

Now suppose that inhibitory neurons indeed exhibit a Mexican-hat shaped response profile. As explained in the section Mean response of unperturbed neurons, mean inhibitory neuron response must be positive. Given the mean suppression of excitatory neurons, we thus expect less lateral suppression of inhibitory neurons than excitatory neurons. Furthermore, it can be shown that if, and only if, det(***W***) *>* 0, the inhibitory response profile has a greater spatial radius of nearby excitation than the excitatory response profile, that is, 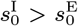, where 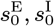 are the distances to the first zero crossing of excitatory and inhibitory neuron responses respectively (Fig. S4; SI section 11D). In particular, for networks with *ρ* ≤ 1, the inequality *s*^I^ *> s*^E^ is always satisfied (Fig. 3B, S4), since the existence of an excitatory zero crossing implies det(***W***) *>* 0 for such networks (Fig. 2B). Furthermore, recall that there is mean suppression of excitatory neurons if and only if det(***W***) *> w*_EE_. Thus, given mean suppression of excitatory neurons, inhibitory neuron response must be less suppressed and exhibit a broader spatial profile than excitatory neuron response.

### Feature-tuning dependence of perturbation response

Upon optogenetic perturbation of a single excitatory neuron, neurons in L2/3 of mouse V1 with similar tuning to that of the perturbed cell (iso-tuned neurons) are, on average over space, more suppressed than those with orthogonal tuning (ortho-tuned neurons) (1). We call this an *opposite-favoring* response, as opposed to a *same-favoring* response in which iso-tuned neurons are more excited than ortho-tuned neurons (Fig. 4A). Since E →E connectivity in L2/3 of mouse V1 is *like-to-like*, (similarly tuned excitatory neurons are preferentially connected) (11, 29), this suggests the need for a like-to-like disynaptic E →I →E motif to obtain preferential suppression of similarly tuned excitatory neurons.

**Fig. 4.**
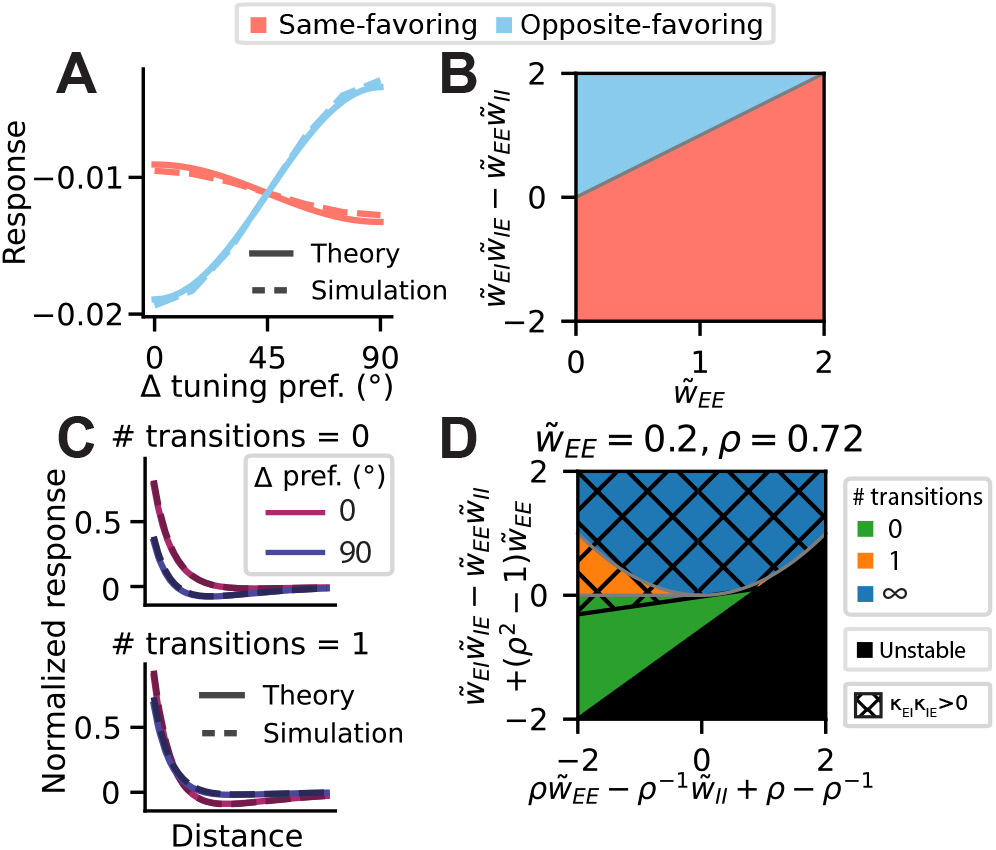
Feature-tuning dependence of excitatory neuron response to single-cell perturbation. **A)** Examples of same-favoring and opposite-favoring responses. **B)** Phase diagram of feature tuning of perturbation response. Red indicates same-favoring response (iso-tuned neurons are more excited than ortho-tuned neurons), and blue indicates opposite-favoring response (the opposite of same-favoring). **C)** Example responses of networks with 0 and 1 transitions between same- and opposite-favoring response with increasing distance, computed either with our theory or with simulations. Theory curves are suitably normalized for comparison, and the same normalization factors are applied to the corresponding simulation curves.**D)** Phase diagram of number of such transitions for a two-dimensional network with 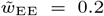 and *ρ* = 0.72. Green, orange, and blue represent 0, 1, and *∞* transitions respectively. Networks in the phase region shaded in black are dynamically unstable. Hatched region indicates like-to-like disynaptic E → I → E inhibition. Orange and blue regions are contained within the hatched region, showing that the presence of at least one transition implies like-to-like disynaptic inhibition.

To determine if this intuition is correct, we integrate Eq. 9 over space to obtain the average perturbation response as a function of feature tuning (Eq. S54). Given like-to-like E → E connectivity, we find that excitatory neuron response is opposite-favoring if and only if 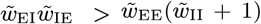 (Fig. 4B; SI section 14A), where 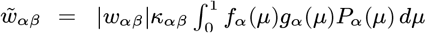 is positive if and only if the connectivity from cell type *β* to *α* is like-to-like, *i*.*e. κ*_*αβ*_ is positive. This condition requires like-to-like E →I →E inhibition (*κ*_EI_*κ*_IE_ *>* 0); while the condition could also be met with *κ*_EI_*κ*_IE_ ≤ 0 if 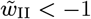, such networks are unstable (SI section 14B). Thus, the observation of opposite-favoring response implies that disynaptic E →I →E connections provide like-to-like inhibition.

The condition for opposite-favoring response is also dependent on the distribution of neuronal feature selectivity *P*_*α*_(*µ*) and the feature-selectivity dependence of connectivity *f*_*α*_(*µ*), *g*_*α*_(*µ*), due to the integral in the definition of 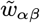. This integral indicates how inhomogeneity of feature selectivity could affect the feature tuning of perturbation responses. For example, a smaller fraction of tuned inhibitory neurons would result in weaker feature dependence of E →I and I →I connections on average (leading to smaller 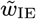 and 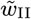), which could in turn result in same-favoring rather than opposite-favoring responses. Note that the connectivity and the perturbation response have identical functional dependence on feature selectivity (compare Eq. 8 to Eq. 9). Thus, the functions that characterize the dependence of connectivity on feature selectivity, *f*_*α*_(*µ*), *g*_*α*_(*µ*), can be easily inferred from suitably designed optogenetic perturbation experiments.

### Modulation of feature-tuning dependence by distance

The single-cell perturbation response measured experimentally is not only opposite-favoring on average, but also opposite-favoring at all distances beyond 25 µm, if one computes tuning similarity as signal correlation (1). We find that in models with two or more spatial dimensions and like-to-like E → E connections, sufficiently nearby excitatory neurons always exhibit same-favoring response (SI section 15). Thus, in order to explain the data, our model should exhibit a very nearby transition from same-favoring to opposite-favoring response with increasing distance from the perturbed neuron (Fig. 4C). Indeed, E-I networks with two or more spatial dimensions and connectivity width that depends only on presynaptic cell type can exhibit 0, 1, or *∞* number of transitions between same- and opposite-favoring response (SI section 15). The number of such transitions is determined by combinations of 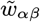 and *ρ* (Fig. 4D). Interestingly, given like-to-like E →E connectivity and the presence of at least one such transition (which is required to explain the data), disynaptic E → I →E connectivity must be like-to-like (Fig. 4D; SI Theorem 15.4). In other words, given that sufficiently nearby neurons have same-favoring responses (as predicted by our theory), if the perturbation response is opposite-favoring at any distance, then the disynaptic E →I →E inhibition must be like-to-like. Note that this finding is stronger than our previous finding that a response whose mean across distance is opposite-favoring implies like-to-like E → I → E connectivity.

### Modulation of perturbation response by neuronal gain

The single-cell optogenetic perturbation experiment that our analysis is based on was carried out with the simultaneous presentation of visual stimuli (drifting gratings) (1). However, a different optogenetic perturbation experiment in mouse V1 has been performed without the presentation of visual stimuli (grey screen) (2), which is likely to result in a lower firing rate of neurons during the experiment. If cortical cells have supralinear input/output functions (30–32, but see 33), then their gain – the change in rate for a given change in input – would be reduced for lower firing rates. This in turn would decrease the effective connection strengths which, in a model linearized about a fixed point, are given by the gains times the synaptic weights. As we have shown that connection strengths play a major role in determining the qualitative features of single-cell perturbation responses, we reason that the results of single-cell perturbation experiments performed without a visual stimulus may significantly deviate from those performed with visual stimulus. Thus we study how various perturbation response properties are modulated by neuronal gain, and predict the results of single-cell perturbation experiments performed in the absence of a visual stimulus.

### Modulation of mean perturbation response by neuronal gain

First we study how changes in neuronal gain (*g*), which in our model effectively scales all connectivity weights by *g*, modulate the mean response. We find that if the unperturbed excitatory neurons exhibit mean suppression, then increasing neuronal gain always results in stronger suppression (SI section 8). Similarly, reducing neuronal gain always results in weaker suppression or, for sufficiently small gain, mean excitation (Fig. 5A). Note that the derivative of the mean response with respect to gain is non-monotonic, such that if the unperturbed excitatory neurons exhibit mean excitation, then increasing the gain may result in stronger excitation instead.

**Fig. 5.**
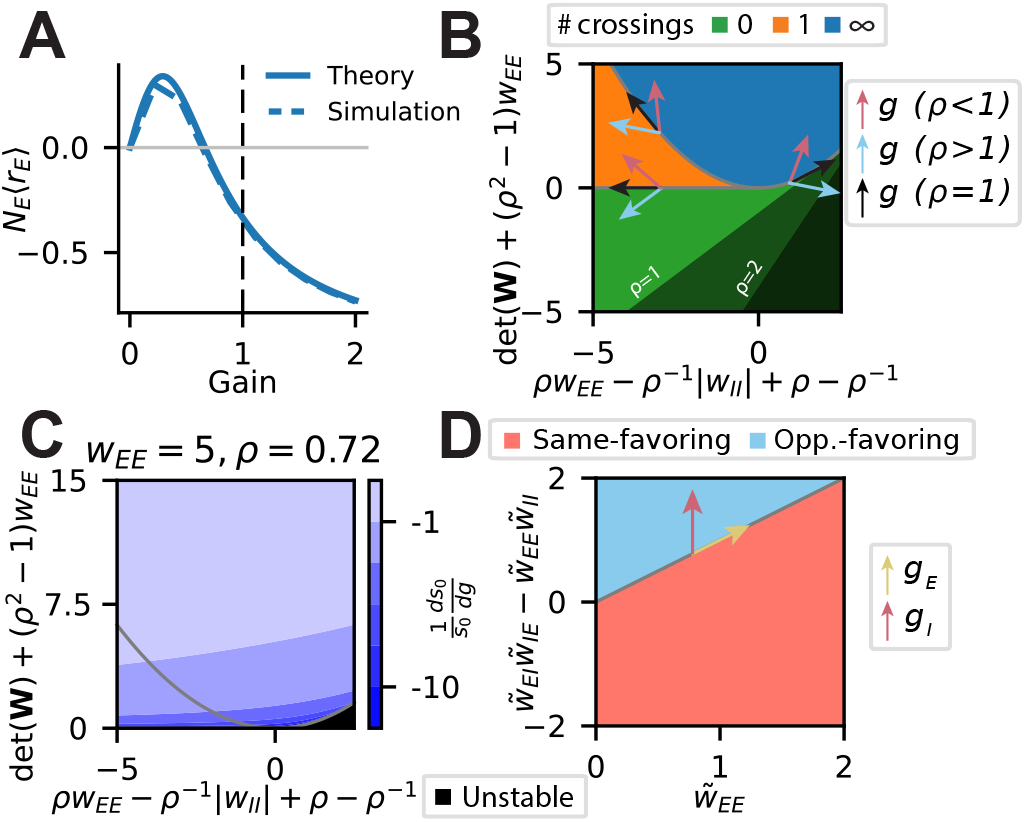
Modulation of excitatory neuron response to single-cell perturbation by neuronal gain in E-I ISNs with 2 spatial dimensions. **A)** Sum of responses of unperturbed excitatory neurons as a function of gain, for a network with *w*_EE_ = 2, *w*_II_ = −1, det(***W***) = 3, computed either with our theory or with numerical simulations. **B)** Phase diagram of number of zero crossings from Fig. 2B. Arrows indicate changes in number of zero crossings induced by increasing gain at the phase boundaries for *ρ* = 1, *ρ >* 1, or *ρ <* 1. **C)** Derivative of distance to the first zero crossing with respect to gain, divided by the distance, for *w*_EE_ = 5, *ρ* = 0.72. **D)** Phase diagram of feature tuning of perturbation response, with red indicating same-favoring response and blue indicating opposite-favoring response. Arrows indicate movement in phase space induced by increasing excitatory and inhibitory neuron gain respectively at the phase boundary.

### Modulation of the spatial profile of the response by neuronal gain

Next, we study the modulation of the number of spatial zero crossings by neuronal gain. We find that the changes in the number of spatial zero crossing due to increasing gain depend entirely on the value of *ρ* (Fig. 5B; SI section 10D): if *ρ* = 1, then an increase in gain does not change the number of spatial zero crossings of the response; while if *ρ <* 1 or *ρ >* 1, then, if starting from near a phase boundary, increasing gain increases or decreases, respectively, the number of zero crossings.

We next study the effect of gain on the location of the first zero crossing, *s*_0_. We find that when *ρ* = 1, the derivative of *s*_0_ with respect to the gain *g* is always negative, *i*.*e*. increasing the gain always produces narrower nearby excitation (SI section 11C). Numerically, we find that this also holds when *ρ <* 1 (Fig. 5C), and is mostly true when *ρ >* 1 (Fig. S5). Thus, given our estimate of *ρ* ≈ 0.72 in experimental data, we predict that single-cell perturbation experiments performed without the presence of visual stimulus should result in a broader response profile with less suppression and the same or a decrease in number of zero crossings.

### Modulation of feature dependence by neuronal gain

Finally, we consider how neuronal gain may affect the feature tuning dependence of perturbation response. We find that increasing gain may result in a transition from same-favoring to opposite-favoring excitatory neuron response (SI section 14C). Furthermore, if we selectively increase the gain of excitatory or inhibitory neurons only, we find that this transition is mediated by an increase in gain of inhibitory neurons *g*_I_ (*i*.*e*. effective scaling of all connections onto inhibitory neurons by *g*_I_), whereas increasing the gain of excitatory neurons *g*_E_ cannot yield such a transition (Fig. 5D). More importantly, it can be shown that a transition from opposite-to same-favoring response can always be induced by sufficiently decreasing the gain of inhibitory neurons (SI section 14C). Thus, we predict that single-cell perturbation experiments performed without visual stimuli may potentially result in same-favoring rather than opposite-favoring responses, suggesting that a supralinear transfer function of neurons (or at least of inhibitory neurons) may be important for switching between two qualitatively distinct computations.

### Validation of theoretical insights in fitted models

So far most of our theoretical analyses of the linear response equation have relied on two simplifying assumptions. First, we assumed, for theoretical tractability, that the connectivity width depends only on presynaptic cell type, *i*.*e*. the models obey the symmetries *σ*_EE_ = *σ*_IE_ and *σ*_EI_ = *σ*_II_. However, recent connection probability data from L2/3 of mouse V1 suggests that the connection-probability length-scales instead satisfy *σ*_EE_ ≈ *σ*_II_ *> σ*_EI_ ≈ *σ*_IE_ (12). Second, we have analyzed the single-cell perturbation response in (1) under the simplifying assumption that all measured responses are of excitatory neurons, while in the experiment, both excitatory and inhibitory neurons were measured. To test the robustness of our findings, we relax these assumptions and fit models so that the mix of 85% excitatory and 15% inhibitory cells match the perturbation response from (1), both as a function of distance and as a function of orientation tuning preference. We also constrain the models such that *σ*_EE_ and *σ*_II_ are broader than *σ*_EI_ and *σ*_IE_, and that the parameter *κ*_EE_ is within two standard deviations of our estimate from data of (11) (Materials and Methods). All neurons in the models are equally well tuned – effectively, we replace the term *f*_*α*_(*µ*)*g*_*β*_(*ν*) in Eq. 8 by 1 – because we know of no data on how perturbation response or connectivity probability/strength depends on feature selectivity, so the choice of functions *f*_*α*_(*µ*), *g*_*α*_(*µ*) is unconstrained.

From the 200 fitted models, we select the 50 best-fitting models for analysis (Materials and Methods). Consistent with our theoretical analysis of the spatial profile of perturbation response, all fitted models exhibit a positive determinant of the weight matrix ***W***, and the determinant and trace of ***W*** are correlated across models (Fig. 6A; compare Fig. 2D, E). Most fitted models (47/50) also exhibit like-to-like disynaptic E →I →E inhibition as suggested by our theory (Fig. 6B; compare Fig. 4D). Furthermore, we find that the three exceptions nonetheless confirm our prediction that negative *κ*_IE_*κ*_EI_ implies same-favoring excitatory responses; in all three cases, the same-favoring behavior of the excitatory cells is weak enough, and the opposite-favoring behavior of the inhibitory cells strong enough, such that the average over the population matches the opposite-favoring behavior of the data. Despite large variances in model parameters (Fig. S6A-C), perturbation responses of all the fitted models closely match experimental data (Fig. 6C, D). On average, the fitted models display opposite-favoring responses across almost the entire range of experimentally measured distances (Fig. 6E), consistent with the findings of (1).

**Fig. 6.**
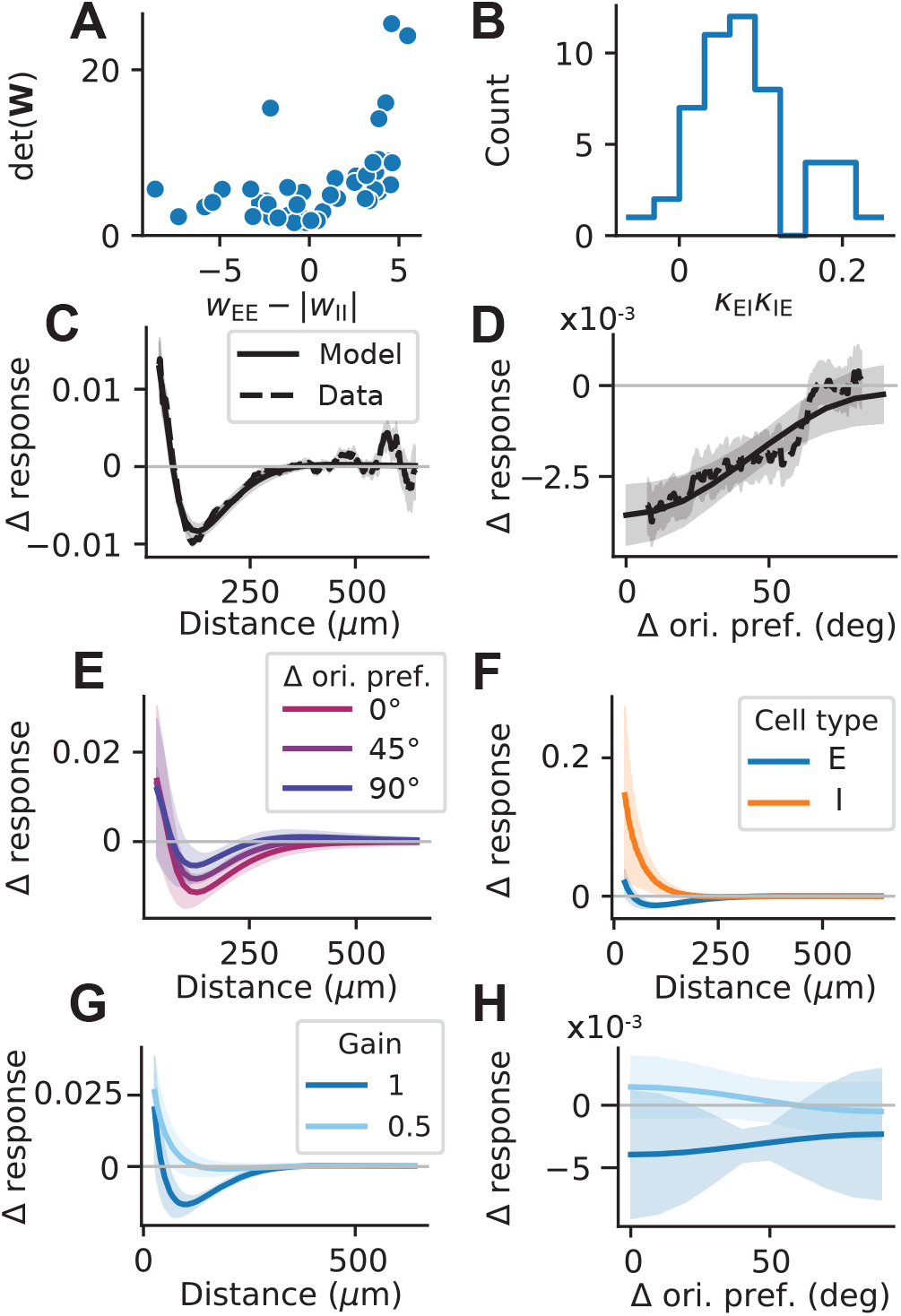
Validation of theoretical insights in fitted models. 200 models are fitted to the single cell-perturbation response curve as a function of distance from (1), the top 50 of which are plotted. **A)** Distribution of the fitted model parameters, where each point is a fitted model. **B)** Histogram of the product of fitted parameters *κ*_EI_*κ*_IE_, which is positive if and only if disynaptic E → I → E inhibition is like-to-like. **C-E)** Perturbation response of all neurons in the model (including E and I) **C-D)** Comparison between the perturbation response of the fitted models and experimental data. Error bars of data represents standard error. Error bars of model represents standard deviation across fitted models. To match the data analysis procedure of (1), bin widths of 60 µm for C and 25^*°*^ for D are used. **E)** Perturbation response of fitted models as a function of distance to the perturbed neuron, for different tuning preferences. Same bin width as C). **F-H)** Simulations support analytical predictions. Smaller bin widths than C-E are used for more accurate results (2 µm bins for F, G and 10^*°*^ bins for H). **F)** Comparison between excitatory and inhibitory neuron response in fitted models. **G-H)** Effect of reducing neuronal gain on the responses of excitatory neurons. Models are fitted with a gain of 1. Note that this shows the response of E neurons rather the average response of all neurons, *i*.*e*. including both E and I neurons, as in C-E.

We then test three of our theoretical predictions on these fitted models: 1) inhibitory neurons should exhibit a broader perturbation response profile than excitatory neurons (Fig. 6F), 2) when overall neuronal gain is lowered, excitatory neuron response should be broader and less suppressed (Fig. 6G), and 3) when overall neuronal gain is sufficiently weak, excitatory neuron response transitions from opposite-favoring to same-favoring (Fig. 6H). These predictions hold true in all the fitted models, despite the large variances in model parameters and despite the fact that these models violate the symmetry assumptions in our theory, suggesting that these are robust effects that can be expected from experimental measurements. Finally, the connectivity matrices of all fitted models have spectral radius larger than 1 (Fig. S6D), suggesting that biological networks are indeed in a strongly-coupled regime where the typical analytical approximation 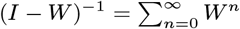 fails to apply.

All fitted models are fully connected, which raises the question of whether our theory also applies to models with realistically sparse connectivity. To test this, we construct a sparsely connected version of each fitted model with parameters chosen to match available data (Fig. S7A-D; Materials and Methods). In simulations, perturbation responses of the sparse models closely match those of the fully connected models (Fig. S7E-J). Thus, our theory captures the average behavior of these realistically sparse models.

We have assumed that neuronal gain is uniform across all neurons of a given cell type. However, presentation of a large drifting grating, as in Ref. (1), preferentially increases the firing rate of, and thus the gain of, iso-tuned neurons, with firing rates decaying slowly with distance from the stimulus center. To understand how this may affect our results, we simulate our fitted models with inhomogeneous space- and/or feature-dependent gain (Materials and Methods). We find that a slow decay of neuronal gain with distance has little effect on perturbation response (Fig. S8A). Similarly, feature dependence of neuronal gain causes excitatory neuron responses to be only slightly less opposite-favoring on average over the fitted models (Fig. S8B,C), though some particular models change to a same-favoring response (Fig. S8D).

## Discussion

In this paper, we developed novel theory for understanding the link between recurrent connectivity structure and single-cell optogenetic perturbation responses. We introduced an exponential-type kernel for describing connectivity as a function of distance that for the first time allows an exact solution for a space- and feature-dependent linear network that is valid in all coupling regimes. We showed that this kernel can well capture the spatial dependence of the connectivity in the data, defined as the product of the connection probability and its strength, and used this to exactly solve for the network’s steady-state response to a single-cell perturbation.

Analysis of the solution for E-I ISN networks revealed five main results. (1) A positive determinant of the 2 × 2 connectivity weight matrix ***W*** is necessary (assuming inhibitory projections are narrower than excitatory) to explain experimental observations of a perturbation response that is excitatory for nearby cells and suppressive at larger distances. The larger the determinant, the shorter the spatial radius of nearby excitation. (2) The response at larger distances can either remain negative or be oscillatory in space, and spatial oscillation frequency is negatively correlated with the strength of I →I connections. (3) We predict that the spatial profile of the perturbation responses of inhibitory neurons is qualitatively like that of excitatory neurons, but with a larger spatial radius of nearby excitation than for excitatory neurons. (4) Feature-specific disynaptic inhibition (E →I → E) that is like-to-like (*i*.*e*., that couples neurons with similar preferred features) is necessary to explain observations of “opposite-favoring” responses, *i*.*e*. that neurons with feature preferences opposite to a perturbed neuron are less suppressed or more excited on average than neurons with similar feature preferences. In fact, such a like-to-like connectivity motif is necessary if the perturbation response is opposite-favoring at any distance. (5) We predict that a decrease in neuronal gain would cause perturbation response to be less suppressive and have a broader spatial radius of excitation, and that the response becomes same-favoring rather than opposite-favoring for sufficiently weak neuronal gain. All of these analytic results except the fourth were obtained on the assumption that connectivity width depends only on presynaptic cell type, but they hold in simulations without this assumption.

To the best of our knowledge, this is the first exactly solvable model of a recurrent network with space- and feature-dependent recurrent connectivity. It is straight-forward to obtain an exact analytic solution of a linear translation-invariant model in Fourier space, but in general this cannot be inverted to obtain responses as a function of distance. Nonetheless, previous works were able to obtain some information analytically, *e*.*g*. using the Fourier-space solutions to compute the spatial resonant frequencies of the network, from which experimental predictions were made (14). Alternatively, one may obtain an approximate expression for the steady-state solution by assuming that all activity patterns have a Gaussian shape (19, 34), although this assumption, typically applied to visual responses, may not be suitable for describing single-cell perturbation responses. Ref. (15) obtained an exact steady-state solution for an E-I network with a Gaussian spatial connectivity kernel in the tightly balanced regime (15). In this regime, there is a precise cancellation between excitatory and inhibitory synaptic input currents such that *W* |*r*⟩ + |*h*⟩ ≈ 0, so the steady state solution can be approximated as |*r*⟩ ≈ −*W*^−1^ |*h*⟩. However, experimental evidence suggests that the cortex is in a loosely balanced rather than a tightly balanced regime (35), and our model is valid in both regimes.

The Green’s function kernel we introduced for modeling the spatial dependence of connectivity simplifies to an exponential kernel 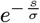 for a 1D network, which has previously studied by (36). Compared to the Gaussian kernel typically used for modeling V1 connectivity (14–20), it has a sharp peak at short distances, which grows sharper in 2D and 3D. It was recently noted that the short spatial radius of nearby excitation in perturbation responses cannot be explained by models with spatial connectivity given by a single Gaussian kernel with realistic length scales, and that a sharp peak must be added to the connectivity kernel to explain the data (2). Our spatial kernel satisfies this condition. Surprisingly, given an exponential-type connectivity kernel, a narrow perturbation response does not necessitate a narrow spatial connectivity kernel, and, conversely, a narrow spatial connectivity kernel does not imply a narrow perturbation response. Instead, for fixed connectivity widths, the spatial radius of nearby excitation can vary over several orders of magnitude depending on the mean connectivity strengths between different cell types (Fig. S1).

Our analysis of both the mean and the spatial profile of perturbation response strongly suggest det(***W***) is positive, *i*.*e*. disynaptic E →I →E inhibition is stronger than the product of E →E and I →I connections. This has important implications for the dynamics of a nonlinear E-I network. Because the linearized dynamics of a nonlinear network around a fixed point is driven by an effective connectivity matrix ***W*** equal to the product of the connectivity ***J*** and a diagonal matrix of (positive) neuronal gains, the determinant of ***W*** and ***J*** have the same sign. Thus, our insight that the determinant of the connectivity matrix ***W*** of the linearized network is positive also implies det(***J***) *>* 0. Theoretical work on networks with power-law input/output functions has shown that the condition det(***J***) *>* 0 guarantees dynamical stability assuming sufficiently fast inhibition (37), and plays an important role in determining aspects of neural dynamics such as bistability, persistent activity, and oscillations (38). Sadeh and Clopath (24) studied the conditions to obtain a suppressive, opposite-favoring mean perturbation response, and also concluded that disynaptic E →I →E connections must be sufficiently strong and like-to-like. Our results extend theirs in several ways. First, we describe the spatial dependence of the response and not only the mean response. Second, we do not assume inhibitory connection strength vastly exceeds E → E strength, and we consider the possibility of like-to-unlike I →I connections. Finally, we note that the single-cell perturbation response data we both are modeling (1) excludes neurons within 25 µm of the perturbed cell in lateral distance. Excluding the strongly excited nearest neurons likely underestimates the mean perturbation response, making analyses based solely on that mean unreliable. By instead examining responses as a function of distance, we show disynaptic E →I → E connections must be sufficiently strong and like-to-like independently of the mean perturbation response.

We inferred parameters of the connectivity from optogenetic perturbation responses based on an explicit expression we derived for response vs. distance and feature preference. Our approach is distinct from the works of (39–45), who inferred individual synaptic connections from whole-cell recordings of postsynaptic currents in response to perturbations of specific cells, based on models of monosynaptic intracellular responses. Other efforts to infer mean, and in some cases variance, of connectivity from responses to visual stimuli (17, 19, 28, 46–48) were either based on fitting by extensive search, or by comparison to expressions for responses that ignored space and/or feature dependence.

There are a few important future directions for our work. First, some of our analysis is restricted to networks with only a single inhibitory cell type and whose connectivity width depends only on presynaptic cell type. Without either restriction, our linear response equation would be composed of a sum over more than two spatial terms, making it difficult or impossible to precisely characterize spatial zero-crossings in the perturbation response. Second, we only considered models with dependence on a single feature. Mathematically it is straightforward to generalize our steady state solution to include an arbitrary number of periodic feature dependencies, but it is unclear how non-periodic features such as spatial and temporal frequency can be incorporated. Third, feature tuning in our connectivity is parametrized by a cosine function, which fixes the feature tuning width. It is important to investigate whether the theory can be adapted for other choices of feature tuning kernel that allow for variable feature tuning width. Finally, we have so far only dealt with single-cell perturbations in a linear network. Linearity is a reasonable approximation since a moderate single-cell perturbation is unlikely to generate significant nonlinear effects. However, many optogenetic experiments perturb an ensemble of neurons (2–5), or use one-photon methods to perturb large numbers of neurons and/or consider the combination of sensory and optogenetic stimuli (*e*.*g*., 20, 28, 49), in which case nonlinear effects cannot be ignored. Furthermore, in nonlinear networks one also needs to consider the effects of connectivity disorder, which would both modify the mean perturbation response and potentially result in chaotic dynamics (20, 50). Thus, it is important to extend our work to consider nonlinear contributions to perturbation response.

## Materials and Methods

### Mathematical notation

Throughout the paper, scalar variables are represented by lowercase letters like *s, k*, vectors by boldface lowercase letters such as ***x, k***, and matrices by boldface uppercase letters such as **Σ, *W***. Given a matrix ***W***, its elements are written as *W*_*ij*_ or [***W***]_*ij*_, where the first notation is preferred whenever possible. Linear operators on vector spaces except ℝ ^*n*^ are represented by uppercase letters such as *W, L, T*. Vectors in vector spaces except ℝ^*n*^ are expressed using the standard bra-ket notation (see SI section 1A for more information), where |*v*⟩ represents a vector with label *v*. For example, in Eq. 4, |*r*⟩ represents the vector of the functions *r*_*α*_(***x***) for all cell types *α*.

### Model setup details

In our model, external input due to the perturbation of a single neuron (*β, ν*, ***y***, *ϕ*) is given by

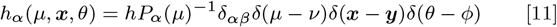

where *h* is a scalar representing the perturbation strength, and the prefactor *P*_*α*_(*µ*)^−1^ ensures that the total input to the network 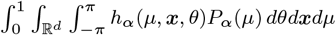 is independent of the feature selectivity of the perturbed neuron *ν*. We assume that the synaptic timescale of each neuron is only dependent on its cell type. Thus, the dynamical equation of our model is given by

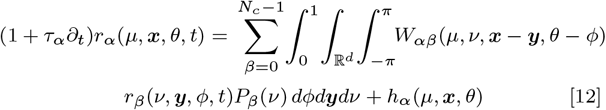

where *τ*_*α*_ is the time constant for cell type *α*. Stability of the network dynamics in general depends on the specific time constants chosen for each cell type. To simplify our analysis, however, we assume that the time constant of inhibitory neurons is sufficiently fast, such that the stability of the network dynamics depends only on the connectivity parameters (SI section 7C).

### Fitting of the spatial connectivity kernel

To compute the product of connection probability data from (11) and connection strength data from (13) in Fig. 1B, we assume that E →I connection probability is the same as I →E since only I →E connection probability is measured in (11). This assumption is supported by another dataset which shows that I →E and E →I connection probabilities have approximately the same spatial spread (12). Given that connection strength is only measured between neurons up to about 100 µm apart in the data from (13), we assume that the connection strength for all distance bins in which no data is available is equal to the connection strength in the last bin in which data is available. The spatial kernel being fitted to this product is given by Eq. 3 with *d* = 2. For a given pair of post- and pre-synaptic cell types *α, β*, there are two free parameters: *w*_*αβ*_ and *σ*_*β*_. These parameters are fitted with non-linear least squares. For further details, see the SI section 1B.

### Estimation of 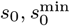 from data

The 95% confidence intervals for 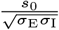 and 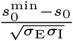 in Fig. 2D and 2E are estimated via bootstrapping. We independently sample each data point of the single-cell perturbation response curve in (1, Fig. 2G) from a Gaussian distribution with its mean and standard error to obtain a random sample of the single-cell perturbation response curve. The confidence intervals are then computed from 100,000 random sample curves. For further details, see SI section 1C. This yields 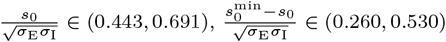.

### Computation of 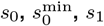, and their derivatives in models

The quantities 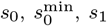, and the derivatives *∂*|*w*_II_|(*s*_1_ − *s*_0_), *∂* _*g*_ *s* _0_ in Figures 2D-F, 3B and 5C do not have closed form expressions in networks with 2 spatial dimensions and thus are evaluated numerically. *s*_0_, *s*^min^, and 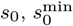 are roots of transcendental equations and are solved with the bisection method (SI section 1D). The derivatives of these quantities are computed numerically by perturbing |*w*_II_| and *g* by 0.001%.

### Comparison of theory and simulations

Here we describe the models used in Figures 1D, 2C, 4A, 4C, 5A, and explain how the theory and simulation curves are obtained. The connectivity function of all these models are described by Eq. 8, with the factor of 2*π* replaced by *π* and cos(*θ − ϕ*) replaced by cos(2(*θ − ϕ*)) since we specifically consider orientation tuning preference which is a variable in 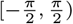 rather than [−*π, π*). OSI-dependence of the connectivity is linear, *i*.*e*. we set *f*_*α*_(*µ*) = *g*_*α*_(*µ*) = *µ* for all cell types *α*, and we assume a uniform distribution of neuronal OSI, *i*.*e. P*_*α*_(*µ*) = 1 for all *α*. In all figures, a neuron with OSI of 1 is perturbed. For other model parameters, see SI section 1E.

For numerical simulations, the model is discretized on a regular grid with *N*_*x*_ = 100 by *N*_*y*_ = 100 spatial locations on a 1 mm × 1 mm torus (*d* = 2), *N*_*θ*_ = 12 feature tuning preferences, and *N*_*µ*_ = 7 feature selectivities. Spatial distances between neurons are measured by toroidal distances. The discretized model connectivity is obtained by multiplying the connectivity function Eq. 8 by 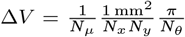. To deal with the divergence of the spatial connectivity kernel *G*_*d*_(*s*; *λ*) for *d* ≥ 2 as *s* →0, we simply set the connectivity strength between neurons at the exact same spatial location to 0. This procedure is justified in SI section 16. In other words, the discretized connectivity matrix ***W*** ^dis^ is defined by

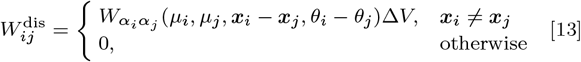

where *α*_*i*_, *µ*_*i*_, ***x***_*i*_, *θ*_*i*_ are the cell type, selectivity, spatial location, and feature preference respectively of neuron *i* ∈ {1, · · ·, *N*} in the model, and *N* := *N*_*c*_*N*_*µ*_*N*_*x*_*N*_*y*_*N*_*θ*_. Perturbation strength is set to *h* = 10000, a large number to avoid numerical issues arising from a small perturbation response. For further details see SI section 1E. The analytical solutions for Fig. 1D and 4C are given by Eq. 9, with the factor of 2*π* replaced by *π* and cos(*θ* − *ϕ*) replaced by cos(2(*θ* − *ϕ*)). They are also scaled by Δ*V* (see SI section 16 for an explanation). Specifically, our analytical solution for the response of neuron *i* to the perturbation of neuron *k* in the discretized model is computed as

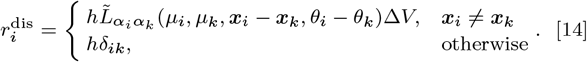

For Fig. 2C, since only the response averaged over feature tuning is plotted, we use the integral of Eq. 9 over feature tuning (Eq. S51) for our analytical solution with Δ*V* = (1 mm^2^)(*N*_*µ*_*N*_*x*_*N*_*y*_*N*_*θ*_)^−1^. Similarly, for Fig. 4A, we use the integral of Eq. 9 over space (Eq. S54) with Δ*V* = *π*(*N*_*µ*_*N*_*x*_*N*_*y*_*N*_*θ*_)^−1^, and for Fig. 5A, we use the integral of Eq. 9 over both space and feature tuning (Eq. S90) with Δ*V* =(*N*_*µ*_*N*_*x*_*N*_*y*_*N*_*θ*_)^−1^.

### Estimation of *κ*_EE_ from data

For the models fitted to experimental data in Fig. 6 and S7, the parameter *κ*_EE_ was constrained by fitting the curve 1 + 2*κ*_EE_ cos(2*θ*) (with an arbitrary scaling factor) to mouse V1 L2/3 connection probability data from (11, Fig. 2H) This yield a best-fit value of *κ*_EE_ = 0.198 ± 0.054 (inset of Fig. S7B). For more details of the fitting see SI section 1F.

### Model fitting to experimental data

The models in Fig. 6 consist of *N* = 24,000 neurons on a 900 µm × 900 µm plane (*d* = 2). The cell type, spatial location, and orientation tuning preference of each neuron is randomly assigned, such that each neuron has 0.85 chance of being an excitatory neuron and 0.15 chance of being an inhibitory neuron, while spatial locations and tuning preferences are uniformly distributed. Tuning selectivity is omitted in models by setting *P*_*α*_(*µ*) = *δ*(*µ* − 1), *i*.*e*. every neuron is perfectly tuned. Inhibitory time constant is set to be half of excitatory, *i*.*e*. 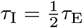.

Model parameters are simultaneously fitted to both experimental data curves in Fig. 6C, D (SI section 1G). There are 13 parameters for fitting: 4 connectivity strengths *w*_*αβ*_, 4 connectivity widths *σ*_*αβ*_, 4 feature tuning parameters *κ*_*αβ*_, and the perturbation strength *h*. Since *h* does not affect the shape of the response curve and only affects its overall amplitude, we eliminate this parameter by normalizing both the model perturbation response as well as the data to unit norm during fitting. 8 constraints (described in SI section 1G) are imposed on the remaining 12 parameters during optimization to ensure realistic model parameters.

Once the optimization algorithm has converged, a normalized validation loss is computed by comparing the results of 50 numerical simulations to data (SI section 1G). We discard the fitted parameters if the validation loss is greater than 0.75. A distribution of fitted model parameters is obtained by bootstrapping. Specifically, the model parameters are fitted to random samples of the perturbation response curves from the data (SI section 1G). The optimization procedure is repeated with different random samples of the perturbation response curve, different random initializations of model parameters, and different random instantiations of cell types, spatial locations, and feature preferences until 200 sets of fitted parameters are obtained, of which the top 50 are analyzed.

### Models with sparse connectivity/inhomogeneous gain

Please see SI section 1H/1I for details.

## Data, Materials, and Software Availability

Code for reproducing all figures is available at https://github.com/hchau630/chau-2024-exact.

## ACKNOWLEDGMENTS

We thank Christopher Harvey and Selmaan Chettih for providing data from (1) for Fig. 6C, D, and Petr Znamenskiy for providing data from (13) for Fig. 1B. We thank Tuan Nguyen and Mora Ogando for helpful discussions. This work is supported by the Gatsby Charitable Foundation (GAT3708; and GAT3850 to A.P), the Kavli Foundation, the NSF (DBI-1707398; and DGE-2036197 to H.Y.C.), the NIH (U01 NS108683z, R01 EY029999, U19 NS107613; and T32 EY013933 to H.Y.C.), and the Simons Foundation (1156607, to A.P).

