## Supporting Information for "Exact linear theory of perturbation response in a space- and feature-dependent cortical circuit model"

##### This PDF file includes:

Supporting text
Figs. S1 to S8
Tables S1 to S4
SI References

### Supporting Information Text

#### 1. Extended methods

**A. Mathematical notation.** In this paper we use the standard bra-ket notation to represent vectors and linear operators on vector spaces other than $\mathbb{R}^n$. In this notation, $|v\rangle$ represents a vector in a Hilbert space with label $v$. $\langle v|$ is the linear functional in the dual space associated with $|v\rangle$ such that $\langle v|(|u\rangle) = (|v\rangle, |u\rangle)$, where $(\cdot,\cdot)$ is the inner product on the Hilbert space. We write $\langle v \,|\, u\rangle$ to denote $\langle v|(|u\rangle)$. Similarly, given an operator $T$, we write $\langle v|T|u\rangle$ to denote $\langle v|(T|u\rangle)$. Given vectors $|v\rangle, |u\rangle$ in vector spaces $V, U$ respectively, the vector $|v, u\rangle$ represents the vector $|v\rangle \otimes |u\rangle$ in the tensor product space $V \otimes U$.

Given cell type index $\alpha \in \mathbb{Z}_{N_c}$, we write $|\alpha\rangle$ to represent the standard basis vector $\boldsymbol{e}_\alpha \in \mathbb{R}^{N_c}$. Given vector $\boldsymbol{y} \in \mathbb{R}^d$, we write $|\boldsymbol{y}\rangle$ to represent the Dirac delta 'function' $\delta(\boldsymbol{x} - \boldsymbol{y})$. Similarly, given $\phi \in \mathbb{S}^1$, we write $|\phi\rangle$ to represent the Dirac delta 'function' $\delta(\theta - \phi)$ on the circle.

**B. Fitting of the spatial connectivity kernel.** Binning of connection probability and connection strength data is performed with bin edges from 0 to 500 µm spaced 25 µm apart. The parameters $w_{\alpha\beta}$ and $\sigma_\beta$ are fitted using the `optimize.curve_fit` function in the `scipy` Python library (1), which performs non-linear least squares. $\sigma_\beta$ is initialized at 100 µm and $w_{\alpha\beta}$ is initialized to match the 2-norm of the data vector. Uncertainty of the fitted parameters is obtained from the default output of the `curve_fit` function, which estimates the covariance of fitted parameters by a linear approximation. This results in the best-fit parameters $\sigma_\mathrm{E} = (150.2 \pm 11.3)$ µm and $\sigma_\mathrm{I} = (107.6 \pm 8.4)$ µm.

**C. Estimation of $s_0, s_0^{\min}$ from data.** Here we describe more details of our bootstrapping procedure for computing the 95% confidence interval of $\frac{s_0}{\sqrt{\sigma_\mathrm{E}\sigma_\mathrm{I}}}$ and $\frac{s_0^{\min} - s_0}{\sqrt{\sigma_\mathrm{E}\sigma_\mathrm{I}}}$. For each of the randomly sampled single-cell perturbation response curves, we compute $s_0$ by linearly interpolating between the first two consecutive data points which exhibit a sign change. However, this would introduce a slight bias towards a smaller $s_0$ since the sampled curve may exhibit multiple crossings around $s_0$ and we are taking the first crossing. To address this bias we filter out all sampled curves with more than one crossing within 100 µm. We compute $s_0^{\min}$ as the location of the minimum of the sampled curve. However, the large standard errors in the data at large distances creates spurious minima in the sampled curve and thus introduces a small bias towards larger $s_0^{\min}$. To address this we simply consider the minimum of the sampled curve within 300 µm. We repeat the above procedures to obtain 100,000 samples of $s_0$ and $s_0^{\min}$. Finally, we divide each sample of $s_0$ and $s_0^{\min}$ by an independent sample of $\sqrt{\sigma_\mathrm{E}\sigma_\mathrm{I}}$ using the mean and uncertainty of $\sigma_\mathrm{E}$ and $\sigma_\mathrm{I}$ as estimated in the previous subsection.

**D. Computation of $s_0$, $s_0^{\min}$, $s_1$ in the model.** Here we describe how to use the bisection method to accurately compute the quantities $s_0$, $s_0^{\min}$, and $s_1$ for different choices of model parameters. For a model with $d$-spatial dimensions, the zero crossings $s_0$, $s_1$ of the response of cell type $\alpha$ to the perturbation of cell type $\beta$ are given by the first two roots of the equation $\frac{G_d(s;\lambda_1)}{G_d(s;\lambda_0)} = c_{\alpha\beta}$, where $\lambda_0, \lambda_1$ are given by Eq. S95, S96 and $c_{\alpha\beta}$ is given by Eq. S99. $s_0^{\min}$ is given by the first root of the equation $\frac{G_{d+2}(s;\lambda_1)}{G_{d+2}(s;\lambda_0)} = c_{\alpha\beta}$ (SI section 12).

To use the bisection method we need to establish a lower bound and an upper bound of each of these quantities. This is simple when the network is in the phase region where there is only 1 zero crossing for the chosen cell types $\alpha, \beta$ (in which case $s_1$ is undefined), but when there are infinitely many zero crossings we need to ensure that the interval defined by our bounds contains only the root that we desire. To address this we establish the following inequality:

$$0 < s_0^{(2)} < s_0^{(3)} < s_0^{(4)} < s_1^{(2)} < s_1^{(3)} < s_1^{(4)} < s_2^{(2)} < \cdots \qquad \text{[S1]}$$

where $s_n^{(d)}$ represents the $(n+1)$-th root of $\frac{G_d(s;\lambda_1)}{G_d(s;\lambda_0)} = c_{\alpha\beta}$ (assuming there are infinitely many roots). To see this, note that since $s_n^{(2)}$ is the $n$-th zero crossing of Eq. S94 for $d = 2$ and $s_n^{(4)}$ is the $n$-th critical point of the same equation (by the same argument in SI section 12), we have $s_n^{(2)} < s_n^{(4)} < s_{n+1}^{(2)}$ for all $n$. But now if we allow $d$ to be any arbitrary real number and consider the derivative of $s_n^{(d)}$ with respect to $d$, we find that $\partial_d s_n^{(d)} > 0$ (Theorem 17.20), so we have $s_n^{(2)} < s_n^{(3)} < s_n^{(4)}$. Combining these two inequalities then yields Eq. S1. Now, because a closed-form analytical expression for $s_n^{(3)}$ exists (Eq. S130), this inequality implies that for a network with $d = 2$ spatial dimensions, we can choose our bounds for $s_0$ (which is equal to $s_0^{(2)}$) to be $(0, s_0^{(3)})$, and the bounds for $s_0^{\min}$ and $s_1$ (which are equal to $s_0^{(4)}$ and $s_1^{(2)}$ respectively) to be $(s_0^{(3)}, s_1^{(3)})$. If the network is in a regime with only 1 zero crossing for the chosen cell types $\alpha, \beta$, then $s_1^{(3)}$ is undefined, so we modify the upper bound for $s_0^{\min}$ to be $2s_0^{(3)}$ (this is just a heuristic that seems to work well within the phase regions that we explored).

$s_n^{(d)}$ is then numerically computed by applying the bisection method on the function

$$f_d(s) := \mathrm{Re}(G_d(s;\lambda_1) - c_{\alpha\beta}G_d(s;\lambda_0))$$

with the bounds we just found. There is just one little additional detail that we need to verify: clearly all roots of $G_d(s;\lambda_1) - c_{\alpha\beta}G_d(s;\lambda_0)$ are roots of $f_d(s)$, but it is not obvious that all roots of $f_d(s)$ are roots of $G_d(s;\lambda_1) - c_{\alpha\beta}G_d(s;\lambda_0)$ when $\lambda_0, \lambda_1, c_{\alpha\beta}$ are complex numbers (or equivalently, when the equation has infinitely many roots). To see that this is indeed the case, we note that when there are infinitely many zero crossings, $\lambda_1 = \overline{\lambda}_0$ and $c_{\alpha\beta} = -\frac{z^+_{\alpha\beta}}{\overline{z}^+_{\alpha\beta}}$, where $z^+_{\alpha\beta}$ is the numerator of the definition of $c_{\alpha\beta}$ in Eq. S97. Thus $f_d(s)$ can be written as

$$f_d(s) = \mathrm{Re}\left(\frac{\overline{z}^+_{\alpha\beta}G_d(s;\overline{\lambda}_0) + z^+_{\alpha\beta}G_d(s;\lambda_0)}{\overline{z}^+_{\alpha\beta}}\right) = \mathrm{Re}\left(\frac{\overline{z^+_{\alpha\beta}G_d(s;\lambda_0)} + z^+_{\alpha\beta}G_d(s;\lambda_0)}{\overline{z}^+_{\alpha\beta}}\right) = \frac{\overline{z^+_{\alpha\beta}G_d(s;\lambda_0)} + z^+_{\alpha\beta}G_d(s;\lambda_0)}{\mathrm{Re}(\overline{z}^+_{\alpha\beta})}$$

where the second equality follows from Lemma 17.6. Thus if $s^*$ is a root of $f_d$, then $\overline{z^+_{\alpha\beta}G_d(s^*;\lambda_0)} + z^+_{\alpha\beta}G_d(s^*;\lambda_0) = 0$. But since we also have

$$G_d(s;\lambda_1) - c_{\alpha\beta}G_d(s;\lambda_0) = \frac{\overline{z^+_{\alpha\beta}G_d(s;\lambda_0)} + z^+_{\alpha\beta}G_d(s;\lambda_0)}{\overline{z}^+_{\alpha\beta}}$$

this means $s^*$ is also a root of $G_d(s;\lambda_1) - c_{\alpha\beta}G_d(s;\lambda_0)$, proving our claim.

**E. Comparison of theory and simulations.** Connectivity strengths for Figures 1D, 4A, 4C, 5A are given by the following: $w_{\mathrm{EE}} = 3, |w_{\mathrm{EI}}| = w_{\mathrm{IE}} = 4, |w_{\mathrm{II}}| = 5.25$ for Figure 1D, $w_{\mathrm{EE}} = 1, |w_{\mathrm{EI}}| = w_{\mathrm{IE}} = 4, |w_{\mathrm{II}}| = 0$ for Figure 4A, $w_{\mathrm{EE}} = 1.5, |w_{\mathrm{EI}}| = w_{\mathrm{IE}} = 3, |w_{\mathrm{II}}| = 5$ for Figure 4C, and $w_{\mathrm{EE}} = 2, |w_{\mathrm{EI}}| = w_{\mathrm{IE}} = \sqrt{3}, |w_{\mathrm{II}}| = 1$ for Figure 5A. Connectivity widths for Figures 1D, 4A, 5A are given by $\sigma_{\mathrm{EE}} = 125\,\mu\mathrm{m}, \sigma_{\mathrm{EI}} = 90\,\mu\mathrm{m}, \sigma_{\mathrm{IE}} = 85\,\mu\mathrm{m}, \sigma_{\mathrm{II}} = 110\,\mu\mathrm{m}$, while connectivity widths for Figure 2C and Figure 4C are given by $\sigma_{\alpha\beta} = 40\,\mu\mathrm{m}$ and $\sigma_{\alpha\beta} = 100\,\mu\mathrm{m}$ respectively. Feature tuning parameters for Figures 1D and 5A are given by $\kappa_{\mathrm{EE}} = 0.5, \kappa_{\mathrm{EI}} = \kappa_{\mathrm{IE}} = -0.25, \kappa_{\mathrm{II}} = 0.25$, while tuning parameters for Figure 2C are given by $\kappa_{\mathrm{EE}} = 0.2, \kappa_{\mathrm{EI}} = \kappa_{\mathrm{IE}} = 0.1, \kappa_{\mathrm{II}} = 0$. Inhibitory neuron time constant is set to be half of excitatory neuron time constant ($\tau_{\mathrm{I}} = \frac{1}{2}\tau_{\mathrm{E}}$) except for Figure 1D, for which $\tau_{\mathrm{I}} = \tau_{\mathrm{E}}$, and Figure 2C, for which $\tau_{\mathrm{I}} = \frac{1}{4}\tau_{\mathrm{E}}$. Note that excitatory and inhibitory time constants are set to be equal in Figure 1D because the stability region plotted in Figure 1C (corresponding to $\mathrm{Re}(\lambda) < 1$) assumes equal time constants. The rest of the model parameters are given by Tables S1, S2, S3.

For simulations, network dynamics given by the discretized version of Eq. 12 in the main text is numerically integrated with order-5 Dormand-Prince method using the `torchdiffeq` package until convergence to steady state (2, 3), which is numerically determined by the condition $\left|\frac{\mathrm{d}r_i}{\mathrm{d}t}\right| \leq \epsilon_{\mathrm{rel}}|r_i| + \epsilon_{\mathrm{abs}}$ being satisfied for all $i \in \{1, \cdots, N\}$, where $r_i$ is the firing rate of neuron $i$, $\epsilon_{\mathrm{abs}} = 10^{-6}$, and $\epsilon_{\mathrm{rel}} = 10^{-5}$, except for Figure 5A in which $\epsilon_{\mathrm{rel}} = 5 \times 10^{-5}$.

**Table S1. Connectivity strengths for Figure 2C**

| # of crossings | Parameter | Value | # of crossings | Parameter | Value | # of crossings | Parameter | Value |
|---|---|---|---|---|---|---|---|---|
| 0 | $w_{\mathrm{EE}}$ | 1.5 | 1 | $w_{\mathrm{EE}}$ | 1.5 | $\infty$ | $w_{\mathrm{EE}}$ | 4.5 |
| | $\lvert w_{\mathrm{EI}}\rvert$ | 2 | | $\lvert w_{\mathrm{EI}}\rvert$ | 3 | | $\lvert w_{\mathrm{EI}}\rvert$ | 3 |
| | $w_{\mathrm{IE}}$ | 2.5 | | $w_{\mathrm{IE}}$ | 2.5 | | $w_{\mathrm{IE}}$ | 2 |
| | $\lvert w_{\mathrm{II}}\rvert$ | 4 | | $\lvert w_{\mathrm{II}}\rvert$ | 4 | | $\lvert w_{\mathrm{II}}\rvert$ | 0 |

**Table S2. Feature tuning parameters for Figure 4A**

| Kind | Parameter | Value | Kind | Parameter | Value |
|---|---|---|---|---|---|
| Same-fav. | $\kappa_{\mathrm{EE}}$ | 0.5 | Opp.-fav. | $\kappa_{\mathrm{EE}}$ | 0.5 |
| | $\kappa_{\mathrm{EI}}, \kappa_{\mathrm{IE}}$ | 0.5 | | $\kappa_{\mathrm{EI}}, \kappa_{\mathrm{IE}}$ | 0.25 |
| | $\kappa_{\mathrm{II}}$ | 0 | | $\kappa_{\mathrm{II}}$ | 0 |

**F. Estimation of $\kappa_{\mathrm{EE}}$ from data.** We perform non-linear least squares using the `optimize.curve_fit` function in the `scipy` Python library (1) to fit the parameters $a, \kappa_{\mathrm{EE}}$ of the curve $\theta \mapsto a(1 + 2\kappa_{\mathrm{EE}}\cos(2\theta))$ to the connection probability data from (4, Figure 2H). Each data point is weighted inversely proportional to its standard error. The parameter $a$ is initialized as the mean connection probability, while $\kappa_{\mathrm{EE}}$ is initialized at 0. $\kappa_{\mathrm{EE}}$ is constrained to be between $-0.5$ and 0.5. Uncertainty of the parameters is taken from the output of `curve_fit`.

**Table S3. Feature tuning parameters for Figure 4C**

| # of transitions | Parameter | Value | # of transitions | Parameter | Value |
|---|---|---|---|---|---|
| 0 | $\kappa_{\mathrm{EE}}$ | 0.15 | 1 | $\kappa_{\mathrm{EE}}$ | 0.15 |
| | $\kappa_{\mathrm{EI}}$ | 0.3 | | $\kappa_{\mathrm{EI}}$ | 0.5 |
| | $\kappa_{\mathrm{IE}}$ | 0.15 | | $\kappa_{\mathrm{IE}}$ | 0.4 |
| | $\kappa_{\mathrm{II}}$ | 0.5 | | $\kappa_{\mathrm{II}}$ | 0.5 |

**G. Model fitting to experimental data.** Here we describe more details about our model fitting procedure. First, to avoid numerical issues due to the divergence of the spatial kernel $G_d(s;\lambda)$ as $s \to 0$, we require that all pairwise distances between neurons be at least 3 µm. This is achieved by resampling the spatial location of one of the neurons from each pair of neurons whose pairwise distance is less than 3 µm, and repeating until the requirement is satisfied. During model fitting, a random excitatory neuron within a 680 µm × 680 µm window centered at the origin of the 900 µm × 900 µm simulation region is chosen for perturbation. During model fitting, the steady-state response of the network is computed using the analytical solution. Specifically the steady state response of neuron $i$ to perturbation of neuron $k$ is computed as

$$r_i^{\mathrm{dis}} = \begin{cases} h\tilde{L}_{\alpha_i\alpha_k}(1,1,\boldsymbol{x}_i - \boldsymbol{x}_k, \theta_i - \theta_k)\Delta V_{\alpha_k}, & \boldsymbol{x}_i \neq \boldsymbol{x}_k \\ h\delta_{ik}, & \text{otherwise} \end{cases} \qquad \text{[S2]}$$

where $\Delta V_\alpha = \frac{1\,\mathrm{mm}^2 \cdot \pi}{p_\alpha N}$, $\alpha_i, \boldsymbol{x}_i$ are the cell type and spatial location of neuron $i$ respectively, $p_\alpha$ is the probability that a given neuron is of cell type $\alpha$ (thus $p_{\mathrm{E}} = 0.85$ and $p_{\mathrm{I}} = 0.15$), and $\tilde{L}_{\alpha\beta}$ is the linear response equation Eq. 9 with the factor of $2\pi$ replaced by $\pi$ and $\cos(\theta-\phi)$ replaced by $\cos(2(\theta-\phi))$. Neurons beyond a 680 µm × 680 µm window centered at the origin are excluded to mimic the field-of-view of the experiment as well as to minimize boundary effects. Following the data analysis procedure of (5), neurons within 25 µm of the perturbed neuron are also excluded, and the mean responses of neurons within bins with bin widths of 60 µm are taken for fitting to the distance curve, while the mean responses of neurons within bins with bin widths of 25° are taken for fitting to the tuning preference curve. Since there are more data points for the distance curve and the y-values of the distance curve are an order of magnitude larger than the y-values of the tuning preference curve, to ensure both curves are equally well-fitted, we compute a weighted root-mean-square loss where the data points on each curve are weighted inversely proportional to the number of data points as well as the variance of the corresponding curve.

8 constraints are imposed on the 12 fitted model parameters to ensure that the model is biologically realistic. Specifically we constraint: 1) the signs of $w_{\alpha\beta}$ ($w_{\alpha\mathrm{E}} > 0$, $w_{\alpha\mathrm{I}} < 0$), 2) the magnitudes of $w_{\alpha\beta}$ to prevent unrealistically strong connections ($|w_{\alpha\beta}| < 10$), 3) the magnitudes of $\kappa_{\alpha\beta}$ to ensure compliance with Dale's law ($|\kappa| < 0.5$) 4) $\sigma_{\mathrm{IE}}$ and $\sigma_{\mathrm{EI}}$ to be within 2 standard deviations of the estimated values of $\sigma_{\mathrm{E}}$ and $\sigma_{\mathrm{I}}$ respectively as obtained from the Methods subsection Fitting of the spatial connectivity kernel, 4) $\sigma_{\mathrm{EE}}$ and $\sigma_{\mathrm{II}}$ to be between 75 µm and 175 µm, 5) $\min\{\sigma_{\mathrm{EE}}, \sigma_{\mathrm{II}}\} > \max\{\sigma_{\mathrm{EI}}, \sigma_{\mathrm{IE}}\}$, based on connection probability data (6), 6) $\kappa_{\mathrm{EE}}$ to be within 2 standard deviations of the estimate value from the Methods subsection Estimation of $\kappa_{\mathrm{EE}}$ from data, 7) the network being an ISN ($w_{\mathrm{EE}} > 1$), and 8) the stability of the network dynamics (see SI section 7 on how the stability condition is approximately computed). Constrained optimization is performed using the `optimize.minimize` function in the `scipy` library (1) with the SLSQP (Sequential Least SQuares Programming) method, with the gradient vector being computed with PyTorch's automatic differentiation engine (7).

The validation loss is computed as the weighted root-mean-square error (using the same weights as previously described) between the data and the average single-cell perturbation response obtained with 50 numerical simulations (5 random single-cell perturbations in 10 random instantiations of the model; a random "instantiation" of the model refers to a resampling of the cell types, spatial locations, and feature tuning preferences of neurons). This validation loss is further normalized such that a value of 1 is achieved by a model predicting zero perturbation response for every neuron. A random instantiation of the model is defined as a random assignment of the cell type and spatial location of each neuron, with connectivity strength from neuron $j$ to neuron $i$ defined by

$$W_{ij}^{\mathrm{dis}} = \begin{cases} W_{\alpha_i\alpha_j}(1,1,\boldsymbol{x}_i - \boldsymbol{x}_j, \theta_i - \theta_j)\Delta V_{\alpha_j}, & \boldsymbol{x}_i \neq \boldsymbol{x}_j \\ 0, & \text{otherwise} \end{cases} \qquad \text{[S3]}$$

where $W_{\alpha\beta}$ is the connectivity function Eq. 8 with $2\pi$ replaced by $\pi$ and $\cos(\theta-\phi)$ replaced by $\cos(2(\theta-\phi))$. Each numerical simulation is performed using the same procedure as described in the Methods subsection Comparison of theory and simulations. If the network dynamics fail to converge for any one of the 50 simulations, the fitted parameters are discarded. This may occur despite the stability constraint imposed during optimization since the randomness of each neuron's spatial location causes variance in the spectral abscissa of the Jacobian that cannot be accounted for by our analysis of the continuum model.

To generate a reasonable distribution of fitted model parameters, instead of fitting the parameters directly to the mean perturbation response curve, we fit the parameters to a randomly sampled curve defined by the collection of points $\{(x_i, y_i)\}_i$, where $y_i$ is an independent sample from the Gaussian distribution $\mathcal{N}(\mu_i, \sigma_i)$ and $\mu_i, \sigma_i$ are the mean and standard error of the perturbation response at distance $x_i$ respectively. Due to the large bin widths used in the data analysis procedure by (5), nearby data points on the perturbation response curves are correlated. This is addressed by simply only fitting the model to data points which are separated roughly 60 µm apart for the distance curve and 25° apart for the tuning preference curve.

**H. Sparsely connected model.** The probability that two neurons in our sparse models are connected is given by the following equation:

$$p_{\alpha\beta}(\mu, \nu, \boldsymbol{x}-\boldsymbol{y}, \theta-\phi) = \frac{p^{\max}_{\alpha\beta}}{\varsigma^2_{\alpha\beta}} G_0(s; \varsigma_{\alpha\beta}{}^{-2}) \left(1 + 2\kappa^{\text{prob}}_{\alpha\beta} \cos(2(\theta-\phi))\right) \qquad \text{[S4]}$$

where $s = \|\boldsymbol{x}-\boldsymbol{y}\|$, $G_0$ is the Green's function kernel (Eq. 2) with $d = 0$, and $p^{\max}_{\alpha\beta}, \varsigma_{\alpha\beta}, \kappa^{\text{prob}}_{\alpha\beta}$ are given by Table S4. Note that since $\varsigma^{-2}_{\alpha\beta} G_0(0; \varsigma^{-2}_{\alpha\beta}) = 1$, $p^{\max}_{\alpha\beta}$ represents the connection probability from a neuron with cell type $\beta$ to a neuron with cell type $\alpha$ at the same spatial location. The values of $p^{\max}_{\alpha\beta}$ are based on mouse L2/3 data reported in Figure S1A of (6), where data for individual inhibitory subtypes are combined using a 0.289 : 0.210 : 0.501 ratio of PV: SST: VIP neurons, which in turn is based on the number of mouse L2/3 neurons for each inhibitory subtypes in the biophysical model by (8). The values of $\varsigma_{\text{EE}}, \varsigma_{\text{EI}}$ are obtained by fitting the curve $G_0(s; \varsigma^{-2}_{\alpha\beta})$ (with an arbitrary scaling factor) to the spatial connection probability data from (4) (Figure S7A). During fitting, each data point is weighted inversely proportional to the size of its 95% confidence interval computed by bootstrapping ($n = 17$ postsynaptic neurons). Furthermore, since we believe the connected probability is underestimated for pairs of neurons separated more than 300 µm apart due to a limited field-of-view (reported to be between 500 µm-850 µm), the curve is only fitted to data points within 300 µm. Finally, we assume $\varsigma_{\text{IE}} = \varsigma_{\text{EI}}$ and $\varsigma_{\text{II}} = \varsigma_{\text{EE}}$ based on data from (6). The value of $\kappa^{\text{prob}}_{EE}$ is obtained with the procedure in Methods subsection Estimation of $\kappa_{\text{EE}}$ from data. $\kappa^{\text{prob}}_{\alpha\beta}$ is set to 0 for other cell type combinations following existing models (8) and experimental data (4, 9–12).

Connection strength between a pair of connected neurons $i, j$ in the these sparse model is conceptually defined by $\frac{W^{\text{dis}}_{ij}}{p_{ij}}$, where $p_{ij} := p_{\alpha_i\alpha_j}(\mu_i, \mu_j, \boldsymbol{x}_i - \boldsymbol{x}_j, \theta_i - \theta_j)$ and $\boldsymbol{W}^{\text{dis}}$ is given by Eq. 13. However, this naive definition has a problem: since $K_\eta(z) \sim \sqrt{\frac{\pi}{2z}} e^{-z}$ as $|z| \to \infty$ (13, Equation 9.7.2), $G_d(s; \lambda) \sim s^{-\eta} e^{-\sqrt{\lambda} s}$ as $s \to \infty$. But the spatial component of $W_{\alpha\beta}$ is given by $G_d(s; \sigma^{-2}_{\alpha\beta})$, while the spatial component of $p_{\alpha\beta}$ is given by $G_0(s; \varsigma^{-2}_{\alpha\beta})$, so the spatial component of the connectivity strength, which is given by the ratio of these two functions, scales as $s^{-\eta-1} e^{-s(\sigma^{-1}_{\alpha\beta} - \varsigma^{-1}_{\alpha\beta})}$. Thus, the connectivity strength diverges as $s \to \infty$ whenever $\sigma_{\alpha\beta} > \varsigma_{\alpha\beta}$. This is clearly not biologically realistic. To address this issue, we define a regularized version of this ratio by

$$\tilde{R}_d(s; \sigma, \varsigma) := \begin{cases} \frac{G_d(s;\sigma^{-2})}{G_0(s;\varsigma^{-2})} & \sigma \le \varsigma \text{ or } s < s_{\min}(\sigma, \varsigma) \\ \frac{G_d(s_{\min}(\sigma,\varsigma);\sigma^{-2})}{G_0(s_{\min}(\sigma,\varsigma);\varsigma^{-2})} & \text{otherwise,} \end{cases} \qquad \text{[S5]}$$

$$s_{\min}(\sigma, \varsigma) := \min_{s>0} \frac{G_d(s; \sigma^{-2})}{G_0(s; \varsigma^{-2})}. \qquad \text{[S6]}$$

We note that the minimum $s_{\min}(\sigma, \varsigma)$ exists and is nonzero when $\sigma > \varsigma$ since it can be shown that the derivative of the ratio $\frac{G_d(s;\sigma^{-2})}{G_0(s;\varsigma^{-2})}$ with respect to $s$ is negative as $s \to 0$. But also this derivative is positive as $s \to \infty$ since the ratio diverges when $\sigma > \varsigma$, so that by the Intermediate Value Theorem $s_{\min}(\sigma, \varsigma)$ must be finite and nonzero. Setting connectivity strength based on the regularized ratio $\tilde{R}_d(s; \sigma, \varsigma)$ fixes the divergence, but it also causes the overall connectivity strength to be reduced. To compensate for this reduction, we simply scale up the connectivity strength by defining

$$R_{d,p}(s; \sigma, \varsigma) := \tilde{R}_d(s; \sigma, \varsigma) \frac{\|G_d(\|\cdot\|; \sigma^{-2})\|_p}{\|G_0(\|\cdot\|; \varsigma^{-2}) \tilde{R}_d(\|\cdot\|; \sigma, \varsigma)\|_p} \qquad \text{[S7]}$$

where $\|\cdot\|_p$ represents the $p$-norm of a function. This ensures that the spatial component of the product of connection probability and connection strength, $G_0(s; \varsigma^{-2}) R_{d,p}(s; \sigma, \varsigma)$, has the same norm as the connectivity of the fully-connected model, $G_d(s; \sigma^{-2})$. Empirically, we find that setting $p = 1.5$ yields a good match between the perturbation response of the sparse model and that of the fully-connected model. In summary, the connectivity strength between a pair of connected neurons $i, j$ in the sparse model is given by

$$\frac{\Delta V_{\alpha_j} w_{\alpha_i\alpha_j} \varsigma^2_{\alpha_i\alpha_j}}{2\pi p^{\max}_{\alpha_i\alpha_j} \sigma^2_{\alpha_i\alpha_j}} R_{d,p}(s_{ij}; \sigma_{\alpha_i\alpha_j}, \varsigma_{\alpha_i\alpha_j}) \frac{1 + 2\kappa_{\alpha_i\alpha_j} \cos(2(\theta_i - \theta_j))}{1 + 2\kappa^{\text{prob}}_{\alpha_i\alpha_j} \cos(2(\theta_i - \theta_j))} \tag{S8}$$

if $s_{ij} \neq 0$, and 0 otherwise, where $s_{ij} = \|\boldsymbol{x}_i - \boldsymbol{x}_j\|$. This is illustrated by Figure S7C, which plots the connection strength of $\text{E} \to \text{E}$ and $\text{I} \to \text{E}$ connections as a function of distance in our sparse models.

Finally, we note that we used the function $G_0(s; \varsigma^{-2})$ rather than a Gaussian $\exp(-\frac{s^2}{2\varsigma^2})$ to model connection probability because the Gaussian decays much more quickly than $G_0(s; \varsigma^{-2})$ ($\mathcal{O}(e^{-\frac{1}{2}(\varsigma^{-1}s)^2})$ compared to $\mathcal{O}(se^{-\varsigma^{-1}s})$). Thus, the issue of connection strength divergence becomes much more severe with a Gaussian, and our attempts to treat the divergence via the same approach results in a much worse match between the perturbation response of the sparse model and that of the fully-connected model.

**Table S4. Connection probability parameters**

| Parameter | Value | Parameter | Value |
|---|---|---|---|
| $p^{\max}_{\text{EE}}$ | 0.11 | $\varsigma_{\text{EE}}, \varsigma_{\text{II}}$ | 110.6 µm |
| $p^{\max}_{\text{EI}}$ | 0.331 | $\varsigma_{\text{EI}}, \varsigma_{\text{IE}}$ | 73.1 µm |
| $p^{\max}_{\text{IE}}$ | 0.578 | $\kappa^{\text{prob}}_{\text{EE}}$ | 0.198 |
| $p^{\max}_{\text{II}}$ | 0.284 | $\kappa^{\text{prob}}_{\text{EI}}, \kappa^{\text{prob}}_{\text{IE}}, \kappa^{\text{prob}}_{\text{II}}$ | 0 |

**I. Models with inhomogeneous gain.** To model the inhomogeneity in neuronal gain that is caused by drifting gratings, we assume that neuronal gain can be parametrized by

$$g(\mathbf{x}, \theta; \psi) = g_0 + (1 - g_0) \exp\left(-\frac{\|x\|^2}{2\sigma_g^2}\right)(1 + 2\kappa_g \cos(\theta - \psi)) \tag{S9}$$

where $\psi$ is the direction of the drifting grating, and $g_0, \sigma_g, \kappa_g$ are scalar parameters. The inhomogeneous gain is then incorporated into the 50 best-fit models in Figure 6 by setting the connection strength between post-synaptic neuron $(\alpha, \mu, \boldsymbol{x}, \theta)$ and pre-synaptic neuron $(\beta, \nu, \boldsymbol{y}, \phi)$ as $g(\mathbf{x}, \theta; \psi) W_{\alpha\beta}(\mu, \nu, \boldsymbol{x}, \boldsymbol{y}, \theta, \phi)$, and we numerically simulate the single-cell perturbation responses of these models. Since Ref. (5) measured the single-cell perturbation response averaged over all different directions $\psi$ of drifting grating, we need to compute the perturbation response averaged over different values of $\psi$. However, we note that the same average can equivalently be computed by fixing $\psi = 0$ and averaging over perturbations of neurons with different feature tuning preferences $\phi$. We took this approach since it is computationally more efficient. Specifically, for each of the top 50 fitted models, we set $\psi = 0$, simulated a total of 200 random single-cell perturbations (20 random single-cell perturbations in 10 random instantiations of the model; see SI Section 1G for the meaning of a random "instantiation"), and computed the average perturbation response. Since we simulated network response with a uniform gain of 0.5 in Figures 6G-H in the main text, we set $g_0 = 0.5$ and $\kappa_g = 0.5$ so that the minimum neuronal gain is also 0.5. We set $\sigma_g = 968\,\mu\text{m}$ because Ref. (5) used drifting gratings with a Gaussian window of 44°, and the cortical magnification factor (CMF) in mouse V1 has been measured to be roughly between 0.0005 and $0.002\,\text{mm}^2/\text{deg}^2$. Taking the square root of $0.0005\,\text{mm}^2/\text{deg}^2$ yields a magnification factor of roughly 22 µm/deg, and thus we get $\sigma_g = 22\,\mu\text{m/deg} \times 44° = 968\,\mu\text{m}$. 2 out of the 50 best-fit models are no longer dynamically stable when $\kappa_g = 0.5$, so we simply exclude the perturbation responses of these two models.

#### 2. The spatial kernel $G_d(s; \lambda)$

In the main text, we stated that $G_d(s; \lambda)$, defined by

$$G_d(s; \lambda) = \frac{1}{(2\pi)^{\frac{d}{2}}} \left(\frac{\sqrt{\lambda}}{s}\right)^{\eta} K_\eta(\sqrt{\lambda} s), \qquad \eta = \frac{d}{2} - 1 \tag{S10}$$

is the unique Green's function of the operator $\lambda - \nabla^2$, a fact which is used to derive the linear response equation for a simplified model (Eq. 7). Owing to the fundamental nature of the Laplacian in mathematics and physics, this is a fairly standard result (14, Section 7, Corollary 2). Nonetheless, we provide a simple, informal proof here for completeness. To start with, we provide a proof sketch of a well-known result about the Fourier transform of spherically symmetric functions, also known as the Hankel transform:

**Lemma 2.1.** *For all $d \in \mathbb{N}$, $\boldsymbol{k} \in \mathbb{R}^d$, and L1-integrable functions $f : (0, \infty) \to \mathbb{C}$, we have*

$$\int_{\mathbb{R}^d} e^{i\boldsymbol{k}\cdot\boldsymbol{x}} f(\|\boldsymbol{x}\|)\, d\boldsymbol{x} = (2\pi)^{\frac{d}{2}} k^{-\eta} \int_0^\infty s^{\eta+1} f(s) J_\eta(ks)\, ds$$

*where $k = \|\boldsymbol{k}\|$, $\eta = \frac{d}{2} - 1$ and $J_\eta$ is the Bessel function of the first kind of order $\eta$.*

*Proof.* One can check the $d = 1$ case easily by using the fact that $J_{-\frac{1}{2}}(z) = \sqrt{\frac{2}{\pi}} \frac{\cos(z)}{\sqrt{z}}$ (13), so we will focus on the case $d \geq 2$. Pick a polar coordinates system for our integration variable $\boldsymbol{x} = (s, \theta, \phi_1, \cdots, \phi_{d-2})$, such that $\boldsymbol{k} \cdot \boldsymbol{x} = ks\cos\theta$. Integrating over $\phi_1, \cdots, \phi_{d-2}$ yields a factor of $\Omega_{d-1}$, the solid angle subtended by the $(d-2)$-sphere (or in other words, the surface area of a unit $(d-2)$-sphere). $\Omega_d$ is well-known to be

$$\Omega_d = \frac{2\pi^{\frac{d}{2}}}{\Gamma(\frac{d}{2})}$$

where $\Gamma$ is the gamma function. Thus we have

$$\begin{aligned}\int_{\mathbb{R}^d} e^{i\boldsymbol{k}\cdot\boldsymbol{x}} f(\|\boldsymbol{x}\|)\, d\boldsymbol{x} &= \Omega_{d-1} \int_0^\infty \int_0^\pi e^{iks\cos\theta} f(s) s^{d-1} \sin^{d-2}\theta\, d\theta ds \\ &= \Omega_{d-1} \int_0^\infty \left( \int_0^\pi \cos(ks\cos\theta) \sin^{2\eta}\theta d\theta \right) f(s) s^{d-1}\, ds.\end{aligned}$$

Note that in the last step, we used the fact that the imaginary part of $e^{iks\cos\theta}$ vanishes under the integral since by applying the transformation $\theta \to \theta + \frac{\pi}{2}$, we have

$$\int_0^\pi \sin(ks\cos\theta) \sin^{2\eta}\theta\, d\theta = -\int_{-\frac{\pi}{2}}^{\frac{\pi}{2}} \sin(ks\sin\theta) \cos^{2\eta}\theta\, d\theta$$

and we see that the transformed integrand is an odd function and thus integrates to 0. But now the integral over $\theta$ can be written in terms of the Bessel function of the first kind due to the integral representation (13, Equation 9.1.20)

$$J_\eta(z) = \frac{(\frac{z}{2})^\eta}{\sqrt{\pi}\Gamma(\eta + \frac{1}{2})} \int_0^\pi \cos(z\cos\theta) \sin^{2\eta}\theta\, d\theta$$

which is valid for $\mathrm{Re}(\eta) > -\frac{1}{2}$. Thus

$$\begin{aligned}\int_{\mathbb{R}^d} e^{i\boldsymbol{k}\cdot\boldsymbol{x}} f(\boldsymbol{x})\, d\boldsymbol{x} &= \frac{2\pi^{\eta+\frac{1}{2}}}{\Gamma(\eta+\frac{1}{2})} \int_0^\infty \frac{\sqrt{\pi}\Gamma(\eta+\frac{1}{2})}{(\frac{ks}{2})^\eta} J_\eta(ks) f(s) s^{2\eta+1}\, ds \\ &= (2\pi)^{\frac{d}{2}} k^{-\eta} \int_0^\infty s^{\eta+1} f(s) J_\eta(ks)\, ds.\end{aligned}$$

□

**Proposition 2.2.** *Let $\nabla^2$ be the Laplacian operator on $\mathbb{R}^d$. For all $\lambda \in \mathbb{C} \setminus (-\infty, 0]$, the inverse $(\lambda - \nabla^2)^{-1}$ exists and*

$$(\lambda - \nabla^2)^{-1} f(\boldsymbol{x}) = \int_{\mathbb{R}^d} G_d(\|\boldsymbol{x} - \boldsymbol{y}\|; \lambda) f(\boldsymbol{y})\, d\boldsymbol{y}.$$

*In other words, $G_d(\|\boldsymbol{x} - \boldsymbol{y}\|; \lambda)$ is the unique Green's function of $\lambda - \nabla^2$.*

*Proof.* We want to show that for all $f$,

$$(\lambda - \nabla^2) \int_{\mathbb{R}^d} G_d(\|\boldsymbol{x} - \boldsymbol{y}\|; \lambda) f(\boldsymbol{y})\, d\boldsymbol{y} = f(\boldsymbol{x}).$$

Let $\hat{G}_d(\boldsymbol{k}; \lambda)$, $\hat{f}(\boldsymbol{k})$ denote the Fourier transform of $G_d(\|\boldsymbol{x}\|; \lambda)$, $f(\boldsymbol{x})$ respectively, and let $k = \|\boldsymbol{k}\|$. Then since $\nabla \mapsto i\boldsymbol{k}$ under the Fourier transform, we have

$$(\lambda + k^2) \hat{G}_d(\boldsymbol{k}; \lambda) \hat{f}(\boldsymbol{k}) = \hat{f}(\boldsymbol{k})$$

where we applied the convolution theorem to the integral. Thus we just need to show that $\hat{G}_d(\boldsymbol{k};\lambda) = (\lambda + k^2)^{-1}$, which is well defined for all $\boldsymbol{k} \in \mathbb{R}^d$ due to the condition $\lambda \notin (-\infty, 0]$. By lemma 2.1, we have

$$\int_{\mathbb{R}^d} e^{i\boldsymbol{k}\cdot\boldsymbol{x}} G_d(\|\boldsymbol{x}\|;\lambda)\,d\boldsymbol{x} = k^{-\eta}\int_0^\infty s^{\eta+1}\left(\frac{\sqrt{\lambda}}{s}\right)^\eta K_\eta\left(\sqrt{\lambda}s\right)J_\eta(ks)\,ds$$

$$= \left(\frac{\sqrt{\lambda}}{k}\right)^\eta \int_0^\infty K_\eta\left(\sqrt{\lambda}s\right)J_\eta(ks)s\,ds.$$

The last integral is known to be given by (15, Section 6.521, Equation 2)

$$\int_0^\infty K_\eta(as)J_\eta(bs)s\,ds = \left(\frac{b}{a}\right)^\eta \frac{1}{a^2+b^2}.$$

which is valid for $\mathrm{Re}(a) > 0$, $b > 0$, and $\mathrm{Re}(\eta) > -1$. Thus we immediately have

$$\hat{G}_d(\boldsymbol{k};\lambda) = \int_{\mathbb{R}^d} e^{i\boldsymbol{k}\cdot\boldsymbol{x}} G_d(\|\boldsymbol{x}\|;\lambda)\,d\boldsymbol{x} = \frac{1}{\lambda + k^2} \qquad \text{[S11]}$$

so we are done. □

#### 3. Derivation of perturbation response for the simplified model

In the section Derivation for a simplified model of the main text, we outlined the steps for deriving the linear response in a simplified model whose connectivity depends only on cell type and space, and whose connectivity width depends only on presynaptic cell type. Here we fill in some intermediate steps of the derivation. We start with the definition of the connectivity kernel (Eq. 5 in the main text)

$$W_{\alpha\beta}(\boldsymbol{x}-\boldsymbol{y}) = \frac{w_{\alpha\beta}}{\sigma_\beta^2} G_d(s;\sigma_\beta^{-2}). \qquad \text{[S12]}$$

where $s = \|\boldsymbol{x}-\boldsymbol{y}\|$. Our first step is to define the connectivity operator $W$ associated with this kernel. This operator should act on the firing rates $r_\beta(\boldsymbol{y})$ such that the recurrent input to neuron $(\alpha, \boldsymbol{x})$ is given by

$$\sum_{\beta=0}^{N_c-1}\int_{\mathbb{R}^d} W_{\alpha\beta}(\boldsymbol{x}-\boldsymbol{y})r_\beta(\boldsymbol{y})\,d\boldsymbol{y}. \qquad \text{[S13]}$$

Because $G_d(s;\sigma^{-2})$ is the Green's function of $\sigma^{-2} - \nabla^2$, convolution of any function by $G_d(s;\sigma^{-2})$ is equivalent to applying the inverse operator $(\sigma^{-2} - \nabla^2)^{-1}$ to that function (Proposition 2.2). Thus Eq. S13 can be written as

$$\sum_{\beta=0}^{N_c-1}\int_{\mathbb{R}^d} W_{\alpha\beta}(\boldsymbol{x}-\boldsymbol{y})r_\beta(\boldsymbol{y})\,d\boldsymbol{y}$$

$$= \sum_{\beta=0}^{N_c-1}\frac{w_{\alpha\beta}}{\sigma_\beta^2}\int_{\mathbb{R}^d} G_d(\|\boldsymbol{x}-\boldsymbol{y}\|;\sigma_\beta^{-2})r_\beta(\boldsymbol{y})\,d\boldsymbol{y}$$

$$= \sum_{\beta=0}^{N_c-1}\frac{w_{\alpha\beta}}{\sigma_\beta^2}(\sigma_\beta^{-2} - \nabla^2)^{-1}r_\beta(\boldsymbol{x}).$$

Define $\boldsymbol{W}$ as the $N_c \times N_c$ matrix of elements $w_{\alpha\beta}$ and $\boldsymbol{\Sigma}$ as the $N_c \times N_c$ diagonal matrix with elements $\sigma_\beta^2$. Furthermore, let $\boldsymbol{\Sigma}^{-1} - \nabla^2\boldsymbol{I}$ denote an $N_c \times N_c$ diagonal matrix of spatial operators with elements $\sigma_\beta^{-2} - \nabla^2$. Because $\boldsymbol{\Sigma}^{-1} - \nabla^2\boldsymbol{I}$ is a diagonal matrix, its inverse, $(\boldsymbol{\Sigma}^{-1} - \nabla^2\boldsymbol{I})^{-1}$ is also a diagonal matrix but with elements $(\sigma_\beta^{-2} - \nabla^2)^{-1}$. That is, $(\boldsymbol{\Sigma}^{-1} - \nabla^2\boldsymbol{I})^{-1}$ acts on the firing rates by

$$\begin{bmatrix} (\sigma_0^{-2} - \nabla^2)^{-1} & 0 & \cdots & 0 \\ 0 & (\sigma_1^{-2} - \nabla^2)^{-1} & \cdots & 0 \\ 0 & \vdots & \ddots & 0 \\ 0 & 0 & \cdots & (\sigma_{N_c-1}^{-2} - \nabla^2)^{-1} \end{bmatrix} \begin{bmatrix} r_0(\boldsymbol{x}) \\ r_1(\boldsymbol{x}) \\ \vdots \\ r_{N_c-1}(\boldsymbol{x}) \end{bmatrix} = \begin{bmatrix} (\sigma_0^{-2} - \nabla^2)^{-1}r_0(\boldsymbol{x}) \\ (\sigma_1^{-2} - \nabla^2)^{-1}r_1(\boldsymbol{x}) \\ \vdots \\ (\sigma_{N_c-1}^{-2} - \nabla^2)^{-1}r_{N_c-1}(\boldsymbol{x}) \end{bmatrix} \qquad \text{[S14]}$$

Thus, we may write the connectivity operator $W$ compactly as

$$W = \boldsymbol{W}\boldsymbol{\Sigma}^{-1}(\boldsymbol{\Sigma}^{-1} - \nabla^2 \boldsymbol{I})^{-1}. \tag{S15}$$

We note that this notation is non-standard and informal. In the next section where we derive the perturbation response in the full model, we shall use a more formal definition of the connectivity operator using tensor products and the bra-ket notation.

As we noted in the main text, we need to compute the linear response operator $L = (I - W)^{-1}$. In the main text we computed $L$ by appealing to the Woodbury matrix (operator) identity since it reduces the number of intermediate steps required and since the derivation of the full model also uses the Woodbury matrix identity. But in fact the operator $L$ can be derived using only basic algebraic manipulations, which we show here for completeness. Indeed,

$$\begin{aligned}
L = (I - W)^{-1} &= \left(I - \boldsymbol{W}\boldsymbol{\Sigma}^{-1}(\boldsymbol{\Sigma}^{-1} - \nabla^2 \boldsymbol{I})^{-1}\right)^{-1} \\
&= \left((\boldsymbol{\Sigma}^{-1} - \nabla^2 \boldsymbol{I})(\boldsymbol{\Sigma}^{-1} - \nabla^2 \boldsymbol{I})^{-1} - \boldsymbol{W}\boldsymbol{\Sigma}^{-1}(\boldsymbol{\Sigma}^{-1} - \nabla^2 \boldsymbol{I})^{-1}\right)^{-1} \\
&= (\boldsymbol{\Sigma}^{-1} - \nabla^2 \boldsymbol{I})(\boldsymbol{\Sigma}^{-1} - \nabla^2 \boldsymbol{I} - \boldsymbol{W}\boldsymbol{\Sigma}^{-1})^{-1} \\
&= (\boldsymbol{\Sigma}^{-1} - \nabla^2 \boldsymbol{I} - \boldsymbol{W}\boldsymbol{\Sigma}^{-1} + \boldsymbol{W}\boldsymbol{\Sigma}^{-1})(\boldsymbol{\Sigma}^{-1} - \nabla^2 \boldsymbol{I} - \boldsymbol{W}\boldsymbol{\Sigma}^{-1})^{-1} \\
&= I + \boldsymbol{W}\boldsymbol{\Sigma}^{-1}(\boldsymbol{\Sigma}^{-1} - \nabla^2 \boldsymbol{I} - \boldsymbol{W}\boldsymbol{\Sigma}^{-1})^{-1} \\
&= I + \boldsymbol{W}\boldsymbol{\Sigma}^{-1}((\boldsymbol{I} - \boldsymbol{W})\boldsymbol{\Sigma}^{-1} - \nabla^2 \boldsymbol{I})^{-1}
\end{aligned}$$

As in the main text, we now diagonalize the matrix $(\boldsymbol{I} - \boldsymbol{W})\boldsymbol{\Sigma}^{-1}$ as $\boldsymbol{Q}\boldsymbol{\Lambda}\boldsymbol{Q}^{-1}$. Then

$$\begin{aligned}
L = (I - W)^{-1} &= I + \boldsymbol{W}\boldsymbol{\Sigma}^{-1}(\boldsymbol{Q}\boldsymbol{\Lambda}\boldsymbol{Q}^{-1} - \nabla^2 \boldsymbol{I})^{-1} \\
&= I + \boldsymbol{W}\boldsymbol{\Sigma}^{-1}\boldsymbol{Q}(\boldsymbol{\Lambda} - \nabla^2 \boldsymbol{I})^{-1}\boldsymbol{Q}^{-1}
\end{aligned}$$

where in the last line we note that $\nabla^2$ commutes with $\boldsymbol{Q}$ because $\nabla^2$ is a scalar operator rather than a matrix operator.

Similar to before, $\boldsymbol{\Lambda} - \nabla^2 \boldsymbol{I}$ represents a diagonal matrix of spatial operators $\lambda_\gamma - \nabla^2$, and thus its inverse $(\boldsymbol{\Lambda} - \nabla^2 \boldsymbol{I})^{-1}$ is also a diagonal matrix of inverse operators $(\lambda_\gamma - \nabla^2)^{-1}$. Thus, if we apply the linear response operator $L$ to an arbitrary perturbation $|h\rangle$, we obtain the perturbation response as

$$\begin{aligned}
r_\alpha(\boldsymbol{x}) &= h_\alpha(\boldsymbol{x}) + \sum_{\beta',\gamma=0}^{N_c-1} [\boldsymbol{W}\boldsymbol{\Sigma}^{-1}\boldsymbol{Q}]_{\alpha\gamma}(\lambda_\gamma - \nabla^2)^{-1}[\boldsymbol{Q}^{-1}]_{\gamma\beta'} h_{\beta'}(\boldsymbol{x}) \\
&= h_\alpha(\boldsymbol{x}) + \sum_{\beta',\gamma=0}^{N_c-1} [\boldsymbol{W}\boldsymbol{\Sigma}^{-1}\boldsymbol{Q}]_{\alpha\gamma}[\boldsymbol{Q}^{-1}]_{\gamma\beta'}(\lambda_\gamma - \nabla^2)^{-1} h_{\beta'}(\boldsymbol{x}).
\end{aligned}$$

But again, since $G_d(s; \lambda_\gamma)$ is the Green's function of $\lambda_\gamma - \nabla^2$, the application of the inverse operator $(\lambda_\gamma - \nabla^2)^{-1}$ to $h_{\beta'}(\boldsymbol{x})$ is given by the convolution of $G_d(s; \lambda_\gamma)$ with $h_{\beta'}(\boldsymbol{x})$. Thus we have

$$r_\alpha(\boldsymbol{x}) = h_\alpha(\boldsymbol{x}) + \sum_{\beta',\gamma=0}^{N_c-1} [\boldsymbol{W}\boldsymbol{\Sigma}^{-1}\boldsymbol{Q}]_{\alpha\gamma}[\boldsymbol{Q}^{-1}]_{\gamma\beta'} \int_{\mathbb{R}^d} G_d(\|\boldsymbol{x} - \boldsymbol{z}\|; \lambda_\gamma) h_{\beta'}(\boldsymbol{z})\, d\boldsymbol{z}. \tag{S16}$$

In particular, for a unit-amplitude single-cell perturbation of neuron $(\beta, \boldsymbol{y})$ (represented by the delta function $h_{\beta'}(\boldsymbol{z}) = \delta_{\beta'\beta}\delta(\boldsymbol{z} - \boldsymbol{y})$), the perturbation response of an unperturbed neuron $(\alpha, \boldsymbol{x})$, denoted by $\tilde{L}_{\alpha\beta}(\boldsymbol{x} - \boldsymbol{y})$, is thus determined by

$$\tilde{L}_{\alpha\beta}(\boldsymbol{x} - \boldsymbol{y}) = \sum_{\gamma=0}^{N_c-1} [\boldsymbol{W}\boldsymbol{\Sigma}^{-1}\boldsymbol{Q}]_{\alpha\gamma}[\boldsymbol{Q}^{-1}]_{\gamma\beta} G_d(\|\boldsymbol{x} - \boldsymbol{y}\|; \lambda_\gamma). \tag{S17}$$

#### 4. Derivation of perturbation response for the full model

In this section we show how to extend our derivation for the simplified model to the general model which includes feature tuning preference and selectivity, and allows for arbitrary connectivity widths. Unlike the pervious section, we will use the bra-ket notation freely and extensively throughout this section. For an explanation of the bra-ket notation, see Materials and Methods in the main text.

Consider our connectivity function for the full model,

$$W_{\alpha\beta}(\mu,\nu,\boldsymbol{x}-\boldsymbol{y},\theta-\phi)=\frac{w_{\alpha\beta}}{2\pi\sigma_{\alpha\beta}^2}G_d(s;\sigma_{\alpha\beta}^{-2})(1+2\kappa_{\alpha\beta}f_\alpha(\mu)g_\beta(\nu)\cos(\theta-\phi)). \quad \text{[S18]}$$

Since our steady state equation is given by

$$r_\alpha(\mu,\boldsymbol{x},\theta)=\sum_{\beta=0}^{N_c-1}\int_0^1\int_{\mathbb{R}^d}\int_{-\pi}^{\pi}W_{\alpha\beta}(\mu,\nu,\boldsymbol{x}-\boldsymbol{y},\theta-\phi)r_\beta(\nu,\boldsymbol{y},\phi)P_\beta(\nu)\,d\phi d\boldsymbol{y}d\nu+h_\alpha(\mu,\boldsymbol{x},\theta), \quad \text{[S19]}$$

we see that if we define the integral operator $W$ by

$$\langle\alpha,\mu,\boldsymbol{x},\theta|W|\beta,\nu,\boldsymbol{y},\phi\rangle=W_{\alpha\beta}(\mu,\nu,\boldsymbol{x}-\boldsymbol{y},\theta-\phi)P_\beta(\nu) \quad \text{[S20]}$$

where $|\beta,\nu,\boldsymbol{y},\phi\rangle$ represents a delta function centered at neuron $(\beta,\nu,\boldsymbol{y},\phi)$, and let $|r\rangle,|h\rangle$ represent the firing rate and external input functions (as vectors in an infinite-dimensional vector space, such that $\langle\alpha,\mu,\boldsymbol{x},\theta\,|\,r\rangle=r_\alpha(\mu,\boldsymbol{x},\theta)$ and $\langle\alpha,\mu,\boldsymbol{x},\theta\,|\,h\rangle=h_\alpha(\mu,\boldsymbol{x},\theta)$), then the steady state equation reduces to the abstract form

$$|r\rangle=W|r\rangle+|h\rangle. \quad \text{[S21]}$$

As explained in the main text, since $|r\rangle$ can be solved abstractly as $(I-W)^{-1}|h\rangle$, our goal is to compute the linear response operator $L=(I-W)^{-1}$.

The first observation is that the linear response can be computed for each Fourier mode of $\theta$ independently. Specifically, let $|n\rangle$ denote the $n$-th Fourier mode on the circle, i.e. $\langle\theta\,|\,n\rangle=\frac{1}{\sqrt{2\pi}}e^{in\theta}$. Then because the connectivity function depends on the pre- and post-synaptic feature tuning preference $\theta$ and $\phi$ via the cosine term $\cos(\theta-\phi)$, we can decompose $W$ as

$$W=\sum_{n=-1}^{1}W_n\otimes|n\rangle\langle n| \quad \text{[S22]}$$

where each $W_n$ is a linear operator on the tensor product space $X:=\mathbb{R}^{N_c}\otimes L^2([0,1])\otimes L^2(\mathbb{R}^d)$, i.e. it acts on cell type, feature selectivity, and space, but not feature preference. Note that we must have $W_{-1}=W_1$ in order for the connectivity function to be real-valued and symmetric. To see this, we simply note that

$$\begin{aligned}\langle\alpha,\mu,\boldsymbol{x},\theta|W|\beta,\nu,\boldsymbol{y},\phi\rangle&=\sum_{n=-1}^{1}\langle\theta\,|\,n\rangle\langle n\,|\,\phi\rangle\langle\alpha,\mu,\boldsymbol{x}|W_n|\beta,\nu,\boldsymbol{y}\rangle\\&=\frac{1}{2\pi}\sum_{n=-1}^{1}e^{in\theta}e^{-in\phi}\langle\alpha,\mu,\boldsymbol{x}|W_n|\beta,\nu,\boldsymbol{y}\rangle\\&=\frac{1}{2\pi}(\langle\alpha,\mu,\boldsymbol{x}|W_0|\beta,\nu,\boldsymbol{y}\rangle+(e^{i(\theta-\phi)}+e^{-i(\theta-\phi)})\langle\alpha,\mu,\boldsymbol{x}|W_1|\beta,\nu,\boldsymbol{y}\rangle)\\&=\frac{1}{2\pi}(\langle\alpha,\mu,\boldsymbol{x}|W_0|\beta,\nu,\boldsymbol{y}\rangle+2\cos(\theta-\phi)\langle\alpha,\mu,\boldsymbol{x}|W_1|\beta,\nu,\boldsymbol{y}\rangle)\end{aligned}$$

Comparing the last line to Eq. S20, we see that $W_0$ and $W_{\pm1}$ must therefore be defined by

$$\langle\alpha,\mu,\boldsymbol{x}|W_0|\beta,\nu,\boldsymbol{y}\rangle=\frac{w_{\alpha\beta}}{\sigma_{\alpha\beta}^2}P_\beta(\nu)G_d(s;\sigma_{\alpha\beta}^{-2}) \quad \text{[S23]}$$

$$\langle\alpha,\mu,\boldsymbol{x}|W_{\pm1}|\beta,\nu,\boldsymbol{y}\rangle=\frac{w_{\alpha\beta}\kappa_{\alpha\beta}}{\sigma_{\alpha\beta}^2}f_\alpha(\mu)g_\beta(\nu)P_\beta(\nu)G_d(s;\sigma_{\alpha\beta}^{-2}) \quad \text{[S24]}$$

where $s=\|\boldsymbol{x}-\boldsymbol{y}\|$.

The key point is that the linear response operator $L=(I-W)^{-1}$ can be computed in terms of the simpler operators $L_n:=(I_X-W_n)^{-1}$ as

$$L=I+\sum_{n=-1}^{1}(L_n-I_X)\otimes|n\rangle\langle n| \quad \text{[S25]}$$

where $I_X$ is the identity operator on the tensor product space $X$ we defined earlier. Indeed,

$$
\begin{aligned}
(I-W)L &= (I-W)\left(I + \sum_{n=-1}^{1}(L_n - I_X)\otimes |n\rangle\langle n|\right) \\
&= I - W + \left(\sum_{n=-1}^{1}(L_n - I_X)\otimes |n\rangle\langle n|\right) - \left(\sum_{n=-1}^{1} W_n \otimes |n\rangle\langle n|\right)\left(\sum_{n=-1}^{1}(L_n - I_X)\otimes |n\rangle\langle n|\right) \\
&= I - W + \left(\sum_{n=-1}^{1}(L_n - I_X)\otimes |n\rangle\langle n|\right) - \left(\sum_{n=-1}^{1} W_n(L_n - I_X)\otimes |n\rangle\langle n|\right) \\
&= I - W + \sum_{n=-1}^{1}(I_X - W_n)(L_n - I_X)\otimes |n\rangle\langle n| \\
&= I - W + \sum_{n=-1}^{1} W_n \otimes |n\rangle\langle n| \\
&= I
\end{aligned}
$$

where we note that in going from the second to the third line, all the cross terms from the product disappear since $\langle n\,|\,m\rangle = \delta_{nm}$.

Thus, it remains to compute $L_n = (I_X - W_n)^{-1}$. The strategy is similar to that of the derivation in the section Derivation for a simplified model of the main text. We will decompose each $W_n$ into $U_n C^{-1} V_n$ such that each $L_n$ can be computed via the Woodbury matrix (operator) identity (16) as

$$L_n = (I_X - W_n)^{-1} = I_X + U_n(C - V_n U_n)^{-1} V_n$$

To unify our discussion for the case of $n = 0$ and $n = \pm 1$, first let us define the variables $A_{n\alpha\beta}$ and functions $f_{n\alpha}, g_{n\beta}$ by

$$
\begin{aligned}
A_{0\alpha\beta} &= w_{\alpha\beta}\sigma_{\alpha\beta}^{-2}, \quad & A_{1\alpha\beta} &= w_{\alpha\beta}\sigma_{\alpha\beta}^{-2}\kappa_{\alpha\beta} \\
f_{0\alpha}(\mu) &= 1, & f_{1\alpha}(\mu) &= f_\alpha(\mu) \\
g_{0\beta}(\nu) &= P_\beta(\nu), & g_{1\beta}(\nu) &= g_\beta(\nu)P_\beta(\nu)
\end{aligned}
\qquad \text{[S26]}
$$

Then Eq. S23 and Eq. S24 can be written together

$$\langle \alpha, \mu, \boldsymbol{x}|W_n|\beta, \nu, \boldsymbol{y}\rangle = A_{n\alpha\beta} f_{n\alpha}(\mu) g_{n\beta}(\nu) G_d(s; \sigma_{\alpha\beta}^{-2}). \qquad \text{[S27]}$$

Because $G_d(s;\sigma_{\alpha\beta}^{-2})$ is the Green's function of the linear operator $\sigma_{\alpha\beta}^{-2} - \nabla^2$ (see section 2), we can therefore write $W_n$ as

$$
\begin{aligned}
W_n &= \sum_{\alpha,\beta=0}^{N_c-1} A_{n\alpha\beta}|\alpha\rangle\langle\beta| \otimes |f_{n\alpha}\rangle\langle g_{n\beta}| \otimes (\sigma_{\alpha\beta}^{-2} - \nabla^2)^{-1} \\
&= \sum_{\alpha,\beta=0}^{N_c-1} A_{n\alpha\beta}|\alpha, f_{n\alpha}\rangle\langle\beta, g_{n\beta}| \otimes (\sigma_{\alpha\beta}^{-2} - \nabla^2)^{-1}.
\end{aligned}
\qquad \text{[S28]}
$$

where $|f_{n\alpha}\rangle, |g_{n\beta}\rangle$ are abstract vectors representing the functions $f_{n\alpha}(\mu), g_{n\beta}(\mu)$, and $|\alpha, f_{n\alpha}\rangle$, $\langle\beta, g_{n\beta}|$ are shorthands for representing the tensor product vectors $|\alpha\rangle \otimes |f_{n\alpha}\rangle$ and $\langle\beta| \otimes \langle g_{n\beta}|$ respectively. Define the tensor product space $Y := \mathbb{R}^{N_c^2} \otimes L^2(\mathbb{R}^d)$ and the operators $U_n : Y \to X$, $V_n : X \to Y$, and $C : Y \to Y$ by

$$U_n = \sum_{\alpha,\beta=0}^{N_c-1} A_{n\alpha\beta}|\alpha, f_{n\alpha}\rangle\langle N_c\alpha + \beta| \otimes I_{L^2(\mathbb{R}^d)} \qquad \text{[S29]}$$

$$V_n = \sum_{\alpha,\beta=0}^{N_c-1} |N_c\alpha + \beta\rangle\langle\beta, g_{n\beta}| \otimes I_{L^2(\mathbb{R}^d)} \qquad \text{[S30]}$$

$$C = \sum_{\gamma=0}^{N_c^2-1} |\gamma\rangle\langle\gamma| \otimes (\tilde{\sigma}_\gamma^{-2} - \nabla^2) \qquad \text{[S31]}$$

where we defined $\tilde{\sigma}_{N_c\alpha+\beta} := \sigma_{\alpha\beta}$, and $I_{L^2(\mathbb{R}^d)}$ is the identity operator on the Hilbert space $L^2(\mathbb{R}^d)$. Note that $V_n$ maps the $N_c$ discrete dimensions into $N_c^2$ discrete dimensions, and $U_n$ maps $N_c^2$ to $N_c$ discrete dimensions. $U_n, V_n$ are 'sparse' along the discrete dimensions since from their definition we see that only $N_c^2$ out of $N_c^3$ elements are non-zero. The reason we need to go to a higher dimensional space is that we have $N_c^2$ unique connectivity widths $\sigma_{\alpha\beta}$ which we need to put along the 'diagonal' of the operator $C$. This is the 'trick' that allows us to generalize the derivation for the simplified model in the main text, where we assumed the connectivity width depends only on the presynaptic cell type so that there were only $N$ unique connectivity widths. With this definition of $U_n$, $V_n$ and $C$, we get $W_n = U_n C^{-1} V_n$. Indeed,

$$\begin{aligned}
U_n C^{-1} V_n &= \sum_{\alpha,\beta,\alpha',\beta'=0}^{N_c-1} \sum_{\gamma=0}^{N_c^2-1} A_{n\alpha\beta'} |\alpha, f_{n\alpha}\rangle \langle N_c\alpha + \beta' \,|\, \gamma\rangle\langle\gamma \,|\, N_c\alpha' + \beta\rangle\langle\beta, g_{n\beta}| \otimes (\tilde{\sigma}_\gamma^{-2} - \nabla^2)^{-1} \\
&= \sum_{\alpha,\beta,\alpha',\beta'=0}^{N_c-1} A_{n\alpha\beta'} |\alpha, f_{n\alpha}\rangle \langle N_c\alpha + \beta' \,|\, N_c\alpha' + \beta\rangle\langle\beta, g_{n\beta}| \otimes (\tilde{\sigma}_{N_c\alpha+\beta'}^{-2} - \nabla^2)^{-1} \\
&= \sum_{\alpha,\beta=0}^{N_c-1} A_{n\alpha\beta} |\alpha, f_{n\alpha}\rangle\langle\beta, g_{n\beta}| \otimes (\sigma_{\alpha\beta}^{-2} - \nabla^2)^{-1}
\end{aligned}$$

where we used the fact that

$$\langle N_c\alpha + \beta' \,|\, N_c\alpha' + \beta\rangle = \delta_{\alpha\alpha'}\delta_{\beta'\beta}.$$

We now proceed to compute the inverse of $C - V_nU_n$. First, let us define the linear operator $\Sigma : \mathbb{R}^{N_c^2} \to \mathbb{R}^{N_c^2}$ by

$$\Sigma = \sum_{\gamma=0}^{N_c^2-1} \tilde{\sigma}_\gamma^2 |\gamma\rangle\langle\gamma| \quad \text{[S32]}$$

so that we can write $C$ as

$$C = \Sigma^{-1} \otimes I_{L^2(\mathbb{R}^d)} - I_{N_c^2} \otimes \nabla^2 \quad \text{[S33]}$$

where $I_{N_c^2}$ is the identity operator on $\mathbb{R}^{N_c^2}$. Let us also define the linear operators $\tilde{U}_n : \mathbb{R}^{N_c^2} \to \mathbb{R}^{N_c}$, $\tilde{V}_n : \mathbb{R}^{N_c} \to \mathbb{R}^{N_c^2}$, $K_n : \mathbb{R}^{N_c} \to \mathbb{R}^{N_c}$ by

$$\tilde{U}_n = \sum_{\alpha,\beta=0}^{N_c-1} A_{n\alpha\beta} |\alpha\rangle\langle N_c\alpha + \beta| \quad \text{[S34]}$$

$$\tilde{V}_n = \sum_{\alpha,\beta=0}^{N_c-1} |N_c\alpha + \beta\rangle\langle\beta| \quad \text{[S35]}$$

$$K_n = \sum_{\alpha=0}^{N_c-1} \langle g_{n\alpha} \,|\, f_{n\alpha}\rangle |\alpha\rangle\langle\alpha| \quad \text{[S36]}$$

so that the product $V_nU_n$ can be simplified as

$$\begin{aligned}
V_nU_n &= \left(\sum_{\alpha,\beta=0}^{N_c-1} |N_c\alpha + \beta\rangle\langle\beta, g_{n\beta}|\right)\left(\sum_{\alpha,\beta=0}^{N_c-1} A_{n\alpha\beta}|\alpha, f_{n\alpha}\rangle\langle N_c\alpha + \beta|\right) \otimes I_{L^2(\mathbb{R}^d)} \\
&= \left(\sum_{\alpha,\beta=0}^{N_c-1} |N_c\alpha + \beta\rangle\langle\beta|\right)\left(\sum_{\alpha=0}^{N_c-1} \langle g_{n\alpha} \,|\, f_{n\alpha}\rangle|\alpha\rangle\langle\alpha|\right)\left(\sum_{\alpha,\beta=0}^{N_c-1} A_{n\alpha\beta}|\alpha\rangle\langle N_c\alpha + \beta|\right) \otimes I_{L^2(\mathbb{R}^d)} \\
&= \tilde{V}_n K_n \tilde{U}_n \otimes I_{L^2(\mathbb{R}^d)}.
\end{aligned} \quad \text{[S37]}$$

Combining Eq. S33 and Eq. S37, we thus get

$$C - V_nU_n = (\Sigma^{-1} - \tilde{V}_n K_n \tilde{U}_n) \otimes I_{L^2(\mathbb{R}^d)} - I_{N_c^2} \otimes \nabla^2. \quad \text{[S38]}$$

Let $\boldsymbol{\Sigma}, \tilde{\boldsymbol{V}}_n, \boldsymbol{K}_n, \tilde{\boldsymbol{U}}_n$ be the matrix representations of $\Sigma, \tilde{V}_n, K_n, \tilde{U}_n$ in the standard basis respectively, such that $\tilde{\boldsymbol{U}}_n \in \mathbb{R}^{N_c \times N_c^2}, \tilde{\boldsymbol{V}}_n \in \mathbb{R}^{N_c^2 \times N_c}$ are given by

$$\tilde{U}_{n\alpha\gamma} = \sum_{\beta=0}^{N_c-1} A_{n\alpha\beta}\delta_{N_c\alpha+\beta,\gamma}, \quad \tilde{V}_{n\gamma\beta} = \sum_{\alpha=0}^{N_c-1} \delta_{N_c\alpha+\beta,\gamma} \tag{S39}$$

$\boldsymbol{K}_n \in \mathbb{R}^{N_c \times N_c}$ is given by

$$K_{0\alpha\beta} = \delta_{\alpha\beta}, \quad K_{1\alpha\beta} = \delta_{\alpha\beta}\int_0^1 f_\beta(\mu)g_\beta(\mu)P_\beta(\mu)\,d\mu, \tag{S40}$$

and $\boldsymbol{\Sigma} \in \mathbb{R}^{N_c^2 \times N_c^2}$ is given by

$$\Sigma_{\gamma\gamma'} = \delta_{\gamma\gamma'} \sum_{\alpha,\beta=0}^{N_c-1} \sigma_{\alpha\beta}^2 \delta_{N_c\alpha+\beta,\gamma}. \tag{S41}$$

Assume that $\boldsymbol{\Sigma}^{-1} - \tilde{\boldsymbol{V}}_n\boldsymbol{K}_n\tilde{\boldsymbol{U}}_n$ is diagonalizable as $\boldsymbol{Q}_n\boldsymbol{\Lambda}_n\boldsymbol{Q}_n^{-1}$. Let $\lambda_{n\rho}$ be the eigenvalues of $\boldsymbol{\Sigma}^{-1} - \tilde{\boldsymbol{V}}_n\boldsymbol{K}_n\tilde{\boldsymbol{U}}_n$, *i.e.*

$$\lambda_{n\rho} = [\boldsymbol{\Lambda}_n]_{\rho\rho}, \tag{S42}$$

and define $|\rho\rangle_{\mathcal{B}} = \sum_{\gamma=0}^{N_c^2-1}[\boldsymbol{Q}_n]_{\gamma\rho}|\gamma\rangle$, $\langle\rho|_{\mathcal{B}} = \sum_{\gamma=0}^{N_c^2-1}[\boldsymbol{Q}_n^{-1}]_{\rho\gamma}\langle\gamma|$. Then $\Sigma^{-1} - \tilde{V}_nK_n\tilde{U}_n = \sum_{\rho=0}^{N_c^2-1}\lambda_{n\rho}|\rho\rangle_{\mathcal{B}}\langle\rho|_{\mathcal{B}}$, and $I_{N_c^2} = \sum_{\rho=0}^{N_c^2-1}|\rho\rangle_{\mathcal{B}}\langle\rho|_{\mathcal{B}}$. Thus

$$\begin{aligned} C - V_nU_n &= (\Sigma^{-1} - \tilde{V}_nK_n\tilde{U}_n) \otimes I_{L^2(\mathbb{R}^d)} - I_{N_c^2} \otimes \nabla^2 \\ &= \sum_{\rho=0}^{N_c^2-1} \lambda_{n\rho}|\rho\rangle_{\mathcal{B}}\langle\rho|_{\mathcal{B}} \otimes I_{L^2(\mathbb{R}^d)} - I_{N_c^2} \otimes \nabla^2 \\ &= \sum_{\rho=0}^{N_c^2-1} |\rho\rangle_{\mathcal{B}}\langle\rho|_{\mathcal{B}} \otimes (\lambda_{n\rho} - \nabla^2) \end{aligned} \tag{S43}$$

and so we have

$$\begin{aligned} L_n &= I_X + U_n(C - V_nU_n)^{-1}V_n \\ &= I_X + U_n\left(\sum_{\rho=0}^{N_c^2-1}|\rho\rangle_{\mathcal{B}}\langle\rho|_{\mathcal{B}} \otimes (\lambda_{n\rho} - \nabla^2)^{-1}\right)V_n \end{aligned} \tag{S44}$$

where we used the property that $\langle\rho\,|\,\rho'\rangle_{\mathcal{B}} = \delta_{\rho\rho'}$ for obtaining the inverse of $C - V_nU_n$. The existence of the inverse $(\lambda_{n\rho} - \nabla^2)^{-1}$ requires $\lambda_{n\rho} \notin (-\infty, 0]$ for $\rho \in \mathbb{Z}_{N_c^2-1}$ (Proposition 2.2), but as we show in section 7, this is guaranteed for stable networks. Finally, plugging Eq. S44 back into Eq. S25 we get

$$\begin{aligned} L &= I + \sum_{n=-1}^{1}(L_n - I_X) \otimes |n\rangle\langle n| \\ &= I + \sum_{n=-1}^{1} U_n\left(\sum_{\rho=0}^{N_c^2-1}|\rho\rangle_{\mathcal{B}}\langle\rho|_{\mathcal{B}} \otimes (\lambda_{n\rho} - \nabla^2)^{-1}\right)V_n \otimes |n\rangle\langle n|. \end{aligned} \tag{S45}$$

Having computed the linear response operator, we can now compute the linear response of neuron $(\alpha, \mu, \boldsymbol{x}, \theta)$ to a unit-amplitude perturbation of neuron $(\beta, \nu, \boldsymbol{y}, \phi)$. When $(\alpha, \boldsymbol{x}, \theta, \mu) \neq (\beta, \boldsymbol{y}, \phi, \nu)$, this is represented by the function $\tilde{L}_{\alpha\beta}(\mu, \nu, \boldsymbol{x} - \boldsymbol{y}, \theta - \phi)$, which we define as the solution $r_\alpha(\mu, \boldsymbol{x}, \theta)$ of the steady state Eq. S19 with external input $h_\alpha(\mu, \boldsymbol{x}, \theta) = hP_\alpha(\mu)^{-1}\delta_{\alpha\beta}\delta(\mu - \nu)\delta(\boldsymbol{x} - \boldsymbol{y})\delta(\theta - \phi)$ (Eq. 11 in the main text), where the scalar parameter $h$ is set to 1 and the factor of $P_\alpha(\mu)^{-1}$ ensures the total input $\int_0^1\int_{\mathbb{R}^d}\int_{-\pi}^{\pi} h_\alpha(\mu, \boldsymbol{x}, \theta)P_\alpha(\mu)\,d\theta d\boldsymbol{x}d\mu$ is independent of $\mu$. In terms of the linear operator $L = (I - W)^{-1}$, it can be written as

$$\tilde{L}_{\alpha\beta}(\mu, \nu, \boldsymbol{x} - \boldsymbol{y}, \theta - \phi) = P_\beta(\nu)^{-1}\langle\alpha, \mu, \boldsymbol{x}, \theta|L - I|\beta, \nu, \boldsymbol{y}, \phi\rangle \tag{S46}$$

where the factor of $P_\beta(\nu)^{-1}$ comes from the definition of $h_\alpha(\mu, \boldsymbol{x}, \theta)$. The identity operator can be subtracted from $L$ since we specified that $(\alpha, \boldsymbol{x}, \theta, \mu) \neq (\beta, \boldsymbol{y}, \phi, \nu)$. We thus have

$$\begin{aligned}
\tilde{L}_{\alpha\beta}(\mu, \nu, \boldsymbol{x}-\boldsymbol{y}, \theta-\phi) &= P_\beta(\nu)^{-1}\langle \alpha, \mu, \boldsymbol{x}, \theta | L - I | \beta, \nu, \boldsymbol{y}, \phi\rangle \\
&= \frac{P_\beta(\nu)^{-1}}{2\pi} \sum_{n=-1}^{1} e^{in(\theta-\phi)} \sum_{\gamma,\gamma',\rho=0}^{N_c^2-1} [\boldsymbol{Q}_n]_{\gamma\rho}[\boldsymbol{Q}_n^{-1}]_{\rho\gamma'} \langle \alpha, \mu, \boldsymbol{x} | U_n \left(|\gamma\rangle\langle\gamma'| \otimes (\lambda_{n\rho} - \nabla^2)^{-1}\right) V_n | \beta, \nu, \boldsymbol{y}\rangle \\
&= \frac{P_\beta(\nu)^{-1}}{2\pi} \sum_{n=-1}^{1} \sum_{\gamma,\gamma',\rho=0}^{N_c^2-1} f_{n\alpha}(\mu)[\tilde{\boldsymbol{U}}_n]_{\alpha\gamma}[\boldsymbol{Q}_n]_{\gamma\rho}[\boldsymbol{Q}_n^{-1}]_{\rho\gamma'}[\tilde{\boldsymbol{V}}_n]_{\gamma'\beta} g_{n\beta}(\nu) G_d(s; \lambda_{n\rho}) e^{in(\theta-\phi)} \\
&= \frac{P_\beta(\nu)^{-1}}{2\pi} \sum_{n=-1}^{1} \sum_{\rho=0}^{N_c^2-1} [\tilde{\boldsymbol{U}}_n \boldsymbol{Q}_n]_{\alpha\rho}[\boldsymbol{Q}_n^{-1}\tilde{\boldsymbol{V}}_n]_{\rho\beta} f_{n\alpha}(\mu) g_{n\beta}(\nu) G_d(s; \lambda_{n\rho}) e^{in(\theta-\phi)} \\
&= \frac{1}{2\pi} \sum_{\rho=0}^{N_c^2-1} \Bigg( [\tilde{\boldsymbol{U}}_0 \boldsymbol{Q}_0]_{\alpha\rho}[\boldsymbol{Q}_0^{-1}\tilde{\boldsymbol{V}}_0]_{\rho\beta} G_d(s; \lambda_{0\rho}) \\
&\qquad + 2[\tilde{\boldsymbol{U}}_1 \boldsymbol{Q}_1]_{\alpha\rho}[\boldsymbol{Q}_1^{-1}\tilde{\boldsymbol{V}}_1]_{\rho\beta} f_\alpha(\mu) g_\beta(\nu) G_d(s; \lambda_{1\rho}) \cos(\theta-\phi) \Bigg).
\end{aligned} \quad \text{[S47]}$$

In particular, if we let $\tilde{L}_{n\alpha\beta}(s)$ be the distance-dependent parts of this linear response kernel, i.e. define

$$\tilde{L}_{n\alpha\beta}(s) = \sum_{\rho=0}^{N_c^2-1} l_{n\alpha\beta\rho} G_d(s; \lambda_{n\rho}) \quad \text{[S48]}$$

$$l_{n\alpha\beta\rho} = [\tilde{\boldsymbol{U}}_n \boldsymbol{Q}_n]_{\alpha\rho}[\boldsymbol{Q}_n^{-1}\tilde{\boldsymbol{V}}_n]_{\rho\beta} \quad \text{[S49]}$$

then we can write Eq. S47 as

$$\tilde{L}_{\alpha\beta}(\mu, \nu, \boldsymbol{x}-\boldsymbol{y}, \theta-\phi) = \frac{1}{2\pi}\left(\tilde{L}_{0\alpha\beta}(s) + 2\tilde{L}_{1\alpha\beta}(s) f_\alpha(\mu) g_\beta(\nu)\cos(\theta-\phi)\right) \quad \text{[S50]}$$

which is equivalent to Eq. 9 given in the main text.

#### 5. Mean perturbation response over space and/or feature-tuning

In some of our analysis we will study the response of neurons averaged over space or feature-tuning. Thus in this section we compute the integral of $\tilde{L}_{\alpha\beta}(\mu, \nu, \boldsymbol{x}-\boldsymbol{y}, \theta-\phi)$ (Eq. S50) over space and feature-tuning respectively. The integral over $\theta, \mu$ is trivial: since the integral $\int_{-\pi}^{\pi} \cos(\theta-\phi) d\theta = 0$, the cosine term in Eq. S50 vanishes, so we have

$$\tilde{L}_{\alpha\beta}(\boldsymbol{x}-\boldsymbol{y}) = \tilde{L}_{0\alpha\beta}(s) = \sum_{\rho=0}^{N_c^2-1} [\tilde{\boldsymbol{U}}_0 \boldsymbol{Q}_0]_{\alpha\rho}[\boldsymbol{Q}_0^{-1}\tilde{\boldsymbol{V}}_0]_{\rho\beta} G_d(s; \lambda_{0\rho}). \quad \text{[S51]}$$

On the other hand, for the integral over $\boldsymbol{x}$, we note that since $\int_{\mathbb{R}^d} f(\boldsymbol{x})\, d\boldsymbol{x} = \hat{f}(\boldsymbol{0})$ for any function $f$ with Fourier transform $\hat{f}$, and since $\hat{G}_d(\boldsymbol{k}; \lambda) = (\lambda + k^2)^{-1}$ (Eq. S11), we have

$$\int_{\mathbb{R}^d} G_d(\|\boldsymbol{x}-\boldsymbol{y}\|; \lambda)\, d\boldsymbol{x} = \lambda^{-1}.$$

Thus the integral of Eq. S48 over space is given by $[\tilde{\boldsymbol{U}}_n \boldsymbol{Q}_n \boldsymbol{\Lambda}_n^{-1} \boldsymbol{Q}_n^{-1} \tilde{\boldsymbol{V}}_n]_{\alpha\beta} = [\tilde{\boldsymbol{U}}_n(\boldsymbol{\Sigma}^{-1} - \tilde{\boldsymbol{V}}_n \boldsymbol{K}_n \tilde{\boldsymbol{U}}_n)^{-1}\tilde{\boldsymbol{V}}_n]_{\alpha\beta}$, so the integral of Eq. S50 over $\boldsymbol{x}$ is

$$\tilde{L}_{\alpha\beta}(\mu, \nu, \theta-\phi) = \frac{1}{2\pi}\left([\tilde{\boldsymbol{U}}_0(\boldsymbol{\Sigma}^{-1} - \tilde{\boldsymbol{V}}_0 \boldsymbol{K}_0 \tilde{\boldsymbol{U}}_0)^{-1}\tilde{\boldsymbol{V}}_0]_{\alpha\beta} + 2[\tilde{\boldsymbol{U}}_1(\boldsymbol{\Sigma}^{-1} - \tilde{\boldsymbol{V}}_1 \boldsymbol{K}_1 \tilde{\boldsymbol{U}}_1)^{-1}\tilde{\boldsymbol{V}}_1]_{\alpha\beta}\cos(\theta-\phi) f_\alpha(\mu) g_\beta(\nu)\right). \quad \text{[S52]}$$

By the Woodbury matrix identity, $(\boldsymbol{I} - \boldsymbol{UCV})^{-1} = \boldsymbol{I} + \boldsymbol{U}(\boldsymbol{C}^{-1} - \boldsymbol{VU})^{-1}\boldsymbol{V}$, so if we apply it in reverse, we get

$$\tilde{\boldsymbol{U}}_n(\boldsymbol{\Sigma}^{-1} - \tilde{\boldsymbol{V}}_n \boldsymbol{K}_n \tilde{\boldsymbol{U}}_n)^{-1}\tilde{\boldsymbol{V}}_n = \boldsymbol{K}_n^{-1}((\boldsymbol{I} - \boldsymbol{K}_n \tilde{\boldsymbol{U}}_n \boldsymbol{\Sigma} \tilde{\boldsymbol{V}}_n)^{-1} - \boldsymbol{I}).$$

Let us define $\boldsymbol{W}$ as the matrix of elements $w_{\alpha\beta}$ and $\tilde{\boldsymbol{W}}$ as the matrix of elements $w_{\alpha\beta}\kappa_{\alpha\beta}$. From the definitions of $\tilde{\boldsymbol{U}}_n, \tilde{\boldsymbol{V}}_n$, one can compute that $\tilde{\boldsymbol{U}}_0\boldsymbol{\Sigma}\tilde{\boldsymbol{V}}_0 = \boldsymbol{W}$ and $\tilde{\boldsymbol{U}}_1\boldsymbol{\Sigma}\tilde{\boldsymbol{V}}_1 = \tilde{\boldsymbol{W}}$. Furthermore, since $\boldsymbol{K}_0$ is just the identity matrix, let us write $\boldsymbol{K}$ to denote $\boldsymbol{K}_1$, and let $k_\alpha$ be the diagonal entries of $\boldsymbol{K}$, so that

$$k_\alpha = \langle g_{1\alpha} \,|\, f_{1\alpha}\rangle = \int_0^1 f_\alpha(\mu) g_\alpha(\mu) P_\alpha(\mu)\, d\mu. \tag{S53}$$

Thus we have

$$\tilde{L}_{\alpha\beta}(\mu,\nu,\theta-\phi) = \frac{1}{2\pi}\left([(\boldsymbol{I}-\boldsymbol{W})^{-1}-\boldsymbol{I}]_{\alpha\beta} + 2[(\boldsymbol{I}-\boldsymbol{K}\tilde{\boldsymbol{W}})^{-1}-\boldsymbol{I}]_{\alpha\beta} k_\alpha^{-1} f_\alpha(\mu) g_\beta(\nu)\cos(\theta-\phi)\right). \tag{S54}$$

Finally, consider the response integrated over both space and feature-tuning. Clearly, since $\int_{-\pi}^{\pi}\cos(\theta-\phi)d\theta = 0$, integrating Eq. S54 over feature-tuning yields

$$\tilde{L}_{\alpha\beta} = [(\boldsymbol{I}-\boldsymbol{W})^{-1}-\boldsymbol{I}]_{\alpha\beta}. \tag{S55}$$

Notice that Eq. S55 is equivalent to the linear response of a model without space and feature-tuning. This is not a coincidence. In fact, since our connectivity function is translationally invariant in $\boldsymbol{x}$ and $\theta$, Eq. S51 is equivalent to the linear response of a model without feature-tuning, while Eq. S54 is equivalent to the linear response of a model without space. To see this, simply integrate both sides of Eq. 1 over $\boldsymbol{x}$ and $\theta$ respectively.

#### 6. Symmetries in connectivity width simplifies perturbation response

The full linear response Eq. S50 involves a sum of $2N_c^2$ terms. However, in the main text we saw that for a simplified model where there is no feature tuning and whose connectivity width depends only on presynaptic, its linear response only consists of $N_c$ terms. Thus we would also expect Eq. S50 to reduce to a sum of $2N_c$ terms if we impose this symmetry condition of connectivity width on the model. Indeed, if we modify the derivation in section 4 by replacing the operators $U_n$, $V_n$, and $C$ with

$$U_n = \sum_{\alpha,\beta=0}^{N_c-1} A_{n\alpha\beta}|\alpha, f_{n\alpha}\rangle\langle\beta| \otimes I_{L^2(\mathbb{R}^d)} \tag{S56}$$

$$V_n = \sum_{\alpha,\beta=0}^{N_c-1} |\alpha\rangle\langle\beta, g_{n\beta}| \otimes I_{L^2(\mathbb{R}^d)} \tag{S57}$$

$$C = \sum_{\gamma=0}^{N_c-1} |\gamma\rangle\langle\gamma| \otimes (\tilde{\sigma}_\gamma^{-2} - \nabla^2)^{-1} \tag{S58}$$

and propagating the changes, one can show that the simplified linear response kernel is given by

$$\tilde{L}_{\alpha\beta}(\mu,\nu,\boldsymbol{x}-\boldsymbol{y},\theta-\phi) = \frac{1}{2\pi}\left(\tilde{L}_{0\alpha\beta}(s) + 2\tilde{L}_{1\alpha\beta}(s) f_\alpha(\mu) g_\beta(\nu)\cos(\theta-\phi)\right) \tag{S59}$$

where

$$\tilde{L}_{0\alpha\beta}(s) = \sum_{\rho=0}^{N_c-1} [\boldsymbol{W}\tilde{\boldsymbol{\Sigma}}^{-1}\tilde{\boldsymbol{Q}}_0]_{\alpha\rho}[\tilde{\boldsymbol{Q}}_0^{-1}]_{\rho\beta} G_d(s;\tilde{\lambda}_{0\rho}) \tag{S60}$$

$$\tilde{L}_{1\alpha\beta}(s) = \sum_{\rho=0}^{N_c-1} [\tilde{\boldsymbol{W}}\tilde{\boldsymbol{\Sigma}}^{-1}\tilde{\boldsymbol{Q}}_1]_{\alpha\rho}[\tilde{\boldsymbol{Q}}_1^{-1}]_{\rho\beta} G_d(s;\tilde{\lambda}_{1\rho}) \tag{S61}$$

where $\tilde{\boldsymbol{W}}$ is as defined in the previous section, $\tilde{\boldsymbol{\Sigma}} \in \mathbb{R}^{N_c\times N_c}$ is a diagonal matrix of elements $\sigma_{0\beta}^2$, and $\tilde{\boldsymbol{Q}}_n, \tilde{\boldsymbol{\Lambda}}_n$ (where $\tilde{\boldsymbol{\Lambda}}_n$ is the diagonal matrix of elements $\tilde{\lambda}_{n\rho}$) are defined by the diagonalizations $\tilde{\boldsymbol{Q}}_0\tilde{\boldsymbol{\Lambda}}_0\tilde{\boldsymbol{Q}}_0^{-1} = (\boldsymbol{I}-\boldsymbol{W})\tilde{\boldsymbol{\Sigma}}^{-1}$, $\tilde{\boldsymbol{Q}}_1\tilde{\boldsymbol{\Lambda}}_1\tilde{\boldsymbol{Q}}_1^{-1} = (\boldsymbol{I}-\boldsymbol{K}\tilde{\boldsymbol{W}})\tilde{\boldsymbol{\Sigma}}^{-1}$. For convenience, let us also define $\tilde{\boldsymbol{W}}_n$ such that $\tilde{\boldsymbol{W}}_0 = \boldsymbol{W}$ and $\tilde{\boldsymbol{W}}_1 = \tilde{\boldsymbol{W}}$, so we have $\tilde{\boldsymbol{Q}}_n\tilde{\boldsymbol{\Lambda}}_n\tilde{\boldsymbol{Q}}_n^{-1} = (\boldsymbol{I}-\boldsymbol{K}_n\tilde{\boldsymbol{W}}_n)\tilde{\boldsymbol{\Sigma}}^{-1}$.

It is not immediately clear how the $2N_c^2$ terms in Eq. S50 can be simplified to the $2N_c$ terms in Eq. S59 under the symmetry $\sigma_{\alpha\beta} = \sigma_{0\beta}$. Here we show that under this symmetry, $N_c(N_c-1)$ eigenvalues of the matrix $\boldsymbol{\Sigma}^{-1} - \tilde{\boldsymbol{V}}_n\boldsymbol{K}_n\tilde{\boldsymbol{U}}_n$

are equal to the $N_c$ elements $\sigma_{0\beta}^{-2}$ with multiplicity $N_c - 1$, and that the terms corresponding to these eigenvalues vanish due to having a coefficient of 0. Furthermore, the remaining $N_c$ eigenvalues of $\boldsymbol{\Sigma}^{-1} - \tilde{\boldsymbol{V}}_n \boldsymbol{K}_n \tilde{\boldsymbol{U}}_n$ are exactly equal to the eigenvalues of $(\boldsymbol{I} - \boldsymbol{K}_n \tilde{\boldsymbol{W}}_n)\tilde{\boldsymbol{\Sigma}}^{-1}$.

First we prove the relationship between the eigenvalues of $\boldsymbol{\Sigma}^{-1} - \tilde{\boldsymbol{V}}_n \boldsymbol{K}_n \tilde{\boldsymbol{U}}_n$ and $(\boldsymbol{I} - \boldsymbol{K}_n \tilde{\boldsymbol{W}}_n)\tilde{\boldsymbol{\Sigma}}^{-1}$. Consider the characteristic polynomial of $\boldsymbol{\Sigma}^{-1} - \tilde{\boldsymbol{V}}_n \boldsymbol{K}_n \tilde{\boldsymbol{U}}_n$, given by

$$p_n(\lambda) = \det(\lambda \boldsymbol{I} - \boldsymbol{\Sigma}^{-1} + \tilde{\boldsymbol{V}}_n \boldsymbol{K}_n \tilde{\boldsymbol{U}}_n). \tag{S62}$$

By Sylvester's determinant theorem (17, p. 271), $\det(\boldsymbol{A} + \boldsymbol{V}\boldsymbol{U}) = \det(\boldsymbol{I} + \boldsymbol{U}\boldsymbol{A}^{-1}\boldsymbol{V})\det(\boldsymbol{A})$ for matrices $\boldsymbol{A}, \boldsymbol{V}, \boldsymbol{U}$ (assuming $\boldsymbol{A}$ is invertible). Thus, if we let $\boldsymbol{A} = \lambda\boldsymbol{I} - \boldsymbol{\Sigma}^{-1}$, $\boldsymbol{V} = \tilde{\boldsymbol{V}}_n$, $\boldsymbol{U} = \boldsymbol{K}_n \tilde{\boldsymbol{U}}_n$, then we get

$$p_n(\lambda) = \det(\boldsymbol{I} + \boldsymbol{K}_n \tilde{\boldsymbol{U}}_n (\lambda\boldsymbol{I} - \boldsymbol{\Sigma}^{-1})^{-1} \tilde{\boldsymbol{V}}_n) \det(\lambda\boldsymbol{I} - \boldsymbol{\Sigma}^{-1}). \tag{S63}$$

Now due to the symmetry condition, it can be shown that

$$(\lambda\boldsymbol{I} - \boldsymbol{\Sigma}^{-1})^{-1} \tilde{\boldsymbol{V}}_n = \tilde{\boldsymbol{V}}_n (\lambda\boldsymbol{I} - \tilde{\boldsymbol{\Sigma}}^{-1})^{-1}. \tag{S64}$$

Indeed, for all $\alpha, \beta, \beta' \in \mathbb{Z}_{N_c}$, by definition of $\tilde{\boldsymbol{V}}_n$ we have

$$[\tilde{\boldsymbol{V}}_n]_{N_c\alpha+\beta,\beta'} = \sum_{\gamma,\gamma'=0}^{N_c-1} \langle N_c\alpha + \beta \,|\, N_c\gamma + \gamma' \rangle \langle \gamma' \,|\, \beta' \rangle = \sum_{\gamma,\gamma'=0}^{N_c-1} \delta_{\alpha\gamma}\delta_{\beta\gamma'}\delta_{\gamma'\beta'} = \delta_{\beta\beta'}$$

so that

$$[(\lambda\boldsymbol{I} - \boldsymbol{\Sigma}^{-1})^{-1}\tilde{\boldsymbol{V}}_n]_{N_c\alpha+\beta,\beta'} = (\lambda - \sigma_\beta^{-2})^{-1}\delta_{\beta\beta'} = \delta_{\beta\beta'}(\lambda - \sigma_{\beta'}^{-2})^{-1} = [\tilde{\boldsymbol{V}}_n(\lambda\boldsymbol{I} - \tilde{\boldsymbol{\Sigma}}^{-1})^{-1}]_{N_c\alpha+\beta,\beta'}.$$

Thus

$$\begin{aligned} p_n(\lambda) &= \det(\boldsymbol{I} + \boldsymbol{K}_n \tilde{\boldsymbol{U}}_n \tilde{\boldsymbol{V}}_n (\lambda\boldsymbol{I} - \tilde{\boldsymbol{\Sigma}}^{-1})^{-1}) \det(\lambda\boldsymbol{I} - \boldsymbol{\Sigma}^{-1}) \\ &= \det(\lambda\boldsymbol{I} - \tilde{\boldsymbol{\Sigma}}^{-1} + \boldsymbol{K}_n \tilde{\boldsymbol{U}}_n \tilde{\boldsymbol{V}}_n) \det(\lambda\boldsymbol{I} - \tilde{\boldsymbol{\Sigma}}^{-1})^{-1} \det(\lambda\boldsymbol{I} - \boldsymbol{\Sigma}^{-1}). \end{aligned} \tag{S65}$$

But under this symmetry condition, $\tilde{\boldsymbol{U}}_n \tilde{\boldsymbol{V}}_n = \tilde{\boldsymbol{W}}_n \tilde{\boldsymbol{\Sigma}}^{-1}$ and $\det(\lambda\boldsymbol{I} - \boldsymbol{\Sigma}^{-1}) = \det(\lambda\boldsymbol{I} - \tilde{\boldsymbol{\Sigma}}^{-1})^{N_c}$. Thus

$$p_n(\lambda) = \det(\lambda\boldsymbol{I} - (\boldsymbol{I} - \boldsymbol{K}_n \tilde{\boldsymbol{W}}_n)\tilde{\boldsymbol{\Sigma}}^{-1}) \det(\lambda\boldsymbol{I} - \tilde{\boldsymbol{\Sigma}}^{-1})^{N_c - 1}, \tag{S66}$$

so the $N_c^2$ eigenvalues of $\boldsymbol{\Sigma}^{-1} - \tilde{\boldsymbol{V}}_n \boldsymbol{K}_n \tilde{\boldsymbol{U}}_n$ consists of precisely the $N_c$ eigenvalues of $(\boldsymbol{I} - \boldsymbol{K}_n \tilde{\boldsymbol{W}}_n)\tilde{\boldsymbol{\Sigma}}^{-1}$ as well as the $N_c$ elements $\sigma_{0\beta}^{-2}$ with multiplicity $N_c - 1$.

Now we will show that the terms corresponding to the eigenvalues $\sigma_{0\beta}^{-2}$ vanish under this symmetry condition. We claim that $[\boldsymbol{Q}_n^{-1}\tilde{\boldsymbol{V}}_n]_{\rho\alpha} = 0$ for all $\alpha \in \mathbb{Z}_{N_c}$ and $\rho \in \mathbb{Z}_{N_c^2}$ such that $\lambda_{n\rho} = \sigma_{0\beta}^{-2}$ for some $\beta \in \mathbb{Z}_{N_c}$. To see this, let $\boldsymbol{l}_{n\rho}^T$ be the $\rho$-th row of $\boldsymbol{Q}_n^{-1}$. Then $\boldsymbol{l}_{n\rho}^T$ is a left eigenvector of $\boldsymbol{\Sigma}^{-1} - \tilde{\boldsymbol{V}}_n \boldsymbol{K}_n \tilde{\boldsymbol{U}}_n$ with eigenvalue $\lambda_{n\rho}$ by Proposition 17.1. Thus, if $\lambda_{n\rho} = \sigma_{0\beta}^{-2}$, then $\boldsymbol{l}_{n\rho}^T(\sigma_{0\beta}^{-2}\boldsymbol{I} - \boldsymbol{\Sigma}^{-1} + \tilde{\boldsymbol{V}}_n \boldsymbol{K}_n \tilde{\boldsymbol{U}}_n) = \boldsymbol{0}^T$. Since the $(N_c\alpha + \beta)$-th diagonal element of $\boldsymbol{\Sigma}^{-1}$ is $\sigma_{0\beta}^{-2}$, we have $[\boldsymbol{l}_{n\rho}^T(\sigma_{0\beta}^{-2}\boldsymbol{I} - \boldsymbol{\Sigma}^{-1})]_{N_c\alpha+\beta} = 0$. Thus

$$\sum_{\alpha'=0}^{N_c-1} [\boldsymbol{l}_{n\rho}\tilde{\boldsymbol{V}}_n]_{\alpha'} [\boldsymbol{K}_n \tilde{\boldsymbol{U}}_n]_{\alpha', N_c\alpha+\beta} = 0. \tag{S67}$$

But by the definition of $\tilde{U}_n$,

$$[\tilde{\boldsymbol{U}}_n]_{\alpha', N_c\alpha+\beta} = \sum_{\gamma\gamma'=0}^{N_c-1} A_{n\gamma\gamma'} \langle \alpha' \,|\, \gamma \rangle \langle N_c\gamma + \gamma' \,|\, N_c\alpha + \beta \rangle = \sum_{\gamma\gamma'=0}^{N_c-1} A_{n\gamma\gamma'} \delta_{\alpha'\gamma}\delta_{\gamma\alpha}\delta_{\gamma'\beta} = A_{n\alpha'\beta}\delta_{\alpha'\alpha},$$

and since $\boldsymbol{K}_n$ is a diagonal matrix, if we let $k_{n\alpha}$ be its $\alpha$-th diagonal entry, then we have

$$\begin{aligned} \sum_{\alpha'=0}^{N_c-1} [\boldsymbol{l}_{n\rho}\tilde{\boldsymbol{V}}_n]_{\alpha'} k_{n\alpha'} A_{n\alpha'\beta}\delta_{\alpha'\alpha} &= 0 \\ [\boldsymbol{l}_{n\rho}\tilde{\boldsymbol{V}}_n]_{\alpha} k_{n\alpha} A_{n\alpha\beta} &= 0. \end{aligned} \tag{S68}$$

Thus, as long as $k_{n\alpha}A_{n\alpha\beta} \neq 0$, then we must have $[\boldsymbol{Q}_n^{-1}\tilde{\boldsymbol{V}}_n]_{\rho\alpha} = 0$. But by the continuity of eigenvectors with respect to each of its matrix elements (18), we conclude unconditionally that $[\boldsymbol{Q}_n^{-1}\tilde{\boldsymbol{V}}_n]_{\rho\alpha} = 0$.

#### 7. Eigenspectrum and stability

**A. Condition for stability.** In this section we consider the condition for the network dynamics given by Eq. 12 to be stable. First, let us write down the dynamic equation more abstractly as

$$T\frac{\mathrm{d}}{\mathrm{d}t}|r\rangle = -|r\rangle + W|r\rangle + |h\rangle \tag{S69}$$

where $T$ is the time constant operator given by $T = \sum_{\alpha=0}^{N_c-1} \tau_\alpha |\alpha\rangle\langle\alpha| \otimes I_Z$ where $I_Z$ is the identity operator on the tensor product space $Z := L^2([0,1]) \otimes L^2(\mathbb{R}^d) \otimes L^2(\mathbb{S}^1)$. Rewriting Eq. S69 as $\frac{\mathrm{d}}{\mathrm{d}t}|r\rangle = T^{-1}(W - I)|r\rangle + |h\rangle$, we see that if we define

$$J = T^{-1}(W - I) \tag{S70}$$

then this dynamical system is (exponentially) stable if and only if the spectrum $\sigma(J)$ of $J$ satisfies

$$\alpha(J) := \sup\{\mathrm{Re}(\lambda) : \lambda \in \sigma(J)\} < 0. \tag{S71}$$

where $\alpha(J)$ denotes the spectral abscissa of $J$. We note that we have glossed over some subtlety that arises when dealing with non-compact operators on infinite-dimensional vector spaces, see (19, Proposition 3.2).

We propose that the spectrum $\sigma(J)$ is given by

$$\sigma(J) = \bigcup_{k\in\mathbb{R}} \bigcup_{n=-1}^{1} \sigma(\boldsymbol{J}_n(k)) \tag{S72}$$

where $\boldsymbol{J}_n(k)$ is a $N_c \times N_c$ matrix with elements

$$J_{n\alpha\beta}(k) = \tau_\alpha^{-1}\left((1+\sigma_{\alpha\beta}^2 k^2)^{-1}\langle g_{n\beta}\,|\,f_{n\beta}\rangle \tilde{W}_{n\alpha\beta} - \delta_{\alpha\beta}\right) \tag{S73}$$

and $\tilde{W}_{n\alpha\beta}$ are the elements of the matrix $\tilde{\boldsymbol{W}}_n$ defined in previous sections and can be written as $A_{n\alpha\beta}\sigma_{\alpha\beta}^2$. Since a rigorous treatment of the spectrum of linear operators on infinite dimensional spaces requires advanced mathematical machinery, we will simply provide an informal argument of this claim based on intuition from operators on finite-dimensional spaces.

To see this, let us first define $|\boldsymbol{k}\rangle$ to be the Fourier mode of $\mathbb{R}^d$ with frequency $\boldsymbol{k}$, i.e. $\langle \boldsymbol{x}\,|\,\boldsymbol{k}\rangle = \frac{1}{(2\pi)^{\frac{d}{2}}}e^{i\boldsymbol{k}\cdot\boldsymbol{x}}$. Let $\boldsymbol{v}$ be an arbitrary vector in $\mathbb{C}^{N_c}$, and let $v_\alpha$ denote the elements of $\boldsymbol{v}$. We claim that

$$J\sum_{\alpha=0}^{N_c-1} v_\alpha|\alpha, f_{n\alpha}, \boldsymbol{k}, n\rangle = \sum_{\alpha,\beta=0}^{N_c-1} J_{n\alpha\beta}(k)v_\beta|\alpha, f_{n\alpha}, \boldsymbol{k}, n\rangle \tag{S74}$$

for all $n, \boldsymbol{k}$, where $k = \|\boldsymbol{k}\|$. Indeed, since

$$\begin{aligned}
W\sum_{\alpha=0}^{N_c-1} v_\alpha|\alpha, f_{n\alpha}, \boldsymbol{k}, n\rangle &= \left(\sum_{m=-1}^{1} W_m \otimes |m\rangle\langle m|\right)\sum_{\alpha=0}^{N_c-1} v_\alpha|\alpha, f_{n\alpha}, \boldsymbol{k}, n\rangle \\
&= \left(\sum_{\alpha=0}^{N_c-1} v_\alpha W_n|\alpha, f_{n\alpha}, \boldsymbol{k}\rangle\right)\otimes|n\rangle \\
&= \left(\sum_{\alpha=0}^{N_c-1} v_\alpha\left(\sum_{\alpha',\beta=0}^{N_c-1} A_{n\alpha'\beta}|\alpha', f_{n\alpha'}\rangle\langle\beta, g_{n\beta}| \otimes (\sigma_{\alpha'\beta}^{-2} - \nabla^2)^{-1}\right)|\alpha, f_{n\alpha}, \boldsymbol{k}\rangle\right)\otimes|n\rangle \\
&= \sum_{\alpha,\alpha',\beta=0}^{N_c-1} v_\alpha A_{n\alpha'\beta}\delta_{\alpha\beta}\langle g_{n\beta}\,|\,f_{n\alpha}\rangle(\sigma_{\alpha'\beta}^{-2} + k^2)^{-1}|\alpha', f_{n\alpha'}, \boldsymbol{k}, n\rangle \\
&= \sum_{\alpha',\beta=0}^{N_c-1} (1+\sigma_{\alpha'\beta}^2 k^2)^{-1}\langle g_{n\beta}\,|\,f_{n\beta}\rangle\tilde{W}_{n\alpha'\beta}v_\beta|\alpha', f_{n\alpha'}, \boldsymbol{k}, n\rangle,
\end{aligned} \tag{S75}$$

we have

$$J\sum_{\alpha=0}^{N_c-1} v_\alpha|\alpha, f_{n\alpha}, \boldsymbol{k}, n\rangle = \sum_{\alpha,\beta=0}^{N_c-1} \tau_\alpha^{-1}\left((1+\sigma_{\alpha\beta}^2 k^2)^{-1}\langle g_{n\beta} \mid f_{n\beta}\rangle \tilde{W}_{n\alpha\beta} - \delta_{\alpha\beta}\right) v_\beta|\alpha, f_{n\alpha}, \boldsymbol{k}, n\rangle$$

$$= \sum_{\alpha,\beta=0}^{N_c-1} J_{n\alpha\beta}(k) v_\beta|\alpha, f_{n\alpha}, \boldsymbol{k}, n\rangle.$$

Now let $\lambda_n(k)$ be an eigenvalue of $\boldsymbol{J}_n(k)$ with eigenvector $\boldsymbol{v}_n(k)$, and let $v_{n\alpha}(k)$ denote the $\alpha$-th element of $\boldsymbol{v}_n(k)$. We claim that $\lambda_n(k)$ is also an eigenvalue of $J$ with eigenvector $\sum_{\alpha=0}^{N_c-1} v_{n\alpha}(k)|\alpha, f_{n\alpha}, \boldsymbol{k}, n\rangle$ (rigorously speaking they are not actually eigenvalues and eigenvectors of $J$ since $|\boldsymbol{k}\rangle$ is not a square-integrable function and thus not a vector in $L^2(\mathbb{R}^d)$). Indeed, by Eq. S74, we have

$$J\sum_{\alpha=0}^{N_c-1} v_{n\alpha}(k)|\alpha, f_{n\alpha}, \boldsymbol{k}, n\rangle = \sum_{\alpha,\beta=0}^{N_c-1} J_{n\alpha\beta}(k) v_{n\beta}(k)|\alpha, f_{n\alpha}, \boldsymbol{k}, n\rangle = \lambda_n(k)\sum_{\alpha=0}^{N_c-1} v_{n\alpha}(k)|\alpha, f_{n\alpha}, \boldsymbol{k}, n\rangle, \tag{S76}$$

so $\lambda_n(k)$ is indeed an element of $\sigma(J)$. Conversely, suppose $\lambda$ is an eigenvalue of $J$ with eigenvector $|v\rangle$. In general, $|v\rangle$ can be written as $\sum_{\alpha=0}^{N_c-1}\sum_{n=-1}^{1}\int_{\mathbb{R}^d} d\boldsymbol{k}\, v_{n\alpha}(\boldsymbol{k})|\alpha, f_{n\alpha}, \boldsymbol{k}, n\rangle$ where $v_{n\alpha}(\boldsymbol{k}) \in \mathbb{C}$ for all $n, \alpha, \boldsymbol{k}$, so that by Eq. S74, we have

$$J|v\rangle = \sum_{\alpha=0}^{N_c-1}\sum_{n=-1}^{1}\int_{\mathbb{R}^d} d\boldsymbol{k} \sum_{\beta=0}^{N_c-1}\left(J_{n\alpha\beta}(k) v_{n\beta}(\boldsymbol{k})\right)|\alpha, f_{n\alpha}, \boldsymbol{k}, n\rangle = \lambda\sum_{\alpha=0}^{N_c-1}\sum_{n=-1}^{1}\int_{\mathbb{R}^d} d\boldsymbol{k}\, v_{n\alpha}(\boldsymbol{k})|\alpha, f_{n\alpha}, \boldsymbol{k}, n\rangle. \tag{S77}$$

Comparing coefficients, we see that we must have

$$\sum_{\beta=0}^{N_c-1} J_{n\alpha\beta}(k) v_{n\beta}(\boldsymbol{k}) = \lambda v_{n\alpha}(k) \tag{S78}$$

for all $n, \boldsymbol{k}$. Thus $\boldsymbol{v}_n(\boldsymbol{k})$ is either a zero vector or an eigenvector of $\boldsymbol{J}_n(k)$ with eigenvalue $\lambda$ for all $n, \boldsymbol{k}$. But $\boldsymbol{v}_n(\boldsymbol{k})$ must be nonzero for some $n, \boldsymbol{k}$ (otherwise $|v\rangle$ would be a zero vector and therefore not an eigenvector of $J$), so $\lambda$ must be an eigenvalue of $\boldsymbol{J}_n(k)$ for some $n, k$. This completes our informal argument.

Since we do not have a rigorous proof of Eq. S72, we validate it numerically by comparing the spectral abscissa and spectral radius computed using Eq. S72 and that of a large matrix obtained by discretizing $J$ in space and feature. The union over $k \in \mathbb{R}$ in Eq. S72 is approximated by computing the eigenvalues of $\boldsymbol{J}_n(k)$ for 1000 uniformly spaced values of $k \in \left[0, 100\frac{\sum_{\alpha\beta}\sigma_{\alpha\beta}^{-2}}{N_c^2}\right]$. The two approaches agree with each other numerically.

**B. Two necessary conditions.** The requirement of stability imposes constraints on the model parameters. However, the condition of stability is quite complicated as currently stated, so it is unclear what those constraints are. Here we consider two necessary conditions of stability which provide important constraints for our analysis of perturbation response in stable networks in other sections of this document.

First, notice that since $-\boldsymbol{J}_n(k)$ is a real matrix, its eigenvalues are either real or come in complex conjugate pairs. Since stability implies that the real part of every eigenvalue of $\boldsymbol{J}_n(k)$ is negative, the real eigenvalues of $-\boldsymbol{J}_n(k)$ are positive. Furthermore, since $z\bar{z} = |z|^2$, the product of each pair of complex conjugate eigenvalues is positive. Thus $\det(-\boldsymbol{J}_n(k)) > 0$ for all $k \in \mathbb{R}$, since the determinant is the product of eigenvalues. Let $\tilde{\boldsymbol{W}}_n$ be the matrix of elements $\tilde{W}_{n\alpha\beta}$, so that we can write $\boldsymbol{J}_n(0) = \boldsymbol{T}^{-1}(\tilde{\boldsymbol{W}}_n\boldsymbol{K}_n - \boldsymbol{I})$. Then $\det(\boldsymbol{T}^{-1})\det(\boldsymbol{I} - \tilde{\boldsymbol{W}}_n\boldsymbol{K}_n) > 0$, but since $\det(\boldsymbol{T}^{-1}) > 0$, we conclude that

$$\det(\boldsymbol{I} - \tilde{\boldsymbol{W}}_n\boldsymbol{K}_n) > 0 \tag{S79}$$

The second necessary condition is that the eigenvalues $\lambda_{n\rho}$ of matrix $\boldsymbol{\Sigma}^{-1} - \tilde{\boldsymbol{V}}_n\boldsymbol{K}_n\tilde{\boldsymbol{U}}_n$ do not lie on the negative real line, i.e. $\lambda_{n\rho} \notin (-\infty, 0]$ for all $\rho \in \mathbb{Z}_{N_c^2-1}$. This is a key condition that ensures the linear response kernel we derived in section 4 is valid for all stable networks, as the inverse of $\lambda_{n\rho} - \nabla^2$, taken in Eq. S44, exists if and only if $\lambda_{n\rho} \notin (-\infty, 0]$. The key is to observe that $\lambda_{n\rho} \notin (-\infty, 0]$ if and only if $\lambda_{n\rho} + k^2 \neq 0$ for all $k \in \mathbb{R}$. But $\lambda_{n\rho} + k^2$ are the eigenvalues of $\boldsymbol{\Sigma}^{-1} - \tilde{\boldsymbol{V}}_n\boldsymbol{K}_n\tilde{\boldsymbol{U}}_n + k^2\boldsymbol{I}$, so $\lambda_{n\rho} \notin (-\infty, 0]$ for all $\rho \in \mathbb{Z}_{N_c^2-1}$ if and only if $\det(\boldsymbol{\Sigma}^{-1} + k^2\boldsymbol{I} - \tilde{\boldsymbol{V}}_n\boldsymbol{K}_n\tilde{\boldsymbol{U}}_n) \neq 0$ for all $k \in \mathbb{R}$. Now by Sylvester's determinant theorem one has

$$\det(\boldsymbol{\Sigma}^{-1} + k^2\boldsymbol{I} - \tilde{\boldsymbol{V}}_n\boldsymbol{K}_n\tilde{\boldsymbol{U}}_n) = \det(\boldsymbol{I} - \tilde{\boldsymbol{U}}_n(\boldsymbol{\Sigma}^{-1} + k^2\boldsymbol{I})^{-1}\tilde{\boldsymbol{V}}_n\boldsymbol{K}_n)\det(\boldsymbol{\Sigma}^{-1} + k^2\boldsymbol{I}) \tag{S80}$$

Since $\det(\boldsymbol{\Sigma}^{-1}+k^2\boldsymbol{I})>0$ for all $k\in\mathbb{R}$, $\det(\boldsymbol{\Sigma}^{-1}+k^2\boldsymbol{I}-\tilde{\boldsymbol{V}}_n\boldsymbol{K}_n\tilde{\boldsymbol{U}}_n)\neq 0$ for all $k\in\mathbb{R}$ if and only if $\det(\boldsymbol{M}_n(k))\neq 0$ for all $k\in\mathbb{R}$, where we defined $\boldsymbol{M}_n(k)=\boldsymbol{I}-\tilde{\boldsymbol{U}}_n(\boldsymbol{\Sigma}^{-1}+k^2\boldsymbol{I})^{-1}\tilde{\boldsymbol{V}}_n\boldsymbol{K}_n$. But the entries of $\boldsymbol{M}_n(k)$ is given by

$$\begin{aligned}
M_{n\alpha\beta}(k)&=\delta_{\alpha\beta}-\sum_{\gamma=0}^{N_c^2-1}\sum_{\alpha',\beta'=0}^{N_c-1}A_{n\alpha\beta'}\langle N_c\alpha+\beta'\,|\,\gamma\rangle(\tilde{\sigma}_\gamma^{-2}+k^2)^{-1}\langle\gamma\,|\,N_c\alpha'+\beta\rangle\langle g_{n\beta}\,|\,f_n\rangle\\
&=\delta_{\alpha\beta}-\sum_{\alpha',\beta'=0}^{N_c-1}A_{n\alpha\beta'}(\sigma_{\alpha\beta}^{-2}+k^2)^{-1}\langle N_c\alpha+\beta'\,|\,N_c\alpha'+\beta\rangle\langle g_{n\beta}\,|\,f_n\rangle\\
&=\delta_{\alpha\beta}-A_{n\alpha\beta}(\sigma_{\alpha\beta}^{-2}+k^2)^{-1}\langle g_{n\beta}\,|\,f_n\rangle\\
&=\delta_{\alpha\beta}-\langle g_{n\beta}\,|\,f_n\rangle(1+k^2\sigma_{\alpha\beta}^2)^{-1}\tilde{W}_{n\alpha\beta}\\
&=-\tau_\alpha J_{n\alpha\beta}(k)
\end{aligned}\qquad\text{[S81]}$$

Thus $\det(\boldsymbol{M}_n(k))=\det(-\boldsymbol{J}_n(k))\prod_{\alpha=0}^{N_c-1}\tau_\alpha$. But $\tau_\alpha>0$ for all $\alpha$, and the stability constraint implies that $\det(-\boldsymbol{J}_n(k))>0$, so $\det(\boldsymbol{M}_n(k))\neq 0$ and thus we are done. Furthermore, by the eigenvalue relation proved in section 6, we see that if the connectivity widths depend only on presynaptic cell type, then the eigenvalues of $(\boldsymbol{I}-\boldsymbol{K}_n\tilde{\boldsymbol{W}}_n)\tilde{\boldsymbol{\Sigma}}^{-1}$ also do not lie on the negative real line.

**C. Simplified stability conditions for E-I networks with sufficiently fast inhibition.** In general, stability of a network depends on the synaptic time constants $\tau_\alpha$ for each cell type, as seen from Eq. S73. However, in an E-I networks with sufficiently fast inhibition time constant, the stability condition simplifies such that it no longer depends on the precise values of E and I time constants. Let the indices $\alpha=0,1$ denote the excitatory cell type and inhibitory cell type respectively (note that these are distinct from the indices $n=0,1$ which denote the zeroth and first Fourier mode on the circle). We claim that for E-I networks with $\langle g_{11}\,|\,f_{11}\rangle\kappa_{11}w_{11}\neq 1$ and $\frac{\tau_0}{\tau_1}>M$, where

$$M=\begin{cases}\max\limits_{n=0,1}\sup\limits_{k\in\mathbb{R}}\frac{(1+\sigma_{00}^2k^2)^{-1}\langle g_{n0}\,|\,f_{n0}\rangle\tilde{W}_{n00}-1}{1-(1+\sigma_{11}^2k^2)^{-1}\langle g_{n1}\,|\,f_{n1}\rangle\tilde{W}_{n11}}, & \langle g_{11}\,|\,f_{11}\rangle\kappa_{11}w_{11}<1\\ \frac{\langle g_{10}\,|\,f_{10}\rangle\tilde{W}_{100}-1}{1-\langle g_{11}\,|\,f_{11}\rangle\tilde{W}_{111}}, & \langle g_{11}\,|\,f_{11}\rangle\kappa_{11}w_{11}>1\end{cases},\qquad\text{[S82]}$$

the network is stable if and only if $\lambda_{n\rho}\notin(-\infty,0]$ and $\langle g_{11}\,|\,f_{11}\rangle\kappa_{11}w_{11}<1$. In other words, unless $\langle g_{11}\,|\,f_{11}\rangle\kappa_{11}w_{11}=1$, an E-I network with sufficiently fast time inhibitory time constant is stable if and only if its I → I connections are not strongly anti-like-to-like and $\lambda_{n\rho}\notin(-\infty,0]$.

To see this, let us first rewrite our claim in a simpler form by defining

$$\tilde{J}_{n\alpha\beta}(k)=(1+\sigma_{\alpha\beta}^2k^2)^{-1}\langle g_{n\beta}\,|\,f_{n\beta}\rangle\tilde{W}_{n\alpha\beta}-\delta_{\alpha\beta}\qquad\text{[S83]}$$

which is simply equal to $\tau_\alpha J_{n\alpha\beta}(k)$. Then our claim is that for E-I networks with $\tilde{J}_{111}(0)\neq 0$ and $\frac{\tau_0}{\tau_1}>M$, where

$$M=\begin{cases}\max\limits_{n=0,1}\sup\limits_{k\in\mathbb{R}}-\frac{\tilde{J}_{n00}(k)}{\tilde{J}_{n11}(k)}, & \tilde{J}_{111}(0)<0\\ -\frac{\tilde{J}_{100}(0)}{\tilde{J}_{111}(0)}, & \tilde{J}_{111}(0)>0\end{cases},\qquad\text{[S84]}$$

the network is stable if and only if $\lambda_{n\rho}\notin(-\infty,0]$ and $\tilde{J}_{111}(0)<0$. Next, we note that for an E-I network, the characteristic polynomial of $\boldsymbol{J}_n(k)$ is given by $p(t)=t^2-\mathrm{tr}(\boldsymbol{J}_n(k))t+\det(\boldsymbol{J}_n(k))$. By the Liénard-Chipart criterion (20, p. 221), the roots of $p$ have negative real part if and only if $\mathrm{tr}(\boldsymbol{J}_n(k))<0$ and $\det(\boldsymbol{J}_n(k))>0$. Thus, an E-I network is stable if and only if $\mathrm{tr}(\boldsymbol{J}_n(k))<0$ and $\det(\boldsymbol{J}_n(k))>0$ for all $n,k$.

Now we show that stability implies $\lambda_{n\rho}\notin(-\infty,0]$ and $\tilde{J}_{111}(0)<0$. We already proved that stability implies $\lambda_{n\rho}\notin(-\infty,0]$ in the previous subsection, so it remains to show that stability implies $\tilde{J}_{111}(0)<0$. Suppose for contradiction that $\tilde{J}_{111}(0)>0$ (it cannot be 0 by assumption). Then our time constant condition becomes $\frac{\tau_0}{\tau_1}>-\frac{\tilde{J}_{100}(0)}{\tilde{J}_{111}(0)}$, so we have $\mathrm{tr}(\boldsymbol{J}_1(0))=\tau_0^{-1}\tilde{J}_{100}(0)+\tau_1^{-1}\tilde{J}_{111}(0)>0$, which violates the stability condition of an E-I network. Thus we must have $\tilde{J}_{111}(0)<0$.

Next we show that $\lambda_{n\rho}\notin(-\infty,0]$ and $\tilde{J}_{111}(0)<0$ implies stability. We do this by showing that $\tilde{J}_{111}(0)<0$ implies $\mathrm{tr}(\boldsymbol{J}_n(k))<0$ for all $n,k$ and that $\lambda_{n\rho}\notin(-\infty,0]$ implies $\det(\boldsymbol{J}_n(k))>0$ for all $n,k$. Indeed, if $\tilde{J}_{111}(0)<0$, then $\langle g_{11}\,|\,f_{11}\rangle\tilde{W}_{111}<1$, so we must have $\tilde{J}_{111}(k)=(1+\sigma_{11}^2k^2)^{-1}\langle g_{11}\,|\,f_{11}\rangle\tilde{W}_{111}-1<0$ for all $k$. Furthermore, since $w_{11}<0$, we have $\tilde{J}_{011}(k)=(1+\sigma_{11}^2k^2)^{-1}w_{11}-1<0$ for all $k$. Thus $\tilde{J}_{n11}(k)<0$ for all $n,k$. But since $\tilde{J}_{111}(0)<0$, our time constant condition becomes $\frac{\tau_0}{\tau_1}>-\frac{\tilde{J}_{n00}(k)}{\tilde{J}_{n11}(k)}$ for all $n,k$, so we have $\mathrm{tr}(\boldsymbol{J}_n(k))=\tau_0^{-1}\tilde{J}_{n00}(k)+\tau_1^{-1}\tilde{J}_{n11}(k)<0$

for all $n, k$. Thus it remains to show that $\lambda_{n\rho} \notin (-\infty, 0]$ implies $\det(\boldsymbol{J}_n(k)) > 0$. Since we have only two cell types ($N_c = 2$), $\det(\boldsymbol{J}_n(k)) = \det(-\boldsymbol{J}_n(k))$. Furthermore, since we showed in the previous subsection that $\det(\boldsymbol{M}_n(k)) = \det(-\boldsymbol{J}_n(k)) \prod_{\alpha=0}^{N_c-1} \tau_\alpha$, it suffices to show that $\det(\boldsymbol{M}_n(k)) > 0$. Since $\boldsymbol{M}_n(k)$ is a real matrix, it suffices to show that all the real eigenvalues of $\boldsymbol{M}_n(k)$ are positive, but this is true if $\lambda_{n\rho} \notin (-\infty, 0]$, so we are done.

**D. ISN condition.** Finally, let us consider the condition for the network to be an inhibition-stabilized network (ISN). ISNs, by definition, are unstable without inhibitory neurons. Suppose there is only a single excitatory cell type $\alpha = 0$. Then the excitatory subnetwork is unstable if and only if

$$\max_{n=0,1} \sup_{k\in\mathbb{R}} \mathrm{Re}(J_{n00}(k)) = \tau_0^{-1} \max_{n=0,1} \sup_{k\in\mathbb{R}} \left(\langle g_{n0} \mid f_{n0}\rangle (1 + \sigma_{00}^2 k^2)^{-1} \tilde{W}_{n00} - 1\right) > 0 \qquad \text{[S85]}$$

Since $\sup_{k\in\mathbb{R}} (1 + \sigma_{00}^2 k^2)^{-1} = 1$, the network is an ISN if and only if

$$\max_{n=0,1} \langle g_{n0} \mid f_{n0}\rangle \tilde{W}_{n00} = \max\{w_{00}, \langle g_{10} \mid f_{10}\rangle w_{00} \kappa_{00}\} > 1 \qquad \text{[S86]}$$

But since $\langle g_{10} \mid f_{10}\rangle \leq 1$ and $\kappa_{00} \leq 1$, the network is an ISN if and only if

$$w_{00} > 1 \qquad \text{[S87]}$$

#### 8. Mean response of unperturbed neurons

In this section we consider the properties of the mean response of unperturbed neurons to a single-cell perturbation in an E-I network. By Eq. S55, the mean response is given by (here $\tilde{\boldsymbol{L}}$ is the matrix of elements $\tilde{L}_{\alpha\beta}$)

$$\tilde{\boldsymbol{L}} = \begin{bmatrix} \frac{1-w_{11}}{d} - 1 & \frac{w_{01}}{d} \\ \frac{w_{10}}{d} & \frac{1-w_{00}}{d} - 1 \end{bmatrix} = \frac{1}{d} \begin{bmatrix} w_{00} - \det(\boldsymbol{W}) & w_{01} \\ w_{10} & w_{11} - \det(\boldsymbol{W}) \end{bmatrix} \qquad \text{[S88]}$$

where $d = \det(\boldsymbol{I} - \boldsymbol{W}) = 1 - \mathrm{tr}(\boldsymbol{W}) + \det(\boldsymbol{W})$, and the indices $\alpha, \beta = 0, 1$ represent excitatory and inhibitory cell types respectively. Since stability constraint implies $d > 0$ (see Eq. S79), we see that the mean response of unperturbed excitatory neurons to single-cell excitatory neuron perturbation must be negative if and only if

$$\det(\boldsymbol{W}) > w_{00} \qquad \text{[S89]}$$

Furthermore, the mean response of inhibitory neurons to single-cell excitatory neuron perturbation must always be positive.

Now let us consider the effect of gain on the mean response of unperturbed excitatory neurons to single-cell excitatory neuron perturbation, assuming that this mean response is negative at gain $g = 1$. We will show that decreasing the gain always results in less suppression, and increasing the gain always results in more suppression. More precisely, let us we define

$$\tilde{L}_{00}(g) = \frac{g w_{00} - g^2 \det(\boldsymbol{W})}{1 - g\mathrm{tr}(\boldsymbol{W}) + g^2 \det(\boldsymbol{W})} \qquad \text{[S90]}$$

which is the mean response of unperturbed excitatory neurons as a function of gain $g$, and suppose that $\tilde{L}_{00}(1) < 0$. Then we claim that $\tilde{L}_{00}(g) > \tilde{L}_{00}(1)$ for $g \in (0, 1)$ and $\tilde{L}_{00}(g) < \tilde{L}_{00}(1)$ for $g > 1$. To show this, first we take the derivative with respect to the gain $g$ to get

$$\tilde{L}'_{00}(g) = \frac{w_{00} - g(2 - g w_{11})\det(\boldsymbol{W})}{(1 - g\mathrm{tr}(\boldsymbol{W}) + g^2 \det(\boldsymbol{W}))^2}. \qquad \text{[S91]}$$

Note that for all $g > \frac{1}{2}$, $g(2 - g w_{11}) > 1$, so $\tilde{L}'_{00}(g) < 0$. This proves that $\tilde{L}_{00}(g) < \tilde{L}_{00}(1)$ for $g > 1$. For the first part of our claim, note that $\tilde{L}'_{00}(g) = 0$ for all $g$ such that

$$w_{11} g^2 - 2g + \frac{w_{00}}{\det(\boldsymbol{W})} = 0. \qquad \text{[S92]}$$

Since the $g \geq 0$, we are only looking for real and non-negative solutions. But by Descartes' rule of signs, since $w_{11} < 0$ and $\det(\boldsymbol{W}) > w_{00} > 0$, the polynomial $w_{11} g^2 - 2g + \frac{w_{00}}{\det(\boldsymbol{W})}$ has only 1 change of sign of its coefficients, so it must have exactly 1 positive real root. In particular, since $\tilde{L}'_{00}(g) < 0$ for all $g > \frac{1}{2}$, this root $g_0$ must be in the interval $(0, 1)$, and that it must be a local maximum of $\tilde{L}_{00}(g)$. Thus within the interval $(0, 1)$, $\tilde{L}_{00}(g)$ must attain its minimum at either 0 or 1. But $\tilde{L}_{00}(g) = 0 > \tilde{L}_{00}(1)$, so $\tilde{L}_{00}(g) > \tilde{L}_{00}(1)$ for all $g \in (0, 1)$.

#### 9. Local distance dependence of perturbation response

In this section we consider the condition under which the linear response integrated over feature-tuning, $\tilde{L}_{\alpha\beta}(s)$, given by Eq. S51, is locally excitatory, i.e. $\lim_{s\to 0}\tilde{L}_{\alpha\beta}(s) > 0$. By Lemma 9.1, for all $d \geq 2 \in \mathbb{N}$, we have

$$
\begin{aligned}
\tilde{L}_{\alpha\beta}(s) &\sim G_d(s;\lambda_{00}) \sum_{\rho=0}^{N_c^2-1} [\tilde{\boldsymbol{U}}_0\boldsymbol{Q}_0]_{\alpha\rho}[\boldsymbol{Q}_0^{-1}\tilde{\boldsymbol{V}}_0]_{\rho\beta} \\
&= [\tilde{\boldsymbol{U}}_0\tilde{\boldsymbol{V}}_0]_{\alpha\beta} G_d(s;\lambda_{00}) \\
&= w_{\alpha\beta}\sigma_{\alpha\beta}^{-2} G_d(s;\lambda_{00})
\end{aligned}
$$

as $s \to 0$. The last step is due to

$$
\begin{aligned}
[\tilde{\boldsymbol{U}}_n\tilde{\boldsymbol{V}}_n]_{\alpha\beta} &= \langle\alpha|\tilde{U}_n\tilde{V}_n|\beta\rangle \\
&= \sum_{\alpha',\beta'=0}^{N_c-1} A_{n\alpha\beta'}\langle N_c\alpha+\beta' \,|\, N_c\alpha'+\beta\rangle \\
&= A_{n\alpha\beta}
\end{aligned}
$$

where $\tilde{U}_n, \tilde{V}_n$ are given by equations S34, S35 and $A_{n\alpha\beta}$ is given by S26. Plugging in the asymptotic expression for $G_d(s;\lambda_{00})$ given by Lemma 17.8, we then get

$$
\tilde{L}_{\alpha\beta}(s) \sim w_{\alpha\beta}\sigma_{\alpha\beta}^{-2} \begin{cases} \dfrac{1}{2\pi}\ln(s^{-1}), & d=2 \\ \dfrac{2^{\eta-1}}{(2\pi)^{\frac{d}{2}}}\Gamma(\eta)s^{-2\eta}, & d\geq 3 \end{cases} . \tag{S93}
$$

as $s \to 0$, where $\eta = \frac{d}{2}-1$. From this expression it is clear that $\lim_{s\to 0}\tilde{L}_{\alpha\beta}(s) > 0$ if and only if

$$
w_{\alpha\beta} > 0.
$$

Thus for $d > 1$ the local behavior is very simple - nearby neurons are excited if an excitatory neuron is perturbed, and inhibited if an inhibitory neuron is perturbed. On the other hand, for $d = 1$ we have

$$
\tilde{L}_{\alpha\beta}(0) = \sum_{\rho=0}^{N_c^2-1} \frac{1}{2\sqrt{\lambda_{0\rho}}}[\tilde{\boldsymbol{U}}_0\boldsymbol{Q}_0]_{\alpha\rho}[\boldsymbol{Q}_0^{-1}\tilde{\boldsymbol{V}}_0]_{\rho\beta} = \frac{1}{2}[\tilde{\boldsymbol{U}}_0\boldsymbol{Q}_0\boldsymbol{\Lambda}_0^{-\frac{1}{2}}\boldsymbol{Q}_0^{-1}\tilde{\boldsymbol{V}}_0]_{\alpha\beta}
$$

Thus the condition for locally excitatory response for the one-dimensional case is much more complicated than higher dimensional cases.

**Lemma 9.1.** *Given positive integer $n$ and $z_i \in \mathbb{C}$, $\lambda_i \in \mathbb{C}\setminus(-\infty,0]$ for $i \in \mathbb{Z}_n$ such that $\sum_{i=0}^{n-1} z_i \neq 0$, we have*

$$
\sum_{i=0}^{n-1} z_i G_d(s;\lambda_i) \sim G_d(s;\lambda_0)\sum_{i=0}^{n-1} z_i
$$

*as $s \to 0$ for all integers $d > 1$.*

*Proof.*

$$
\lim_{s\to 0}\frac{\sum_{i=0}^{n-1} z_i G_d(s;\lambda_i)}{G_d(s;\lambda_0)\sum_{i=0}^{n-1} z_i} = \frac{1}{\sum_{i=0}^{n-1} z_i}\left(z_0 + \sum_{i=1}^{n-1} z_i \lim_{s\to 0}\frac{G_d(s;\lambda_i)}{G_d(s;\lambda_0)}\right) = 1
$$

where we applied Lemma 17.10 to get $\lim_{s\to 0}\frac{G_d(s;\lambda_i)}{G_d(s;\lambda_0)} = 1$ for all $i \in \{1,\cdots,n-1\}$. □

#### 10. Distance dependence of perturbation response - number of zero crossings

**A. Conditions for different number of zero crossings.** Consider the linear response integrated over feature-tuning, $\tilde{L}_{\alpha\beta}(\boldsymbol{x}-\boldsymbol{y})$, given by Eq. S51. We would like to study conditions under which this function exhibits zero-crossing.

However, in this general form the analysis is intractable due to the large number of sums. Thus, we restrict our analysis to a E-I network where connectivity width depends only on presynaptic cell type, i.e. $\sigma_{\alpha\beta} = \sigma_{0\beta}$ for all $\alpha, \beta$. By integrating Eq. S59 over feature-tuning similar to what we did in section 5, we get

$$\tilde{L}_{\alpha\beta}(\boldsymbol{x}-\boldsymbol{y}) = \tilde{L}_{0\alpha\beta}(s) = \sum_{\gamma=0}^{1}[\boldsymbol{W}\boldsymbol{\Sigma}^{-1}\boldsymbol{Q}]_{\alpha\gamma}[\boldsymbol{Q}^{-1}]_{\gamma\beta}G_d(s;\lambda_\gamma) \tag{S94}$$

where $\boldsymbol{\Sigma}$ is the diagonal matrix of elements $\sigma_{0\beta}^2$, $\boldsymbol{Q}\boldsymbol{\Lambda}\boldsymbol{Q}^{-1} = (\boldsymbol{I}-\boldsymbol{W})\boldsymbol{\Sigma}^{-1}$, and $\lambda_\gamma$ are the diagonal entries of $\boldsymbol{\Lambda}$, i.e. the eigenvalues. Note that we have simplified the notation from Eq. S60 by dropping some tildes and subscripts for convenience. Let us define $\boldsymbol{M} = (\boldsymbol{I}-\boldsymbol{W})\boldsymbol{\Sigma}^{-1}$. Since $\boldsymbol{M}$ is a real matrix, its two eigenvalues $\lambda_\gamma$ must be either form a complex conjugate pair or both be real. In the rest of the supplementary text we will order the two eigenvalues such that

$$\lambda_0 = \frac{1}{2}\left(\mathrm{tr}(\boldsymbol{M}) - \sqrt{\mathrm{tr}(\boldsymbol{M})^2 - 4\det(\boldsymbol{M})}\right) \tag{S95}$$

$$\lambda_1 = \frac{1}{2}\left(\mathrm{tr}(\boldsymbol{M}) + \sqrt{\mathrm{tr}(\boldsymbol{M})^2 - 4\det(\boldsymbol{M})}\right) \tag{S96}$$

so $\lambda_0 \le \lambda_1$ in the case of real eigenvalues and $\mathrm{Im}(\lambda_0) \le 0$, $\mathrm{Im}(\lambda_1) \ge 0$ in the case of complex conjugate eigenvalues. Note that in the real case we must also have $\lambda_0, \lambda_1 > 0$ since stability of the network implies $\lambda_0, \lambda_1 \notin (-\infty, 0]$ (SI section 7B).

The number of zero crossings in the case of complex conjugate eigenvalues is straightforward: since $[\boldsymbol{Q}]_{\alpha 1}[\boldsymbol{Q}^{-1}]_{1\beta} = \overline{[\boldsymbol{Q}]_{\alpha 0}[\boldsymbol{Q}^{-1}]_{0\beta}}$ by Proposition 17.2, and since $G_d(s;\bar{\lambda}) = \overline{G_d(s;\lambda)}$ by Lemma 17.6, we have

$$\begin{aligned}\tilde{L}_{\alpha\beta}(\boldsymbol{x}-\boldsymbol{y}) &= [\boldsymbol{W}\boldsymbol{\Sigma}^{-1}\boldsymbol{Q}]_{\alpha 0}[\boldsymbol{Q}^{-1}]_{0\beta}G_d(s;\lambda_0) + \overline{[\boldsymbol{W}\boldsymbol{\Sigma}^{-1}\boldsymbol{Q}]_{\alpha 0}[\boldsymbol{Q}^{-1}]_{0\beta}G_d(s;\lambda_0)} \\ &= 2\mathrm{Re}([\boldsymbol{W}\boldsymbol{\Sigma}^{-1}\boldsymbol{Q}]_{\alpha 0}[\boldsymbol{Q}^{-1}]_{0\beta}G_d(s;\lambda_0)).\end{aligned}$$

But since $K_\eta(z) \sim \sqrt{\frac{\pi}{2z}}e^{-z}$ as $|z| \to \infty$ (13, Equation 9.7.2), $\mathrm{Re}(cK_\eta(\sqrt{\lambda_0}s))$ is an oscillatory function for all nonzero constants $c \in \mathbb{C}$, meaning it has infinitely many zero-crossings, and thus so does $\tilde{L}_{\alpha\beta}(\boldsymbol{x}-\boldsymbol{y})$. A more formal argument is presented in Theorem 17.18.

In the case of real eigenvalues, each term of $\tilde{L}_{\alpha\beta}$ is an exponentially decaying function. As a sum of two exponentially decaying functions, one would naturally expect there is either zero or one zero-crossing depending on the signs and relative amplitudes of the two terms. Assuming $\lambda_0 \neq \lambda_1$ and the coefficient of each term is non-zero (we will freely invoke these assumptions in the rest of the text since the set of model parameters which violate these assumptions has measure zero and the behavior of the linear response is trivial and uninteresting under these parameters), this is indeed true as we show in Theorem 17.19, which further shows that $\tilde{L}_{\alpha\beta}$ has exactly one zero-crossing if and only if

$$c_{\alpha\beta} := -\frac{[\boldsymbol{W}\boldsymbol{\Sigma}^{-1}\boldsymbol{Q}]_{\alpha 0}[\boldsymbol{Q}^{-1}]_{0\beta}}{[\boldsymbol{W}\boldsymbol{\Sigma}^{-1}\boldsymbol{Q}]_{\alpha 1}[\boldsymbol{Q}^{-1}]_{1\beta}} \in \begin{cases}(0,1), & d \ge 2 \\ (0, \sqrt{\frac{\lambda_0}{\lambda_1}}), & d = 1\end{cases}. \tag{S97}$$

Since $\boldsymbol{M}$ is a $2\times 2$ matrix, $\boldsymbol{Q}$ can be expressed in a rather simple form in terms of the eigenvalues $\lambda_\gamma$ of $\boldsymbol{M}$ as

$$\boldsymbol{Q} = \begin{bmatrix}\lambda_0 - M_{11} & \lambda_1 - M_{11} \\ M_{10} & M_{10}\end{bmatrix}. \tag{S98}$$

Plugging this into the expression for $c_{\alpha\beta}$ (the algebra can be slightly simplified by noting that $\boldsymbol{W}\boldsymbol{\Sigma}^{-1}\boldsymbol{Q} = \boldsymbol{\Sigma}^{-1}\boldsymbol{Q} - \boldsymbol{Q}\boldsymbol{\Lambda}$), one finds remarkably simple expressions for $c_{\alpha\beta}$, especially for $c_{01}$ and $c_{10}$, given by

$$\begin{aligned} c_{00} &= \frac{(1-\sigma_0^2\lambda_0)(1-\sigma_1^2\lambda_0 - w_{11})}{(1-\sigma_0^2\lambda_1)(1-\sigma_1^2\lambda_1 - w_{11})}, \qquad & c_{01} &= \frac{1-\sigma_0^2\lambda_0}{1-\sigma_0^2\lambda_1}, \\ c_{10} &= \frac{1-\sigma_1^2\lambda_0}{1-\sigma_1^2\lambda_1}, & c_{11} &= \frac{(1-\sigma_1^2\lambda_0)(1-\sigma_1^2\lambda_1 - w_{11})}{(1-\sigma_1^2\lambda_1)(1-\sigma_1^2\lambda_0 - w_{11})}. \end{aligned} \tag{S99}$$

The zero crossing conditions given by Eq. S97 can be further simplified. Let us define $c_{\alpha\beta}^+, c_{\alpha\beta}^-$ to be the numerator and the denominator of each expression in Eq. S99 respectively. Then we claim that

**Theorem 10.1.** *For all $d \in \mathbb{N}$ and $(\alpha,\beta) \in \{(0,1),(1,0)\}$, $\tilde{L}_{\alpha\beta}$ has exactly one zero crossing if and only if $c_{\alpha\beta}^+ < 0$.*

*Proof.* Define $f_\alpha : (0, \infty) \to \mathbb{R}$ by

$$f_\alpha(\lambda) = \begin{cases} 1 - \sigma_\alpha^2 \lambda, & d \geq 2 \\ \frac{1-\sigma_\alpha^2 \lambda}{\sqrt{\lambda}}, & d = 1 \end{cases}$$

Then $\tilde{L}_{\alpha\beta}$ has exactly one zero crossing if and only if $\frac{f_\alpha(\lambda_0)}{f_\alpha(\lambda_1)} \in (0, 1)$. Since

$$\partial_\lambda f_\alpha(\lambda) = \begin{cases} -\sigma_\alpha^2, & d \geq 2 \\ -\frac{1+\sigma_\alpha^2 \lambda}{2\lambda^{\frac{3}{2}}} < 0, & d = 1 \end{cases}$$
$$< 0,$$

$f_\alpha$ is a strictly monotonically decreasing function, and since $0 < \lambda_0 < \lambda_1$, we have $f_\alpha(\lambda_0) > f_\alpha(\lambda_1)$. Thus $\frac{f_\alpha(\lambda_0)}{f_\alpha(\lambda_1)} \in (0, 1)$ if and only if $f_\alpha(\lambda_0) < 0$. Finally, note that $f_\alpha(\lambda_0) < 0$ if and only if $c_{\alpha\beta}^+ < 0$, so we are done. □

If we consider only networks with two or more spatial dimensions, then we can simplify the zero crossing conditions for the cases $(\alpha, \beta) \in \{(0, 0), (1, 1)\}$ as well. Specifically, we claim that

**Theorem 10.2.** *For all $d \geq 2 \in \mathbb{N}$ and $\alpha, \beta \in \{0, 1\}$, $\tilde{L}_{\alpha\beta}$ has exactly one zero crossing if and only if $c_{\alpha\beta}^+ < 0$.*

*Proof.* For $d \geq 2$, $\tilde{L}_{\alpha\beta}$ has exactly one zero crossing if and only if $\frac{c_{\alpha\beta}^+}{c_{\alpha\beta}^-} \in (0, 1)$. We already proved the statement for the cases where $\alpha \neq \beta$, so it remains to consider the cases where $\alpha = \beta$. Now notice that

$$c_{00}^+ - c_{00}^- = (\lambda_1 - \lambda_0)\sigma_0^2 \sigma_1^2 \sum_{\rho=0}^{1} [\boldsymbol{W}\boldsymbol{\Sigma}^{-1}\boldsymbol{Q}]_{0\rho}[\boldsymbol{Q}^{-1}]_{\rho 0} = (\lambda_1 - \lambda_0)\sigma_1^2 w_{00}, \quad \text{[S100]}$$

$$c_{11}^+ - c_{11}^- = (\lambda_0 - \lambda_1)\sigma_1^4 \sum_{\rho=0}^{1} [\boldsymbol{W}\boldsymbol{\Sigma}^{-1}\boldsymbol{Q}]_{1\rho}[\boldsymbol{Q}^{-1}]_{\rho 1} = (\lambda_0 - \lambda_1)\sigma_1^2 w_{11}. \quad \text{[S101]}$$

Since $\lambda_1 > \lambda_0$ and $w_{00} > 0$, $w_{11} < 0$, we have $c_{\alpha\alpha}^+ > c_{\alpha\alpha}^-$. Thus $c_{\alpha\alpha} \in (0, 1)$ if and only if $c_{\alpha\alpha}^+ < 0$. □

We can simplify the zero crossing conditions for the cases $(\alpha, \beta) \in \{(0, 0), (1, 1)\}$ even further under some mild assumptions on the model parameters. Specifically, we claim that

**Theorem 10.3.** *If $M_{00} - M_{11} < 0$, then for all $d \geq 2 \in \mathbb{N}$ and $\alpha, \beta \in \{0, 1\}$, $\tilde{L}_{\alpha\beta}$ has exactly one zero crossing if and only if $1 - \sigma_\alpha^2 \lambda_0 < 0$.*

*Proof.* Note that we have already proved this statement for $\alpha \neq \beta$ since in these cases $c_{\alpha\beta}^+ = 1 - \sigma_\alpha^2 \lambda_0$. So consider the cases $\alpha = \beta$. Note that since $c_{\alpha\alpha}^+ = (1 - \sigma_\alpha^2 \lambda_0)(1 - \sigma_1^2 \lambda_\alpha - w_{11})$, it is sufficient to show that $1 - \sigma_1^2 \lambda_\alpha - w_{11} > 0$, so that $c_{\alpha\alpha}^+ < 0$ if and only if $1 - \sigma_\alpha^2 \lambda_0 < 0$. But since $M_{11} = \frac{1 - w_{11}}{\sigma_1^2}$, we have $1 - \sigma_1^2 \lambda_\alpha - w_{11} > 0$ if and only if $\lambda_\alpha < M_{11}$, i.e.

$$\frac{1}{2}\left(M_{00} + M_{11} \pm \sqrt{M_{00}^2 + M_{11}^2 + 4M_{01}M_{10} - 2M_{00}M_{11}}\right) < M_{11}$$
$$M_{00} - M_{11} < \mp\sqrt{(M_{00} - M_{11})^2 + 4M_{01}M_{10}}.$$

Regardless of the sign of the square root, the last condition is satisfied if and only if $M_{00} - M_{11} < 0$ since $M_{01}M_{10} = \sigma_0^{-2}\sigma_1^{-2} w_{01} w_{10} < 0$, so we are done. □

**Corollary 10.4.** *If the network is an ISN ($w_{00} > 1$, see SI section 7D) or its excitatory connections are wider than inhibitory connections ($\sigma_0 > \sigma_1$), then for all $d \geq 2 \in \mathbb{N}$ and $\alpha, \beta \in \{0, 1\}$, $\tilde{L}_{\alpha\beta}$ has exactly one zero crossing if and only if $1 - \sigma_\alpha^2 \lambda_0 < 0$, i.e. $\lambda_0 > \sigma_\alpha^{-2}$.*

*Proof.* Since

$$M_{00} - M_{11} = \sigma_0^{-2}(1 - w_{00}) - \sigma_1^{-2}(1 - w_{11}) = \sigma_0^{-2} - \sigma_1^{-2} - \sigma_0^{-2} w_{00} + \sigma_1^{-2} w_{11},$$

we see that if either $w_{00} > 1$ or $\sigma_0 > \sigma_1$, then $M_{00} - M_{11} < 0$, so we are done. □

Finally, since the number of zero crossings is a unitless quantity, it should be dependent only on the ratio between $\sigma_0$ and $\sigma_1$ and not their absolute magnitudes. To make this explicit, define

$$\rho = \frac{\sigma_1}{\sigma_0}, \tag{S102}$$

$$\tilde{\boldsymbol{\Sigma}} = \sigma_0^{-1}\sigma_1^{-1}\boldsymbol{\Sigma} = \begin{bmatrix} \rho^{-1} & 0 \\ 0 & \rho \end{bmatrix}, \tag{S103}$$

$$\tilde{\boldsymbol{M}} = \sigma_0\sigma_1\boldsymbol{M} = (\boldsymbol{I} - \boldsymbol{W})\tilde{\boldsymbol{\Sigma}}^{-1} \tag{S104}$$

$$\tilde{\lambda}_0 = \frac{1}{2}\left(\mathrm{tr}(\tilde{\boldsymbol{M}}) - \sqrt{\mathrm{tr}(\tilde{\boldsymbol{M}})^2 - 4\det(\tilde{\boldsymbol{M}})}\right) \tag{S105}$$

$$\tilde{\lambda}_1 = \frac{1}{2}\left(\mathrm{tr}(\tilde{\boldsymbol{M}}) + \sqrt{\mathrm{tr}(\tilde{\boldsymbol{M}})^2 - 4\det(\tilde{\boldsymbol{M}})}\right), \tag{S106}$$

so $\tilde{\lambda}_0, \tilde{\lambda}_1$ are the eigenvalues of $\tilde{\boldsymbol{M}}$ and $\lambda_\gamma = \sigma_0^{-1}\sigma_1^{-1}\tilde{\lambda}_\gamma$. Using this notation, the zero crossing conditions for ISNs in $d \geq 2$ in the case of real eigenvalues are precisely $\tilde{\lambda}_0 > \rho$ for $\alpha = 0$ and $\tilde{\lambda}_0 > \rho^{-1}$ for $\alpha = 1$.

**B. Phase diagram of zero crossings.** We have found very simple conditions for the number of zero crossings of $\tilde{L}_{\alpha\beta}$ in terms of the eigenvalue $\tilde{\lambda}_0$ for ISNs in $d \geq 2$: $\tilde{L}_{\alpha\beta}$ has infinitely many zero crossings if and only if $\tilde{\lambda}_0$ has non-zero imaginary part, and $\tilde{L}_{\alpha\beta}$ has exactly one zero crossing if and only if $\tilde{\lambda}_0 > \rho^{\pm 1}$ (the choice of $\pm$ sign depends on whether $\alpha = 0$ or $\alpha = 1$). We now want to define two real variables such that these simple conditions can be illustrated by a phase diagram of these two variables. This phase diagram will be characterized by the phase boundaries $\mathrm{Im}(\tilde{\lambda}_0) = 0$ and $\tilde{\lambda}_0 - \rho^{\pm 1} = 0$, which, in terms of the trace and determinant of $\tilde{\boldsymbol{M}}$, are respectively given by

$$\mathrm{tr}(\tilde{\boldsymbol{M}})^2 - 4\det(\tilde{\boldsymbol{M}}) = 0, \tag{S107}$$

$$\frac{1}{2}\left(\mathrm{tr}(\tilde{\boldsymbol{M}}) - \sqrt{\mathrm{tr}(\tilde{\boldsymbol{M}})^2 - 4\det(\tilde{\boldsymbol{M}})}\right) - \rho^{\pm 1} = 0. \tag{S108}$$

The latter phase boundary can be simplified as

$$\begin{aligned}
\mathrm{tr}(\tilde{\boldsymbol{M}}) - 2\rho^{\pm 1} &= \sqrt{\mathrm{tr}(\tilde{\boldsymbol{M}})^2 - 4\det(\tilde{\boldsymbol{M}})} \\
\mathrm{tr}(\tilde{\boldsymbol{M}})^2 - 4\mathrm{tr}(\tilde{\boldsymbol{M}})\rho^{\pm 1} + 4\rho^{\pm 2} &= \mathrm{tr}(\tilde{\boldsymbol{M}})^2 - 4\det(\tilde{\boldsymbol{M}}) \\
\rho^{\pm 1}(\mathrm{tr}(\tilde{\boldsymbol{M}}) - \rho^{\pm 1}) - \det(\tilde{\boldsymbol{M}}) &= 0.
\end{aligned} \tag{S109}$$

Note that in the first line the left hand side must be non-negative, so the latter phase boundary is valid only if $\mathrm{tr}(\tilde{\boldsymbol{M}}) - 2\rho^{\pm 1} \geq 0$. Thus it is natural to define one of the variables as

$$x_\pm = 2\rho^{\pm 1} - \mathrm{tr}(\tilde{\boldsymbol{M}}) \tag{S110}$$

so we have the condition $x_\pm \leq 0$ for the latter phase boundary. Substituting this definition of $x_\pm$ into Eq. S107 and Eq. S109 respectively, we get

$$\begin{aligned}
\rho^{\pm 2} - \rho^{\pm 1}x_\pm - \det(\tilde{\boldsymbol{M}}) + \frac{x_\pm^2}{4} &= 0, \\
\rho^{\pm 2} - \rho^{\pm 1}x_\pm - \det(\tilde{\boldsymbol{M}}) &= 0.
\end{aligned}$$

Thus, it is natural to define our second variable as

$$\begin{aligned}
y_\pm &= \det(\tilde{\boldsymbol{M}}) + \rho^{\pm 1}x_\pm - \rho^{\pm 2} \\
&= \det(\tilde{\boldsymbol{M}}) - \rho^{\pm 1}\mathrm{tr}(\tilde{\boldsymbol{M}}) + \rho^{\pm 2}
\end{aligned} \tag{S111}$$

so that our two phase boundaries are given respectively by

$$y_\pm = \frac{x_\pm^2}{4}, \tag{S112}$$

$$y_\pm = 0. \tag{S113}$$

In terms of the model parameters $w_{\alpha\beta}$ and $\rho$, our two variables $x_\pm$, $y_\pm$ can be written as

$$\begin{aligned} x_\pm &= 2\rho^{\pm 1} - \rho(1 - w_{00}) - \rho^{-1}(1 - w_{11}) \\ &= \rho w_{00} - \rho^{-1}|w_{11}| \pm (\rho - \rho^{-1}) \end{aligned} \quad \text{[S114]}$$

$$\begin{aligned} y_+ &= 1 - w_{00} - w_{11} + \det(\boldsymbol{W}) - \rho^2(1 - w_{00}) - (1 - w_{11}) + \rho^2 \\ &= \det(\boldsymbol{W}) + (\rho^2 - 1)w_{00} \end{aligned} \quad \text{[S115]}$$

$$\begin{aligned} y_- &= 1 - w_{00} - w_{11} + \det(\boldsymbol{W}) - (1 - w_{00}) - \rho^{-2}(1 - w_{11}) + \rho^{-2} \\ &= \det(\boldsymbol{W}) + (1 - \rho^{-2})|w_{11}| \end{aligned} \quad \text{[S116]}$$

As we have shown in SI section 7B, stability implies that the eigenvalues of $\tilde{\boldsymbol{M}}$ cannot be real and negative, i.e. either the eigenvalues are complex ($y_\pm > \frac{x_\pm^2}{4}$) or they are both positive. The latter is satisfied if and only if $\tilde{\lambda}_0 > 0$, which in turn is satisfied if and only if $\mathrm{tr}(\tilde{\boldsymbol{M}}) \geq 0$ and $\det(\tilde{\boldsymbol{M}}) > 0$, i.e. $x_\pm \leq 2\rho^{\pm 1}$ and $y_\pm > \rho^{\pm 1}x_\pm - \rho^{\pm 2}$. Thus in summary, our phase diagram is characterized by the following boundaries

$$y_\pm = \begin{cases} 0, x_\pm \leq 0 & \text{boundary between 0 and 1 crossings} \quad \text{[S117]} \\ \dfrac{x_\pm^2}{4}, x_\pm \leq 0 & \text{boundary between 1 and } \infty \text{ crossings} \quad \text{[S118]} \\ \dfrac{x_\pm^2}{4}, 0 < x_\pm \leq 2\rho^{\pm 1} & \text{boundary between 0 and } \infty \text{ crossings} \quad \text{[S119]} \\ \rho^{\pm 1}x_\pm - \rho^{\pm 2}, x_\pm \leq 2\rho^{\pm 1} & \text{boundary between 0 crossings and instability} \quad \text{[S120]} \\ \dfrac{x_\pm^2}{4}, x_\pm > 2\rho^{\pm 1} & \text{boundary between } \infty \text{ crossings and instability} \quad \text{[S121]} \end{cases}$$

**C. Changes in number of zero crossings due to perturbations of model parameters.** Now let us understand how the different model parameters affect the qualitative features of the linear response by studying the changes in number of zero crossings due to an infinitesimal perturbation of each parameter along the each of the phase boundaries given by equations S117, S118, S119. More concretely, for each model parameter $\theta$, we shall compute the derivatives $\partial_\theta y_\pm$ and $\partial_\theta z$, where

$$z := 4y_\pm - x_\pm^2 = 4\det(\tilde{\boldsymbol{M}}) - \mathrm{tr}(\tilde{\boldsymbol{M}})^2. \quad \text{[S122]}$$

If $\partial_\theta y_\pm > 0$ along boundary S117, then an increase of $\theta$ along this boundary results in the formation of a zero crossing. If $\partial_\theta z > 0$ along boundaries S119 and S118, then an increase of $\theta$ along these boundaries results in an increase of number of zero crossings from 0 or 1 to $\infty$. As in the previous subsection, we shall only consider ISNs with $d \geq 2$.

**Perturbing E-I/I-E connection strength** Since

$$\begin{aligned} \partial_{w_{10}} x_\pm &= 0, & \partial_{|w_{01}|} x_\pm &= 0, \\ \partial_{w_{10}} y_\pm &= |w_{01}|, & \partial_{|w_{01}|} y_\pm &= w_{10}, \end{aligned}$$

we see that for $\theta \in \{w_{10}, |w_{01}|\}$, we have $\partial_\theta y_\pm > 0$ and $\partial_\theta z = 4\partial_\theta y_\pm > 0$ everywhere, i.e. increasing E $\to$ I or I $\to$ E connection strength at the phase boundaries always result an increase in number of zero-crossings.

**Perturbing $\rho$** Since

$$\partial_\rho y_+ = 2\rho w_{00}, \qquad \partial_\rho y_- = 2\rho^{-3}|w_{11}|,$$

we have $\partial_\rho y_\pm > 0$ everywhere. On the other hand, since $\det(\tilde{\boldsymbol{M}})$ is independent of $\rho$, we have

$$\partial_\rho z = -\partial_\rho \mathrm{tr}(\tilde{\boldsymbol{M}})^2 = -2\mathrm{tr}(\tilde{\boldsymbol{M}})\partial_\rho \mathrm{tr}(\tilde{\boldsymbol{M}}).$$

Since boundaries S118, S119 satisfies $x_\pm < 2\rho^{\pm 1}$, we have $\mathrm{tr}(\tilde{\boldsymbol{M}}) = 2\rho^{\pm 1} - x_\pm > 0$. On the other hand, since $\mathrm{tr}(\tilde{\boldsymbol{M}}) = \rho(1 - w_{00}) + \rho^{-1}(1 + |w_{11}|)$, we have

$$\partial_\rho \mathrm{tr}(\tilde{\boldsymbol{M}}) = 1 - w_{00} - \rho^{-2}(1 + |w_{11}|)$$

which is negative for ISNs. Thus $\partial_\rho z > 0$ along the boundaries S118, S119, so increasing $\rho$ at the phase boundaries always results in an increase in number of zero-crossings.

**Perturbing E-E/I-I connection strength** We have the derivatives

$$\partial_{w_{00}} y_+ = \rho^2 - 1 - |w_{11}|, \qquad \partial_{|w_{11}|} y_+ = -w_{00},$$
$$\partial_{w_{00}} y_- = -|w_{11}|, \qquad \partial_{|w_{11}|} y_- = 1 - \rho^{-2} - w_{00}.$$

Since ISNs satisfy $w_{00} > 1$, we have $\partial_{|w_{11}|} y_- < 0$. Furthermore, along the boundary S117, we have $x_+ \leq 0$, which results in the inequality

$$\rho^2 - 1 - |w_{11}| \leq -\rho^2 w_{00}$$

so $\partial_{w_{00}} y_+ < 0$. Thus for $\theta \in \{w_{00}, |w_{11}|\}$, we have $\partial_\theta y_\pm < 0$ along the boundary S117.

On the other hand, since

$$\partial_{w_{00}} \mathrm{tr}(\tilde{\boldsymbol{M}}) = -\rho, \qquad \partial_{|w_{11}|} \mathrm{tr}(\tilde{\boldsymbol{M}}) = \rho^{-1},$$
$$\partial_{w_{00}} \det(\tilde{\boldsymbol{M}}) = -1 - |w_{11}|, \qquad \partial_{|w_{11}|} \det(\tilde{\boldsymbol{M}}) = 1 - w_{00},$$

we have

$$\begin{aligned}
\partial_{w_{00}} z &= -4(1 + |w_{11}|) + 2\mathrm{tr}(\tilde{\boldsymbol{M}})\rho \\
&= -4(1 + |w_{11}|) + 2(\rho^2(1 - w_{00}) + 1 + |w_{11}|) \\
&= -2(1 + |w_{11}|) - 2\rho^2(w_{00} - 1), \\
\partial_{|w_{11}|} z &= -4(w_{00} - 1) - 2\mathrm{tr}(\tilde{\boldsymbol{M}})\rho^{-1} \\
&= -4(w_{00} - 1) - 2(1 - w_{00} + \rho^{-2}(1 + |w_{11}|)) \\
&= -2(w_{00} - 1) - 2\rho^{-2}(1 + |w_{11}|).
\end{aligned}$$

Since $w_{00} - 1 > 0$ for ISNs, for $\theta \in \{w_{00}, |w_{11}|\}$ we have $\partial_\theta z < 0$ everywhere. Thus increasing E $\to$ E or I $\to$ I connection strength at the phase boundaries always results in a decrease in number of zero-crossings.

**D. Changes in number of zero crossings due to perturbation of neuronal gain.** Let us also study the effect of baseline activity firing rate on the linearized perturbation response in a nonlinear network. Increasing baseline activity changes the gain of neurons $g$, which in turn changes the linearized connectivity weights $g\boldsymbol{W}$ and thus changes the number of zero crossings in the linearized perturbation response as a function of space. Since the connectivity weights are now scaled by $g$, the two axes of our phase diagram, $x_\pm, y_\pm$ (equations S114-S116), are now functions of $g$. Specifically,

$$\begin{aligned}
x_\pm(g) &= g(\rho w_{00} - \rho^{-1}|w_{11}|) \pm (\rho - \rho^{-1}) \\
y_+(g) &= g^2 \det(\boldsymbol{W}) + g(\rho^2 - 1)w_{00} \\
y_-(g) &= g^2 \det(\boldsymbol{W}) + g(1 - \rho^{-2})|w_{11}|,
\end{aligned}$$

so we have

$$\begin{aligned}
g\partial_g x_\pm &= g(\rho w_{00} - \rho^{-1}|w_{11}|) = x_\pm \mp (\rho - \rho^{-1}) \\
g\partial_g y_+ &= 2g^2 \det(\boldsymbol{W}) + g(\rho^2 - 1)w_{00} = 2y_+ - g(\rho^2 - 1)w_{00} \\
g\partial_g y_- &= 2g^2 \det(\boldsymbol{W}) + g(1 - \rho^{-2})|w_{11}| = 2y_- - g(1 - \rho^{-2})|w_{11}|.
\end{aligned}$$

Along the boundary S117, $y_\pm = 0$, so $\partial_g y_\pm = 0$ if $\rho = 1$, $\partial_g y_\pm < 0$ if $\rho > 1$, and $\partial_g y_\pm > 0$ otherwise. On the other hand, the variable $z$ defined by equation S122 is now also a function of $g$ and is given by

$$z(g) = 4y_+(g) - x_+^2(g).$$

Thus we have

$$\begin{aligned}
g\partial_g z &= 4g\partial_g y_+ - 2x_+ g\partial_g x_+ \\
&= 8y_+ - 4g(\rho^2 - 1)w_{00} - 2x_+(x_+ - (\rho - \rho^{-1})),
\end{aligned}$$

so along the boundaries S118, S119 where $y_+ = \frac{x_+^2}{4}$, we have

$$\begin{aligned}
g\partial_g z &= -4g(\rho^2 - 1)w_{00} + 2x_+(\rho - \rho^{-1}) \\
&= 2(\rho - \rho^{-1})(x_+ - 2\rho g w_{00}).
\end{aligned}$$

Since $x_+ < 2\rho$ along the two boundaries and $gw_{00} > 1$ for an ISN, we have $x_+ - 2\rho g w_{00} < 0$. Thus $\partial_g z = 0$ if $\rho = 1$, $\partial_g z < 0$ if $\rho > 1$ and $\partial_g z > 0$ otherwise.

In summary, if $\rho = 1$, there is no change in number of zero crossings with an increase in gain. If $\rho > 1$, the number of zero crossing decreases, whereas if $\rho < 1$, the number of zero crossing increases.

**E. Relationship between the number of zero crossings of inhibitory and excitatory neurons.** As we showed earlier, in E-I ISNs with two or more spatial dimensions, there are infinitely many zero crossings if and only if the eigenvalues of $(\boldsymbol{I} - \boldsymbol{W})\boldsymbol{\Sigma}^{-1}$ are complex conjugates. This is true regardless of cell types, so that inhibitory neurons exhibit infinitely many zero crossings if and only if excitatory neurons exhibit infinitely many zero crossings. However, the condition for exactly one zero crossing is different for excitatory and inhibitory neurons. Here we characterize the condition under which excitatory neurons exhibit exactly one zero crossing but inhibitory neurons do not.

In the following we will always be assuming single-cell excitatory neuron perturbations. By Corollary 10.4, excitatory neurons exhibit one zero crossing while inhibitory neurons exhibit no zero crossing if and only if $\lambda_0 > \sigma_{\mathrm{E}}^{-2}$ and $\lambda_0 < \sigma_{\mathrm{I}}^{-2}$, i.e. $\sigma_{\mathrm{E}}^{-2} < \lambda_0 < \sigma_{\mathrm{I}}^{-2}$. Note that this is only possible if $\sigma_{\mathrm{I}} < \sigma_{\mathrm{E}}$, i.e. $\rho < 1$. Now we want to express this in terms of our phase diagram variables, $x_+, y_+$. From our phase diagram analysis, we found that excitatory neurons exhibit exactly one zero crossing if and only if $x_+ < 0$ and $0 < y_+ < \frac{x_+^2}{4}$. Similarly, the same condition for inhibitory neurons is that $x_- < 0$ and $0 < y_- < \frac{x_-^2}{4}$. Given the definitions of $x_\pm$ and $y_\pm$ (equations S114, S115, S116), we have

$$\begin{aligned} x_+ - x_- &= 2(\rho - \rho^{-1}), \\ y_+ - y_- &= (\rho^2 - 1)w_{00} - (1 - \rho^{-2})|w_{11}|. \end{aligned}$$

This implies

$$\begin{aligned} x_- < 0 &\iff x_+ < 2(\rho - \rho^{-1}), \\ y_- > 0 &\iff y_+ > (\rho^2 - 1)w_{00} - (1 - \rho^{-2})|w_{11}|. \end{aligned}$$

But by rearranging the definition of $x_+$, we have

$$|w_{11}| = \rho(\rho w_{00} + (\rho - \rho^{-1}) - x_+).$$

Thus

$$\begin{aligned} y_- > 0 &\iff y_+ > (\rho^2 - 1)w_{00} - (\rho - \rho^{-1})(\rho w_{00} + (\rho - \rho^{-1}) - x_+) \\ &\iff y_+ > (\rho - \rho^{-1})(x_+ - (\rho - \rho^{-1})). \end{aligned}$$

Furthermore, since

$$y_- < \frac{x_-^2}{4} \iff y_+ < \frac{x_+^2}{4},$$

inhibitory neurons exhibit exactly one zero crossing if and only if

$$\begin{cases} x_+ < 2(\rho - \rho^{-1}) \\ (\rho - \rho^{-1})(x_+ - (\rho - \rho^{-1})) < y_+ < \frac{x_+^2}{4} \end{cases}. \quad \text{[S123]}$$

This yields the phase boundaries illustrated in Figure 3 and the left column of Figure S4.

#### 11. Distance dependence of perturbation response - location of zero crossings

**A. Analytical solution of zero crossing locations in 1/3D.** In general there is no simple expression for the location of zero crossings of the linear response integrated over feature-tuning, even in the case of E-I networks with connectivity width depending only on presynaptic cell type, as studied in the previous section. However, for $d = 1, 3$ we can write down a simple expression due to the fact that $K_{\pm\frac{1}{2}}(z)$ is expressible in terms of the exponential function. Using the notation from the previous section, the zero-crossings of $\tilde{L}_{\alpha\beta}(s)$ are given by solutions of

$$\frac{G_d(s; \lambda_1)}{G_d(s; \lambda_0)} = c_{\alpha\beta}. \quad \text{[S124]}$$

For $d = 1, 3$ we can plug in equations S163 and S164 and get

$$e^{-(\sqrt{\lambda_1}-\sqrt{\lambda_0})s} = z_{d\alpha\beta}$$

$$s = \frac{\log z_{d\alpha\beta}}{\sqrt{\lambda_0}-\sqrt{\lambda_1}} \qquad \text{[S125]}$$

where

$$z_{d\alpha\beta} = c_{\alpha\beta}\begin{cases} \frac{\sqrt{\lambda_1}}{\sqrt{\lambda_0}}, & d=1 \\ 1, & d=3 \end{cases}. \qquad \text{[S126]}$$

Equation S125 is the exact expression for the location of the zero crossing for the case of real eigenvalues $\lambda_0, \lambda_1$. However, for complex conjugate $\lambda_0, \lambda_1$, the logarithm here is multi-valued, and thus further simplification is required. In particular, let us derive the exact expression for the location of the first zero crossing $s_0$ for the case of complex conjugate $\lambda_0, \lambda_1$. Let

$$z^+_{d\alpha\beta} = c^+_{\alpha\beta}\begin{cases} \sqrt{\lambda_1}, & d=1 \\ 1, & d=3 \end{cases} \qquad \text{[S127]}$$

where $c^+_{\alpha\beta}$ is as defined in the previous section, so that $z_{d\alpha\beta} = \frac{z^+_{d\alpha\beta}}{\overline{z}^+_{d\alpha\beta}}$. Then since $\log z \in \{\ln|z| + i(\mathrm{Arg}(z) + 2\pi m)\}_{m\in\mathbb{Z}}$ where $\mathrm{Arg}(z) \in (-\pi, \pi]$, we have

$$\log z_{d\alpha\beta} \in \left\{\ln 1 + \left(\mathrm{Arg}\left(\frac{z^+_{d\alpha\beta}}{\overline{z}^+_{d\alpha\beta}}\right) + 2\pi m\right) i\right\}_{m\in\mathbb{Z}} = \left\{2(\mathrm{Arg}(z^+_{d\alpha\beta}) + \pi m) i\right\}_{m\in\mathbb{Z}}.$$

Furthermore, since

$$\sqrt{\lambda_0} - \sqrt{\lambda_1} = \sqrt{\lambda_0} - \overline{\sqrt{\lambda_0}} = 2\mathrm{Im}(\sqrt{\lambda_0})i,$$

we have

$$s \in \left\{\frac{\mathrm{Arg}(z^+_{d\alpha\beta}) + \pi m}{\mathrm{Im}(\sqrt{\lambda_0})}\right\}_{m\in\mathbb{Z}}. \qquad \text{[S128]}$$

By definition of $\lambda_0$ (Eq. S95), $\mathrm{Im}(\sqrt{\lambda_0}) < 0$. Thus the first zero crossing must correspond to $m = -1$ if $\mathrm{Arg}(z^+_{d\alpha\beta}) \in (0, \pi]$ and $m = 0$ if $\mathrm{Arg}(z^+_{d\alpha\beta}) \in (-\pi, 0]$. We now claim that

**Lemma 11.1.** *If $d \geq 2$ or $\alpha \neq \beta$, then* $\mathrm{Arg}(z^+_{d\alpha\beta}) \in (0, \pi]$.

*Proof.* First, note that it suffices to show $\mathrm{Im}(z^+_{d\alpha\beta}) > 0$, since $\mathrm{Im}(z) > 0$ if and only if $\mathrm{Arg}(z) \in (0, \pi)$.

**Case $d \geq 2$, $\alpha \neq \beta$:** In this case, $z^+_{d\alpha\beta} = 1 - \sigma_\alpha^2\lambda_0$, so $\mathrm{Im}(z^+_{d\alpha\beta}) > 0$.

**Case $d = 1$, $\alpha \neq \beta$:** In this case, $z^+_{d\alpha\beta} = (1 - \sigma_\alpha^2\lambda_0)\sqrt{\lambda_0} = \sqrt{\lambda_0} - \sigma_\alpha^2|\lambda_0|\sqrt{\lambda_0}$, so $\mathrm{Im}(z^+_{d\alpha\beta}) > 0$ as well.

**Case $d \geq 2$, $\alpha = \beta$:** Using the algebraic relations given by equations S100, S101, we have for $d \geq 2$,

$$2\mathrm{Im}(z^+_{d00})i = c^+_{00} - \overline{c}^+_{00} = c^+_{00} - c^-_{00} = (\lambda_1 - \lambda_0)\sigma_1^2 w_{00} = 2\mathrm{Im}(\lambda_1)i\sigma_1^2 w_{00}$$

$$2\mathrm{Im}(z^+_{d11})i = c^+_{11} - \overline{c}^+_{11} = c^+_{11} - c^-_{11} = (\lambda_0 - \lambda_1)\sigma_1^2 w_{11} = 2\mathrm{Im}(\lambda_0)i\sigma_1^2 w_{11}$$

and since $\mathrm{Im}(\lambda_0) < 0$, $\mathrm{Im}(\lambda_1) > 0$, $w_{00} > 0$, $w_{11} < 0$, we have $\mathrm{Im}(z^+_{d00}), \mathrm{Im}(z^+_{d11}) > 0$ for $d \geq 2$. □

Thus for the case of complex conjugate $\lambda_0, \lambda_1$, if $d = 3$ or $\alpha \neq \beta$ then the first zero crossing is given by

$$s_0 = \frac{\mathrm{Arg}(z^+_{d\alpha\beta}) - \pi}{\mathrm{Im}(\sqrt{\lambda_0})}. \qquad \text{[S129]}$$

From this we may also immediately deduce the equation for the $(n+1)$-th zero-crossing:

$$s_n = \frac{\mathrm{Arg}(z^+_{d\alpha\beta}) - (n+1)\pi}{\mathrm{Im}(\sqrt{\lambda_0})}. \qquad \text{[S130]}$$

**B. Frequency of spatial oscillation in 1/3D.** From Eq. S128, we see that in 1/3D, for the case of complex conjugate eigenvalues $\lambda_0, \lambda_1$, the distance between successive zero crossings can be found as

$$\Delta s := s_1 - s_0 = \frac{\pi}{\mathrm{Im}(\sqrt{\lambda_1})}. \tag{S131}$$

$\Delta s$ is inversely proportional to the frequency of spatial oscillations in the perturbation response. We will also consider the unitless version of the same quantity given by

$$\Delta\tilde{s} := \frac{\Delta s}{\sqrt{\sigma_0\sigma_1}} = \frac{\pi}{\mathrm{Im}(\sqrt{\tilde{\lambda}_1})}. \tag{S132}$$

where $\tilde{\lambda}_1$ is the unitless version of $\lambda_1$ as defined by Eq. S106. Note that since $\Delta\tilde{s}$ depends only on $\tilde{\lambda}_1$, it is completely characterized by the two axes of the zero crossing phase diagram $(x_\pm, y_\pm)$ and $\rho$. Interestingly, $\Delta\tilde{s}$ is intimately tied to the strength of I $\to$ I connections (Figure 2F in main text). Their relationship can be analyzed mathematically by computing the derivative of $\Delta\tilde{s}$ with respect to $|w_{11}|$: if it is positive, then increasing I $\to$ I connection strength increases $\Delta\tilde{s}$, i.e. it reduces the spatial oscillation frequency. To do this, let $a = \frac{1}{2}\mathrm{tr}(\tilde{\boldsymbol{M}})$ and $b = \frac{1}{2}\sqrt{4\det(\tilde{\boldsymbol{M}}) - \mathrm{tr}(\tilde{\boldsymbol{M}})^2}$, such that $\tilde{\lambda}_1 = a + bi$. Let $\theta = \mathrm{Arg}(\tilde{\lambda}_1)$, so we may also write $\tilde{\lambda}_1 = |\tilde{\lambda}_1|e^{i\theta}$. Then

$$\mathrm{Im}\left(\sqrt{\tilde{\lambda}_1}\right) = \mathrm{Im}(\sqrt{|\tilde{\lambda}_1|}e^{i\frac{\theta}{2}}) = \sqrt{|\tilde{\lambda}_1|}\sin\left(\frac{\theta}{2}\right) = \sqrt{\frac{|\tilde{\lambda}_1|(1-\cos\theta)}{2}} = \sqrt{\frac{\sqrt{a^2+b^2}-a}{2}} = \frac{1}{2}\sqrt{2\sqrt{\det(\tilde{\boldsymbol{M}})} - \mathrm{tr}(\tilde{\boldsymbol{M}})}$$

But now since

$$\partial_{|w_{11}|}\det(\tilde{\boldsymbol{M}}) = 1 - w_{00}$$
$$\partial_{|w_{11}|}\mathrm{tr}(\tilde{\boldsymbol{M}}) = \rho^{-1}$$

we get

$$\partial_{|w_{11}|}\Delta\tilde{s} = \partial_{|w_{11}|}\frac{2\pi}{\sqrt{2\sqrt{\det(\tilde{\boldsymbol{M}})} - \mathrm{tr}(\tilde{\boldsymbol{M}})}} = \pi\left(2\sqrt{\det(\tilde{\boldsymbol{M}})} - \mathrm{tr}(\tilde{\boldsymbol{M}})\right)^{-\frac{3}{2}}\left(\rho^{-1} - \frac{1-w_{00}}{\sqrt{\det(\tilde{\boldsymbol{M}})}}\right)$$

Thus, for ISNs ($w_{00} > 1$), the derivative $\partial_{|w_{11}|}\Delta\tilde{s}$ is positive everywhere, and thus increasing I $\to$ I connection strength always reduces the spatial oscillation frequency.

**C. Modulation of zero crossings by neuronal gain.** Now let us study how the baseline firing rate in a nonlinear network affects the location of zero-crossing of the linearized perturbation response in space. As discussed in SI section 10D, a change in baseline firing rate changes the gain of neurons $g$, which in turn changes the linearized connectivity matrix $g\boldsymbol{W}$. Let us consider the special case of $\rho = 1$. We claim that in this case, increasing the gain $g$ always reduces the spatial radius of nearby excitation/inhibition, regardless of the perturbed cell type or the measured cell type.

To see this, first note that when $\rho = 1$, the eigenvalues can be written as a function of gain as $\lambda_\alpha(g) = \sigma^{-2}(1 - g\lambda_\alpha^W)$, where $\sigma = \sigma_0 = \sigma_1$, and $\lambda_\alpha^W$ are the corresponding eigenvalues of $\boldsymbol{W}$. Furthermore, let us define $f_{d\alpha\beta} : \mathbb{C}\times\mathbb{C} \to \mathbb{C}$ by

$$f_{d\alpha\beta}(g, s) = \frac{G_d(s; \lambda_1(g))}{G_d(s; \lambda_0(g))} - c_{\alpha\beta}$$

where $c_{\alpha\beta}$ is given by

$$c_{00} = \frac{\lambda_0^W(\lambda_0^W - w_{11})}{\lambda_1^W(\lambda_1^W - w_{11})}, \qquad c_{01} = c_{10} = \frac{\lambda_0^W}{\lambda_1^W}, \qquad c_{11} = \frac{\lambda_0^W(\lambda_1^W - w_{11})}{\lambda_1^W(\lambda_0^W - w_{11})}.$$

These expressions for $c_{\alpha\beta}$ is obtained by replacing $\lambda_\alpha$ and $w_{11}$ in Eq. S99 with $\lambda_\alpha(g)$ and $gw_{11}$ respectively, and then simplifying by plugging in $\rho = 1$. Given a fixed $\boldsymbol{W}$ and gain $g_0 > 0$, suppose there exists a zero crossing $s_0 > 0$, i.e. $f_{d\alpha\beta}(g_0, s_0) = 0$. Since $K_\eta(z)$ is holomorphic on $\mathbb{C}\setminus(-\infty, 0]$ and has no zeros in the right-half plane $\mathrm{Re}(z) > 0$ (13, section 9.6), there exists open neighborhoods $U, V \in \mathbb{C}$ containing $g_0$, $s_0$ such that $f_{d\alpha\beta}$ is holomorphic on $U \times V$. Furthermore, as part of our derivation later, it can be shown that $\partial_s f_{d\alpha\beta}(g, s) \neq 0$ for all $g, s > 0$. Thus, by the holomorphic implicit function theorem (21, chapter I, theorem 7.6) there exists a neighborhood $\tilde{U} \subset U$ of

$g_0$ and a holomorphic function $h : \tilde{U} \to \mathbb{C}$ such that $h(g_0) = s_0$ and $f_{d\alpha\beta}(g, h(g)) = 0$ for all $g \in \tilde{U}$. This implies that $h$ is a function that maps the gain $g$ to the corresponding location of the first zero crossing, $h(g)$. Thus, if $h'(g_0) < 0$, then increasing gain reduces the distance to the first zero crossing, i.e. reduces the spatial radius of nearby excitation/inhibition. The implicit function theorem yields the following equation for $h'(g)$

$$h'(g) = -\frac{\partial_g f_{d\alpha\beta}(g, h(g))}{\partial_s f_{d\alpha\beta}(g, h(g))}$$

By Lemma 17.15

$$\partial_s f_{d\alpha\beta}(g, s) = -2\pi s \frac{G_d(s; \lambda_0(g))G_{d+2}(s; \lambda_1(g)) - G_d(s; \lambda_1(g))G_{d+2}(s; \lambda_0(g))}{G_d(s; \lambda_0(g))^2},$$

and by Lemma 17.14,

$$\partial_g f_{d\alpha\beta}(g, s) = -\frac{1}{4\pi} \frac{\lambda_1'(g)G_d(s; \lambda_0(g))G_{d-2}(s; \lambda_1(g)) - \lambda_0'(g)G_d(s; \lambda_1(g))G_{d-2}(s; \lambda_0(g))}{G_d(s; \lambda_0(g))^2}.$$

Since $g\lambda_\alpha'(g) = -\sigma^{-2}g\lambda_\alpha^W = \lambda_\alpha(g) - \sigma^{-2}$, we thus have

$$\frac{\partial_g f_{d\alpha\beta}(g, s)}{\partial_s f_{d\alpha\beta}(g, s)} = \frac{1}{8\pi^2 gs} \frac{(\lambda_1(g) - \sigma^{-2})G_d(s; \lambda_0(g))G_{d-2}(s; \lambda_1(g)) - (\lambda_0(g) - \sigma^{-2})G_d(s; \lambda_1(g))G_{d-2}(s; \lambda_0(g))}{G_d(s; \lambda_0(g))G_{d+2}(s; \lambda_1(g)) - G_d(s; \lambda_1(g))G_{d+2}(s; \lambda_0(g))}.$$

Applying Lemma 17.17, this simplifies as

$$\frac{\partial_g f_{d\alpha\beta}(g, s)}{\partial_s f_{d\alpha\beta}(g, s)} = \frac{1}{8\pi^2 gs} \left( 4\pi^2 s^2 - \sigma^{-2} \frac{\frac{G_{d-2}(s;\lambda_1(g))}{G_d(s;\lambda_1(g))} - \frac{G_{d-2}(s;\lambda_0(g))}{G_d(s;\lambda_0(g))}}{\frac{G_{d+2}(s;\lambda_1(g))}{G_d(s;\lambda_1(g))} - \frac{G_{d+2}(s;\lambda_0(g))}{G_d(s;\lambda_0(g))}} \right).$$

But by Lemma 17.12, the ratio in the second term is always a negative real number (for both the cases of real eigenvalues and complex conjugate eigenvalues), so the second term is always positive. Thus

$$\frac{\partial_g f_{d\alpha\beta}(g, s)}{\partial_s f_{d\alpha\beta}(g, s)} > \frac{s}{2g}$$

$$h'(g_0) < -\frac{s_0}{2g_0} \qquad \text{[S133]}$$

so increasing gain always reduces the spatial radius of nearby excitation/inhibition, regardless of cell types.

**D. Relationship between the zero crossing locations of inhibitory and excitatory neurons.** Let us also consider the zero crossings of inhibitory neurons compared to that of excitatory neurons. Define $f_d : (0, \infty) \to \mathbb{C}$ by

$$f_d(s) = \frac{G_d(s; \lambda_1)}{G_d(s; \lambda_0)}.$$

Let $s_0^E, s_0^I$ be the first zero-crossing of excitatory neurons and inhibitory neurons respectively in response to the perturbation of a single excitatory neuron, i.e. $s_0^E$ is the smallest positive solution of $f_d(s_0^E) = c_{00}$ while $s_0^I$ is the smallest positive solution of $f_d(s_0^I) = c_{10}$. We claim that for ISNs in two or higher dimensions, $s_0^E < s_0^I$ if and only $\det(\boldsymbol{W}) > 0$.

To see this, first consider the case of real eigenvalues, where $0 < \lambda_0 < \lambda_1$ and $c_{\alpha\beta} \in (0, 1)$. By lemma 17.11 and 17.16, $f_d$ is strictly monotonically decreasing on $(0, \infty)$ with image $(0, 1)$, so its inverse $f_d^{-1} : (0, 1) \to (0, \infty)$ exists and is also strictly monotonically decreasing. Thus $s_0^E < s_0^I$ if and only if $f_d^{-1}(c_{00}) < f_d^{-1}(c_{10})$, so it is sufficient to show that $c_{00} > c_{10}$ if and only if $\det(\boldsymbol{W}) > 0$. To see this, note that the existence of zero crossing for inhibitory neurons implies $c_{10} > 0$, so $c_{00} > c_{10}$ if and only if $\frac{c_{00}}{c_{10}} > 1$. From the definition of $c_{00}$ and $c_{10}$, we see that $\frac{c_{00}}{c_{10}} = \frac{[\boldsymbol{W\Sigma}^{-1}\boldsymbol{Q}]_{00}}{[\boldsymbol{W\Sigma}^{-1}\boldsymbol{Q}]_{01}} \frac{[\boldsymbol{W\Sigma}^{-1}\boldsymbol{Q}]_{11}}{[\boldsymbol{W\Sigma}^{-1}\boldsymbol{Q}]_{10}}$. Now note that $[\boldsymbol{W\Sigma}^{-1}\boldsymbol{Q}]_{01}[\boldsymbol{W\Sigma}^{-1}\boldsymbol{Q}]_{10} = \frac{(1-\sigma_0^2\lambda_1)(1-\sigma_1^2\lambda_0)(1-\sigma_1^2\lambda_1-w_{11})}{\sigma_1^4 w_{10}} > 0$ since for ISNs in two or higher dimensions such that $\tilde{L}_{\alpha\beta}(s)$ exhibits a zero crossing, Corollary 10.4 shows that $1 - \sigma_\alpha^2\lambda_0 < 0$ (which also implies $1 - \sigma_\alpha^2\lambda_1 < 0$ since $0 < \lambda_0 < \lambda_1$), and the proof of Theorem 10.3 shows that $1 - \sigma_1^2\lambda_\alpha - w_{11} > 0$. Thus $c_{00} > c_{10}$ if and only if $\det(\boldsymbol{W\Sigma}^{-1}\boldsymbol{Q}) > 0$. But $\det(\boldsymbol{\Sigma}^{-1}) = \sigma_0^{-2}\sigma_1^{-2} > 0$ and $\det(\boldsymbol{Q}) = \frac{\lambda_0 - \lambda_1}{M_{10}} = \frac{\lambda_1 - \lambda_0}{\sigma_0^{-2} w_{10}} > 0$. Thus $c_{00} > c_{10}$ if and only if $\det(\boldsymbol{W}) > 0$.

Now consider the case of complex conjugate eigenvalues. In this case, both $c_{\alpha\beta}$ and $f_d(s)$ are of the form $z\bar{z}^{-1}$, so $|c_{\alpha\beta}| = |f_d(s)| = 1$, i.e. $c_{\alpha\beta}$ lies on the unit circle and the image of $f_d$ is also the unit circle. Thus $s_0^{\mathrm{E}}, s_0^{\mathrm{I}}$ may be equivalently defined as the smallest positive solutions satisfying $\mathrm{Arg}(f_d(s_0^{\mathrm{E}})) = \mathrm{Arg}(c_{00})$ and $\mathrm{Arg}(f_d(s_0^{\mathrm{I}})) = \mathrm{Arg}(c_{10})$, where $\mathrm{Arg}(z)$ is the argument of $z$ such that $\mathrm{Arg}(z) \in [0, 2\pi)$ (note that this is different from the convention of $\mathrm{Arg}(z) \in (-\pi, \pi]$). We claim that there exists $s_0 > \max\{s_0^{\mathrm{E}}, s_0^{\mathrm{I}}\}$ such that $\mathrm{Arg}(f_d(s))$ is a strictly monotonically decreasing function on the interval $(0, s_0)$, and thus $s_0^{\mathrm{E}} < s_0^{\mathrm{I}}$ if and only if $\mathrm{Arg}(c_{00}) > \mathrm{Arg}(c_{10})$. Indeed, by lemma 17.15, we have

$$\begin{aligned} f_d'(s) &= -2\pi s \frac{G_{d+2}(s;\lambda_1)G_d(s;\lambda_0) - G_{d+2}(s;\lambda_0)G_d(s;\lambda_1)}{G_d(s;\lambda_0)^2} \\ &= -2\pi s f_d(s)\left(\frac{G_{d+2}(s;\lambda_1)}{G_d(s;\lambda_1)} - \frac{G_{d+2}(s;\lambda_0)}{G_d(s;\lambda_0)}\right) \\ &= 4\pi s f_d(s) \mathrm{Im}\left(\frac{G_{d+2}(s;\lambda_0)}{G_d(s;\lambda_0)}\right) i. \end{aligned}$$

But by Lemma 17.12, $\mathrm{Im}\left(\frac{G_{d+2}(s;\lambda_0)}{G_d(s;\lambda_0)}\right) < 0$ for all $d \in \mathbb{N}$ and $s > 0$. Furthermore, on the unit circle we have $\mathrm{Arg}(z) = i^{-1}\mathrm{Log}(z)$ where $\mathrm{Log}(z)$ is the logarithm of $z$ such that $\mathrm{Im}(\mathrm{Log}(z)) \in [0, 2\pi)$. Thus

$$\partial_s \mathrm{Arg}(f_d(s)) = \frac{f_d'(s)}{f_d(s) i} < 0$$

for all $s > 0$ such that $f_d(s) \neq 1$ (we need to exclude these points since $\mathrm{Arg}(z)$, as currently defined, has a branch cut along $[0, \infty)$). Now define $S = \{s \in \mathbb{R} : f_d(s) = 1, s > 0\}$, which is non-empty due to Theorem 17.18. By the Inverse Function Theorem (viewing $f_d$ as a map between two one-dimensional differentiable manifolds, $(0, \infty)$ and $\mathbb{S}$), for all $s' \in S$ there exists some $\epsilon_{s'} > 0$ such that $f_d$ is one-to-one on $U_{s'} := (s' - \epsilon_{s'}, s' + \epsilon_{s'})$. Since $U_{s'}$ does not contain any point in $S$ other than $s'$ (otherwise $f_d$ would not be one-to-one), we have $\partial_s \mathrm{Arg}(f_d(s)) < 0$ for all $s \in U_{s'} \setminus \{s'\}$, and thus $\mathrm{Arg}(f_d(s))$ is strictly monotonically decreasing on both $(s' - \epsilon_{s'}, s')$ and $(s', s' + \epsilon_{s'})$. But since $f_d(s') = 1$, this is only possible if $\mathrm{Arg}(f_d(s))$ is discontinuous at $s'$ and $\lim_{s \to s'^-} \mathrm{Arg}(f_d(s)) = 0$, $\lim_{s \to s'^+} \mathrm{Arg}(f_d(s)) = 2\pi$. Thus, for all $s_1, s_2 \in S$ such that $(s_1, s_2) \cap S = \emptyset$, $\mathrm{Arg}(f_d(s))$ is continuous on $(s_1, s_2)$ with image $(0, 2\pi)$, so by the Intermediate Value Theorem there exists a point $t \in (s_1, s_2)$ such that $\mathrm{Arg}(f_d(t)) = \pi$. Define $T$ to be the set of all such $t$. Now let $s_0 = \inf S$. We claim that $s_0 > 0$. Indeed, suppose for contradiction that $s_0 = 0$. Then we must also have $\inf T = 0$, so we can construct a sequence $(t_n)_{n \in \mathbb{N}}$ such that $\lim_{n \to \infty} t_n = 0$ and $\lim_{n \to \infty} \mathrm{Arg}(f_d(t_n)) = \pi$. But this is impossible since $\lim_{s \to 0^+} f_d(s) = 1$ for $d \geq 2$ by Lemma 17.10. Thus $s_0 > 0$. Since $\mathrm{Arg}(f_d(s))$ is strictly monotonically decreasing on $(0, s_0)$ and $\lim_{s \to 0^+} f_d(s) = 1$, we must have $\lim_{s \to 0^+} \mathrm{Arg}(f_d(s)) = 2\pi$. Thus $\mathrm{Arg}(f_d(s))$ is a bijective mapping from $(0, s_0)$ to $(0, 2\pi)$. But this implies $s_0^{\mathrm{E}}, s_0^{\mathrm{I}}$ must be located in $(0, s_0)$, so we are done.

Since we proved that $s_0^{\mathrm{E}} < s_0^{\mathrm{I}}$ if and only if $\mathrm{Arg}(c_{00}) > \mathrm{Arg}(c_{10})$, it remains to show that $\mathrm{Arg}(c_{00}) > \mathrm{Arg}(c_{10})$ if and only if $\det(\boldsymbol{W}) > 0$. Lemma 11.1 shows that $\mathrm{Arg}(c_{\alpha\beta}^+) \in (0, \pi)$ for all $\alpha, \beta$. Thus we have $\mathrm{Arg}(c_{\alpha\beta}) = 2\mathrm{Arg}(c_{\alpha\beta}^+)$ (note that this is not necessarily true if we took the principal branch). Thus $\mathrm{Arg}(c_{00}) > \mathrm{Arg}(c_{10})$ if and only if $\mathrm{Arg}(c_{00}^+) > \mathrm{Arg}(c_{10}^+)$. But this is true if and only if $\mathrm{Arg}\left(\frac{c_{00}^+}{c_{10}^+}\right) \in (0, \pi)$, or that equivalently, $\mathrm{Im}\left(\frac{c_{00}^+}{c_{10}^+}\right) > 0$. Now $\mathrm{Im}\left(\frac{c_{00}^+}{c_{10}^+}\right) = \frac{1}{i}\left(\frac{c_{00}^+}{c_{10}^+} - \frac{\bar{c}_{00}^+}{\bar{c}_{10}^+}\right) = \frac{1}{i}\left(\frac{(1-\sigma_0^2\lambda_0)(1-\sigma_1^2\lambda_0 - w_{11})}{1-\sigma_1^2\lambda_0} - \frac{(1-\sigma_0^2\lambda_1)(1-\sigma_1^2\lambda_1 - w_{11})}{1-\sigma_1^2\lambda_1}\right)$. It can be shown with some algebra that this is exactly equal to $\frac{1}{i}\det(\boldsymbol{W}\boldsymbol{\Sigma}^{-1}\boldsymbol{Q})$. But $\det(\boldsymbol{\Sigma}^{-1}) = \sigma_0^{-2}\sigma_1^{-2} > 0$ and $\frac{1}{i}\det(\boldsymbol{Q}) = \frac{1}{i}\frac{\lambda_0 - \lambda_1}{M_{10}} = \frac{2\mathrm{Im}(\lambda_1)}{\sigma_0^{-2} w_{10}} > 0$. Thus $\mathrm{Arg}(c_{00}) > \mathrm{Arg}(c_{10})$ if and only if $\det(\boldsymbol{W}) > 0$, so we are done.

#### 12. Distance dependence of perturbation response - location of maximum suppression

Another useful characteristic of the linear response is the distance to its first local minimum. As it turns out, the location of the local mimina are precisely the zero crossings of a model in $d+2$-dimensions. Indeed, taking the derivative of Eq. S94 using Lemma 17.15 and setting the result to zero, we see that for $s > 0$, $s$ must satisfy

$$\sum_{\gamma=0}^{1} [\boldsymbol{W}\boldsymbol{\Sigma}^{-1}\boldsymbol{Q}]_{\alpha\gamma}[\boldsymbol{Q}^{-1}]_{\gamma\beta} G_{d+2}(s;\lambda_\gamma) = 0$$

$$\frac{G_{d+2}(s;\lambda_1)}{G_{d+2}(s;\lambda_0)} = c_{\alpha\beta} \qquad \text{[S134]}$$

so the local minima are the zero crossings of a model in $(d+2)$-dimensions. In particular, this yields an exact closed-form solution for the local minima of a 1D model, since they correspond to the zero crossings of the corresponding 3D model, for which we have already found a closed-form expression in the previous section.

#### 13. Asymptotic distance dependence of perturbation response

Recall that the linear response integrated over feature-tuning, $\tilde{L}_{\alpha\beta}(\boldsymbol{x}-\boldsymbol{y})$, given by Eq. S51, has the form

$$\tilde{L}_{\alpha\beta}(\boldsymbol{x}-\boldsymbol{y}) = \sum_{\rho=0}^{N_c^2-1} \tilde{L}_{0\alpha\beta\rho} G_d(s;\lambda_{0\rho}).$$

where $\tilde{L}_{0\alpha\beta\rho} = [\tilde{\boldsymbol{U}}_0\boldsymbol{Q}_0]_{\alpha\rho}[\boldsymbol{Q}_0^{-1}\tilde{\boldsymbol{V}}_0]_{\rho\beta}$. Due to the asymptotic expansion $K_\eta(z) \sim \sqrt{\frac{\pi}{2z}}e^{-z}\left(1+\frac{4\eta^2-1}{8z}+\cdots\right)$ as $|z|\to\infty$ (13, Equation 9.7.2), we have:

$$\begin{aligned}
\tilde{L}_{\alpha\beta}(s) &= \frac{1}{(2\pi)^{\frac{d}{2}}}\sum_{\rho=0}^{N_c^2-1}\tilde{L}_{0\alpha\beta\rho}\left(\frac{\sqrt{\lambda_{0\rho}}}{s}\right)^{\eta} K_\eta(\sqrt{\lambda_{0\rho}}s) \\
&= \frac{1}{(2\pi)^{\frac{d}{2}}}\sum_{\rho=0}^{N_c^2-1}\tilde{L}_{0\alpha\beta\rho}\left(\frac{\sqrt{\lambda_{0\rho}}}{s}\right)^{\eta}\left(\sqrt{\frac{\pi}{2\sqrt{\lambda_{0\rho}}s}}e^{-\sqrt{\lambda_{0\rho}}s}+\mathcal{O}\left(\frac{e^{-\sqrt{\lambda_{0\rho}}s}}{s^{\frac{3}{2}}}\right)\right) \\
&= \sqrt{\frac{\pi}{2}}\frac{1}{(2\pi)^{\frac{d}{2}}}\sum_{\rho=0}^{N_c^2-1}\tilde{L}_{0\alpha\beta\rho}\sqrt{\lambda_{0\rho}}^{\eta-\frac{1}{2}}e^{-i\mathrm{Im}(\sqrt{\lambda_{0\rho}})s}\frac{e^{-\mathrm{Re}(\sqrt{\lambda_{0\rho}})s}}{s^{\eta+\frac{1}{2}}}+\mathcal{O}\left(\frac{e^{-\mathrm{Re}(\sqrt{\lambda_{0\rho^*}})s}}{s^{\eta+\frac{3}{2}}}\right)
\end{aligned}$$

where

$$\rho^* = \mathrm{argmin}_\rho \mathrm{Re}(\sqrt{\lambda_{0\rho}}). \qquad \text{[S135]}$$

Define $\mathcal{I}_{\mathbb{R}} := \{\rho : \lambda_{0\rho}\in\mathbb{R}\}$ as the set of indices whose corresponding eigenvalues are real, and define $\mathcal{I}_{\mathbb{C}_+} := \{\rho : \mathrm{Im}(\lambda_{0\rho})>0\}$ as the set of indices whose corresponding eigenvalues are in the upper-half complex plane. Assume all eigenvalues have multiplicity of 1 (the following also works without this assumption but is more cumbersome). Let $\bar{\rho}$ denote the index such that $\lambda_{0\bar{\rho}} = \overline{\lambda_{0\rho}}$. Then by Proposition 17.2, $\tilde{L}_{0\alpha\beta\bar{\rho}} = \overline{\tilde{L}_{0\alpha\beta\rho}}$, so

$$\begin{aligned}
\tilde{L}_{0\alpha\beta\rho}\sqrt{\lambda_{0\rho}}^{\eta-\frac{1}{2}}e^{-i\mathrm{Im}(\sqrt{\lambda_{0\rho}})s}+\tilde{L}_{0\alpha\beta\bar{\rho}}\sqrt{\lambda_{0\bar{\rho}}}^{\eta-\frac{1}{2}}e^{-i\mathrm{Im}(\sqrt{\lambda_{0\bar{\rho}}})s} &= 2\mathrm{Re}\left(\tilde{L}_{0\alpha\beta\rho}\sqrt{\lambda_{0\rho}}^{\eta-\frac{1}{2}}e^{-i\mathrm{Im}(\sqrt{\lambda_{0\rho}})s}\right) \\
&= 2a_{0\rho}\cos\left(\mathrm{Im}(\sqrt{\lambda_{0\rho}})s-\phi_{0\rho}\right)
\end{aligned}$$

where $a_{0\rho} = \left|\tilde{L}_{0\alpha\beta\rho}\sqrt{\lambda_{0\rho}}^{\eta-\frac{1}{2}}\right|$ and $\phi_{0\rho} = \mathrm{Arg}\left(\tilde{L}_{0\alpha\beta\rho}\sqrt{\lambda_{0\rho}}^{\eta-\frac{1}{2}}\right)$. Thus

$$\begin{aligned}
\tilde{L}_{\alpha\beta}(s) = \sqrt{\frac{\pi}{2}}\frac{1}{(2\pi)^{\frac{d}{2}}}&\left(\sum_{\rho\in\mathcal{I}_{\mathbb{R}}}\tilde{L}_{0\alpha\beta\rho}\sqrt{\lambda_{0\rho}}^{\frac{d-1}{2}}\frac{e^{-\sqrt{\lambda_{0\rho}}s}}{s^{\frac{d-1}{2}}}+2\sum_{\rho\in\mathcal{I}_{\mathbb{C}_+}}a_{0\rho}\cos\left(\mathrm{Im}(\sqrt{\lambda_{0\rho}})s-\phi_{0\rho}\right)\frac{e^{-\mathrm{Re}(\sqrt{\lambda_{0\rho}})s}}{s^{\frac{d-1}{2}}}\right) \\
&+\mathcal{O}\left(\frac{e^{-\mathrm{Re}(\sqrt{\lambda_{0\rho^*}})s}}{s^{\frac{d+1}{2}}}\right)
\end{aligned} \qquad \text{[S136]}$$

Thus, if $\lambda_{0\rho^*}$ is real, then

$$\tilde{L}_{\alpha\beta}(s) = \sqrt{\frac{\pi}{2}}\frac{1}{(2\pi)^{\frac{d}{2}}}\tilde{L}_{0\alpha\beta\rho}\sqrt{\lambda_{0\rho^*}}^{\frac{d-1}{2}}\frac{e^{-\sqrt{\lambda_{0\rho^*}}s}}{s^{\frac{d-1}{2}}}+\mathcal{O}\left(\frac{e^{-\sqrt{\lambda_{0\rho^*}}s}}{s^{\frac{d+1}{2}}}\right). \qquad \text{[S137]}$$

Otherwise,

$$\tilde{L}_{\alpha\beta}(s) = \sqrt{\frac{\pi}{2}}\frac{2}{(2\pi)^{\frac{d}{2}}}a_{0\rho^*}\cos\left(\mathrm{Im}(\sqrt{\lambda_{0\rho^*}})s-\phi_{0\rho^*}\right)\frac{e^{-\mathrm{Re}(\sqrt{\lambda_{0\rho^*}})s}}{s^{\frac{d-1}{2}}}+\mathcal{O}\left(\frac{e^{-\mathrm{Re}(\sqrt{\lambda_{0\rho^*}})s}}{s^{\frac{d+1}{2}}}\right). \qquad \text{[S138]}$$

In both cases, $\tilde{L}_{\alpha\beta}(s)$ decays asymptotically as

$$\tilde{L}_\infty(s) := \frac{e^{-\mathrm{Re}(\sqrt{\lambda_{0\rho^*}})s}}{s^{\frac{d-1}{2}}}. \quad [S139]$$

Note that here by 'asymptotic decay' we are NOT referring to the mathematical notion of asymptotic equivalence, i.e. $\tilde{L}_{\alpha\beta}(s) \sim \tilde{L}_\infty(s)$ as $s \to \infty$; rather, the notion of 'asymptotic decay' here is precisely defined by equations S137 and S138 themselves.

Let us define the 'decay length scale' by

$$\sigma_\infty := \frac{1}{\mathrm{Re}(\sqrt{\lambda_{0\rho^*}})}. \quad [S140]$$

It is clear that the $\sigma_\infty \to \infty$ as $\lambda_{0\rho^*}$ approaches the negative real line (since $\sqrt{\lambda_{0\rho^*}}$ approaches the imaginary axis). But also recall that assuming sufficiently fast inhibition and $\langle g_1 \mid f\rangle \kappa_{11} w_{11} < 1$, the network is stable if and only if none of the eigenvalues lie on the negative real line (SI section 7C). Thus, this shows that for networks with sufficiently fast inhibition and $\langle g_1 \mid f\rangle \kappa_{11} w_{11} < 1$, as the network approaches instability, its decay length scale tends to infinity.

#### 14. Feature-tuning dependence of perturbation response

**A. Condition for same-favoring response.** We now consider the perturbation response integrated over space, $\tilde{L}_{\alpha\beta}(\mu, \nu, \theta - \phi)$, and study the condition under which iso-tuned neurons are more excited relative to ortho-tuned neurons, i.e. the condition under which the response is same-favoring. By inspection of Eq. S54, this is the case if and only if

$$\tilde{c}_{\alpha\beta} := k_\alpha^{-1}[(\boldsymbol{I} - \boldsymbol{K}\tilde{\boldsymbol{W}})^{-1} - \boldsymbol{I}]_{\alpha\beta} > 0. \quad [S141]$$

where $k_\alpha$ are the diagonal entries of $\boldsymbol{K}$ and are given by Eq. S53. Since in general $\tilde{c}_{\alpha\beta}$ are quite complicated, we will again specifically study the case of a E-I network. In this case, we have

$$\tilde{c}_{00} = \frac{1}{k_0}\left(\frac{1 - k_1\kappa_{11}w_{11}}{D} - 1\right), \qquad \tilde{c}_{01} = \frac{\kappa_{01}w_{01}}{D},$$
$$\tilde{c}_{10} = \frac{\kappa_{10}w_{10}}{D}, \qquad \tilde{c}_{11} = \frac{1}{k_1}\left(\frac{1 - k_0\kappa_{00}w_{00}}{D} - 1\right),$$

where $D = \det(\boldsymbol{I} - \boldsymbol{K}\tilde{\boldsymbol{W}})$. For convenience let us define

$$\tilde{w}_{\alpha\beta} := k_\alpha \kappa_{\alpha\beta}|w_{\alpha\beta}|. \quad [S142]$$

Note that $\tilde{w}_{\alpha\beta}$ are distinct from the entries of the matrix $\tilde{\boldsymbol{W}}$ (unfortunately we are running out of variable names), and that $\kappa_{\alpha\beta} > 0 \iff \tilde{w}_{\alpha\beta} > 0$. Thus we have the following respective conditions for $\tilde{L}_{00}, \tilde{L}_{01}, \tilde{L}_{10}, \tilde{L}_{11}$ to exhibit same-favoring response:

$$-\tilde{w}_{11} < 1 - D, \qquad \kappa_{01} < 0, \qquad \kappa_{10} > 0, \qquad \tilde{w}_{00} < 1 - D, \quad [S143]$$

where we used the fact that stability constraint implies $D > 0$ (see Eq. S79 and note that $\det(\boldsymbol{I} - \boldsymbol{K}\tilde{\boldsymbol{W}}) = \det(\boldsymbol{I} - \tilde{\boldsymbol{W}}\boldsymbol{K})$ by Sylvester's determinant theorem), and that $w_{10} > 0$, $w_{01} < 0$.

Note that the conditions for $\tilde{L}_{01}, \tilde{L}_{10}$ to exhibit same-favoring response is very simple. As one would expect from intuition, the response of E neurons to perturbations of I neurons is same-favoring if and only if I $\to$ E connections are anti-like-to-like, while the response of I neurons to perturbations of E neurons is same-favoring if and only if E $\to$ I connections are like-to-like. On the other hand, the conditions for $\tilde{L}_{00}, \tilde{L}_{11}$ can be written out more explicitly by noting that

$$\begin{aligned} 1 - D &= 1 - (1 - k_0\kappa_{00}w_{00} - k_1\kappa_{11}w_{11} + k_0k_1\kappa_{00}\kappa_{11}w_{00}w_{11} - k_0k_1\kappa_{01}\kappa_{10}w_{01}w_{10}) \\ &= \tilde{w}_{00} - \tilde{w}_{11} + \tilde{w}_{00}\tilde{w}_{11} - \tilde{w}_{10}\tilde{w}_{01}, \end{aligned}$$

so the conditions for $\tilde{L}_{00}, \tilde{L}_{11}$ to exhibit same-favoring response respectively becomes

$$\tilde{w}_{01}\tilde{w}_{10} < \tilde{w}_{00}(\tilde{w}_{11} + 1), \quad [S144]$$
$$\tilde{w}_{01}\tilde{w}_{10} < \tilde{w}_{11}(\tilde{w}_{00} - 1). \quad [S145]$$

**B. Opposite-favoring response implies like-to-like disynaptic E $\to$ I $\to$ E inhibition.** Suppose E $\to$ E connectivity is like-to-like, a fact which is well-supported by experimental data, so that $\kappa_{00} > 0$ and thus $\tilde{w}_{00} > 0$. We claim that if the network is stable and the response of excitatory neurons to perturbation of a single-excitatory neuron is opposite-favoring, i.e. $\tilde{w}_{01}\tilde{w}_{10} > \tilde{w}_{00}(\tilde{w}_{11}+1)$, then we must have like-to-like E $\to$ I $\to$ E inhibition, i.e. $\kappa_{01}\kappa_{10} > 0$. Indeed, suppose for the contrary that $\kappa_{01}\kappa_{10} \leq 0$, which also implies $\tilde{w}_{01}\tilde{w}_{10} \leq 0$. Then

$$\begin{aligned} \tilde{w}_{00}(\tilde{w}_{11}+1) &< \tilde{w}_{01}\tilde{w}_{10} \\ &\leq 0 \\ \tilde{w}_{11} &< -1. \end{aligned}$$

But now we claim that $\tilde{w}_{01}\tilde{w}_{10} \leq 0$ and $\tilde{w}_{11} < -1$ together implies instability, so we have a contradiction. Indeed, in SI section 7C, we showed that an E-I network is stable if and only if $\mathrm{tr}(\boldsymbol{J}_n(k)) < 0$ and $\det(\boldsymbol{J}_n(k)) > 0$ for all $n, k$, where $\boldsymbol{J}_n(k)$ is defined by Eq. S73. This yields a necessary condition for stability as $\mathrm{tr}(\boldsymbol{J}_1(0)) < 0$. On the other hand, we have another necessary condition for stability given by $D > 0$. As we now show, we cannot satisfy both stability conditions simultaneously. Indeed, since

$$\mathrm{tr}(\boldsymbol{J}_1(0)) = \tau_0^{-1}(k_0\kappa_{00}w_{00} - 1) + \tau_1^{-1}(k_1\kappa_{11}w_{11} - 1) = \tau_0^{-1}(\tilde{w}_{00} - 1) - \tau_1^{-1}(\tilde{w}_{11} + 1),$$

the first stability condition along with $\tilde{w}_{11} < -1$ thus implies

$$\begin{aligned} \tau_0^{-1}(\tilde{w}_{00} - 1) &< \tau_1^{-1}(\tilde{w}_{11} + 1) \\ &< 0 \\ \tilde{w}_{00} &< 1. \end{aligned}$$

But $\tilde{w}_{01}\tilde{w}_{10} \leq 0$, $\tilde{w}_{11} < -1$, and $\tilde{w}_{00} < 1$ together implies

$$\begin{aligned} D &= 1 - \tilde{w}_{00} + \tilde{w}_{11} - \tilde{w}_{00}\tilde{w}_{11} + \tilde{w}_{01}\tilde{w}_{10} \\ &\leq (1 - \tilde{w}_{00})(1 + \tilde{w}_{11}) \\ &< 0. \end{aligned}$$

Thus the other stability condition cannot be simultaneously satisfied, so we are done.

**C. Modulation of feature-tuning dependence by neuronal gain.** Now let us study how the baseline firing rate in a nonlinear network affects the feature-tuning dependence of the linearized perturbation response. As discussed in SI section 10D and 11C, a change in baseline firing rate changes the gain of neurons $g$, which in turn changes the linearized connectivity matrix $g\boldsymbol{W}$. For a more fine-grained understanding, we allow the gains of excitatory neurons ($g_0$) and inhibitory neurons ($g_1$) to be independent of each other. Define

$$z_\alpha(g_0, g_1) := g_0 g_1(\tilde{w}_{01}\tilde{w}_{10} - \tilde{w}_{00}\tilde{w}_{11}) - g_\alpha k_\alpha \kappa_{\alpha\alpha} w_{\alpha\alpha} \qquad \text{[S146]}$$

such that $\tilde{L}_{\alpha\alpha}$ is same-favoring if and only if $z_\alpha < 0$. Then increasing the gain of cell type $\beta$ ($g_\beta$) at the phase boundary $z_\alpha(g_0, g_1) = 0$ results in a transition from same-favoring to opposite-favoring response if and only if $\partial_{g_\beta} z_\alpha(g_0, g_1) > 0$. But along the phase boundary we have

$$\begin{aligned} g_\beta \partial_{g_\beta} z_\alpha(g_0, g_1) &= g_0 g_1(\tilde{w}_{01}\tilde{w}_{10} - \tilde{w}_{00}\tilde{w}_{11}) - \delta_{\alpha\beta} g_\beta k_\alpha \kappa_{\alpha\alpha} w_{\alpha\alpha} \\ &= z_\alpha(g_0, g_1) + (1 - \delta_{\alpha\beta}) g_\alpha k_\alpha \kappa_{\alpha\alpha} w_{\alpha\alpha} \\ &= \begin{cases} 0, & \alpha = \beta \\ g_\alpha k_\alpha \kappa_{\alpha\alpha} w_{\alpha\alpha}, & \alpha \neq \beta \end{cases} \end{aligned}$$

Thus increasing the gain of cell type $\alpha$ cannot result in a transition from same-favoring to opposite-favoring response for $\tilde{L}_{\alpha\alpha}$. Furthermore, since we assume $\kappa_{00} > 0$, we see that increasing the gain of inhibitory neurons $g_1$ at the phase boundary results in a transition from same-favoring to opposite-favoring response for $\tilde{L}_{00}$. On the other hand, increasing the gain of excitatory neurons $g_0$ at the phase boundary results in a transition from same-favoring to opposite-favoring response for $\tilde{L}_{11}$ if and only if $\kappa_{11} < 0$.

A stronger statement can be made if $\kappa_{00} > 0$ and the response of excitatory neurons is opposite-favoring, i.e. $z_0(g_0, g_1) > 0$, for some given gains $g_0, g_1$. This is the regime which the data suggests we are in. Since

$$z_0(g_0, g_1) = g_0\left(g_1(\tilde{w}_{01}\tilde{w}_{10} - \tilde{w}_{00}\tilde{w}_{11}) - k_0\kappa_{00}w_{00}\right) \qquad \text{[S147]}$$

we see that $z_0(g_0, g_1) > 0$ and $\kappa_{00} > 0$ implies $\tilde{w}_{01}\tilde{w}_{10} - \tilde{w}_{00}\tilde{w}_{11} > 0$. But this in turn implies that there exists a sufficiently small $g_1'$ such that $z_0(g_0, g_1') < 0$. Thus, not only does decreasing the gain of inhibitory neurons at the phase boundary result in a transition from opposite- to same-favoring excitatory neuron response, but also it is the case that a transition from opposite- to same-favoring excitatory neuron response must occur given sufficiently weak inhibitory neuron gain.

#### 15. Modulation of feature-tuning dependence by distance

Finally, we study how the feature-tuning dependence of perturbation response is modulated by distance. Specifically we consider two questions: 1) whether perturbation response is locally same-favoring or opposite-favoring, and 2) whether or not perturbation response at different distances may have different feature-tuning dependences, e.g. nearby response is same-favoring but far-away response is opposite-favoring. By inspection of Eq. S50, we see that these two questions are determined by the behavior of $\tilde{L}_{1\alpha\beta}(s)$ (Eq. S48): the first question is determined by the sign of $\lim_{s\to 0} \tilde{L}_{1\alpha\beta}(s)$, while the second question is determined by the number of zero crossings of $\tilde{L}_{1\alpha\beta}(s)$. But notice that $\tilde{L}_{1\alpha\beta}(s)$ has the same structure as the perturbation response integrated over feature-tuning, $\tilde{L}_{\alpha\beta}(s)$ (Eq. S51), for which we have already addressed both questions in SI sections 9 and 10. Thus we just need to slightly adapt our analysis from those sections.

For the first question, we note that the perturbation response is locally same-favoring if and only if $\lim_{s\to 0} \tilde{L}_{1\alpha\beta}(s) > 0$. But applying the same reasoning as in SI section 9, we find that for $d \geq 2 \in \mathbb{N}$, $\lim_{s\to 0} \tilde{L}_{1\alpha\beta}(s) > 0$ if and only if

$$[\tilde{\boldsymbol{U}}_1 \tilde{\boldsymbol{V}}_1]_{\alpha\beta} = w_{\alpha\beta}\sigma_{\alpha\beta}^{-2}\kappa_{\alpha\beta} > 0 \qquad \text{[S148]}$$

Thus, if $\beta$ is an excitatory cell type, then $\lim_{s\to 0} \tilde{L}_{1\alpha\beta}(s) > 0$ if and only if $\kappa_{\alpha\beta} > 0$. Similarly, if $\beta$ is an inhibitory cell type, then the condition is that $\kappa_{\alpha\beta} < 0$. In particular, given E $\to$ E is experimentally determined to be like-to-like, i.e. $\kappa_{00} > 0$, this implies perturbation of a single excitatory neuron must produce locally same-favoring response.

For the second question, let us consider the number of zero crossings of $\tilde{L}_{1\alpha\beta}(s)$. As in SI section 10, we will consider a E-I network where connectivity width depends only on presynaptic cell type, such that $\tilde{L}_{1\alpha\beta}(s)$ simplifies to Eq. S61, given by

$$\tilde{L}_{1\alpha\beta}(s) = \sum_{\rho=0}^{1} [\tilde{\boldsymbol{W}}\boldsymbol{\Sigma}^{-1}\tilde{\boldsymbol{Q}}]_{\alpha\rho}[\tilde{\boldsymbol{Q}}^{-1}]_{\rho\beta} G_d(s; \tilde{\lambda}_{1\rho}) \qquad \text{[S149]}$$

where we dropped the tilde from $\tilde{\boldsymbol{\Sigma}}$ and dropped the subscripts of 1 from $\tilde{\boldsymbol{Q}}_1$, $\tilde{\lambda}_{1\rho}$ in Eq. S61 for notational simplicity. By the same analysis as in SI section 10A, $\tilde{L}_{1\alpha\beta}(s)$ has infinitely many zero crossings if and only if the eigenvalues $\tilde{\lambda}_0, \tilde{\lambda}_1$ of $\boldsymbol{M} := (\boldsymbol{I} - \boldsymbol{K}\tilde{\boldsymbol{W}})\boldsymbol{\Sigma}^{-1}$ are complex conjugates. For real eigenvalues, the analysis from SI section 10A can be recycled by substituting instances of $\boldsymbol{W}$ with $\boldsymbol{K}\tilde{\boldsymbol{W}}$ and instance of $w_{\alpha\beta}$ with $k_\alpha \kappa_{\alpha\beta} w_{\alpha\beta}$. Specifically, $\tilde{L}_{1\alpha\beta}(s)$ has exactly one zero-crossing if and only if

$$\tilde{c}_{\alpha\beta} := -\frac{[\boldsymbol{K}\tilde{\boldsymbol{W}}\boldsymbol{\Sigma}^{-1}\tilde{\boldsymbol{Q}}]_{\alpha 0}[\tilde{\boldsymbol{Q}}^{-1}]_{0\beta}}{[\boldsymbol{K}\tilde{\boldsymbol{W}}\boldsymbol{\Sigma}^{-1}\tilde{\boldsymbol{Q}}]_{\alpha 1}[\tilde{\boldsymbol{Q}}^{-1}]_{1\beta}} \in \begin{cases} (0,1), & d \geq 2 \\ (0, \sqrt{\frac{\tilde{\lambda}_0}{\tilde{\lambda}_1}}), & d = 1 \end{cases}. \qquad \text{[S150]}$$

where

$$\tilde{c}_{00} = \frac{(1-\sigma_0^2\tilde{\lambda}_0)(1-\sigma_1^2\tilde{\lambda}_0 - k_1\kappa_{11}w_{11})}{(1-\sigma_0^2\tilde{\lambda}_1)(1-\sigma_1^2\tilde{\lambda}_1 - k_1\kappa_{11}w_{11})}, \qquad \tilde{c}_{01} = \frac{1-\sigma_0^2\tilde{\lambda}_0}{1-\sigma_0^2\tilde{\lambda}_1},$$

$$\tilde{c}_{10} = \frac{1-\sigma_1^2\tilde{\lambda}_0}{1-\sigma_1^2\tilde{\lambda}_1}, \qquad \tilde{c}_{11} = \frac{(1-\sigma_1^2\tilde{\lambda}_0)(1-\sigma_1^2\tilde{\lambda}_1 - k_1\kappa_{11}w_{11})}{(1-\sigma_1^2\tilde{\lambda}_1)(1-\sigma_1^2\tilde{\lambda}_0 - k_1\kappa_{11}w_{11})}. \qquad \text{[S151]}$$

However, the analogue of some theorems in SI section 10A are only conditionally valid here. The analogue of Theorem 10.1 still holds, but the analogues of Theorem 10.2 and 10.3 require additional assumptions since the inequalities $w_{\alpha 0} > 0$, $w_{\alpha 1} < 0$ were used in the proofs, but the analogues of these inequalities are only true if $\kappa_{\alpha\beta} > 0$ for all $\alpha, \beta$. Specifically, if we let $\tilde{c}_{\alpha\beta}^+$ be the numerators of Eq. S151, then we have the following modified theorems:

**Theorem 15.1.** *For all $d \in \mathbb{N}$ and $(\alpha, \beta) \in \{(0,1),(1,0)\}$, $\tilde{L}_{1\alpha\beta}$ has exactly one zero crossing if and only if $\tilde{c}_{\alpha\beta}^+ < 0$.*

**Theorem 15.2.** *For all $d \geq 2 \in \mathbb{N}$ and $\alpha \in \{0,1\}$, if $\kappa_{\alpha\alpha} > 0$, then $\tilde{L}_{1\alpha\alpha}$ has exactly one zero crossing if and only if $\tilde{c}_{\alpha\alpha}^+ < 0$.*

**Theorem 15.3.** *Suppose $M_{00} - M_{11} < 0$ and $\kappa_{01}\kappa_{10} > 0$. For all $d \geq 2 \in \mathbb{N}$ and $\alpha \in \{0,1\}$, if $\kappa_{\alpha\alpha} > 0$, then $\tilde{L}_{1\alpha\alpha}$ has exactly one zero crossing if and only if $1 - \sigma_\alpha^2\tilde{\lambda}_0 < 0$.*

We omit the proofs since they are completely analogous to Theorems 10.1, 10.2, and 10.3.

A natural question to ask is what happens if $\kappa_{\alpha\beta} < 0$ for some $\alpha, \beta$. In particular, let us consider what happens to Theorem 15.3 if the condition $\kappa_{01}\kappa_{10} > 0$ is violated, i.e. if the disynaptic E $\to$ I $\to$ E inhibition is anti-like-to-like rather than like-to-like. We find that

**Theorem 15.4.** *If $\kappa_{00} > 0$ and $\kappa_{01}\kappa_{10} < 0$, then for all $d \geq 2 \in \mathbb{N}$, $\tilde{L}_{100}$ has no zero crossing.*

*Proof.* We need to show two things: 1) the eigenvalues $\tilde{\lambda}_0, \tilde{\lambda}_1$ are real, and 2) $\tilde{c}_{00}^+ > 0$. Our claim then follows from Theorem 15.2 and the fact that there is at most one zero crossing when the eigenvalues are real (Theorem 17.19).

First, the eigenvalues are real if $\mathrm{tr}(\boldsymbol{M})^2 - 4\det(\boldsymbol{M}) > 0$. But

$$\mathrm{tr}(\boldsymbol{M})^2 - 4\det(\boldsymbol{M}) = M_{00}^2 + 2M_{00}M_{11} + M_{11}^2 - 4(M_{00}M_{11} - M_{01}M_{10}) = (M_{00} - M_{11})^2 + 4M_{01}M_{10}$$

and

$$M_{01}M_{10} = \sigma_0^{-2}\sigma_1^{-2}k_0k_1\kappa_{01}\kappa_{10}w_{01}w_{10} > 0$$

since $\kappa_{01}\kappa_{10} < 0$. Thus

$$\mathrm{tr}(\boldsymbol{M})^2 - 4\det(\boldsymbol{M}) > (M_{00} - M_{11})^2 > 0.$$

so indeed $\tilde{\lambda}_0, \tilde{\lambda}_1$ must be real.

Second, we claim that $1 - \sigma_0^2\tilde{\lambda}_0 > 0$ and $1 - \sigma_1^2\tilde{\lambda}_0 - k_1\kappa_{11}w_{11} > 0$, which implies $\tilde{c}_{00}^+ > 0$. For the first claim, notice that since $M_{01}M_{10} > 0$, we have

$$\begin{aligned}\tilde{\lambda}_0 &= \frac{1}{2}(M_{00} + M_{11} - \sqrt{(M_{00} - M_{11})^2 + 4M_{01}M_{10}}) \\ &< \frac{1}{2}(M_{00} + M_{11} - |M_{00} - M_{11}|).\end{aligned}$$

If $M_{00} > M_{11}$, then $\tilde{\lambda}_0 < M_{11} < M_{00}$. On the other hand, if $M_{00} \leq M_{11}$, then $\tilde{\lambda}_0 < \frac{1}{2}(M_{00} + M_{11} + M_{00} - M_{11}) = M_{00}$. In either case we have $\tilde{\lambda}_0 < M_{00}$. But $M_{00} = \sigma_0^{-2}(1 - k_0\kappa_{00}w_{00})$, and since we assumed $\kappa_{00} > 0$, we must have $\tilde{\lambda}_0 < \sigma_0^{-2}$, and thus $1 - \sigma_0^2\tilde{\lambda}_0 > 0$. For the second claim, notice that since $M_{11} = \sigma_1^{-2}(1 - k_1\kappa_{11}w_{11})$, $1 - \sigma_1^2\tilde{\lambda}_0 - k_1\kappa_{11}w_{11} > 0$ if and only if $\tilde{\lambda}_\alpha < M_{11}$, i.e.

$$\begin{aligned}\frac{1}{2}\left(M_{00} + M_{11} - \sqrt{(M_{00} - M_{11})^2 + 4M_{01}M_{10}}\right) &< M_{11} \\ M_{00} - M_{11} &< \sqrt{(M_{00} - M_{11})^2 + 4M_{01}M_{10}}.\end{aligned}$$

But this is always true since $M_{01}M_{10} > 0$. □

#### 16. Discretization of continuum model

In Figure 1D we compared our analytical linear response equation to numerical simulations. This requires discretizing our continuum model into a model with a finite number of neurons. Here we explain how we discretize our equations which contain Dirac delta functions and show that the response of unperturbed neurons to single-cell perturbation scales as $N^{-1}$. We also explain and justify our procedure for dealing with the divergence in the spatial kernel $G_d(s;\lambda)$ during discretization.

**A. Unperturbed neuron response scales as $N^{-1}$.** For simplicity, we will consider the case where feature selectivity is uniformly distributed, i.e. $P_\alpha(\mu) = 1$. The steady state equation of the continuum model is then given by

$$r_\alpha(\mu, \boldsymbol{x}, \theta) = \sum_{\beta=0}^{N_c-1}\int_0^1\int_{\mathbb{R}^d}\int_{-\pi}^{\pi} W_{\alpha\beta}(\mu, \nu, \boldsymbol{x} - \boldsymbol{y}, \theta - \phi)r_\beta(\nu, \boldsymbol{y}, \phi)\, d\phi d\boldsymbol{y} d\nu + h_\alpha(\mu, \boldsymbol{x}, \theta). \quad \text{[S152]}$$

Let us discretize the continuum model on a regular grid of neurons composed of $N_\mu$ feature selectivities, $N_1 \times \cdots \times N_d$ spatial locations within a $l_1 \times \cdots \times l_d$ rectangular region, and $N_\theta$ feature tuning preferences. Thus in total we have $N := N_c N_\mu N_\theta \prod_{i=1}^d N_i$ neurons. For all $i \in \{1, \cdots, N\}$, let us define $\alpha_i, \mu_i, \boldsymbol{x}_i, \theta_i$ to be the cell type, feature selectivity, spatial location, and feature tuning preference of neuron $i$.

Given $k \in \{1, \cdots, N\}$, consider the perturbation of neuron $(\alpha_k, \mu_k, \boldsymbol{x}_k, \theta_k)$ in the continuum model. By Eq. 11 in the main text, we have

$$h_\alpha(\mu, \boldsymbol{x}, \theta) = h\delta_{\alpha\alpha_k}\delta(\mu - \mu_k)\delta(\boldsymbol{x} - \boldsymbol{x}_k)\delta(\theta - \theta_k),$$

and the steady-state solution is given by

$$\begin{aligned}
r_\alpha(\mu, \boldsymbol{x}, \theta) &= \langle \alpha, \mu, \boldsymbol{x}, \theta \,|\, r\rangle \\
&= \langle \alpha, \mu, \boldsymbol{x}, \theta | L | h\rangle \\
&= h\langle \alpha, \mu, \boldsymbol{x}, \theta | (I + \tilde{L}) | \alpha_k, \mu_k, \boldsymbol{x}_k, \phi_k\rangle \\
&= h(\langle \alpha, \mu, \boldsymbol{x}, \theta \,|\, \alpha_k, \mu_k, \boldsymbol{x}_k, \phi_k\rangle + \langle \alpha, \mu, \boldsymbol{x}, \theta | \tilde{L} | \alpha_k, \mu_k, \boldsymbol{x}_k, \phi_k\rangle) \\
&= h(\delta_{\alpha\alpha_k}\delta(\mu - \mu_k)\delta(\boldsymbol{x} - \boldsymbol{x}_k)\delta(\theta - \theta_k) + \tilde{L}_{\alpha\alpha_k}(\mu, \mu_k, \boldsymbol{x} - \boldsymbol{x}_k, \theta - \theta_k)) \\
&= h\delta_{\alpha\alpha_k}\delta(\mu - \mu_k)\delta(\boldsymbol{x} - \boldsymbol{x}_k)\delta(\theta - \theta_k) + \tilde{r}_\alpha(\mu, \boldsymbol{x}, \theta)
\end{aligned}$$

where

$$\tilde{r}_\alpha(\mu, \boldsymbol{x}, \theta) := h\tilde{L}_{\alpha\alpha_k}(\mu, \mu_k, \boldsymbol{x} - \boldsymbol{x}_k, \theta - \theta_k). \tag{S153}$$

denotes the continuous part of the perturbation response. Then the integral term in S152 can be approximated as

$$\begin{aligned}
&\sum_{\beta=0}^{N_c-1} \int_0^1 \int_{\mathbb{R}^d} \int_{-\pi}^{\pi} W_{\alpha\beta}(\mu, \nu, \boldsymbol{x} - \boldsymbol{y}, \theta - \phi) r_\beta(\nu, \boldsymbol{y}, \phi)\, d\phi d\boldsymbol{y} d\nu \\
&= hW_{\alpha\alpha_k}(\mu, \mu_k, \boldsymbol{x} - \boldsymbol{x}_k, \theta - \theta_k) + \sum_{\beta=0}^{N_c-1} \int_0^1 \int_{\mathbb{R}^d} \int_{-\pi}^{\pi} W_{\alpha\beta}(\mu, \nu, \boldsymbol{x} - \boldsymbol{y}, \theta - \phi) \tilde{r}_\beta(\nu, \boldsymbol{y}, \phi)\, d\phi d\boldsymbol{y} d\nu \\
&\approx hW_{\alpha\alpha_k}(\mu, \mu_k, \boldsymbol{x} - \boldsymbol{x}_k, \theta - \theta_k) + \sum_{j=1}^{N} W_{\alpha\alpha_j}(\mu, \mu_j, \boldsymbol{x} - \boldsymbol{x}_j, \theta - \theta_j) \tilde{r}_{\alpha_j}(\mu_j, \boldsymbol{x}_j, \theta_j)\Delta V \\
&= \sum_{j=1}^{N} W_{\alpha\alpha_j}(\mu, \mu_j, \boldsymbol{x} - \boldsymbol{x}_j, \theta - \theta_j) \left(\tilde{r}_{\alpha_j}(\mu_j, \boldsymbol{x}_j, \theta_j)\Delta V + h\delta_{jk})\right) \\
&= \sum_{j=1}^{N} W_{\alpha\alpha_j}(\mu, \mu_j, \boldsymbol{x} - \boldsymbol{x}_j, \theta - \theta_j) r_j^{\text{dis}}
\end{aligned}$$

where

$$\Delta V := \left(\frac{1}{N_\mu}\right)\left(\frac{l_1}{N_1} \cdots \frac{l_d}{N_d}\right)\left(\frac{2\pi}{N_\theta}\right) = \frac{2\pi N_c \prod_{i=1}^d l_i}{N} \tag{S154}$$

$$r_i^{\text{dis}} := \tilde{r}_{\alpha_i}(\mu_i, \boldsymbol{x}_i, \theta_i)\Delta V + h\delta_{ik} \tag{S155}$$

For all $i \in \{1, \cdots, N\}$, let $U_i$ the grid cell containing neuron $(\alpha_i, \mu_i, \boldsymbol{x}_i, \theta_i)$, and apply the operator

$$\sum_{\alpha=0}^{N_c-1} \delta_{\alpha\alpha_i} \iiint_{U_i} d\mu d\boldsymbol{x} d\theta$$

to both sides of Eq. S152. Defining

$$\begin{aligned}
W_{ij}^{\text{dis}} &:= \iiint_{U_i} W_{\alpha_i\alpha_j}(\mu, \mu_j, \boldsymbol{x} - \boldsymbol{x}_j, \theta - \theta_j)\, d\mu d\boldsymbol{x} d\theta \\
&\approx W_{\alpha_i\alpha_j}(\mu_i, \mu_j, \boldsymbol{x}_i - \boldsymbol{x}_j, \theta_i - \theta_j)\Delta V,
\end{aligned} \tag{S156}$$

and noting that

$$\begin{aligned}
\iiint_{U_i} r_{\alpha_i}(\mu, \boldsymbol{x}, \theta)\, d\mu d\boldsymbol{x} d\theta &= h\delta_{ik} + \iiint_{U_i} \tilde{r}_{\alpha_i}(\mu, \boldsymbol{x}, \theta)\, d\mu d\boldsymbol{x} d\theta \\
&\approx r_{\alpha_i}(\mu_i, \boldsymbol{x}_i, \theta_i)\Delta V + h\delta_{ik} \\
&= r_i^{\text{dis}},
\end{aligned}$$

$$\iiint_{U_i} h_{\alpha_i}(\mu, \boldsymbol{x}, \theta)\, d\mu d\boldsymbol{x} d\theta = h\delta_{ik},$$

we get, for all $i \in \{1, \cdots, N\}$,

$$r_i^{\text{dis}} \approx \sum_{j=1}^{N} W_{ij}^{\text{dis}} r_j^{\text{dis}} + h\delta_{ik}. \quad \text{[S157]}$$

Thus, for a discrete model with connectivity given by $W_{ij}^{\text{dis}}$, the steady state response to the single cell perturbation $h\delta_{ik}$ is approximately given by $r_i^{\text{dis}}$. From the definition of $r_i^{\text{dis}}$ (Eq. S155), we see that the responses of unperturbed neurons scale proportionally to $\Delta V \propto N^{-1}$.

**B. Treatment of the divergence of $G_d(s;\lambda)$ during discretization.** We note that our above derivative has one slight issue for $d \geq 2$: the spatial kernel $G_d(s;\lambda)$ diverges as $s \to 0$, and thus $W_{ij}^{\text{dis}}$ is not well-defined when $\boldsymbol{x}_i = \boldsymbol{x}_j$ and $r_i^{\text{dis}}$ is not well-defined when $\boldsymbol{x}_i = \boldsymbol{x}_k$. To deal with this issue, in our numerical simulations we simply set $W_{ij}^{\text{dis}} = 0$ if $\boldsymbol{x}_i = \boldsymbol{x}_j$ and $r_i^{\text{dis}} = h\delta_{ik}$ if $\boldsymbol{x}_i = \boldsymbol{x}_k$. This procedure is valid for $d = 2, 3$, justified numerically by the fact that our analytics agree with the numerical simulations to a very high degree (the case of $d = 2$ is showcased by Figure 1D; the case of $d = 3$ is validated by our test code). Here we provide an informal argument for why this procedure incurs vanishing error as $N \to \infty$ in 2 and 3 spatial dimensions.

Consider the error in the right hand side of Eq. S157 (or equivalently, the error in $\sum_j W_{ij}^{\text{dis}} r_j^{\text{dis}}$) due to this procedure. Since $r_i^{\text{dis}}$ scales as $N^{-1}$, we want to show that this error decays faster than $N^{-1}$, so that this error is negligible for large $N$. For all $i \in \{1, \cdots, N\}$, let

$$B_i = \left(x_i^1 - \frac{l_1}{2N_1}, x_i^1 + \frac{l_1}{2N_1}\right) \times \cdots \times \left(x_i^d - \frac{l_d}{2N_d}, x_i^d + \frac{l_d}{2N_d}\right) \subset \mathbb{R}^d$$

denote the spatial grid cell centered at $\boldsymbol{x}_i$, where $x_i^1, \cdots, x_i^d$ are the elements of $\boldsymbol{x}_i$. If $\boldsymbol{x}_i \neq \boldsymbol{x}_k$, then the error is given by

$$\Delta V h \sum_{\beta=0}^{N_c-1} \int_0^1 \int_{B_i \cup B_k} \int_{-\pi}^{\pi} W_{\alpha_i\beta}(\mu_i, \nu, \boldsymbol{x}_i - \boldsymbol{y}, \theta_i - \phi) \tilde{L}_{\beta\alpha_k}(\nu, \mu_k, \boldsymbol{y} - \boldsymbol{x}_k, \phi - \theta_k)\, d\phi d\boldsymbol{y} d\nu.$$

Otherwise, then the error is given by

$$\Delta V h \sum_{\beta=0}^{N_c-1} \int_0^1 \int_{B_i} \int_{-\pi}^{\pi} W_{\alpha_i\beta}(\mu_i, \nu, \boldsymbol{x}_i - \boldsymbol{y}, \theta_i - \phi) \tilde{L}_{\beta\alpha_i}(\nu, \mu_i, \boldsymbol{y} - \boldsymbol{x}_i, \phi - \theta_i)\, d\phi d\boldsymbol{y} d\nu.$$

We expect the first error to be less than the second, since in the second case, the divergences in $W$ and $\tilde{L}$ are 'lined-up' together. So let us focus on the second error term. It is clear that this error term must be bounded by

$$N^{-1} M \int_{B_i} G_d(\|\boldsymbol{y} - \boldsymbol{x}_i\|; \lambda_0) G_d(\|\boldsymbol{y} - \boldsymbol{x}_i\|; \lambda_1)\, d\boldsymbol{y}$$

for some constant $M > 0$ and $\lambda_0, \lambda_1 \in \mathbb{C} \setminus (-\infty, 0]$. Consider the simplest case where $\frac{l_1}{N_1} = \cdots = \frac{l_d}{N_d}$. Define $R = \frac{\sqrt{d} l_1}{2N_1}$. Then the error is bounded by

$$N^{-1} M' \int_0^R G_d(s;\lambda_0) G_d(s;\lambda_1) s^{d-1}\, ds$$

for some new constant $M' > 0$. Now apply the asymptotic expressions for $G_d(s;\lambda)$ as $s \to 0$ given by Eq. S165. We can apply this since the upper integration limit $R \to 0$ as $N \to \infty$. Then for $d = 2$, the error upper bound tends to

$$\begin{aligned}
N^{-1} M'' \int_0^R s(\ln s)^2\, ds &= N^{-1} M'' \left( \frac{1}{2}(s \ln s)^2 \Big|_0^R - \int_0^R s \ln s\, ds \right) \\
&= N^{-1} M'' \left( \frac{1}{2}(R \ln R)^2 - \frac{1}{2} s^2 \ln s \Big|_0^R + \frac{1}{2} \int_0^R s\, ds \right) \\
&= N^{-1} \frac{M''}{2} R^2 \left( (\ln R)^2 - \ln R + \frac{1}{2} \right) \\
&= \mathcal{O}\left(N^{-3} (\ln N^{-1})^2\right)
\end{aligned}$$

for some constant $M'' > 0$, and for $d \geq 3$, the error bound tends to

$$N^{-1}M''' \int_0^R (s^{-2\eta})^2 s^{d-1}\, ds = N^{-1}M''' \int_0^R s^{3-d}\, ds = N^{-1}\frac{M'''}{4-d}R^{4-d} = \mathcal{O}(N^{d-5})$$

for some constant $M''' > 0$. Thus we see that for $d = 2, 3$, the error decays faster than $\mathcal{O}(N^{-1})$. This justifies our procedure for setting the $W_{ii}^{\text{dis}} = 0$ and $r_k^{\text{dis}} = h\delta_{ik}$ for $d = 2, 3$.

#### 17. Further mathematical details

Here we prove various technical mathematical statements invoked in earlier sections.

**A. Linear algebra.** Here we establish some basic linear algebra facts about left eigenvectors.

**Proposition 17.1.** *Given a diagonalizable matrix $\boldsymbol{M} \in \mathbb{C}^{n\times n}$ and a diagonalization $\boldsymbol{M} = \boldsymbol{V}\boldsymbol{\Lambda}\boldsymbol{V}^{-1}$, for all $\alpha \in \{1, \cdots, n\}$ the $\alpha$-th row of $\boldsymbol{V}^{-1}$ is a left eigenvector of $\boldsymbol{M}$ with eigenvalue $\lambda_\alpha := \Lambda_{\alpha\alpha}$.*

*Proof.* This follows immediately by observing that $\boldsymbol{V}^{-1}\boldsymbol{M} = \boldsymbol{\Lambda}\boldsymbol{V}^{-1}$. □

**Proposition 17.2.** *Given a real diagonalizable matrix $\boldsymbol{M} \in \mathbb{R}^{n\times n}$ and a diagonalization $\boldsymbol{M} = \boldsymbol{V}\boldsymbol{\Lambda}\boldsymbol{V}^{-1}$, if there exists indices $\gamma, \gamma' \in \{1, \cdots, n\}$ such that $\lambda_{\gamma'} = \overline{\lambda_\gamma}$ and $\lambda_\gamma, \lambda_{\gamma'}$ have multiplicity of 1 where $\lambda_\alpha := \Lambda_{\alpha\alpha}$, then*

$$[\boldsymbol{V}]_{\alpha\gamma'}[\boldsymbol{V}^{-1}]_{\gamma'\beta} = \overline{[\boldsymbol{V}]_{\alpha\gamma}[\boldsymbol{V}^{-1}]_{\gamma\beta}}$$

*for all $\alpha, \beta \in \{1, \cdots, n\}$.*

*Proof.* For all $\alpha \in \{1, \cdots, n\}$, let $\boldsymbol{r}_\alpha, \boldsymbol{l}_\alpha^T$ denote the $\alpha$-th column of $\boldsymbol{V}$ and the $\alpha$-th row of $\boldsymbol{V}^{-1}$ respectively. Then our claim is equivalent to

$$\boldsymbol{r}_{\gamma'}\boldsymbol{l}_{\gamma'}^T = \overline{\boldsymbol{r}_\gamma \boldsymbol{l}_\gamma^T}.$$

By Proposition 17.1, $\boldsymbol{r}_\alpha$ and $\boldsymbol{l}_\alpha^T$ are also right and left eigenvectors of $\boldsymbol{V}$ with eigenvalue $\lambda_\alpha$ respectively, so $\overline{\boldsymbol{r}_\gamma}$, $\overline{\boldsymbol{l}_\gamma^T}$ are right and left eigenvectors of $\boldsymbol{V}$ with eigenvalue $\overline{\lambda_\gamma} = \lambda_{\gamma'}$ respectively. But since $\lambda_{\gamma'}$ has multiplicity of 1, eigenvectors corresponding to $\lambda_{\gamma'}$ are unique up to a scalar multiple, so $\boldsymbol{r}_{\gamma'} = c_r\overline{\boldsymbol{r}_\gamma}$ and $\boldsymbol{l}_{\gamma'}^T = c_l\overline{\boldsymbol{l}_\gamma^T}$ for some $c_r, c_l \in \mathbb{C}$. Finally, since $\boldsymbol{V}^{-1}\boldsymbol{V} = \boldsymbol{I}$, $\boldsymbol{l}_\alpha^T\boldsymbol{r}_\alpha = 1$ for all $\alpha \in \{1, \cdots, n\}$, so we have $c_l c_r = c_l c_r \overline{\boldsymbol{l}_\gamma^T \boldsymbol{r}_\gamma} = \boldsymbol{l}_{\gamma'}^T \boldsymbol{r}_{\gamma'} = 1$. Thus

$$\boldsymbol{r}_{\gamma'}\boldsymbol{l}_{\gamma'}^T = c_r c_l \overline{\boldsymbol{r}_\gamma \boldsymbol{l}_\gamma^T} = \overline{\boldsymbol{r}_\gamma \boldsymbol{l}_\gamma^T}.$$

□

**B. An integral representation of the order derivative of $K_\eta(z)$.** Here we prove an integral representation of the function

$$F_\eta(z) := \frac{\partial_\eta K_\eta(\sqrt{z})}{K_\eta(\sqrt{z})}. \qquad \text{[S158]}$$

This integral representation is later used in proving Lemma 17.13.

**Lemma 17.3.** *For all $z \in \mathbb{C} \setminus (-\infty, 0]$ and $\eta \in \mathbb{R}$,*

$$F_\eta(z) = \int_0^\infty \frac{1}{z+t}\left(\frac{1}{2} + \frac{1}{\pi}\frac{J_\eta(\sqrt{t})\partial_\eta Y_\eta(\sqrt{t}) - Y_\eta(\sqrt{t})\partial_\eta J_\eta(\sqrt{t})}{J_\eta(\sqrt{t})^2 + Y_\eta(\sqrt{t})^2}\right) dt. \qquad \text{[S159]}$$

*Proof.* The proof of this integral identity mimics that of equation (1.4) in (22). First we need to show that 1) $F_\eta(z)$ is holomorphic and 2) $F_\eta(z) = o(|z|^{-1})$ as $|z| \to 0$ and $F_\eta(z) = o(1)$ as $|z| \to \infty$. This would prove that $F_\eta(z)$ is a Stieltjes transform, so that

$$F_\eta(z) = \int_0^\infty \frac{\rho_\eta(t)}{z+t}\, dt \qquad \text{[S160]}$$

where

$$\rho_\eta(t) := \frac{1}{2\pi i}\lim_{\epsilon\to 0}\left(F_\eta(-t - i\epsilon) - F_\eta(-t + i\epsilon)\right) \qquad \text{[S161]}$$

First, since $K_\eta(z)$ is holomorphic in $\eta$ on $\mathbb{C}$ for all $z \neq 0$ and holomorphic in $z$ on $\mathbb{C} \setminus (-\infty, 0]$ for all $\eta \in \mathbb{C}$ (13, section 9.6), we have that $(\eta, z) \mapsto K_\eta(z)$ is holomorphic on $\mathbb{C} \times \mathbb{C} \setminus (-\infty, 0]$ (23, section 0.2, definition I), and thus $z \mapsto \partial_\eta K_\eta(z)$ must also be holomorphic on $\mathbb{C} \setminus (-\infty, 0]$. On the other hand, $K_\eta(\sqrt{z})$ is non-zero for all $\eta \in \mathbb{R}$ and $z \in \mathbb{C} \setminus (-\infty, 0]$ (13, section 9.6), so $F_\eta(z)$ is holomorphic on $\mathbb{C} \setminus (-\infty, 0]$. Now we show that $F_\eta(z)$ satisfies the asymptotic conditions as $|z| \to 0$ and $|z| \to \infty$. For the asymptotic condition as $|z| \to 0$, we separately treat the case of $\eta > 0$, $\eta = 0$, and $\eta < 0$. If $\eta > 0$ we have

$$K_\eta(z) \sim \frac{1}{2}\Gamma(\eta)\left(\frac{z}{2}\right)^{-\eta}$$

as $|z| \to 0$ (13, equation 9.6.9), and thus

$$F_\eta(z) \sim \psi(\eta) - \operatorname{Log}\left(\frac{\sqrt{z}}{2}\right) = o(|z|^{-1})$$

where $\psi(z)$ is the digamma function and is defined by $\psi(z) = \frac{\Gamma'(z)}{\Gamma(z)}$ (13, equation 6.3.1). If $\eta < 0$, then we use the fact that $K_\eta(z) = K_{-\eta}(z)$ for all $\eta, z \in \mathbb{C}$ (13, equation 9.6.6) to get

$$F_\eta(z) \sim \operatorname{Log}\left(\frac{\sqrt{z}}{2}\right) - \psi(-\eta) = o(|z|^{-1}).$$

Finally, if $\eta = 0$, we have $F_0(z) = 0 = o(|z|^{-1})$ since (13, equation 9.6.46)

$$\left.\frac{\partial K_\eta(z)}{\partial \eta}\right|_{\eta=0} = 0.$$

Next, for the asymptotic condition as $|z| \to \infty$, we have the asymptotic expansion

$$K_\eta(z) \sim \sqrt{\frac{\pi}{2z}}e^{-z}\left(1 + \frac{4\eta^2 - 1}{8z} + \cdots\right)$$

as $|z| \to \infty$. Thus

$$\partial_\eta K_\eta(z) \sim \sqrt{\frac{\pi}{2z}}\frac{\eta}{z}e^{-z}$$

and we have

$$F_\eta(z) \sim \frac{\eta}{\sqrt{z}} = o(1).$$

Having shown that $F_\eta(z)$ is indeed a Stieltjes transform, we now compute $\rho(t)$. By (13, equation 9.6.4), we have

$$K_\eta(\sqrt{-t - \epsilon i}) = \frac{i\pi}{2}e^{i\eta\frac{\pi}{2}}H_\eta^{(1)}(\sqrt{t + \epsilon i})$$
$$K_\eta(\sqrt{-t + \epsilon i}) = \frac{-i\pi}{2}e^{-i\eta\frac{\pi}{2}}H_\eta^{(2)}(\sqrt{t - \epsilon i})$$

so that

$$\rho(t) = \frac{1}{2} + \frac{1}{2\pi i}\left(\frac{\partial_\eta H_\eta^{(1)}(\sqrt{t})}{H_\eta^{(1)}(\sqrt{t})} - \frac{\partial_\eta H_\eta^{(2)}(\sqrt{t})}{H_\eta^{(2)}(\sqrt{t})}\right) \tag{S162}$$

But now since $H_\eta^{(2)}(x) = \overline{H}_\eta^{(1)}(x)$ for $x > 0$, and

$$H_\eta^{(1)}(x) = J_\eta(x) + iY_\eta(z),$$

we have

$$\begin{aligned}\rho(t) &= \frac{1}{2} + \frac{1}{\pi}\operatorname{Im}\left(\frac{\partial_\eta H_\eta^{(1)}(\sqrt{t})}{H_\eta^{(1)}(\sqrt{t})}\right)\\ &= \frac{1}{2} + \frac{1}{\pi}\frac{J_\eta(\sqrt{t})\partial_\eta Y_\eta(\sqrt{t}) - Y_\eta(\sqrt{t})\partial_\eta J_\eta(\sqrt{t})}{J_\eta(\sqrt{t})^2 + Y_\eta(\sqrt{t})^2}\end{aligned}$$

so we are done. □

**C. Properties of $G_d(s;\lambda)$.** Below we prove various properties about the function $G_d(s;\lambda)$.

**Lemma 17.4.** *For all $d \in R$, $\lambda > 0$, and $s > 0$, $G_d(s;\lambda)$ is real and positive.*

*Proof.* This follows immediately from the fact that $K_\eta(z)$ is real and positive for all $\eta > -1$ and $z > 0$ (13, section 9.6) and that $K_{-\eta}(z) = K_\eta(z)$ (13, equation 9.6.6). □

**Lemma 17.5.** *For all $d \in \mathbb{R}$, $\lambda \in \mathbb{C} \setminus (-\infty, 0]$, and $s > 0$, $G_d(s;\lambda)$ is non-zero.*

*Proof.* This follows immediately from the fact that $K_\eta(z)$ is non-zero for all $\eta \in \mathbb{R}$ and $z \in \mathbb{C}$ such that $|\mathrm{Arg}(z)| < \frac{\pi}{2}$ (13, section 9.6). □

**Lemma 17.6.** *For all $d \in \mathbb{R}$, $\lambda \in \mathbb{C} \setminus (-\infty, 0]$, and $s > 0$, $G_d(s;\overline{\lambda}) = \overline{G_d(s;\lambda)}$.*

*Proof.* This follows immediately from the fact that $K_\eta(\overline{z}) = \overline{K_\eta(z)}$ for all $\eta \in \mathbb{R}$ (13, equation 9.6.32). □

**Lemma 17.7.** *For all $\lambda \in \mathbb{C} \setminus (-\infty, 0]$ and $s > 0$, we have*

$$G_1(s;\lambda) = \frac{e^{-\sqrt{\lambda}s}}{2\sqrt{\lambda}} \qquad \text{[S163]}$$

$$G_3(s;\lambda) = \frac{e^{-\sqrt{\lambda}s}}{4\pi s} \qquad \text{[S164]}$$

*Proof.* This follows immediately from the fact that $K_{\pm\frac{1}{2}}(z) = \sqrt{\frac{\pi}{2z}}e^{-z}$ for all $z \in \mathbb{C}$ (13, Equation 10.2.16, 10.2.17). □

**Lemma 17.8.** *For all $d \in \mathbb{N}$, $\lambda \in \mathbb{C} \setminus (-\infty, 0]$, we have*

$$G_d(s;\lambda) \sim \begin{cases} \dfrac{1}{2\sqrt{\lambda}}, & d = 1 \\ -\dfrac{1}{2\pi}\ln(s), & d = 2 \\ \dfrac{2^{\eta-1}}{(2\pi)^{\frac{d}{2}}}\Gamma(\eta)s^{-2\eta}, & d \geq 3 \end{cases} \qquad \text{[S165]}$$

*as $s \to 0$, where $s > 0$ and $\eta = \frac{d}{2} - 1$.*

*Proof.* The case of $d = 1$ follows immediately from plugging $s = 0$ into Eq. S163. For the cases $d \geq 2$, we use the following asymptotic expressions for $K_\eta(z)$ as $z \to 0$ (13, Equations 9.6.8, 9.6.9):

$$K_\eta(z) \sim \begin{cases} -\ln(z) & \eta = 0 \\ \frac{1}{2}\Gamma(\eta)\left(\frac{1}{2}z\right)^{-\eta} & \eta > 0 \end{cases}.$$

Then

$$\lim_{s\to 0} \frac{G_2(s;\lambda)}{-\frac{1}{2\pi}\ln(s)} = \lim_{s\to 0} \frac{K_0(\sqrt{\lambda}s)}{-\ln(s)} = \lim_{s\to 0} \frac{K_0(\sqrt{\lambda}s)}{-\ln(\lambda) - \ln(s)} = \lim_{s\to 0} \frac{K_0(\sqrt{\lambda}s)}{-\ln(\sqrt{\lambda}s)} = 1$$

and for $d \geq 3$,

$$\lim_{s\to 0} \frac{G_d(s;\lambda)}{\frac{2^{\eta-1}}{(2\pi)^{\frac{d}{2}}}\Gamma(\eta)s^{-2\eta}} = \lim_{s\to 0} \frac{\sqrt{\lambda}^{\eta}K_\eta(\sqrt{\lambda}s)}{2^{\eta-1}\Gamma(\eta)s^{-\eta}} = \lim_{s\to 0} \frac{K_\eta(\sqrt{\lambda}s)}{\frac{1}{2}\Gamma(\eta)\left(\frac{1}{2}\sqrt{\lambda}s\right)^{-\eta}} = 1.$$

□

**Lemma 17.9.** *For all $d \in \mathbb{R}$ and $\lambda_0, \lambda_1 \in \mathbb{R}$ such that $0 < \lambda_0 < \lambda_1$, we have*

$$\lim_{s\to\infty} \frac{G_d(s;\lambda_1)}{G_d(s;\lambda_0)} = 0.$$

*Proof.* We will make use of the following inequality of the modified Bessel function of the second kind (24, Equation 3.2)

$$\frac{K_\eta(y)}{K_\eta(x)} > e^{x-y}$$

which holds for all $0 < x < y$ and $\eta \in \mathbb{R}$. Thus

$$\frac{G_d(s;\lambda_1)}{G_d(s;\lambda_0)} = \frac{\sqrt{\lambda_1}^\eta K_\eta(\sqrt{\lambda_1}s)}{\sqrt{\lambda_0}^\eta K_\eta(\sqrt{\lambda_0}s)} < \frac{\sqrt{\lambda_1}^\eta}{\sqrt{\lambda_0}^\eta} e^{(\sqrt{\lambda_0}-\sqrt{\lambda_1})s}.$$

Our limit then follows by noting that $\sqrt{\lambda_0} - \sqrt{\lambda_1} < 0$. □

**Lemma 17.10.** *For all $d \in \mathbb{N}$ and $\lambda_0, \lambda_1 \in \mathbb{C} \setminus (-\infty, 0]$, if $d \geq 2$ then*

$$\lim_{s\to 0} \frac{G_d(s;\lambda_1)}{G_d(s;\lambda_0)} = 1.$$

*If $d = 1$, then*

$$\frac{G_1(0;\lambda_1)}{G_1(0;\lambda_0)} = \sqrt{\frac{\lambda_0}{\lambda_1}}.$$

*Proof.* For $d = 1$, we simply plug in Eq. S163 and let $s = 0$. For $d \geq 2$, we simply note that the asymptotic expressions for $G_d(s;\lambda)$ as $s \to 0$ given by Lemma 17.8 are independent of $\lambda$. □

**Lemma 17.11.** *For all $d \in \mathbb{N}$, $s > 0$, and $\lambda_0, \lambda_1 \in \mathbb{R}$ such that $0 < \lambda_0 < \lambda_1$, we have*

$$\frac{G_d(s;\lambda_1)}{G_d(s;\lambda_0)} > 0.$$

*Furthermore, if $d > 1$, then*

$$\frac{G_d(s;\lambda_1)}{G_d(s;\lambda_0)} < 1.$$

*If $d = 1$, then*

$$\frac{G_d(s;\lambda_1)}{G_d(s;\lambda_0)} \leq \sqrt{\frac{\lambda_0}{\lambda_1}}.$$

*Proof.* For the first claim, note that since $K_\eta(x) > 0$ for all $x > 0$, $\eta \in \mathbb{R}$ (13, Section 9.6), we have $G_d(s;\lambda) > 0$ for all $s > 0$, $\lambda > 0$, and positive integers $d$. Thus

$$\frac{G_d(s;\lambda_1)}{G_d(s;\lambda_0)} > 0.$$

For the second claim, we need an inequality of the modified Bessel function of the second kind (24, Equation 3.4)

$$\frac{y^\eta K_\eta(y)}{x^\eta K_\eta(x)} < 1$$

which holds for all $0 < x < y$, $\eta > -\frac{1}{2}$. Thus for all $s > 0$, $d > 1$, $0 < \lambda_0 < \lambda_1$ we have

$$\frac{G_d(s;\lambda_1)}{G_d(s;\lambda_0)} = \frac{(\sqrt{\lambda_1}s)^\eta K_\eta(\sqrt{\lambda_1}s)}{(\sqrt{\lambda_0}s)^\eta K_\eta(\sqrt{\lambda_0}s)} < 1.$$

For the case $d = 1$, we use the fact that $K_{-\frac{1}{2}}(z) = \sqrt{\frac{\pi}{2z}}e^{-z}$ (13, Equation 10.2.16, 10.2.17) and get

$$\frac{G_d(s;\lambda_1)}{G_d(s;\lambda_0)} = \sqrt{\frac{\lambda_0}{\lambda_1}} e^{-(\sqrt{\lambda_1}-\sqrt{\lambda_0})s} \leq \sqrt{\frac{\lambda_0}{\lambda_1}}.$$

□

**Lemma 17.12.** *For all $d \in \mathbb{R}$, $s > 0$, and $\lambda_0, \lambda_1 \in \mathbb{C} \setminus (-\infty, 0]$, if $\lambda_0, \lambda_1$ are real and $0 < \lambda_0 < \lambda_1$, then*

$$\frac{G_{d+2}(s;\lambda_1)}{G_d(s;\lambda_1)} > \frac{G_{d+2}(s;\lambda_0)}{G_d(s;\lambda_0)}. \quad \text{[S166]}$$

*On the other hand, if $\lambda_0 = \overline{\lambda_1}$ and $\mathrm{Im}(\lambda_1) > 0$, then*

$$\mathrm{Im}\left(\frac{G_{d+2}(s;\lambda_1)}{G_d(s;\lambda_1)}\right) > 0 > \mathrm{Im}\left(\frac{G_{d+2}(s;\lambda_0)}{G_d(s;\lambda_0)}\right). \quad \text{[S167]}$$

*Note that these equations are well-defined since $G_d(s;\lambda)$ is non-zero for all $\lambda \in \mathbb{C} \setminus (-\infty, 0]$ (Lemma 17.5).*

*Proof.* **Case $0 < \lambda_0 < \lambda_1$:** Plugging in the definition of $G_d(s;\lambda)$ (Eq. S10), we get the equivalent inequality in terms of $K_\eta$ as

$$\frac{\sqrt{\lambda_1} s K_{\eta+1}(\sqrt{\lambda_1} s)}{K_\eta(\sqrt{\lambda_1} s)} > \frac{\sqrt{\lambda_0} s K_{\eta+1}(\sqrt{\lambda_0} s)}{K_\eta(\sqrt{\lambda_0} s)}$$

where $\eta = \frac{d}{2} - 1$. But for all $\eta \in \mathbb{R}$, $\beta > 0$, the function

$$f_{\eta,\beta}(s) = \frac{s^\beta K_{\eta+\beta}(s)}{K_\eta(s)} \quad \text{[S168]}$$

is strictly monotonically increasing on $(0, \infty)$ (25, Lemma 2.6). Since $\sqrt{\lambda_1} s > \sqrt{\lambda_0} s$, our inequality follows.

**Case $d \geq 0$, $\lambda_0 = \overline{\lambda_1}$:** By Lemma 17.6, we see that it suffices to show

$$\mathrm{Im}\left(\frac{G_{d+2}(s;\lambda_1)}{G_d(s;\lambda_1)}\right) > 0.$$

Plugging in the definition of $G_d(s;\lambda)$, we get the equivalent inequality

$$\mathrm{Im}\left(\frac{\sqrt{\lambda_1} s K_{\eta+1}(\sqrt{\lambda_1} s)}{K_\eta(\sqrt{\lambda_1} s)}\right) > 0. \quad \text{[S169]}$$

Since for all $\mathrm{Re}(z) > 0$ and $\eta \geq -1$ we have the following integral representation (22, Equation 1.4)

$$\frac{K_\eta(z)}{z K_{\eta+1}(z)} = \frac{2}{\pi^2} \int_0^\infty \frac{t^{-1}}{z^2 + t} \left(J_{\eta+1}^2(\sqrt{t}) + Y_{\eta+1}^2(\sqrt{t})\right)^{-1} dt,$$

we see that $\mathrm{Im}\left(\frac{K_\eta(z)}{zK_{\eta+1}(z)}\right) < 0$ if $\mathrm{Im}((z^2+t)^{-1}) < 0$ for all $t > 0$. But $\mathrm{Im}(z) > 0$ if and only if $\mathrm{Im}(z^{-1}) < 0$, so $\mathrm{Im}\left(\frac{zK_{\eta+1}(z)}{K_\eta(z)}\right) > 0$ if $\mathrm{Im}(z^2) > 0$. Plugging in $z = \sqrt{\lambda_1} s$, our inequality thus follows from our assumption that $\mathrm{Im}(\lambda_1) > 0$.

**Case $d < 0$, $\lambda_0 = \overline{\lambda_1}$:** Since $d < 0$, we have $\eta < -1$, so $-\eta = |\eta|$ and $-(\eta+1) = -\eta - 1 = |\eta| - 1$. Moreover, since $K_\eta = K_{-\eta}$ (13, equation 9.6.6), Eq. S169 is thus equivalent to

$$\mathrm{Im}\left(\frac{\sqrt{\lambda_1} s K_{|\eta|-1}(\sqrt{\lambda_1} s)}{K_{|\eta|}(\sqrt{\lambda_1} s)}\right) > 0.$$

Since for all $\mathrm{Re}(z) > 0$ and $\eta' \geq 0$ we have the following integral representation (22, Equation 1.5)

$$\frac{K_{\eta'+1}(z)}{z K_{\eta'}(z)} = \frac{2\eta'}{z^2} + \frac{2}{\pi^2} \int_0^\infty \frac{t^{-1}}{z^2 + t} \left(J_{\eta'}^2(\sqrt{t}) + Y_{\eta'}^2(\sqrt{t})\right)^{-1} dt,$$

we see that $\mathrm{Im}\left(\frac{K_{\eta'+1}(z)}{zK_{\eta'}(z)}\right) < 0$ if $\mathrm{Im}((z^2+t)^{-1}) < 0$ for all $t > 0$. But $\mathrm{Im}(z) > 0$ if and only if $\mathrm{Im}(z^{-1}) < 0$, so $\mathrm{Im}\left(\frac{zK_{\eta'}(z)}{K_{\eta'+1}(z)}\right) > 0$ if $\mathrm{Im}(z^2) > 0$. Plugging in $\eta' = |\eta| - 1$ and $z = \sqrt{\lambda_1} s$, our inequality thus follows from our assumption that $\mathrm{Im}(\lambda_1) > 0$. □

**Lemma 17.13.** *For all $d \in \mathbb{R}$, $s > 0$, and $\lambda_0, \lambda_1 \in \mathbb{C} \setminus (-\infty, 0]$, if $\lambda_0, \lambda_1$ are real and $0 < \lambda_0 < \lambda_1$, then*

$$\frac{\partial_d G_d(s;\lambda_1)}{G_d(s;\lambda_1)} > \frac{\partial_d G_d(s;\lambda_0)}{G_d(s;\lambda_0)}. \qquad \text{[S170]}$$

*On the other hand, if $\lambda_0 = \overline{\lambda_1}$ and $\mathrm{Im}(\lambda_1) > 0$, then*

$$\mathrm{Im}\left(\frac{\partial_d G_d(s;\lambda_1)}{G_d(s;\lambda_1)}\right) > 0 > \mathrm{Im}\left(\frac{\partial_d G_d(s;\lambda_0)}{G_d(s;\lambda_0)}\right). \qquad \text{[S171]}$$

*Note that these equations are well-defined since $G_d(s;\lambda)$ is non-zero for all $\lambda \in \mathbb{C} \setminus (-\infty, 0]$ (Lemma 17.5).*

*Proof.* First, from the definition of $G_d$ (Eq. S10), we have

$$\begin{aligned}\partial_d G_d(s;\lambda) &= \frac{1}{2\pi}\partial_d\left(\left(\frac{\sqrt{\lambda}}{2\pi s}\right)^{\eta} K_\eta(\sqrt{\lambda}s)\right) \\ &= \frac{1}{4\pi}\left(\frac{\sqrt{\lambda}}{2\pi s}\right)^{\eta}\left(\mathrm{Log}\left(\frac{\sqrt{\lambda}}{2\pi s}\right) K_\eta(\sqrt{\lambda}s) + \partial_\eta K_\eta(\sqrt{\lambda}s)\right)\end{aligned}$$

so that

$$\frac{\partial_d G_d(s;\lambda)}{G_d(s;\lambda)} = \frac{1}{2}\left(\mathrm{Log}\left(\sqrt{\lambda}\right) - \ln(2\pi s) + \frac{\partial_\eta K_\eta(\sqrt{\lambda}s)}{K_\eta(\sqrt{\lambda}s)}\right).$$

Plugging in the integral representation S159 proved in Lemma 17.3, we thus have

$$\frac{\partial_d G_d(s;\lambda)}{G_d(s;\lambda)} = \frac{1}{2}\left(\mathrm{Log}\left(\sqrt{\lambda}\right) - \ln(2\pi s) + \int_0^\infty \frac{1}{\lambda s^2 + t}\left(\frac{1}{2} + \frac{1}{\pi}\frac{J_\eta(\sqrt{t})\partial_\eta Y_\eta(\sqrt{t}) - Y_\eta(\sqrt{t})\partial_\eta J_\eta(\sqrt{t})}{J_\eta(\sqrt{t})^2 + Y_\eta(\sqrt{t})^2}\right) dt\right). \qquad \text{[S172]}$$

Next, we note that the Wronskian term inside the integral has, in turn, its own integral representation (26, Equation 6) (27, p. 444):

$$J_\eta(x)\partial_\eta Y_\eta(x) - Y_\eta(x)\partial_\eta J_\eta(x) = -\frac{4}{\pi}\int_0^\infty K_0(2x\sinh t)e^{-2\eta t}\,dt$$

for all $\eta \in \mathbb{C}$ and $x > 0$. In particular, since $K_0(x)$ is real and positive for all $x > 0$ (13, section 9.6), this implies that

$$\frac{J_\eta(\sqrt{t})\partial_\eta Y_\eta(\sqrt{t}) - Y_\eta(\sqrt{t})\partial_\eta J_\eta(\sqrt{t})}{J_\eta(\sqrt{t})^2 + Y_\eta(\sqrt{t})^2} < 0$$

for all $t > 0$. Now we prove the first inequality (Eq. S170). Taking the partial derivative of Eq. S172, we have, for $\lambda > 0$,

$$\begin{aligned}\partial_\lambda\left(\frac{\partial_d G_d(s;\lambda)}{G_d(s;\lambda)}\right) &= \frac{1}{2}\left(\frac{1}{2\lambda} - s^2\int_0^\infty \frac{1}{(\lambda s^2 + t)^2}\left(\frac{1}{2} + \frac{1}{\pi}\frac{J_\eta(\sqrt{t})\partial_\eta Y_\eta(\sqrt{t}) - Y_\eta(\sqrt{t})\partial_\eta J_\eta(\sqrt{t})}{J_\eta(\sqrt{t})^2 + Y_\eta(\sqrt{t})^2}\right) dt\right) \\ &= -\frac{s^2}{2\pi}\int_0^\infty \frac{1}{(\lambda s^2 + t)^2}\frac{J_\eta(\sqrt{t})\partial_\eta Y_\eta(\sqrt{t}) - Y_\eta(\sqrt{t})\partial_\eta J_\eta(\sqrt{t})}{J_\eta(\sqrt{t})^2 + Y_\eta(\sqrt{t})^2}\,dt \\ &> 0.\end{aligned}$$

For the second inequality (Eq. S171), we have

$$\mathrm{Im}\left(\frac{\partial_d G_d(s;\lambda)}{G_d(s;\lambda)}\right) = \frac{1}{2}\left(\frac{1}{2}\mathrm{Arg}(\lambda) - \mathrm{Im}(\lambda)s^2\int_0^\infty \frac{1}{|\lambda s^2 + t|^2}\left(\frac{1}{2} + \frac{1}{\pi}\frac{J_\eta(\sqrt{t})\partial_\eta Y_\eta(\sqrt{t}) - Y_\eta(\sqrt{t})\partial_\eta J_\eta(\sqrt{t})}{J_\eta(\sqrt{t})^2 + Y_\eta(\sqrt{t})^2}\right) dt\right)$$

But for all $z = x + iy$ such that $x, y \in \mathbb{R}$ and $y > 0$, we have

$$\int_0^\infty \frac{1}{|z+t|^2}\,dt = \int_0^\infty \frac{1}{(t+x)^2 + y^2}\,dt = \frac{1}{y}\int_{\frac{x}{y}}^\infty \frac{1}{1+u^2}\,du = \frac{1}{y}\left(\frac{\pi}{2} - \arctan\left(\frac{x}{y}\right)\right) = \frac{\mathrm{Arg}(z)}{\mathrm{Im}(z)}.$$

 

Thus if $\mathrm{Im}(\lambda) > 0$, then

$$\begin{aligned}\mathrm{Im}\left(\frac{\partial_d G_d(s;\lambda)}{G_d(s;\lambda)}\right) &= -\frac{\mathrm{Im}(\lambda)s^2}{2\pi}\int_0^\infty \frac{1}{|\lambda s^2+t|^2}\frac{J_\eta(\sqrt{t})\partial_\eta Y_\eta(\sqrt{t}) - Y_\eta(\sqrt{t})\partial_\eta J_\eta(\sqrt{t})}{J_\eta(\sqrt{t})^2+Y_\eta(\sqrt{t})^2}\,dt \\ &> 0.\end{aligned}$$

□

**Lemma 17.14.** *For all $d \in \mathbb{R}$, $s > 0$, and $\lambda \in \mathbb{C}\setminus(-\infty, 0]$, we have*

$$\partial_\lambda G_d(s;\lambda) = -\frac{1}{4\pi}G_{d-2}(s;\lambda)$$

*Proof.* Noting that (13, Equation 9.6.28)

$$\begin{aligned}\frac{1}{z}\frac{\mathrm{d}}{\mathrm{d}z}\left(z^\eta e^{\eta\pi i}K_\eta(z)\right) &= z^{\eta-1}e^{(\eta-1)\pi i}K_{\eta-1}(z) \\ \frac{\mathrm{d}}{\mathrm{d}z}\left(z^\eta K_\eta(z)\right) &= -z^\eta K_{\eta-1}(z)\end{aligned}$$

we have

$$\begin{aligned}\partial_\lambda G_d(s;\lambda) &= \frac{1}{(2\pi)^{\frac{d}{2}}}s^{-2\eta}\partial_\lambda[(\sqrt{\lambda}s)^\eta K_\eta(\sqrt{\lambda}s)] \\ &= -\frac{1}{(2\pi)^{\frac{d}{2}}}s^{-2\eta}(\sqrt{\lambda}s)^\eta K_{\eta-1}(\sqrt{\lambda}s)\left(\frac{1}{2\sqrt{\lambda}}s\right) \\ &= -\frac{1}{2(2\pi)^{\frac{d}{2}}}\left(\frac{\sqrt{\lambda}}{s}\right)^{\eta-1}K_{\eta-1}(\sqrt{\lambda}s) \\ &= -\frac{1}{4\pi}G_{d-2}(s;\lambda)\end{aligned}$$

□

**Lemma 17.15.** *For all $d \in \mathbb{R}$, $s > 0$, and $\lambda \in \mathbb{C}\setminus(-\infty, 0]$, we have*

$$\partial_s G_d(s;\lambda) = -2\pi r G_{d+2}(s;\lambda)$$

*Proof.* Noting that (13, Equation 9.6.28)

$$\begin{aligned}\frac{1}{z}\frac{\mathrm{d}}{\mathrm{d}z}\left(z^{-\eta} e^{\eta\pi i}K_\eta(z)\right) &= z^{-\eta-1}e^{(\eta+1)\pi i}K_{\eta+1}(z) \\ \frac{\mathrm{d}}{\mathrm{d}z}\left(z^{-\eta} K_\eta(z)\right) &= -z^{-\eta} K_{\eta+1}(z)\end{aligned}$$

we have

$$\begin{aligned}\partial_s G_d(s;\lambda) &= \frac{1}{(2\pi)^{\frac{d}{2}}}\sqrt{\lambda}^{2\eta}\partial_s[(\sqrt{\lambda}s)^{-\eta}K_\eta(\sqrt{\lambda}s)] \\ &= -\frac{1}{(2\pi)^{\frac{d}{2}}}\sqrt{\lambda}^{2\eta}(\sqrt{\lambda}s)^{-\eta}K_{\eta+1}(\sqrt{\lambda}s)\sqrt{\lambda} \\ &= -\frac{1}{(2\pi)^{\frac{d}{2}}}s\left(\frac{\sqrt{\lambda}}{s}\right)^{\eta+1}K_{\eta+1}(\sqrt{\lambda}s) \\ &= -2\pi r G_{d+2}(s;\lambda)\end{aligned}$$

□

**Lemma 17.16.** *For all $d \in \mathbb{R}$ and $\lambda_0, \lambda_1 \in \mathbb{R}$ such that $0 < \lambda_0 < \lambda_1$, the function $s \to \frac{G_d(s;\lambda_1)}{G_d(s;\lambda_0)}$ is strictly monotonically decreasing on $(0, \infty)$.*

*Proof.* By Lemma 17.15,

$$\frac{\mathrm{d}}{\mathrm{d}s}\frac{G_d(s;\lambda_1)}{G_d(s;\lambda_0)} = -2\pi s\frac{G_d(s;\lambda_0)G_{d+2}(s;\lambda_1) - G_d(s;\lambda_1)G_{d+2}(s;\lambda_0)}{G_d(s;\lambda_0)^2},$$

and since the numerator is positive by Lemma 17.12, the derivative is always negative, and thus the function is strictly monotonically decreasing. □

**Lemma 17.17.** *For all $d \in \mathbb{R}$, $s > 0$, and $\lambda_0, \lambda_1 \in \mathbb{C} \setminus (-\infty, 0]$, we have*

$$2\pi s\left(G_d(s;\lambda_0)G_{d+2}(s;\lambda_1) - G_d(s;\lambda_1)G_{d+2}(s;\lambda_0)\right) = (2\pi s)^{-1}\left(\lambda_1 G_d(s;\lambda_0)G_{d-2}(s;\lambda_1) - \lambda_0 G_d(s;\lambda_1)G_{d-2}(s;\lambda_0)\right) \quad \text{[S173]}$$

*Proof.* Let $\eta = \frac{d}{2} - 1$. By the recurrence relation (13, Equation 9.6.26)

$$e^{(\eta-1)\pi i}K_{\eta-1}(z) = e^{(\eta+1)\pi i}K_{\eta+1}(z) + \frac{2\eta}{z}e^{\eta\pi i}K_\eta(z)$$
$$K_{\eta+1}(z) = K_{\eta-1}(z) + \frac{2\eta}{z}K_\eta(z),$$

we have

$$\begin{aligned}
&\sqrt{\lambda_1}K_\eta(\sqrt{\lambda_0}s)K_{\eta+1}(\sqrt{\lambda_1}s) - \sqrt{\lambda_0}K_\eta(\sqrt{\lambda_1}s)K_{\eta+1}(\sqrt{\lambda_0}s)\\
=&\sqrt{\lambda_1}K_\eta(\sqrt{\lambda_0}s)\left(K_{\eta-1}(\sqrt{\lambda_1}s) + \frac{2\eta}{\sqrt{\lambda_1}s}K_\eta(\sqrt{\lambda_1}s)\right) - \sqrt{\lambda_0}K_\eta(\sqrt{\lambda_1}s)\left(K_{\eta-1}(\sqrt{\lambda_0}s) + \frac{2\eta}{\sqrt{\lambda_0}s}K_\eta(\sqrt{\lambda_0}s)\right)\\
=&\sqrt{\lambda_1}K_\eta(\sqrt{\lambda_0}s)K_{\eta-1}(\sqrt{\lambda_1}s) - \sqrt{\lambda_0}K_\eta(\sqrt{\lambda_1}s)K_{\eta-1}(\sqrt{\lambda_0}s).
\end{aligned}$$

Multiplying both sides by $\frac{1}{(2\pi)^{\frac{d}{2}}}\left(\frac{\sqrt{\lambda_0}}{s}\right)^\eta \frac{1}{(2\pi)^{\frac{d}{2}}}\left(\frac{\sqrt{\lambda_1}}{s}\right)^\eta$ then yields Eq. S173. □

**D. Number of zero crossings.** Here we prove statements about the number of zero crossings of the perturbation response as a function of distance.

**Theorem 17.18.** *For all $d \in \mathbb{R}$, $\lambda \in \mathbb{C} \setminus \mathbb{R}$, and $c \in \mathbb{C} \setminus \{0\}$, the function $s \mapsto \mathrm{Re}(cG_d(s;\lambda))$ has infinitely many zero-crossings on $(0, \infty)$.*

*Proof.* First, note that it is sufficient to show that the function $s \mapsto \mathrm{Re}(cK_\eta(ks))$ has infinitely many zero-crossings on $(0, \infty)$, where $k = \sqrt{\lambda}$, since

$$\mathrm{Re}(cG_d(s;\lambda)) = s^{-\eta}\mathrm{Re}\left(\frac{c}{(2\pi)^{\frac{d}{2}}}k^\eta K_\eta(ks)\right) = s^{-\eta}\mathrm{Re}(c'K_\eta(ks))$$

for some $c' \in \mathbb{C}$, and the extra factor of $s^{-\eta}$ does not affect the number of zero-crossings. Now, the modified Bessel function of the second kind $K_\eta(z)$ has the following asymptotic expansion as $|z| \to \infty$ (13, Equation 9.7.2)

$$K_\eta(z) \sim \sqrt{\frac{\pi}{2z}}e^{-z}\left(1 + \frac{4\eta^2 - 1}{8z} + \cdots\right) \quad \text{[S174]}$$

This implies

$$K_\eta(z) - \sqrt{\frac{\pi}{2z}}e^{-z} = \mathcal{O}\left(\frac{e^{-z}}{z^{\frac{3}{2}}}\right),$$

so there exists $M > 0$ and $s_0 > 0$ such that

$$\left|cK_\eta(ks) - c\sqrt{\frac{\pi}{2ks}}e^{-ks}\right| \le M\frac{|e^{-ks}|}{s^{\frac{3}{2}}}$$

 

for all $s > s_0$. But since $|\mathrm{Re}(f(z))| \leq |f(z)|$ for all $f$, we also have

$$\left|\mathrm{Re}(cK_\eta(ks)) - \mathrm{Re}\left(c\sqrt{\frac{\pi}{2ks}}e^{-ks}\right)\right| \leq M\frac{|e^{-ks}|}{s^{\frac{3}{2}}}$$

Now let $k^R$ and $k^I$ be the real and imaginary parts of $k$. Then

$$\begin{aligned}\mathrm{Re}\left(c\sqrt{\frac{\pi}{2ks}}e^{-ks}\right) &= \sqrt{\frac{\pi}{2s}}\mathrm{Re}\left(\left|\frac{c}{\sqrt{k}}\right|e^{-k^R s - i(k^I s - \mathrm{Arg}(\frac{c}{\sqrt{k}}))}\right) \\ &= \sqrt{\frac{\pi}{2s}}\left|\frac{c}{\sqrt{k}}\right|e^{-k^R s}\cos\left(k^I s - \mathrm{Arg}\left(\frac{c}{\sqrt{k}}\right)\right)\end{aligned}$$

which is clearly a function with infinitely many zero crossings. Also noting that $|e^{-ks}| = e^{-k^R s}$, we thus have

$$\left|\rho\sqrt{s}e^{k^R s}\mathrm{Re}(cK_\eta(ks)) - \cos\left(k^I s - \theta\right)\right| \leq \frac{M\rho}{s}$$

where $\rho = \sqrt{\frac{2}{\pi}}\left|\frac{\sqrt{k}}{c}\right|$ and $\theta = \mathrm{Arg}\left(\frac{c}{\sqrt{k}}\right)$. Now fix some $s_1 > s_0$ such that

$$\frac{M\rho}{s_1} \leq \frac{1}{2}$$

Then for all $s \geq s_1$, we have

$$\left|\rho\sqrt{s}e^{k^R s}\mathrm{Re}(cK_\eta(ks)) - \cos\left(k^I s - \theta\right)\right| \leq \frac{1}{2}$$

Now fix $s_1^- \geq s_1$ such that

$$\cos(k^I s_1^- - \theta) = -1$$

and define

$$\begin{aligned}s_{n+1}^- &= s_1^- + \frac{2n\pi}{k^I} \\ s_{n+1}^+ &= s_1^- + \frac{(2n+1)\pi}{k^I}\end{aligned}$$

for all $n \in \mathbb{N}$. Clearly, $\cos(k^I s_n^\pm - \theta) = \pm 1$. Thus we have

$$\begin{aligned}\rho\sqrt{s_n^-}e^{k^R s_n^-}\mathrm{Re}(cK_\eta(ks_n^-)) &\leq -\frac{1}{2} \\ \rho\sqrt{s_n^+}e^{k^R s_n^+}\mathrm{Re}(cK_\eta(ks_n^+)) &\geq \frac{1}{2}\end{aligned}$$

for all $n \in \mathbb{N}$, so by the intermediate value theorem, the function

$$\rho\sqrt{s}e^{k^R s}\mathrm{Re}(cK_\eta(ks))$$

has a zero crossing between $s_n^-$ and $s_n^+$ for all $n \in \mathbb{N}$, and thus $\mathrm{Re}(cK_\eta(ks))$ has infinitely many zero crossings. □

**Theorem 17.19.** *For all $d \in \mathbb{N}$ and $c_0, c_1, \lambda_0, \lambda_1 \in \mathbb{R}$ such that $0 < \lambda_0 < \lambda_1$, the function $l : (0, \infty) \to \mathbb{R}$ given by*

$$l(s) = c_0 G_d(s; \lambda_0) + c_1 G_d(s; \lambda_1) \tag{S175}$$

*has at most one root and at most one critical point. Given some $s_0 > 0$, $s_0$ is a root of $l$ if and only if $s_0$ is zero crossing of $l$ (in the sense that $l(s_0) = 0$, $l'(s_0) \neq 0$). If both the root and the critical point exists, then the critical point must be greater than the root.*

*Suppose further that $c_1 \neq 0$. Then for $d \geq 2$, the zero crossing exists if and only if $-\frac{c_0}{c_1} \in (0, 1)$, while for $d = 1$, the zero crossing exists if and only if $-\frac{c_0}{c_1} \in (0, \sqrt{\frac{\lambda_0}{\lambda_1}})$.*

*Proof.* By Lemma 17.4, $G_d(s;\lambda) > 0$ for all $s > 0$, so the roots of $l$ are given by

$$c_0 + c_1 \frac{G_d(s;\lambda_1)}{G_d(s;\lambda_0)} = 0. \quad \text{[S176]}$$

Similarly, by Lemma 17.15, the critical points of $l$ are given by

$$c_0 + c_1 \frac{G_{d+2}(s;\lambda_1)}{G_{d+2}(s;\lambda_0)} = 0. \quad \text{[S177]}$$

By Lemma 17.16, the left hand side of both equations are strictly monotonically decreasing functions of $s$ on $(0,\infty)$, hence $l$ has at most one root and at most one critical point. Furthermore, by Lemma 17.12, the L.H.S. of Eq. S177 is always greater than the L.H.S. of Eq. S176, so no roots of $l$ can also be a critical point, i.e. all roots of $l$ are also zero crossings. But by definition, zero crossings are also roots, so the roots and zero crossings of $l$ are equivalent. In fact it also shows that if both the root and the critical point exists, then the critical point must be greater than the root.

Now suppose $c_1 \neq 0$. Let $c = -\frac{c_0}{c_1}$. Then Eq. S176 can be rewritten as

$$\frac{G_d(s;\lambda_1)}{G_d(s;\lambda_0)} = c.$$

Consider the case $d \geq 2$. By Lemma 17.9 and 17.10, we have $\lim_{s\to 0} \frac{G_d(s;\lambda_1)}{G_d(s;\lambda_0)} = 1$ and $\lim_{s\to\infty} \frac{G_d(s;\lambda_1)}{G_d(s;\lambda_0)} = 0$. Thus, if $c \in (0,1)$, then a root (and hence a zero crossing) exists by the intermediate value theorem. Conversely, if there exists a zero crossing, then $c = \frac{G_d(s;\lambda_1)}{G_d(s;\lambda_0)}$ for some $s > 0$. But by Lemma 17.11, $\frac{G_d(s;\lambda_1)}{G_d(s;\lambda_0)} \in (0,1)$ for all $s > 0$, so $c \in (0,1)$. Thus, $l$ has a zero crossing if and only if $c \in (0,1)$. But since there is at most zero crossing, there is exactly one zero crossing if and only if $c \in (0,1)$. One can repeat the same analysis for the case $d = 1$ and show that in this case the condition is $c \in (0, \sqrt{\frac{\lambda_0}{\lambda_1}})$. □

**Theorem 17.20.** *Given $c \in \mathbb{C}$ and $\lambda_0, \lambda_1 \in \mathbb{C} \setminus (-\infty, 0]$ such that $\lambda_0 \neq \lambda_1$ and that either $\lambda_0, \lambda_1 \in \mathbb{R}$ or $\lambda_1 = \overline{\lambda}_0$, define the equation*

$$f(d,s) := \frac{G_d(s;\lambda_1)}{G_d(s;\lambda_0)} - c.$$

*Suppose there exists $d_0 \in \mathbb{R}$ and $s_0 > 0$ such that $f(d_0, s_0) = 0$. Then there exists $U \subset \mathbb{R}$ such that $d_0 \in U$ and a function $h : U \to \mathbb{C}$ such that $h(d_0) = s_0$ and $f(d, h(d)) = 0$ for all $d \in U$. In other words, we have a function $h$ which parametrizes the root $s_0$ in a small neighborhood of $d_0$. Furthermore, $h'(d) > 0$, so the root increases with $d$.*

*Proof.* Since $K_\eta(z)$ is holomorphic in $\eta$ for all $z \neq 0$ and holomorphic in $z$ on $\mathbb{C} \setminus (-\infty, 0]$ for all $\eta$ (13, section 9.6), the function $(\eta, z) \mapsto K_\eta(z)$ is holomorphic on $\mathbb{C} \times (\mathbb{C} \setminus (-\infty, 0])$ (23, chapter 0.2, defintion I). Thus there exists neighborhoods $\tilde{U}, \tilde{V} \subset \mathbb{C}$ containing $d_0, s_0$ such that $f$ is holomorphic on $\tilde{U} \times \tilde{V}$. By Lemma 17.15, we have

$$\partial_s f(d,s) = -2\pi s \frac{G_d(s;\lambda_0)G_{d+2}(s;\lambda_1) - G_d(s;\lambda_1)G_{d+2}(s;\lambda_0)}{G_d(s;\lambda_0)^2} = -2\pi s \frac{G_d(s;\lambda_1)}{G_d(s;\lambda_0)} \left( \frac{G_{d+2}(s;\lambda_1)}{G_d(s;\lambda_1)} - \frac{G_{d+2}(s;\lambda_0)}{G_d(s;\lambda_0)} \right).$$

But the last expression is nonzero for all $d \in \mathbb{R}$ and $s > 0$ by Lemma 17.5 and Lemma 17.12, so by the holomorphic implicit function theorem (21, chapter I, theorem 7.6), there exists $U \subset \tilde{U}$ and $h : U \to \mathbb{C}$ such that $h(d_0) = s_0$ and $f(d, h(d)) = 0$ for all $d \in U$, and that

$$h'(d) = -\frac{\partial_d f(d, h(d))}{\partial_s f(d, h(d))}.$$

Now since

$$\partial_d f(d,s) = \frac{G_d(s;\lambda_0)\partial_d G_d(s;\lambda_1) - G_d(s;\lambda_1)\partial_d G_d(s;\lambda_0)}{G_d(s;\lambda_0)^2} = \frac{G_d(s;\lambda_1)}{G_d(s;\lambda_0)} \left( \frac{\partial_d G_d(s;\lambda_1)}{G_d(s;\lambda_1)} - \frac{\partial_d G_d(s;\lambda_0)}{G_d(s;\lambda_0)} \right)$$

we have

$$-\frac{\partial_d f(d,s)}{\partial_s f(d,s)} = \frac{1}{2\pi s} \frac{\frac{\partial_d G_d(s;\lambda_1)}{G_d(s;\lambda_1)} - \frac{\partial_d G_d(s;\lambda_0)}{G_d(s;\lambda_0)}}{\frac{G_{d+2}(s;\lambda_1)}{G_d(s;\lambda_1)} - \frac{G_{d+2}(s;\lambda_0)}{G_d(s;\lambda_0)}}.$$

But by Lemma 17.12 and Lemma 17.13 (and noting that $G_d(s;\lambda_1) = \overline{G_d(s;\lambda_0)}$ in the case of $\lambda_1 = \overline{\lambda}_0$ by Lemma 17.6), the denominator and the numerator always have the same sign. Thus $h'(d) > 0$ for all $\lambda \in \mathbb{C} \setminus \mathbb{R}$. □

 

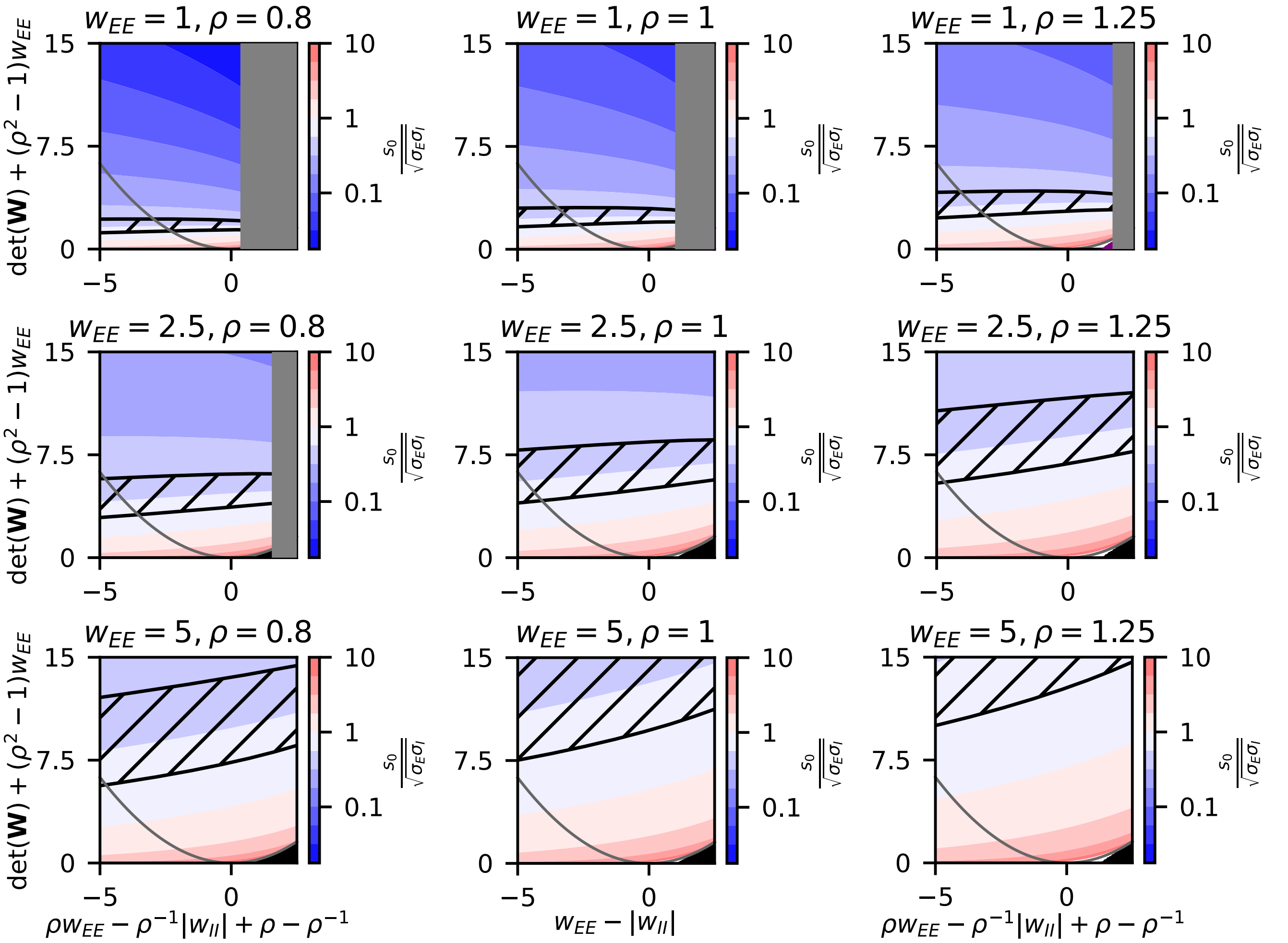


**Fig. S1.** Distance to the first zero-crossing, $s_0$, as a fraction of $\sqrt{\sigma_E \sigma_I}$, for the response of excitatory neurons to single-cell excitatory neuron perturbation in an E-I network with 2 spatial dimensions, for different combinations of $w_{\mathrm{EE}}$ and $\rho = \frac{\sigma_I}{\sigma_E}$. Hatched region indicates 95% confidence interval of $\frac{s_0}{\sqrt{\sigma_E \sigma_I}}$ estimated from experimental data. Networks in the phase region shaded in black are dynamically unstable. Phase region shaded in grey indicates an invalid set of model parameters. For example, in the middle figure of the first row, $x = 2$ is inside the greyed out region because that would require $w_{\mathrm{EE}} - |w_{\mathrm{II}}| = 2$, but since $w_{\mathrm{EE}} = 1$, that implies $|w_{\mathrm{II}}| = -1$, which is impossible.

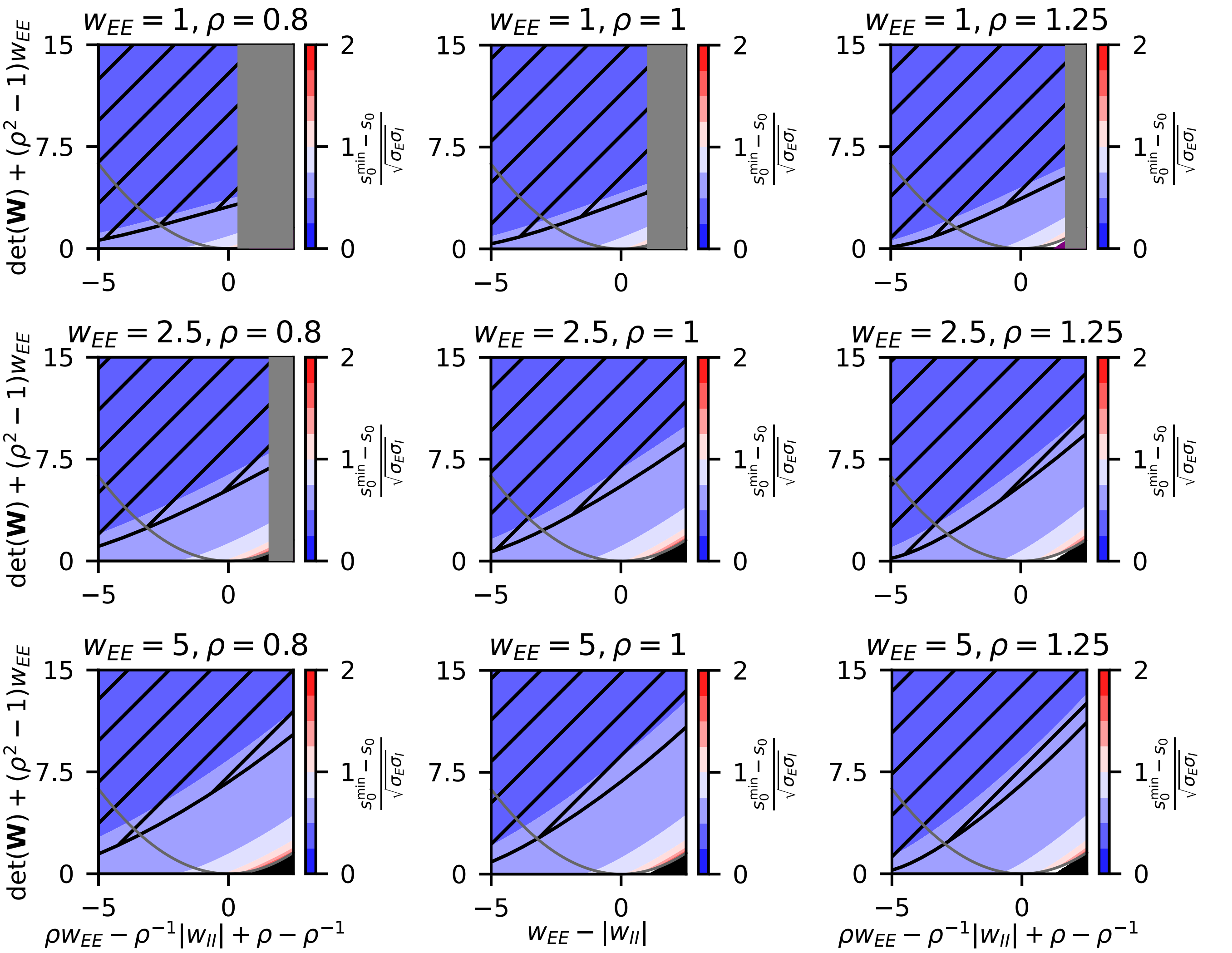


**Fig. S2.** Similar to Figure S1, but the distance from the first zero crossing $s_0$ to the first minimum $r_0^{\min}$ as a fraction of $\sqrt{\sigma_E\sigma_I}$ is plotted

 

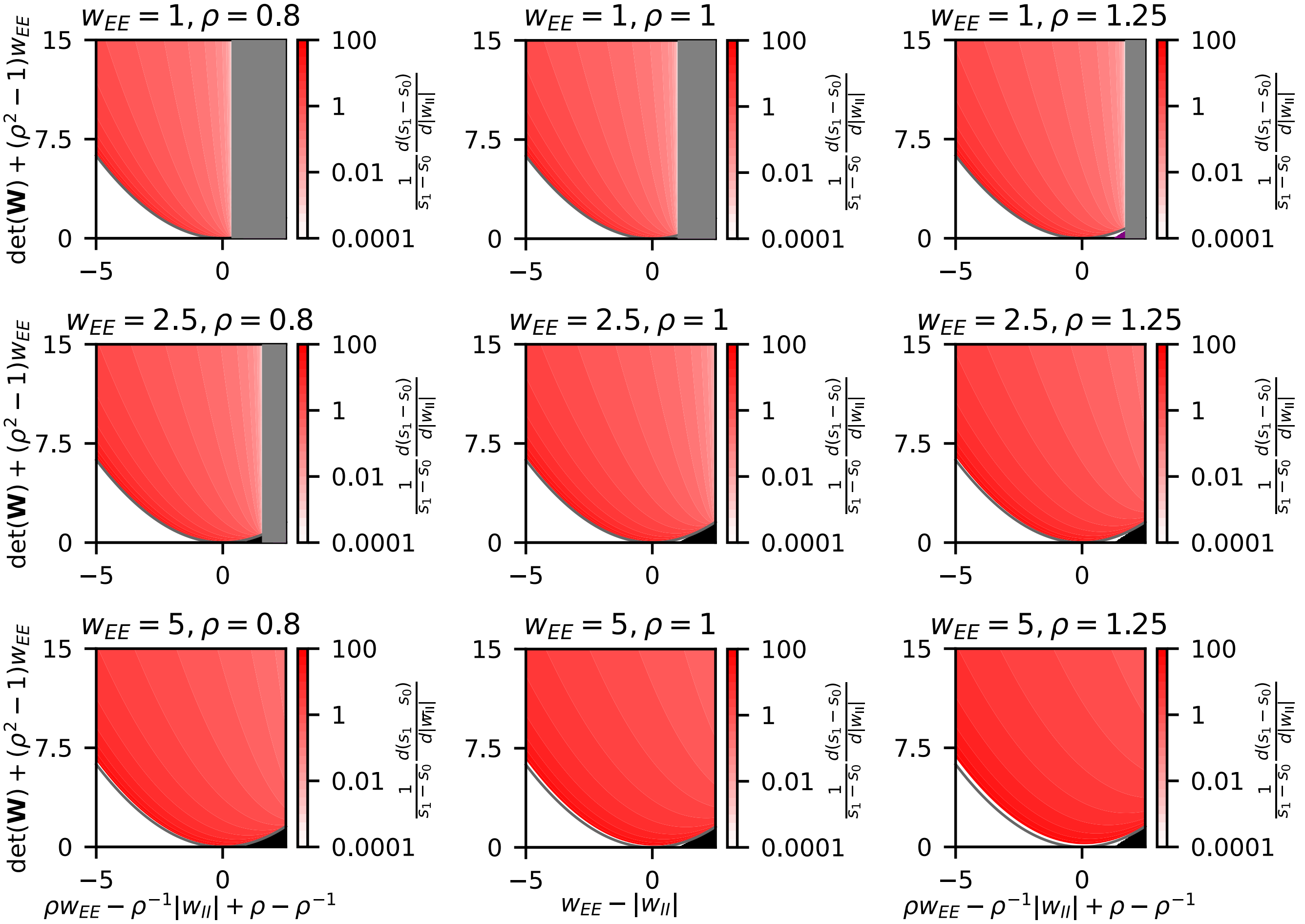


**Fig. S3.** Similar to Figure S1, but the derivative of the distance from the first zero-crossing $s_0$ to the second zero-crossing $s_1$ with respect to recurrent inhibition strength, normalized by this distance, is plotted.

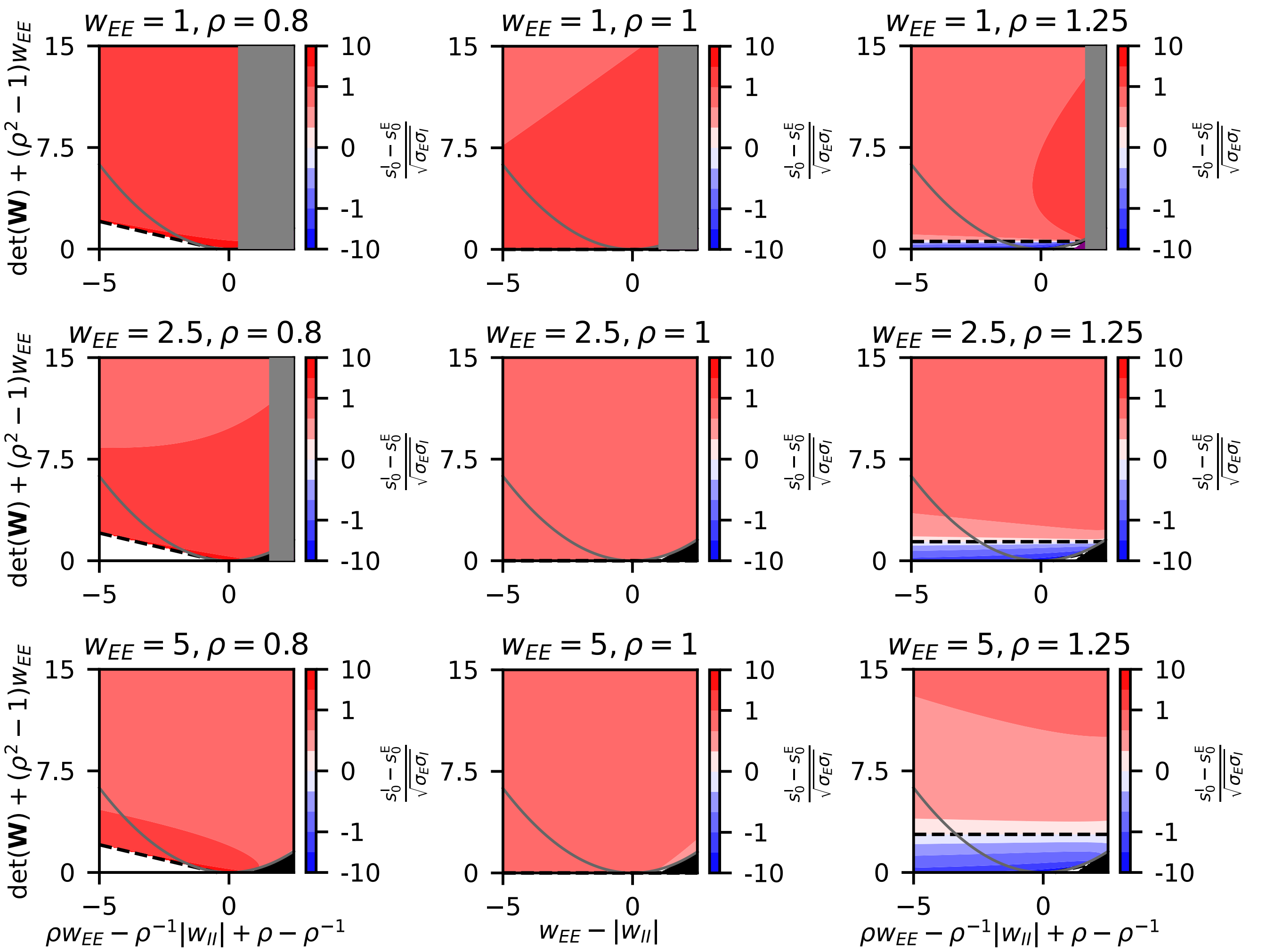


**Fig. S4.** Similar to Figure S1, but the distance between the first zero crossing of the response of excitatory neurons $r_0^{\mathrm{E}}$ and that of inhibitory neurons $r_0^{\mathrm{I}}$ as a fraction of $\sqrt{\sigma_E \sigma_I}$ is plotted. Dashed line in the left column plots is given by the equation $y = (\rho - \rho^{-1})(x - (\rho - \rho^{-1}))$. Dashed line in the right column plots is given by $\det(\boldsymbol{W}) = 0$, i.e. $y = (\rho^2 - 1)w_{\mathrm{EE}}$

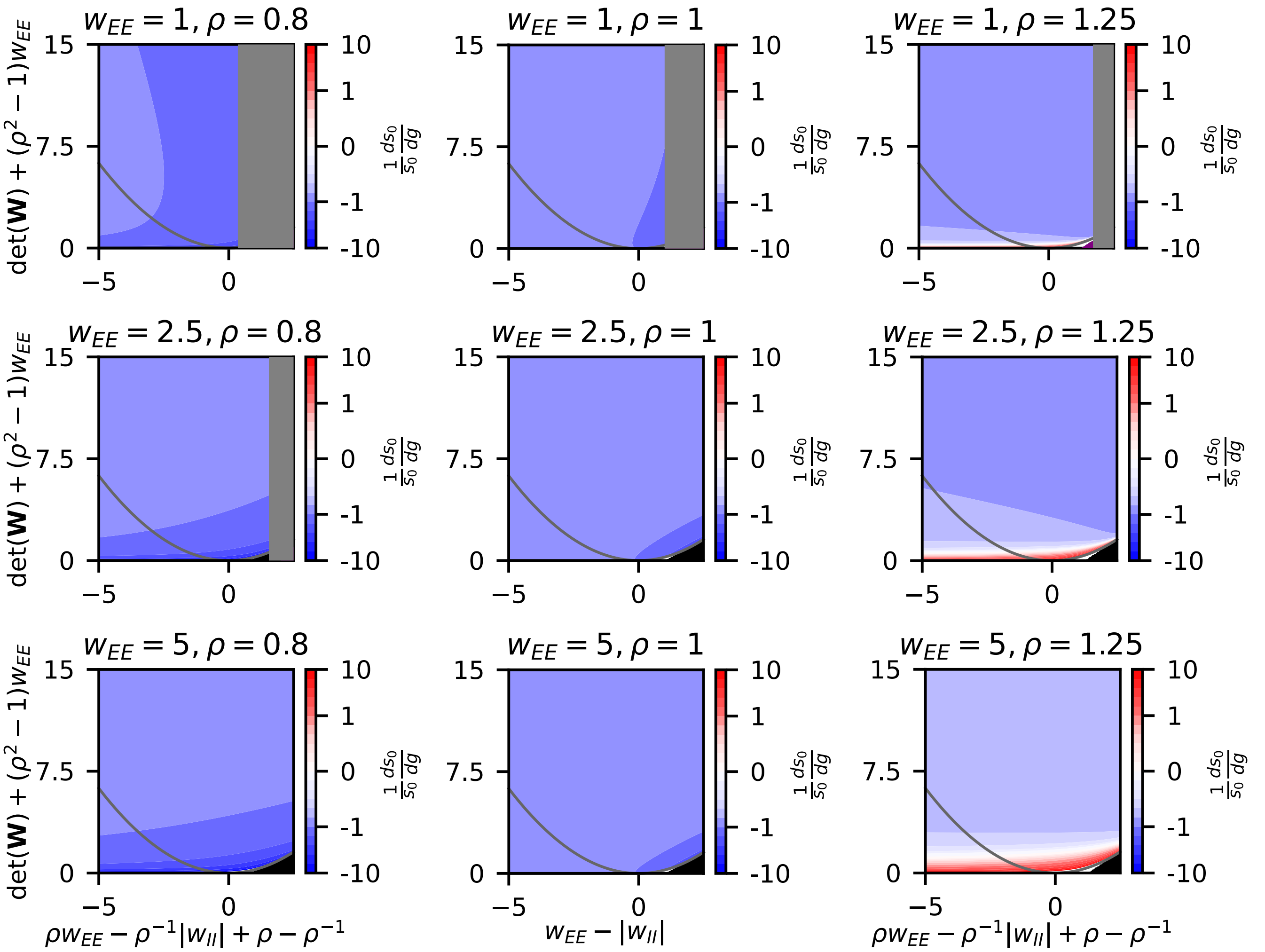


**Fig. S5.** Derivative of distance to the first zero crossing of the response of excitatory neurons to single-cell excitatory neuron perturbation with respect to overall gain, divided by the distance, for different combinations of $w_{\mathrm{EE}}$ and $\rho = \frac{\sigma_I}{\sigma_E}$

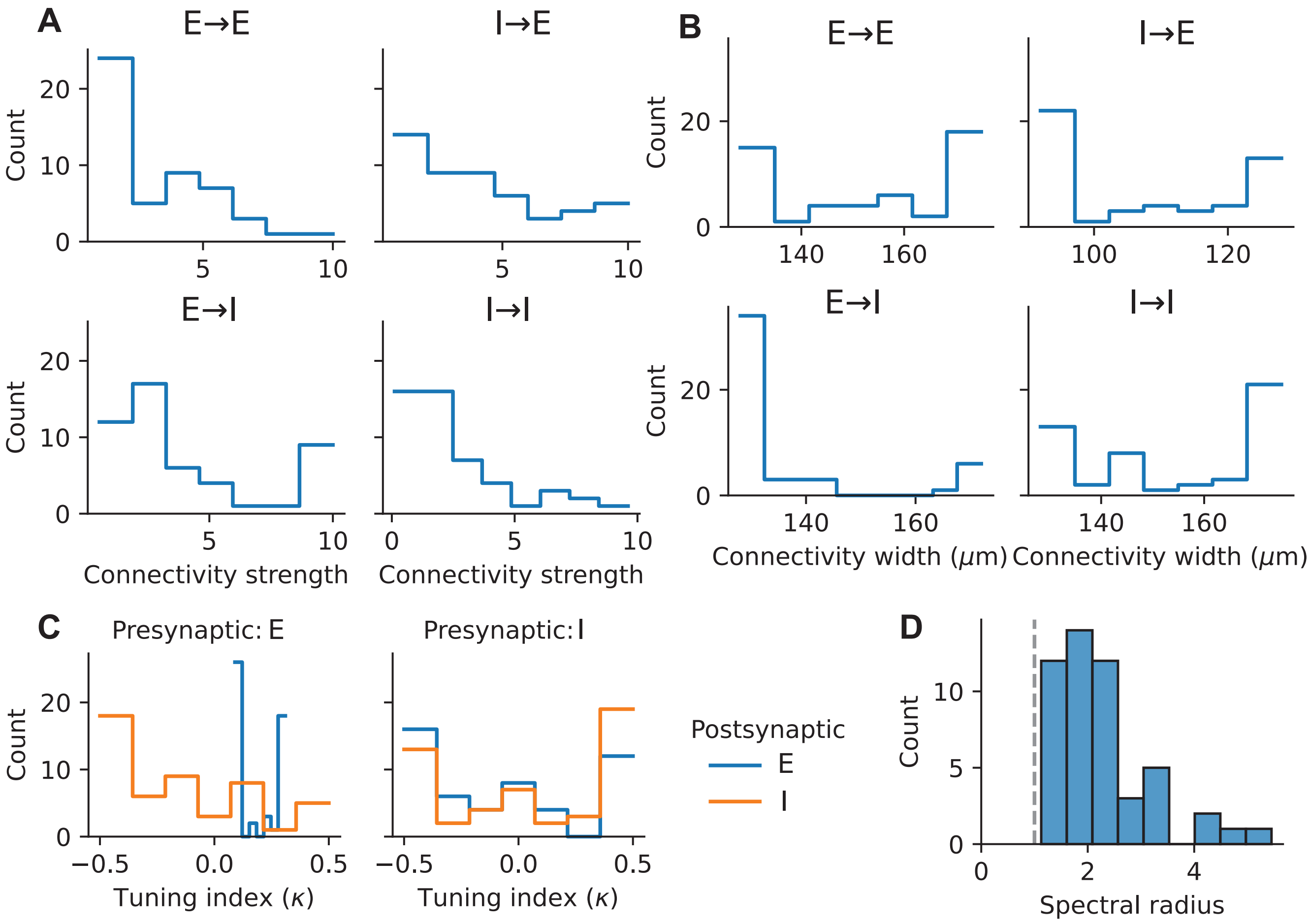


**Fig. S6. Distributon of fitted model parameters. A)** Histogram of the connectivity strength parameter $w_{\alpha\beta}$ for all combinations of cell types $\alpha, \beta$ in the top 50 fitted models. **B)** Similar to A, but for the connectivity width parameter $\sigma_{\alpha\beta}$. **C)** Similar to A, but for the feature tuning parameter $\kappa_{\alpha\beta}$. **D)** Histogram of the spectral radius of the full connectivity matrix of the top 50 fitted models. Vertical dashed line represents spectral radius of $1$.

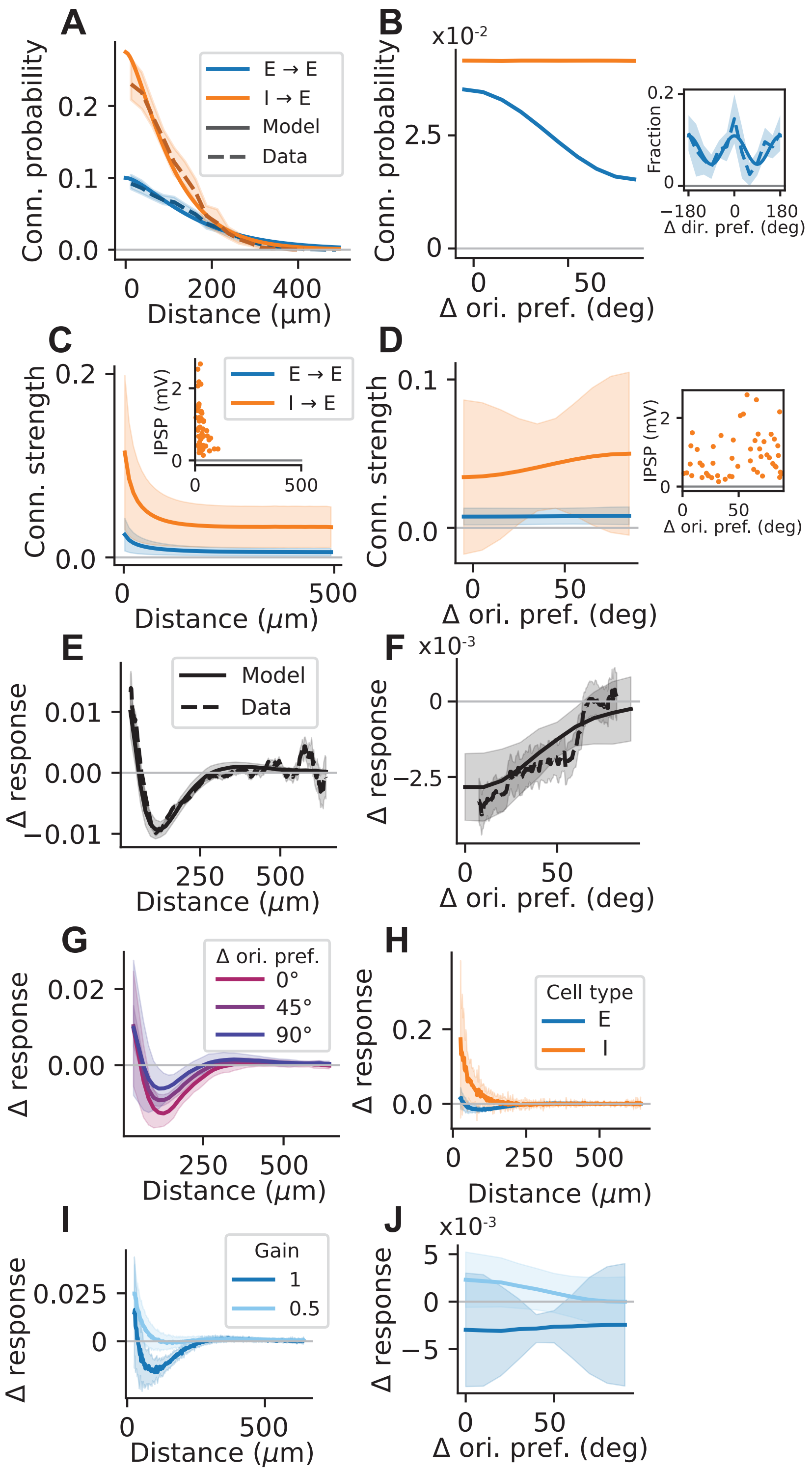


**Fig. S7. Perturbation responses of models with sparse connectivity. A)** Connection probability for E $\to$ E and I $\to$ E as a function of distance in the sparsely connected models (solid lines), compared to scaled data from (4) (dashed lines). Model connection probability is parametrized as $w_{\alpha\beta}\sigma_{\alpha\beta}^{-2}G_0(s;\sigma_{\alpha\beta}^{-2})$, where $G_0$ is the Green's function kernel (Eq. S10) with $d = 0$. As (4) reports the relative but not the absolute connection probabilities, each data curve is scaled such that connection probability at $0\,\mu\mathrm{m}$ agrees with data reported by (6) (Materials and Methods). Shaded area represents 95% confidence interval of the data computed by bootstrapping ($n = 17$). **B)** Connection probability for E $\to$ E and I $\to$ E as a function of difference in preferred orientation in the sparsely connected models. Inset: Comparison of E $\to$ E connection density as a function of preferred direction in the sparse model (solid line) to data from (4) (dashed line). Note that the y-axis of the inset represents the fraction of presynaptic excitatory neurons connecting to an excitatory neuron since (4) reports the relative but not the absolute connection probabilities. Shaded area represents standard error. **C)** Connection strength for E $\to$ E and I $\to$ E as a function of distance in the sparsely connected models. Shaded area represents standard deviation across models. Inset: IPSP of I $\to$ E connections as a function of distance reported by (12), reproduced here for comparison. Notice that in both the model and the data, connection strength decays as a function of distance at short length scales ($< 100\,\mu\mathrm{m}$). **D)** Similar to C), but as a function of difference in preferred orientation. Inset: IPSP of I $\to$ E connections as a function of difference in preferred orientation reported by (12). Notice that in both the model and the data, the connection strength between similarly-tuned neurons is slightly weaker than the connection strength between oppositely-tuned neurons, though this trend is not statistically significant in the data. **E-J)** Same plots as main text Figure 6C-H, but the response of the sparsely connected model rather than the fully connected model is shown.

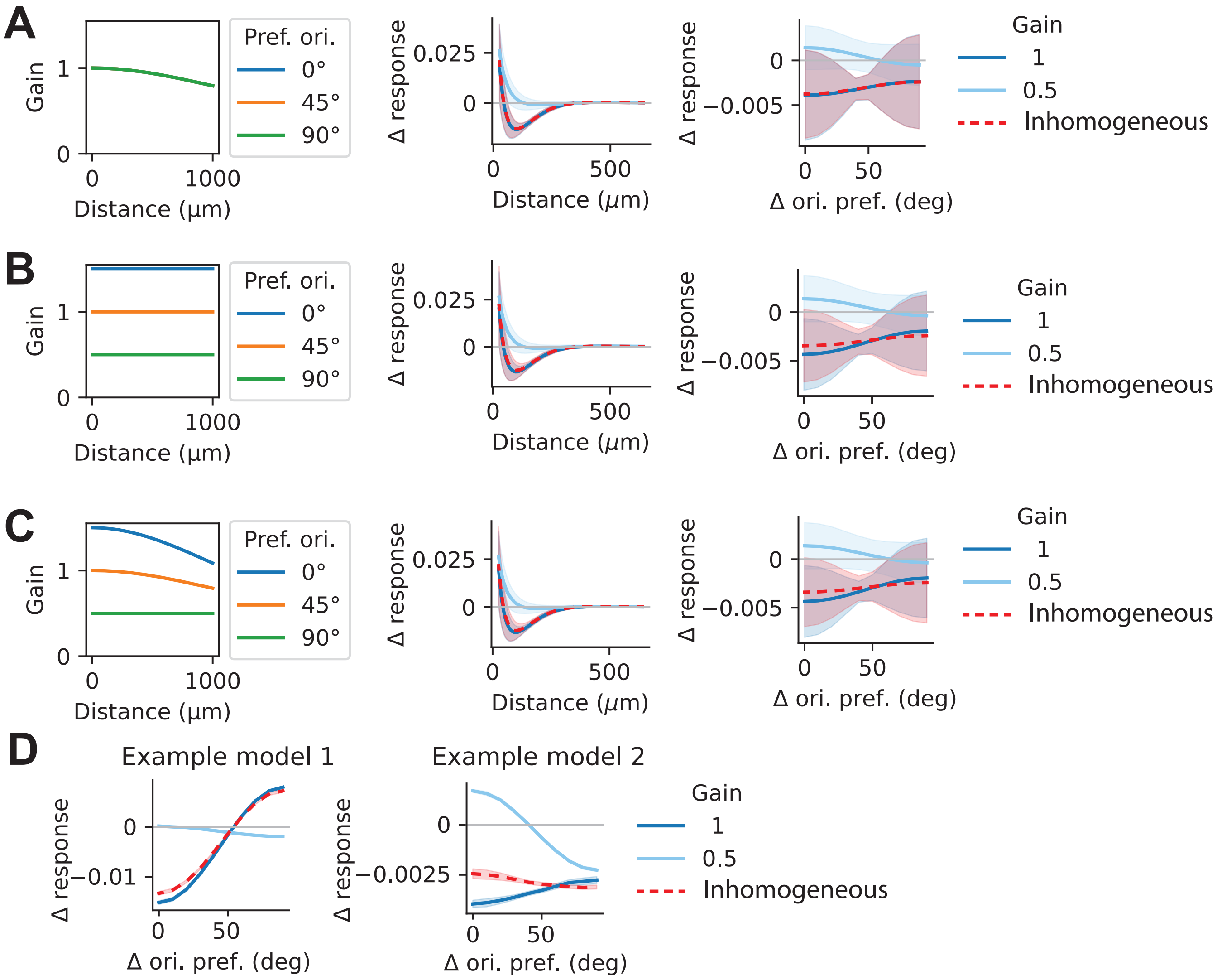


**Fig. S8. Perturbation responses of models with inhomogeneous gain. A)** Left: Neuronal gain as a function of spatial distance from the center of the simulation region and orientation tuning preference. The curves for $0, 45$, and $90$ degrees are overlapping. Mathematically, the gain is given by $g_0 + (1 - g_0)\exp\left(-\frac{r^2}{2\sigma_g^2}\right)(1 + 2\kappa_g \cos(2\theta))$, where $g_0 = 0.5, \sigma_g = 968\,\mu\mathrm{m}, \kappa_g = 0$. Center and right: Mean perturbation responses of excitatory neurons in the top $50$ best-fit models when the gain is equal to $1$ uniformly (dark blue), when the gain is equal to $0.5$ uniformly (light blue), and when the gain is inhomogeneous and is described by the left panel (dashed red line), with shaded region indicating standard deviation across models. Note that the dark blue and light blue curves are the same as those in Figures 6G and 6H in the main text. **B)** Similar to A, but neuronal gain is dependent on orientation tuning but not spatial distance, with the gain parameters given by $g_0 = 0.5, \sigma_g = \infty, \kappa_g = 0.5$. $2$ of the $50$ models are dynamically unstable with this choice of gain, and thus the perturbation responses of these $2$ models are excluded from this subplot. **C)** Similar to B, but neuronal gain is dependent both spatial distance and orientation tuning, with the gain parameters given by $g_0 = 0.5, \sigma_g = \sigma_g = 968\,\mu\mathrm{m}, \kappa_g = 0.5$. Again, $2$ of the $50$ models are dynamically unstable with this choice of gain. **D)** Perturbation responses of excitatory neurons in two example fitted models with the same space- and feature-dependent neuronal gain as in C. Notice that the inhomogeneity in neuronal gain has little effect on the perturbation response of model $1$, while the effect on model $2$ is so strong that the excitatory neuron response changes from opposite-favoring to same-favoring.
